# Extreme spatial plasticity in Italian wolves: ecological determinants of range size and movement patterns across heterogeneous landscapes

**DOI:** 10.64898/2026.06.12.731796

**Authors:** Silvia Cavazza, Rudy Brogi, Michele Zanni, Celeste Buelli, Duccio Berzi, Siriano Luccarini, Liliana Costanzi, Cristian Tirapelle, Nadia Cappai, Paolo Bongi, Marco Del Frate, Enrico Vettorazzo, Marco Apollonio

## Abstract

As wolves (*Canis lupus*) recolonize diverse environments, the extent to which ecological and individual factors shape their spatial behaviour remains insufficiently understood. Italy, where wolf recovery began earlier than elsewhere in Europe and spans highly heterogeneous landscapes, offers an opportunity to explore patterns relevant to future European scenarios.

We analysed variation in home range size and movement patterns using GPS data from 19 wolves monitored between 2019 and 2025 across five study areas representing a broad environmental gradient. Continuous-time movement modelling was used to estimate home range size and distance travelled.

Home range size varied markedly among areas, with annual averages for pack-member wolves ranging from 35 to 185 km^2^. The smallest ranges occurred where wolves had been established for longer and/or where prey availability was higher, suggesting that both intraspecific competition and resource abundance influence space use. Wolves expanded their ranges in winter and in more urbanized landscapes. Pack members consistently had smaller home ranges than floaters, and among floaters, females showed slightly larger ranges than males. Distance travelled also differed significantly across areas (local averages 25–53 km/day) but showed no significant association with individual traits. Three dispersal events were documented, two of which resulted in successful pair formation and subsequent reproduction.

Overall, our results provide a broad overview of wolf spatial behaviour in contemporary Italian landscapes and offer insights relevant to the management of wolf populations across Europe.

## 1. Introduction

The recent recolonization of Europe by wolves (*Canis lupus*), after decades of near-total absence from large portions of its former distribution, was a phenomenon primarily driven by reduced human persecution and the abandonment of mountainous and hilly areas, which has caused an increase in rewilded areas and in populations of wild ungulates (Apollonio et al. 2010; Pereira and Navarro 2015) and protective policies devoted to large carnivores (Boitani 2000). In Italy, as in the rest of Europe, the wolf population declined over the past two centuries (Breitenmoser 1998), with a population remaining confined to the Apennines (Cagnolaro et al. 1974). However, since the 1970s, this negative trend reversed, increasing the Apennine population from some hundreds individuals to the current estimate of 2,388 (95% Confidence Interval (CI): 2,020-2,645; La Morgia et al. 2022) which may still underestimate the actual population size (Merli et al. 2023). This rapid population growth led to wolves gradually saturating large portions of peninsular Italy (e.g., Tuscany; Zanni et al. 2023; Brogi et al. 2025a). Their expansion along the Apennine chain made Italian wolves reach the Western Alps in the last decade of the previous century (Scandura et al. 2001; Fabbri et al. 2007). The Alps recolonization gradually proceeded eastward until reaching the Venetian Alps, where a reproductive pair composed of an Italian female and a Dinaric male was first documented in 2012 (Ražen et al. 2016). Currently, the Alpine wolf population is estimated at 946 individuals (95% CI: 822-1,099; La Morgia et al. 2022).

The wolf, as a highly adaptable species, exhibits spatial behaviour ranging from the occupation of stable home ranges to variable dispersal patterns, in accordance with to the species’ social structure and ecology (Mech and Boitani 2003). Wolves indeed display complex social behaviour, characterized mainly by the formation of territorial packs, but also involving solitary, non-territorial displacements (Fuller et al. 2003; Mech and Boitani 2003). Within the broader category of solitary individuals, two distinct movement strategies can be identified: dispersal and the floater phase. Dispersal refers to the directed, one-way movement of an individual away from its natal or previous territory, typically covering long distances to reach new, unoccupied areas where reproduction may eventually occur (Fritts and Mech 1981; Caniglia et al. 2014; Fabbri et al. 2014; Sanz-Pérez et al. 2018). Empirical evidence indicates that sex-related differences during dispersal are not consistent across studies, with several authors reporting no significant differences between males and females in either dispersal rates or dispersal distances (Peterson et al. 1984; Fuller et al. 1989; Gese and Mech 1991; Ballard et al. 1997; Blanco and Cortés 2007; Jarausch et al. 2021). However, when sex differences are observed, they generally suggest that males tend to disperse more frequently and over longer distances, particularly during long-distance dispersal events (Caniglia et al. 2014; Fabbri et al. 2014; Jiménez et al. 2017; Sanz-Pérez et al. 2018), a pattern consistent with stronger female philopatry reported in wolves and other mammalian species (Greenwood 1980; Pacheco et al. 2024). Genetic evidence supporting sex-biased dispersal further indicates that males are more likely to reproduce in non-natal packs or to establish new packs, whereas females more frequently reproduce within their natal group (Jędrzejewski et al. 2005; Caniglia et al. 2014; Morales-González et al. 2022; Pacheco et al. 2024). In contrast, floaters are non-territorial wolves that, after leaving their natal group, temporarily occupy and move within areas overlapping several pack territories without establishing a stable range. This phase is often characterized by repeated exploratory movements and shorter-term space use while seeking vacant territories or mating opportunities (Mech and Boitani 2003). When analyzing spatial ecology, it is critical to distinguish between these two strategies since floater behaviour entails extended, irregular usage of space across varied time periods, whereas dispersal movements are usually quick and linear.

These differences have important implications for understanding population connectivity, colonization dynamics, and the spatial variability of home range estimates in expanding wolf populations.

Home range sizes of wolves in Europe vary widely depending on geographic and ecological context. In southern and central regions, estimates range from 80 to 460 km^2^ (Okarma et al. 1998; Findo and Chovancova 2004; Kusak et al. 2005; Jędrzejewski et al. 2007; Karamanlidis et al. 2016; Mysłajek et al. 2018), while in Scandinavia, ranges are considerably larger, spanning from 259 to 1,676 km^2^ (Mattisson et al. 2013). In Italy, home range sizes estimated using the Minimum Convex Polygon (MCP, Mohr 1947) method were found to range between 120 and 200 km^2^ in the Apennines (Ciucci et al. 1997; Mancinelli et al. 2018), and around 150 km^2^ in the eastern Alps (Ražen et al. 2016). However, estimates based on alternative methodologies provide indirect cues of smaller home ranges in the Apennines, where wolves have a long history of presence and prey availability is relatively high. For instance, Mattioli et al. (2018), using spatially explicit capture-recapture (SCR) models applied to camera trap data estimated a density of 1.21 packs per 100 km^2^, implying an average territory size of approximately 82.6 km^2^ per pack.

Population density indeed plays an important role in the spatial organization of wolves, in fact, although there is limited information available, it has been observed that as wolf populations increase in colonized or recolonized areas, home range sizes tend to decrease (Fritts and Mech 1981; Mech and Boitani 2003; Mysłajek et al. 2018). This process had also been observed in another canid species as foxes (*Vulpes vulpes*) by Bateman and Fleming (2012). Besides recolonization progress and the resulting intra-specific competition for space, ungulate density may also play a prominent role in influencing wolf spatial behaviour. Wolf home ranges and territories are indeed generally smaller in environments with higher abundance of prey (Mech and Boitani 2003; Jedrzejewski et al. 2007; Mattisson et al. 2013), following the principle of the “rubber disk” firstly proposed by Huxley (1934), as the abundance of prey allows smaller areas to fulfil a pack’s food needs.

Individual factors may further influence wolf home range size, with pack-member wolves usually occupying a more confined and stable area, which could vary throughout the year in relation to their social status, compared to solitary individuals exhibiting more nomadic and unstable behaviour (Fuller et al. 2003; Roffler and Gregorovich 2018).

Among individual traits, sex has been shown to play a context-dependent role in shaping wolf spatial behaviour. Sex differences among pack members can be expected to be absent or negligible under comparable social and reproductive conditions, as male and females share the same territory and have similar requirements within the pack (Mech and Boitani 2003; Fuller et al. 2003).

In addition, human activities could influence wolf spatial behaviour, triggering variations in home range size (Paquet et al. 1996; Rich et al. 2012; Mattisson et al. 2013; Mancinelli et al. 2018). Higher human disturbance induces an overall reduction in the availability of wild prey, fragmenting suitable areas for wolves and compelling them to enlarge their home ranges (Rich et al. 2012; Mancinelli et al. 2018). Conversely, humans may provide anthropogenic food resources, thereby reducing the home range size required to meet wolves’ ecological needs (Paquet et al. 1996; Mattisson et al. 2013).

Besides home ranges, the distance travelled may also contribute to giving a comprehensive understanding of the spatial behaviour of large mammals (Cavazza et al. 2023). In wolves it received relatively less attention as compared with home range size, particularly concerning its variability during the progression of recolonization or across areas offering different prey availabilities. In North America, distance travelled ranged from 1.6 - 9.0 km/day up to 80 km/day (Kolenosky and Johnston 1967; Mech et al. 1995). In Europe, values ranged from 2.5 to 4.4 km/day (Kusak et al. 2005; Jędrzejewski et al. 2007), with 3.3 km/day estimated in central Italy (Ciucci et al. 1997), but estimations based on high-resolution spatial data and accurate analytical approaches are still missing. The distance travelled seems to be primarily influenced by the structure of the environment. In areas with open habitats or rich in anthropogenic linear features such as roads and trails, wolves travel faster than they do in forested habitats (Musiani et al. 1998) and can thus be expected to cover longer distances within the same time interval. Moreover, analogously to the expected resource-dependent variations of home range size, a low availability of food may force wolves to move over longer distances (Jedrzejewski et al. 2001). Regarding the effect of increased wolf abundance on the average distance travelled, two opposite patterns may be expected. On the one hand, wolves might travel shorter distances because smaller home ranges require less movement compared to larger ones, on the other, increased competition for space might cause wolves to patrol territory boundaries more frequently to prevent intrusions from conspecifics, as observed by Zub et al. (2003) in the Bialowieza Primeval Forest.

The aim of this study is to provide a comprehensive description of wolf spatial behaviour in terms of range size, distance travelled, and dispersal or floater dynamics, and to identify key ecological and individual-level drivers underlying variations in range size and distance travelled, considering both individual and pack-level dynamics. From 2019 to 2025, we equipped 19 wolves with GPS collars across five areas of Italy, selected to represent a range of environmental conditions, including differences in prey availability, recolonization history, and landscape anthropization. We investigated the influence of ecological and individual factors on both monthly range and monthly distance travelled. By modelling the spatial behaviour of wolves in relation to these multiple factors, this analysis contributes to a deeper understanding of the ecological flexibility of the species and provides useful information for wolf conservation and management scenarios in European landscapes.

## 2. Material and Methods

### 2.1. Study areas

Spatial data on wolves were collected in five distinct areas of Italy (Fig. 1): the Foreste Casentinesi National Park (FCNP), the north-eastern Italian pre-Alps and Alps, the San Rossore Estate (SRE), a protected area within Regional Park Migliarino-San Rossore-Massaciuccoli, and the Presidential Estate of Castelporziano (PEC). These areas provided a diverse set of environmental conditions to study wolf spatial behaviour, particularly in terms of time elapsed from the wolf return and prey availability.

**Fig. 1.**
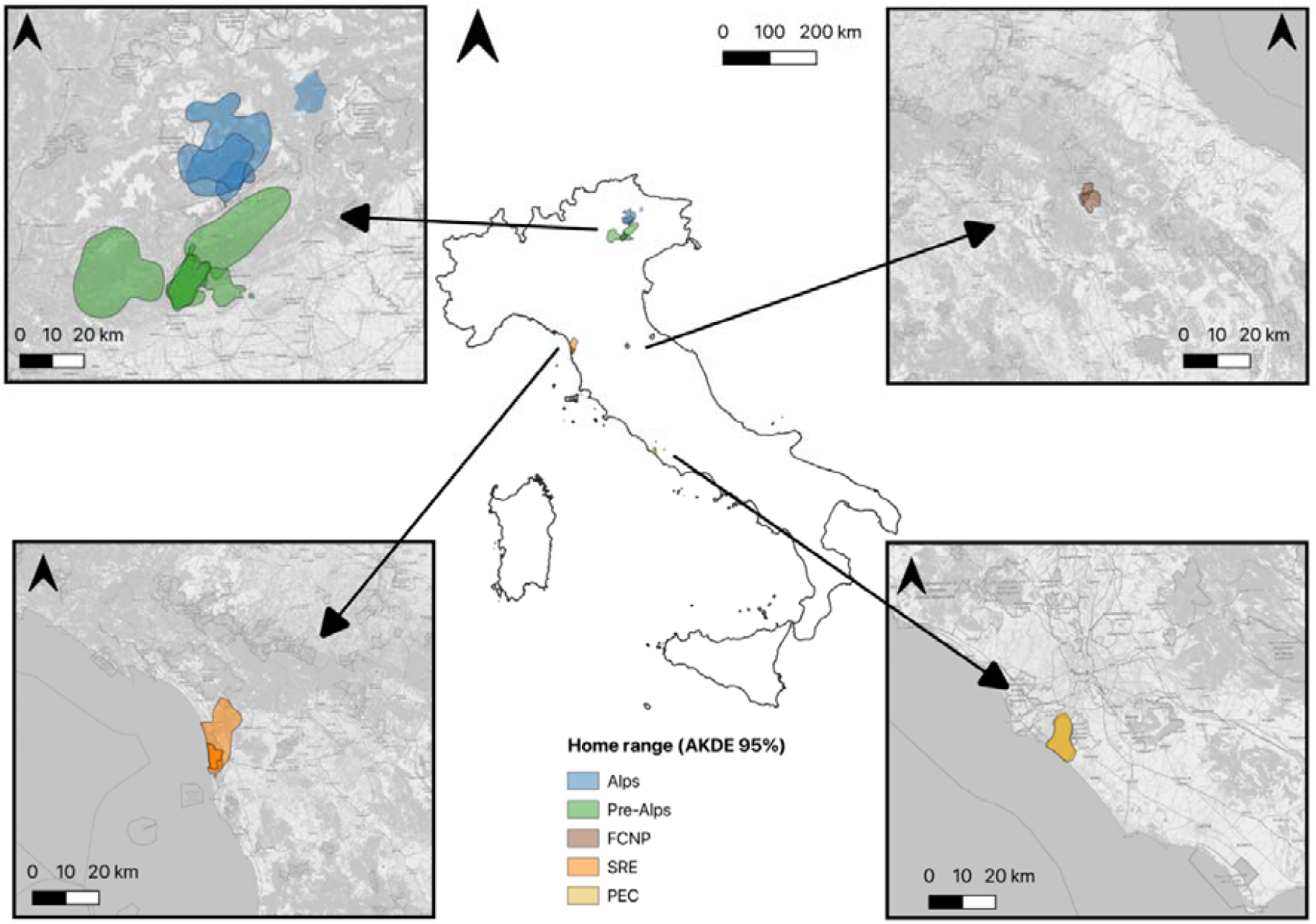
Distribution in Italy of the individual total home ranges of GPS-collared wolves in Alps (blue), pre-Alps (green), Foreste Casentinesi National Park (FCNP, brown), San Rossore Estate (SRE, orange), and Presidential Estate of Castelporziano (PEC, yellow). Alps and pre-Alps were treated as separated study areas on account on their different biotic and abiotic conditions (please see the Study area section for more details)

FCNP is located between the provinces of Arezzo, Firenze and Forlì-Cesena and covers approximately 36,000 ha. Elevation ranges from 330 to 1,658 m a.s.l., with a cool continental temperate climate (Fratianni and Acquaotta 2017). Within the home ranges of the monitored wolves, forests account for 65% of the land cover and are dominated by deciduous species, with a smaller proportion of conifers. Shrubs represent 2%, agricultural areas 30%, and anthropized areas about 3%. The area hosts several ungulate species, including wild boar (*Sus scrofa*), roe deer (*Capreolus capreolus*), red deer (*Cervus elaphus*), and fallow deer (*Dama dama*), amounting to approximately 1,449 kg/km^2^ of available prey biomass (Table S1, see the section 2.2 for more details). The presence of wolves in FCNP is historical, dating back to the early 1990s, with the first confirmed reproduction in 1992 (Apollonio et al., 2004) and possibly even from 1985 (Boscagli 1985).

The North-eastern Italian pre-Alps include the Grappa Massif (straddling the provinces of Vicenza, Belluno, and Treviso) and the Asiago Plateau (province of Vicenza), with elevations ranging from 90 to 2,300 m a.s.l. and a cold-temperate climate. Habitat composition within the wolves’ home ranges consist of approximately 62% forest cover, dominated by deciduous rather than coniferous species. Shrubs, natural open areas, and agricultural areas occupy 4%, 22%, and 5% respectively, while anthropized areas cover around 7% of the area. The ungulate community includes roe deer, red deer, northern chamois (*Rupicapra rupicapra*), and mouflon (*Ovis aries*), with an estimated total prey biomass of approximately 962 kg/km^2^ available in this area (Table S1).

The north-eastern Italian Alps include the Dolomiti Bellunesi (province of Belluno), reaching up to 3,343 m a.s.l. The Dolomiti Bellunesi National Park (31,000 ha) lies within this sector. The climate transitions from cold-temperate at lower elevation to a nival climate above 2,800 meters (Fratianni and Acquaotta 2017). Within the monitored wolves’ ranges, forests cover about 63% of the area, dominated by coniferous rather than deciduous species. Shrublands, natural open areas, and agricultural lands account for approximately 11%, 21%, and 2% of the landscape, respectively, while human-modified areas represent around 2%. The area hosts a rich ungulate community, including roe deer, red deer, northern chamois and mouflon, providing an estimated 1,275 kg/km^2^ of available prey biomass (Table S1).

Both mountainous ranges experience significant human presence due to tourism and livestock grazing, especially during the summer. Wolves recently recolonized these mountainous regions, with the first sightings in 2015 and first confirmed reproduction on the Asiago Plateau in 2016, and on the Grappa Massif in 2017 (Avanzinelli et al. 2017, 2018).

The coastal areas of SRE (province of Pisa) and PEC (province of Rome) are flat protected areas along the Tyrrhenian coast, both characterized by a sub-Mediterranean climate. SRE, which is only partially fenced on its eastern side and bounded by two major river systems to the north and south and by the sea to the west, covers approximately 5,000 ha, while PEC spans about 6,000 ha.

In SRE, forests cover 42% of the area occupied by the wolves’ home ranges and are predominantly composed of deciduous species. Shrubs occupy 4%, natural open areas are around 2%, agricultural areas are 36%, and urbanized areas approximately 16%. The park hosts wild boar with a limited density and fallow deer with a very high density, for a total of 2,167 kg of prey biomass per km^2^ being available to wolves (Table S1).

In PEC, forests cover 73%, followed by 11% of shrubslands, 10% of agricultural areas, 5% of natural open areas, and 1% of urbanized areas. The estate hosts wild boar with a high density, fallow deer, roe deer and red deer with a very limited density, for a total of 1,605 kg of prey biomass per km^2^ (Table S1, Franzetti, 2024).

Both areas host the youngest wolf populations, with the first individuals recorded in 2016. The first wolf reproduction was documented in 2021 in SRE (Del Frate et al. 2023), and in 2022 in PEC (Luccarini et al. unpublished data).

### 2.2. Prey biomass availability estimation

We calculated the available prey biomass, separately for each study area, by multiplying the local densities of ungulate species by the average individual body mass, across sex and age classes, of each species.

For ungulate density estimates in the FCNP, we used data from the most recent Foreste Casentinesi National Park annual report for red deer (Foreste Casentinesi National Park, 2023), results of spring 2022 drive censuses for roe deer and fallow deer (L. Mattioli, pers. comm.), and Random Encounter Model estimates for wild boar derived from a nearby sector with comparable environmental conditions (Guerrasio et al. 2022).

For the Alps and Pre-Alps, density estimates for all four ungulate species were obtained from vantage-point counts conducted by hunters and local game managers (official data from the Provincial Administrations of Treviso, Belluno, and Vicenza, kindly provided by the provincial Wildlife Service offices). These data were complemented with 23 systematic block-count surveys for northern chamois and red deer, and spotlight counts for roe deer, carried out within the Dolomiti Bellunesi National Park (E. Vettorazzo, pers. comm.). Given the large spatial extent (Fig. 1) and environmental heterogeneity of the Alpine and Pre-Alpine areas, we estimated prey biomass separately for each wolf pack. Pack-specific estimates were calculated by averaging ungulate densities within the Autocorrelated Kernel Density Estimation (AKDE) total home ranges of packs (see Section 2.4).

For wild boar and fallow deer densities in the SRE study area, we used the mean values from the 2023 annual vantage-point censuses reported in the Management Plan of the Migliarino–San Rossore– Massaciuccoli Regional Park. For the PEC study area, ungulate density estimates were derived from autumn 2024 distance-sampling surveys combined with infrared thermal-camera observations (ISPRA, 2024).

Average body masses for each species were taken from the European ungulate values reported by Apollonio et al. (2010). For wild boar, however, we used an average mass of 30 kg instead of the 50 kg reported for European populations, to reflect the markedly smaller size of Italian populations (Brogi et al. 2021, Table S1).

### 2.3. Data collection

From 2019 to 2025, we GPS-tracked 19 wolves (nine females and ten males) belonging to 13 different packs (Table 1). We captured all of them by means of foothold traps (Fremont™ Humane Foot Snare Wolf/Cougar 1/8 7×7 and Fremont™ Humane Foot Snare Fox/Coyote 3/32 7×7). Following capture, wolves were chemically immobilised using an anaesthetic protocol based on a ketamine-medetomidine combinations (Holz et al. 1994; Arnemo et al. 2013), administered via syringe blowpipe (Telinject). This procedure allowed the collection of biometric measurements and biological samples, age estimation, and the fitting of GPS collars (VERTEX Plus, Vectronic Aerospace GmbH). All collars were set with a variable fix rate, acquiring 1 GPS position/30 minutes during the night, when wolves are more active, and 1 GPS position/2 hours during the day, when wolf activity is usually reduced (Blaskovic et al. 2022; Ferretti et al. 2023). During the first months following deployment, collars were temporarily programmed with a constant fix rate throughout the day (locations recorded every 1 hour or every 30 minutes) to closely monitor animals immediately post-release. On a monthly level, we classified each wolf’s individual status as pack member, floater or dispersal. We classified as pack-member those wolves belonging to a pack or reproductive pair. Solitary individuals were considered floaters or dispersers: floaters ranged within a defined area, whereas dispersers moved away from their natal range along a predominantly directional path. Both categories could eventually join an existing pack or establish a new one, thus attaining pack-member status. Individual social status, pair or pack formation, and reproductive events (including the presence of pups) were determined through the combined use of telemetry data, camera trapping (6–32 cameras per area) and field inspections. Camera traps were placed in locations suitable for detecting wolf movements and associations across each study area. To support the interpretation of GPS and camera-trap data, field inspections of GPS location clusters were conducted. Clusters were defined as two or more consecutive locations within a 200-m radius and included both diurnal and nocturnal locations, which were visited daily. At each cluster site, the surrounding area was systematically searched for field signs, including prey remains from predation events, scats, resting sites, and other indicators of wolf presence and social activity. Pack-related activity was inferred from behavioural or physical indicators of coordinated action among multiple individuals, such as group hunting signs (e.g., multiple feeding sites around a single carcass), overlapping tracks, or communal resting areas (e.g., multiple resting sites containing wolf hair). By integrating these methods, we could determine the time of any individual status change and assign the appropriate status for each month, which could vary over time. Some wolves transitioned from being floaters or dispersing to joining a pack or forming a pair and vice versa (Table 1).

**Table 1.** Individual data of captured wolves during their monitoring through GPS-tracking.

| ID | Area | Sex | Age | Status | Pack | Start | End | Tracking days |
| --- | --- | --- | --- | --- | --- | --- | --- | --- |
| WF1901 | Pre-Alps | f | >2 yrs | P | Grappa1 | 06-09-2019 | 21-12-2019 | 137 |
| WF2001 | Pre-Alps | f | >2 yrs | P | Grappa1 | 08-08-2020<br>*13-09-2023 | 06-06-2021<br>26-02-2024 | 304<br>166 |
| WF2002 | Pre-Alps | f | 18 mo | F/P | Asiago | 23-11-2020 | 29-06-2022 | 583 |
| WM2101 | Pre-Alps/Alps | m | >2 yrs | P/D/P | Grappa1 / Dubiea | 29-03-2021 | 12-08-2022 | 501 |
| WF2101 | Pre-Alps | f | 17 mo | P/D/P | Grappa1 / Grappa2 | 01-10-2021 | 07-01-2023 | 463 |
| WF2202 | Alps | f | >2 yrs | P | DBNP1 | 13-09-2022 | 08-09-2024 | 726 |
| WM2201 | FCNP | m | >2 yrs | P | FCNP1 | 20-10-2022 | 09-12-2023 | 415 |
| WF2203 | FCNP | f | >2 yrs | P | FCNP2 | 21-11-2022 | 25-11-2023 | 369 |
| WM2301 | Pre-Alps | m | 20 mo | **P/F | Grappa3 | 22-01-2023 | 31-10-2023 | 298 |
| WM2303 | SRE | m | >2 yrs | P | SRE1 | 15-04-2023 | 15-11-2023 | 214 |
| WM2304 | SRE | m | >2 yrs | F/D/P | SRE2 | 17-04-2023 | 20-09-2024 | 522 |
| WM2305 | FCNP | m | >2 yrs | P | FCNP3 | 15-10-2023 | 20-02-2025 | 494 |
| WF2401 | SRE | f | 8 mo | P | SRE1 | 22-01-2024 | 30-04-2025 | 464 |
| WM2401 | SRE | m | 20 mo | P | SRE1 | 22-01-2024 | 30-04-2025 | 464 |
| WM2402 | PEC | m | >2 yrs | P/D | PEC | 13-02-2024 | 07-12-2024 | 298 |
| WF2402 | Alps | f | 16 mo | P/F | DBNP1 | 04-09-2024 | 30-04-2025 | 238 |
| WF2404 | Alps | f | >2 yrs | P | DBNP2 | 25-10-2024 | 30-04-2025 | 187 |
| WM2403 | PEC | m | >2 yrs | P | PEC | 21-11-2024 | 30-04-2025 | 160 |
| WM2404 | PEC | m | 19 mo | P | PEC | 20-12-2024 | 30-04-2025 | 131 |

FCNP: Foreste Casentinesi National Park; DBNP: Dolomiti Bellunesi National Park; SRE: San Rossore Estate; PEC: Presidential Estate of Castelporziano; f: female; m: male; yrs: estimated years of age at the time of capture; mo: estimated months of age at the time of capture; P: pack-member individual; F: floater individual; D: dispersal individual. Start: start date of the monitoring; End: end date of the monitoring. \*Date of second capture event. \*\*Social status is uncertain due to ambiguous spatial behaviour

Finally, kinship and parent-offspring relationships were assessed using genetic analyses based on blood samples collected during capture and non-invasive samples (scats) collected during field monitoring.

### 2.4. Home range size, distance travelled estimation and dispersal patterns

We estimated annual, seasonal, and monthly home ranges for pack-member and floater wolves using Autocorrelated Kernel Density Estimation (AKDE; Fleming et al. 2015; Calabrese et al. 2016) implemented in the “akde()” function of the *ctmm* R package (Calabrese et al. 2016). Distances travelled at annual, seasonal, and monthly scales were calculated using the same package through the “speed()” function and the Continuous-Time Speed and Distance (CTSD) method (Noonan et al. 2019). These approaches account for autocorrelation in telemetry data and are robust to varying sampling schedules. In distance estimation specifically, incorporating autocorrelation enables the simulation of the full range of possible non-linear movement paths, yielding accurate estimates of total distance travelled (Fleming et al. 2014, 2019; Noonan et al. 2019).

Only GPS positions with a Dilution Of Precision (DOP) < 10 were retained. Annual home ranges and distances were calculated for wolves monitored for at least 10 months; for individuals with longer datasets, estimates were restricted to the first 12 months during which social status remained stable. Seasonal home ranges and distances were computed for two biologically meaningful six-month periods: summer (May–October), corresponding to parturition and pup rearing, and winter (November–April), beginning after the abandonment of the rendezvous sites (Messier 1985; Bassi et al. 2012). We, therefore, estimated seasonal home ranges and seasonal distances travelled only for individuals whose monitoring period covered the full six consecutive months of one or both seasons. Monthly home ranges and monthly distances were calculated for months with more than 15 days of data, as these metrics were later used as response variables in the linear mixed models (see Section 2.5). Months corresponding to dispersal periods were excluded because individuals did not occupy a stable home range during those intervals. To facilitate comparison with previous studies, we also computed home ranges using the 95% Minimum Convex Polygon (MCP) and distances using Straight-Line Displacement (SLD). Because these traditional methods are sensitive to sampling frequency and to autocorrelation among successive locations, spatial data were resampled at a coarser six-hour interval before calculating MCP and SLD values.

For dispersal events, we measured the total distance covered by the dispersing wolf, calculated as the linear distance between the centroid of the original home range and either the centroid of the new home range or the last recorded location, the temporal duration of the dispersal period, and its outcome (successful formation of a new reproductive unit or not).

### 2.5. Statistical analyses

At the annual scale, we examined the relationship between wolf space use and prey characteristics by testing for correlations between the annual home range size of pack-member individuals and (i) the average body mass of the main prey species and (ii) the estimated total prey biomass in each study area. Both relationships were assessed using Pearson’s correlation coefficient (cor.test function in R).

At a finer scale, we analysed monthly range size and monthly distance travelled using 229 individual-months out of 251. One month was excluded because it was best described by an Ornstein–Uhlenbeck (OU) rather than an Ornstein–Uhlenbeck Foraging (OUF) movement model, preventing the calculation of distance travelled (Calabrese et al. 2016). Eighteen additional months had insufficient sampling (<15 days), and four were discarded because the individuals (WM2101 and WM2402) were dispersing, showing directed movement rather than range use.

To investigate variation in monthly range size across the five study areas, we modelled log-transformed monthly range size using a linear mixed model (LMM; Baayen 2008), with wolf identity included as a random effect. The dataset consisted of 229 observations from 19 individuals. Fixed effects included study area (five-level factor), individual sex, social status (two-level factor), and their interaction to test for sex-specific status effects. We also included the average human footprint within the monthly range (Mu et al. 2022) as a covariate. We did not include prey biomass in the same model because prey availability was estimated at the study-area level and therefore varied together with study area, which in our dataset also captured broader environmental differences (including recolonization context), making the two effects difficult to disentangle. Seasonal variation in range size was accounted for by modelling the annual cycle using a trigonometric approach (Stolwijk et al. 1999): each observation was assigned a month (1–12), converted to radians, and its sine and cosine included as predictors. To assess whether seasonal patterns differed by social status, we added a status × month interaction. The dispersion parameter (0.53) indicated no evidence of overdispersion in the full model, although mild underdispersion was detected. Model fit was evaluated using marginal and conditional R^2^, which indicated that a substantial proportion of variance was explained (marginal R^2^ = 0.61, conditional R^2^ = 0.91). Collinearity among predictors was evaluated using Pearson correlation coefficients (Zuur et al. 2009) and none exceeded 0.6. Models were fitted with the lmer function in the lme4 package (Bates et al. 2015). To maintain the nominal type-I error rate (0.05), we incorporated random slopes of human footprint, sine(month), and cosine(month) within wolf identity (Schielzeth and Forstmeier 2009; Barr et al. 2013). The human footprint covariate was z-transformed prior to analysis to facilitate model interpretation (Schielzeth 2010).

To evaluate the overall influence of individual and ecological predictors on monthly range size, we compared the full model with a null model containing only random effects using a likelihood-ratio test (anova function in R). Study-area effects were assessed by comparing the full model with a reduced model lacking the study-area term. Interaction significance was evaluated by contrasting the full model with reduced models lacking (i) the sex × status interaction (retaining both main effects) or (ii) the status × month interaction. Predicted effects of study area and other predictors were obtained using the effect function.

The entire analytical procedure was repeated for monthly distance travelled, using an identical model structure and the same dataset (N = 229); residual diagnostics indicated no evidence of dispersion issues (dispersion = 0.99), and the model accounted for a substantial proportion of variance (marginal R^2^ = 0.48, conditional R^2^ = 0.61).

## 3. Results

### 3.1. Individual spatial strategies

Among the 19 collared wolves, 17 were pack members at the beginning of monitoring: five of these changed status during the study period. Three individuals dispersed (WM2101, WF2101, WM2402; Fig. 2a) and two became floaters for several months (WM2301, WF2402; Fig. 2b). An additional two wolves (WF2002, WM2304) were already solitary at capture and behaved as floaters (Fig. 2b).

**Figure 2.**
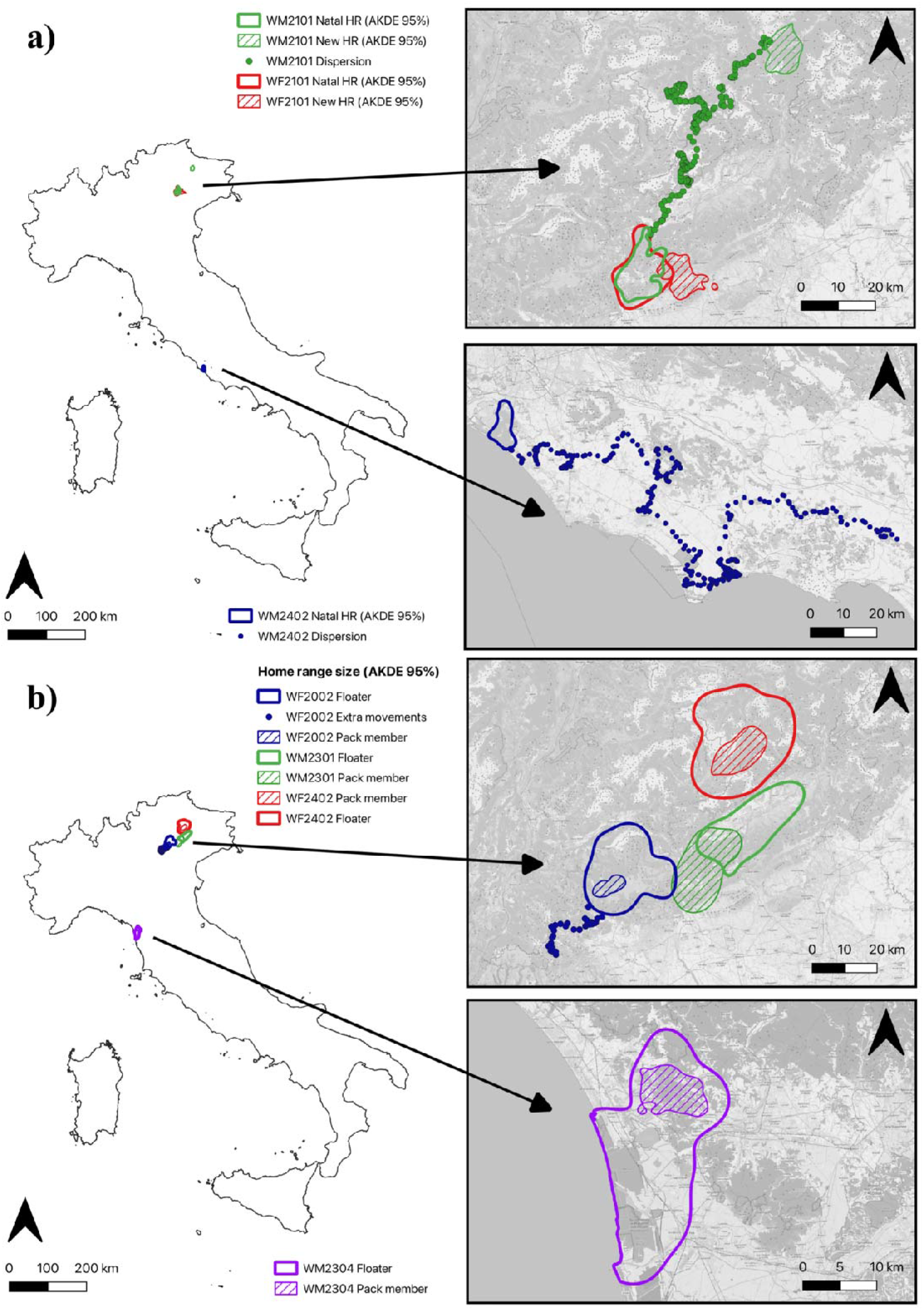
Home range dynamics and dispersal movements of GPS-collared wolves. Panel (a) shows dispersal movements of three individuals (WM2101, WF2101, and WM2402), illustrating routes from natal home ranges (HR) to settlement sites or last known locations. Panel (b) shows home ranges of four individuals (WF2002, WM2301, WM2304, and WF2402), highlighting spatial changes during transitions from floater to pack resident status, or vice versa, within a pack

WM2101 and WF2101, offspring of WF2001 from the Grappa Pack, showed a brief pre-dispersal phase before permanently leaving their natal range. WM2101 remained in the pack for ∼10 months, departed in late January, and established a new territory ∼70 km away within two weeks. In early March, WM2101 was detected forming a new stable pair with an adult female, based on camera trap records. Subsequent spatial behaviour, characterized by localized movements around a restricted area, suggested reproduction, which was later confirmed by camera trap recordings, documenting the presence of five pups in May.

Similarly, WF2101 began detaching from the natal pack in November and completed dispersal in mid-December, settling ∼10 km away within few hours. Shortly after settlement, WF2101 formed a new stable pair. As for WM2101, reproduction was initially inferred from spatial behaviour and subsequently confirmed by camera trap data, documenting the presence of pups in May (three pups; Table 2).

**Table 2.** Summary of the final outcomes of dispersal and floater phases for GPS-tracked wolves. For dispersing individuals, the duration and distance of dispersal movements are also reported.

| **Id** | **Area** | **Sex** | **Status** | **Dispersal duration** | **Distance from natal range (km)** | **Natal pack size** | **Outcome** |
| --- | --- | --- | --- | --- | --- | --- | --- |
| WF2002 | Pre-Alps | f | Floater | / | / | / | Formed a new pack; reproduction confirmed (five pups) |
| WM2101 | Pre-Alps/Alps | m | Dispersal | ~ 14 days | ~ 70 | 10 | Formed a new pack; reproduction confirmed (five pups) |
| WF2101 | Pre-Alps | f | Dispersal | ~ 4 hours | ~ 10 | 11 | Formed a new pack; reproduction confirmed (three pups) |
| WM2301 | Pre-Alps | m | Floater | / | / | 6 | Joined unrelated pack |
| WM2304 | SRE | m | Floater | / | / | / | Formed a new pack |
| WM2402 | PEC | m | Dispersal | ~ 21 days | ~ 120 | 12 | Unknown |
| WF2402 | Alps | f | Floater | / | / | 4 | Dead |

SRE: San Rossore Estate; PEC: Presidential Estate of Castreporziano; f: female; m: male; /: information not available (dispersal-related variables; monitoring started during the floater phase)

WM2402 (PEC) dispersed in mid-November together with another individual, as confirmed by camera traps. He reached his last known location ∼120 km from the natal range after ∼3 weeks; the collar signal was subsequently lost, likely due to poaching or device malfunction (Table 2).

WM2301 (Pre-Alps) displayed prolonged solitary behaviour within his natal territory before joining the Grappa Pack in late October, despite lacking genetic relatedness to its members (unpublished data). WF2402, an offspring of WF2201 (DBNP1 Pack), left her natal pack in late December and adopted a floater-like pattern while occasionally reusing parts of the maternal range, until she was found dead in summer 2025.

WF2002 and WM2304 were collared as solitaries and exhibited extended floater behaviour prior to pair formation. WF2002 ranged widely across the Asiago Plateau for ∼1 year, overlapping multiple pack territories, before pairing in December, forming a new pack, and reproducing in May (five pups). WM2304 (SRE) ranged solitarily for ∼10 months until early February, subsequently paired, formed a new pack, and reproduced in May (three pups). Both individuals showed pronounced reductions in range size when transitioning from floater to pack-member status (WF2002 AKDE 95%: 478.46 ⍰ 42.39 km^2^; WM2304 AKDE 95%: 224.08 ⍰ 48.16 km^2^)

### 3.2. Annual home range size and distance travelled variation

At the annual scale, pack-member wolf home range size exhibited a negative trend with estimated prey biomass, although this relationship was not statistically significant (r = –0.49; p = 0.10; Fig. S5) and showed no association with the average body mass of the main prey species (r = 0.20; p = 0.53; Fig. S6). Annual home ranges varied markedly among GPS-tracked wolves, ranging from 17.12 to 269.52 km^2^.

In FCNP, pack-member wolves exhibited mean annual home ranges of 34.54 ± 27.43 km^2^ (n = 3; AKDE 95%) and 32.8 ± 22.9 km^2^ (MCP 95%), values comparable to those observed in the Tyrrhenian coastal area of SRE (n = 2; 59.29 ± 8.98 km^2^ AKDE; 53.3 ± 7.1 km^2^ MCP) but markedly smaller than those in the Pre-Alps (n =2; 114.80 ± 26.27 km^2^ AKDE; 116.85 ± 19.16 km^2^ MCP) and in the Alps (n = 2; 185.21 ± 119.20 km^2^ AKDE; 162.85 ± 84.64 km^2^ MCP; Table 3). Similar patterns were observed for both summer and winter home ranges (Table S4).

**Table 3.** GPS-tracked wolves’ annual home range sizes estimated by means of Autocorrelation Kernel Density Estimation (AKDE) and Minimum Convex Polygon (MCP) and their annual distance travelled estimated by means of Continuous-Time Speed and Distance (CTSD) and Straight-Line Displacement (SLD)

| **ID** | **Area** | **Sex** | **Status** | **AKDE hrs (km<sup>2</sup>) 95%** | **AKDE hrs CI** | **MCP hrs (km<sup>2</sup>) 95%** | **CTSD dt (km/day)** | **CTSD dt CI** | **SLD dt (km/day)** |
| --- | --- | --- | --- | --- | --- | --- | --- | --- | --- |
| WF2001 | Pre-Alps | f | P | 133.37 | 117.57-150.16 | 130.4 | 32.83 | 32.26-33.40 | 7.15 |
| WF2002 | Pre-Alps | f | F | 468.52 | 372.79-575.03 | 420.7 | 29.53 | 29.24-29.82 | 7.27 |
| WF2101 | Pre-Alps | f | P | 96.22 | 87.16-105.72 | 103.3 | 36.45 | 35.77-37.12 | 5.68 |
| WF2202* | Alps | f | P | 269.52 | 222.86-320.56 | 222.7 | 26.67 | 26.30-27.05 | 6.94 |
|  |  |  |  | 100.89 | 89.48- | 103 | 24.18 | 24.89- | 6.35 |

| ID | Area | Sex | Status | AKDE hrs (km<sup>2</sup>)<br>95% | AKDE hrs<br>CI | MCP hrs<br>(km<sup>2</sup>) 95% | CTSD dt<br>(km/day) | CTSD dt<br>CI | SLD dt<br>(km/day) |
| --- | --- | --- | --- | --- | --- | --- | --- | --- | --- |
|  |  |  |  |  | 112.98 |  |  | 24.18 |  |
| WM2201 | FCNP | m | P | 20.34 | 19.14-21.58 | 21.8 | 23.65 | 23.33-23.97 | 4.54 |
| WF2203 | FCNP | f | P | 17.12 | 16.16-18.09 | 17.5 | 33.60 | 32.85-34.34 | 4.18 |
| WM2304 | SRE | m | F | 333.09 | 255.15-421.26 | 250.2 | 27.18 | 26.77-27.59 | 6.21 |
| WM2305 | FCNP | m | P | 66.16 | 60.12-72.48 | 59.1 | 27.00 | 26.74-27.27 | 6.08 |
| WF2401 | SRE | f | P | 52.93 | 49.07-56.95 | 48.3 | 55.05 | 54.21-55.88 | 7.75 |
| WM2401 | SRE | m | P | 65.64 | 60.19-71.32 | 58.3 | 51.41 | 50.82-51.99 | 7.05 |

FCNP: Foreste Casentinesi National Park; SRE: San Rossore Estate; PEC: Presidential Estate of Castelporziano; hrs: home range size; dt: distance travelled; CI: 95% Confidence Intervals; f: female; m: male; P: pack-member individual; F: floater individual. \*For wolf WF2202 the monitoring duration allowed the calculation of two annual home ranges and distance travelled

In contrast, distance travelled was more homogeneous across individuals, study areas, and seasons (Table 3, S4). Pack member wolves travelled an average of 28.08 ± 5.06 km/day in FCNP, 53.23 ± 2.57 km/day in SRE, 34.64 ± 2.56 km/day in the Pre-Alps, and 25.43 ± 1.76 km/day in the Alps. Overall, CTSD estimates were five times higher than classical SLD estimates (FCNP: 4.93 ± 1.01 km/day; SRE: 7.40 ± 0.50 km/day; Pre-Alps: 6.41 ± 0.74 km/day; Alps: 6.65 ± 0.42 km/day; Table 3).

Monthly range size and distance travelled variation

The full model for monthly range size was highly significant relative to the null model (likelihood ratio test: χ^2^ = 168.50, df = 12, P < 0.0001, Table S2), confirming the overall predictive value of the ecological and individual variables considered. Monthly range size differed significantly among study areas (full vs. area-reduced model: χ^2^ = 12.15, df = 4, P = 0.02), with wolves in the Alpine area exhibiting larger monthly ranges than those in the Apennines (FCNP) and in the coastal areas (SRE and PEC; Fig. 3a, Table 4). Among individual factors, the interaction between sex and social status significantly influenced monthly range size (full vs. interaction-reduced model: χ^2^ = 3.96, df = 1, P = 0.046). Floaters generally occupied larger ranges than pack-member wolves, and this difference was more pronounced in females than in males (Fig. 3b, Table 4). However, seasonal variation in the pack-member – floater difference was not detected (full vs. seasonality interaction-reduced model: χ^2^ = 1.71, df = 2, P = 0.42). The model also revealed a clear seasonal cycle, with a minimum in July and a maximum in January (Fig. 3c). The average human footprint had a weak but significant positive effect, indicating that wolves tended to use larger ranges in more anthropized portions of the study area (Fig. 3d, Table 4).

**Table 4.** Results of the Linear Mixed Model on GPS-tracked wolves’ monthly range size variation. Alps, female, and pack member were used as reference levels for the predictors study area, sex, and individual status, respectively.

| Predictor | Coefficient estimate | SE | P |
| --- | --- | --- | --- |
| Pre-Alps | -0.005 | 0.22 | 0.98 |
| FCNP | -1.67 | 0.57 | 0.01 |
| SRE | -1.24 | 0.54 | 0.04 |
| PEC | -1.09 | 0.68 | 0.13 |
| Sin(month) | 0.17 | 0.05 | <0.01 |
| Cos(month) | 0.18 | 0.06 | <0.01 |
| HF | 0.40 | 0.16 | 0.02 |
| Male | 0.45 | 0.43 | 0.32 |
| Floater | 2.05 | 0.15 | < 0.001 |
| Male x Floater | -0.75 | 0.26 | <0.01 |

FCNP: Foreste Casentinesi National Park; SRE: San Rossore Estate; PEC: Presidential Estate of Castelporziano; month: month of the year (integer, then transformed into radians), inserted as circular variable by means of a sin and a cosine terms; HF: human footprint; SE: standard error of estimated coefficient; P: P-value; value in grey indicate data with limited interpretability. See section 2.5 for more details about model and fixed term structure

**Fig. 3.**
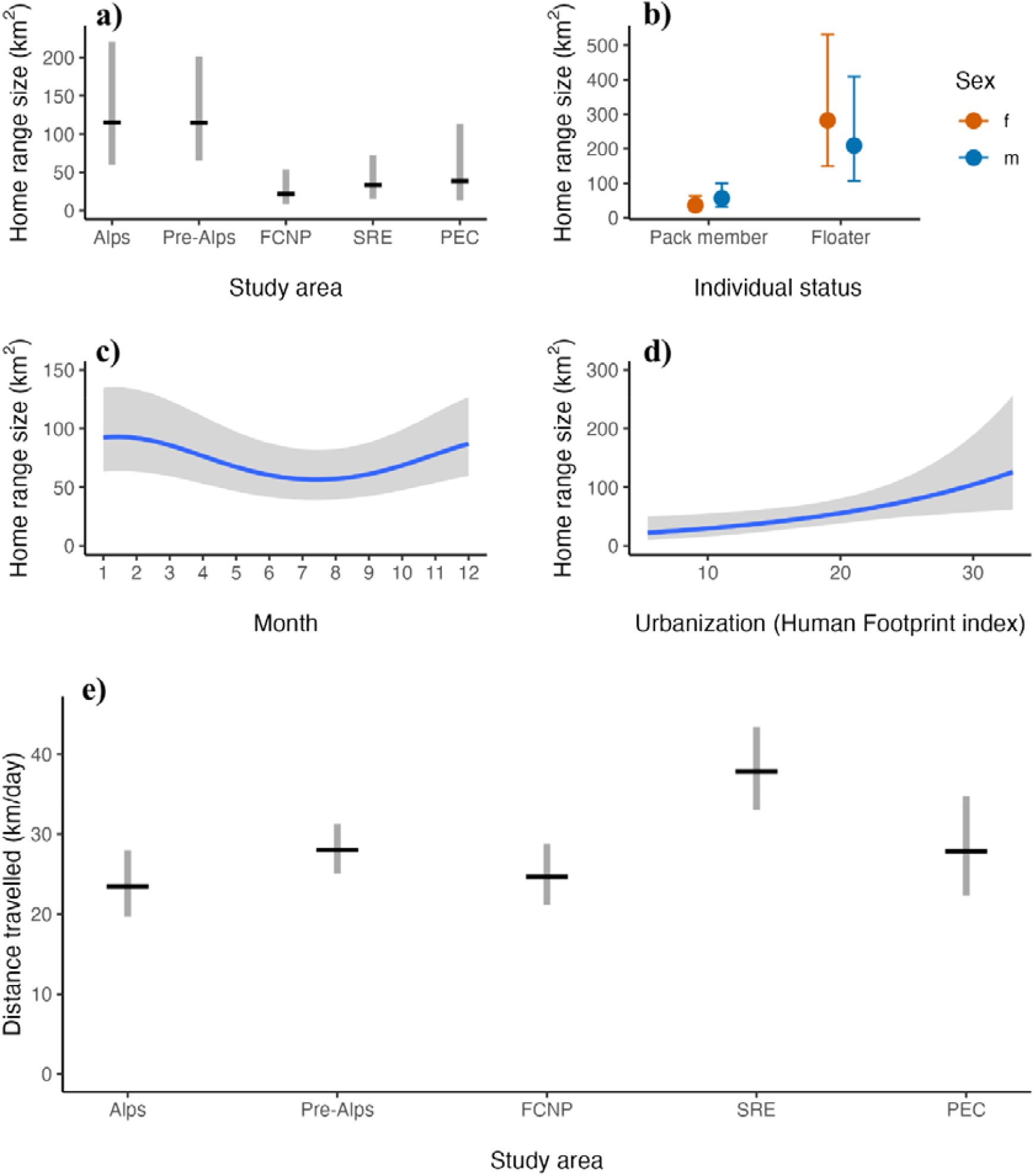
Variation of GPS-tracked wolves’ monthly range size and distance travelled. Panels (a-d) show variation in monthly rage size across a) study areas (FCNP: Foreste Casentinesi National Park; SRE: San Rossore Estate; PEC: Presidential Estate of Castelporziano), b) individual status classes and sexes, c) months the year and d) the of urbanization gradient measured by the Human Footprint Index, as predicted by the Linear Mixed Model. Panel e) shows variation in monthly distance travelled across study areas, as predicted by the Linear Mixed Model

The full model for monthly distance travelled was also significant relative to the null model (χ^2^ = 32.20, df = 12, P < 0.01, Table S3). Study area strongly improved model fit (χ^2^ = 18.28, df = 4, P < 0.01), showing that travelled distances varied across regions. Wolves from SRE travelled significantly farther than those from the other study areas (Fig. 3e, Table 5). Neither the sex × status interaction (χ^2^ = 0.31, df = 1, P = 0.57) nor the status × seasonality interaction (χ^2^ = 1.63, df = 2, P = 0.44) significantly affected travelled distances.

**Table 5.** Results of the Linear Mixed Model on GPS-tracked wolves’ monthly distance travelled variation. Alps, female, and pack member were used as reference levels for the predictors study area, sex, and individual status, respectively.

| Predictor | Coefficient estimate | SE | P |
| --- | --- | --- | --- |
| Pre-Alps | 0.18 | 0.08 | 0.04 |
| FCNP | 0.05 | 0.11 | 0.66 |
| SRE | 0.48 | 0.11 | <0.01 |
| PEC | 0.17 | 0.15 | 0.26 |
| Sin(month) | 0.0002 | 0.03 | 0.99 |
| Cos(month) | 0.06 | 0.03 | 0.07 |
| HF | 0.02 | 0.04 | 0.64 |
| Male | 0.008 | 0.07 | 0.91 |
| Floater | -0.05 | 0.05 | 0.34 |

FCNP: Foreste Casentinesi National Park; SRE: San Rossore Estate; PEC: Presidential Estate Castelporziano; month: month of the year (integer, then transformed into radians), inserted as circular variable by means of a sin and a cosine terms; HF: human footprint; SE: standard error of estimated coefficient; P: P-value. See section 2.5 for more details about model and fixed term structure

## 4. Discussion

Our findings reveal substantial variation in wolf range sizes across study areas. In Alps and pre-Alps, the mean annual home range sizes were consistent with those estimated in the various regions of Central Europe (Okarma et al. 1998; Findo and Chovancova 2004; Kusak et al. 2005; Jędrzejewski et al. 2007; Karamanlidis et al. 2016; Mysłajek et al. 2018; Vorel et al. 2024), while wolves in the other areas occupied exceptionally small territories. These patterns were consistent across monthly, seasonal, and annual estimates, particularly among pack-member individuals (Table 3; S4). Prey biomass availability in FCNP and PEC was higher and in SRE was markedly higher than in Alpine areas (Franceschi et al. 2023), suggesting that resource density was the primary driver of reduced home range size, as wolves can meet their energetic requirements within smaller areas (Fuller 2003; Jedrzejewski et al. 2007; Mattisson et al. 2013). However, our correlation analyses reveal a weak relationship between the annual home range size of pack-member wolves and prey biomass, and no relationship at all with the size of the main prey species. This suggests that, within the contexts considered here, prey availability and prey size alone cannot fully explain the observed variation in range size. Given the limited number of pack-member wolves with annual home ranges available for these analyses, it remains possible that the lack of significance reflects insufficient statistical power rather than the absence of an ecological effect. Recolonization history, however, might have played a secondary, yet substantial, role: FCNP, with the longest continuous wolf presence and likely the highest intra-specific competition for space, exhibited the smallest ranges even though PEC and especially SRE offered richer prey resources (Fig. 3a). Indeed, the home ranges in FCNP are significantly smaller than previous estimates conducted in other Apennine areas, where prey availability was already high even if scavenging played a relevant role (Ciucci et al. 2020; Mancinelli et al. 2018). This suggests that rising wolf density and intensified intra11specific competition might have played a role in reducing home range size in FCNP, a change made possible by ample prey resources. Recently recolonized areas are typically characterized by lower wolf densities due to limited time for population growth, whereas long-established population tend to exhibit higher densities (Pletscher et al. 1997; Mysłajek et al. 2018; Mattioli et al. 2018). Collectively, these findings indicate that wolves have reduced spatial needs in resource11rich environments, but also that increasing intra11specific competition may drive further reductions in territory size.

It remains unclear, however, how continuing population growth will alter spatial behaviour in regions with lower resource availability, such as Alpine areas. Assuming that the prey density of the Alpine regions will remain stable, it will be indeed worth to monitor the possible outcomes of the likely demographic increase of wolves in Alps and pre-Alps in the near future. Packs may be expected to either maintain large home ranges, which may favour dispersive behaviour when local resources are insufficient to sustain all pack members and intra-specific competition increases with pack size, or, alternatively, reduce their home ranges, as observed in FCNP, by exploiting alternative resources, such as domestic livestock. Interestingly, livestock generally contributes more to wolf diet in pre-Alpine regions than in the other areas included in this study, with an increase during the summer grazing period (Cerri et al. 2026). In addition, pack size may mediate intraspecific competition and space use through both within-pack and between-pack processes: as pack size increases, per-capita access to resources may decrease, potentially intensifying within-pack competition and thereby favouring dispersal. Consistently with this possibility, the natal packs of the dispersers documented here were relatively large (WM2101 and WF2101 dispersed from a pack of 11 individuals, and WM2402 from a pack of 12 individuals, including the focal animals, Table 2), suggesting that these dispersal events may have occurred under conditions of high pack size, although the limited number of cases warrants cautious interpretation. Conversely, increasing local density may strengthen inter-pack competition and territorial constraints, which could limit spatial expansion and instead contribute to reduced space use. Targeted studies integrating movement data with independent estimates of pack size and neighbouring pack density will be needed to disentangle these mechanisms and assess how they jointly shape home-range dynamics. A further interesting issue is represented by wolves’ packs living in small, partially fenced, protected areas like in the case of San Rossore (SRE) and Castelporziano Estates (PEC): in both cases a high local ungulate density gives a rich food basis for wolves that just outside the two areas face a substantially reduced prey availability and the presence of large cities like Pisa and Rome. Their home range size is not only small but also shaped by the border of the Estates (Fig. S4) where they bred and spent all their time. Such cases are quite common in a country like Italy, where high human pressure caused pronounced environmental fragmentation and small portion of lands well preserved and rich in wildlife were left for hunting purposes in previous centuries, then becoming protected areas.

The larger ranges of floater wolves compared to pack members (Fig. 3b) align with their social behaviour, leading to less spatially predictable and more wide-ranging movement patterns (Mech and Boitani 2003), as also observed by Blount et al. (2024). WM2304, a floater male of SRE, had range sizes comparable to the floaters inhabiting Alpine areas, despite the pack-member wolf home ranges in the two areas being very different. Overall, sex-related differences were limited among pack-member wolves. However, among floaters, females exhibited slightly larger monthly ranges than males (Fig. 3b). A possible explanation for this apparent discrepancy is that males are more frequently involved in long-distance dispersal events, characterized by direct movements across large spatial scales that do not correspond to a defined range (Fabbri et al. 2014; Sanz-Perez et al. 2018) and were therefore not included in our analysis (e.g., the 70 km and 120 km dispersal of WM2101 and WM2402 respectively). Females, by contrast, may more often adopt a floater strategy, spending extended periods in non-territorial conditions near established packs. Such prolonged exploratory behaviour, involving recurrent movements within and around neighbouring territories, can result in larger and more spatially complex monthly ranges, as observed for WF2002 and WF2402 (Mech and Boitani 2003). This interpretation is consistent with the broader pattern of stronger female philopatry observed in wolves and other mammalian species (Greenwood 1980; Vanholdt et al. 2008; Ausband et al. 2017). Females may invest more time in exploring familiar or adjacent areas to locate reproductive opportunities or vacant territories, while males tend to disperse farther. Nonetheless, given the limited number of floaters and dispersal individuals monitored in this study, we caution against overinterpreting this pattern, which may also reflect individual variability rather than broader behavioural trends. An interesting outcome is related to the fates of floaters and dispersing individuals. Among the three documented dispersal events, two led to successful reproduction through the establishment of a new pack (Fig. 2a). Among the four floater individuals, two formed new packs, and one joined an existing pack, with two of them subsequently reproducing (Fig. 2b). These results confirm that both dispersal and floater phases often lead to breeding success, even within a mature wolf population such as the one in Italy. The dispersal events we documented occurred consistently between October and January, suggesting a seasonal pattern in line with other findings (Gese and Mech 1991). The distances of dispersal observed also varied considerably: two males dispersed approximately 70 km and 120 km from their natal territories, while one female dispersed a much shorter distance of about 10 km. These values fall within the range reported in previous European studies, where dispersal distances can span from a few kilometers to several hundred kilometers, and further support the documented pattern of males dispersing over longer distances than females (Kojola et al. 2006; Fabbri et al. 2014; Sanz-Perez et al. 2018; Marucco et al. 2022). These findings suggest that, despite the risks associated with prolonged solitary phases, both dispersal and floating can represents viable strategies for securing reproductive opportunities and contributing to population expansion.

The variation in monthly range sizes observed throughout the year was consistent with the wolf’s reproductive cycle and social behaviour (Fig. 3c). In summer, when monthly range sizes decreased, pack movements are known to be concentrated around the den area because the pups, born in spring, are not yet able to follow the adults (Jedrzejewski et al. 2007; Kusak et al. 2005; Roffler and Gregovich 2018). The gradual increase in monthly range sizes in early autumn may correspond to the period when the pups start following the adults, allowing the pack to expand its movements beyond the den area (Ciucci et al. 1997). This explanation, however, should apply primarily to pack members. Interestingly, we did not detect a significant interaction between individual status and month of the year, indicating that the reduction in summer range size was also evident in floater wolves. Although we lack direct evidence to fully explain this pattern, we can speculate that the observed seasonal patterns in space use likely reflect the combined influence of seasonal changes in resource availability, social behaviour, and anthropogenic disturbance. In our study, movements outside the natal pack were more frequently observed during late autumn and winter, a period characterized by reduced prey availability and increased intra-pack competition as pups begin to follow the pack, leading to lower food availability per individual (Mech and Boitani 2003; Morales-Gonzalez et al. 2022). These conditions may promote the departure of subordinate individuals in advance of the breeding season and, once solitary, wider-ranging behaviour driven by energetic constraints and mate-searching activity, resulting in larger spatial use (Jimenez et al. 2017). Conversely, during summer, also solitary wolves exhibited smaller home ranges. Increased availability of vulnerable wild prey, including ungulate newborns, and the presence of domestic livestock in open pastures during summer months (Cerri et al. 2026), may allow wolves, regardless of social status, to meet their energetic needs within smaller areas. At the same time, peak anthropogenic disturbance associated with increased tourism and recreational activities may further constrain movements, as previously reported for extraterritorial movements of pack members (Mancinelli and Ciucci 2018).

Finally, we found that, after accounting for study area-level differences, wolf range slightly increased with increasing human footprint (Fig. 3d). This suggests that, when moving to more disturbed and/or fragmented landscapes, wolves may need to use larger areas to meet their ecological requirements. Similar patterns have been observed in other studies, in which home range expansion was associated with reduced habitat quality, lower prey accessibility, or the need to avoid human activity (Rich et al. 2012; Mancinelli et al. 2018). In more urbanized environments, suitable habitats and resources tend to be more dispersed and less predictable, potentially requiring wolves to adopt broader movement strategies. Additionally, high levels of human disturbance may encourage wolves to expand their space use, allowing them to access quieter, less frequented areas farther from human activity.

Annual mean distances travelled calculated with the classical SLD approach, both for pack member inhabiting the Apennines (FCNP) and Alps, were similar to those reported by previous studies in areas with abundant resources (Kolenosky and Johnston 1967; Kusak et al. 2005; Jedrzejewski et al. 2007). However, the distance travelled calculated with the CTSD revealed that wolves likely travelled distances about five times longer (Tab. 3 and S4), which may be considered more realistic since the CTSD estimates account for the spatiotemporal autocorrelation of GPS data and are thus unbiased by the sampling frequency (Fleming et al. 2014; Fleming et al. 2019; Noonan et al. 2019). We found little variation in distance travelled across temporal scales or individual traits, with similar values among most wolves. The only clear exception was SRE, where wolves travelled significantly longer distances despite small home ranges and very high prey availability (Fig. 3e). Although this pattern appears counterintuitive, given that high prey density typically reduces movement requirements (Kolenosky and Johnston 1967; Johnson et al. 2017), the frequent overkilling and partial consumption of fallow deer in SRE (Brogi et al. 2025b) may partly account for it, as high prey availability can favour repeated hunting events and thus increase daily displacement. Other factors might also contribute, including the dense road network (Musiani et al. 1998), the compartmentalized structure of this highly productive landscape, and social movements during the rendezvous period, when CTSD distances peak in SRE (Jedrzejewski et al. 2001; Table 4S). Field inspections of GPS clusters supported this interpretation by documenting repeated pack use of rendezvous areas and associated social activity (e.g., communal resting sites and shared feeding at kill sites). However, interpretation should remain cautious, as the highest values (>40 km/day) are largely driven by two individuals (WM2401 and WF2401), suggesting that individual behavioural variability may play a substantial role. Further investigation is thus needed to disentangle these effects.

Considering that the distances travelled in other areas did not differ significantly (Fig. 3e), the comparison with range size suggested that even individuals with small ranges travelled, in the time unit, the similar average distances travelled by wolves with much larger ranges. These patterns can be attributed to the different effects that increased spatial competition has on range size and distance travelled. Wolves might have been forced to use smaller ranges due to heightened competition for space (Tinbergen 1957), ultimately reducing their need for long displacements. However, increased competition might also lead wolves to patrol their territory boundaries more frequently to prevent intrusions (Zub et al. 2003), potentially counterbalancing the effect of reduced range size on the distance travelled.

The rapid but spatially uneven recovery of the Italian wolf population, exposing wolves to a wide range of ecological conditions, offered a valuable opportunity to assess how environmental context and human influence shape spatial behaviour. The ability of wolves to reduce spatial requirements under abundant resources, amplified by high levels of intraspecific competition, indicates that they may reach higher densities than traditionally assumed, a possibility that may warrant timely re-evaluation of conservation and management strategies. At the same time, we found that patterns of range size and movement vary substantially among regions and environmental settings, sometimes diverging from classical expectations of wolf ecology. Collectively, these results highlight the species’ marked ecological flexibility and emphasize the need for European conservation and management strategies to rely on fine-scale, context-specific information on wolf space use and population dynamics.

## Statements and Declarations

### Ethical statement

This study complies with guidelines of ARRIVE (Animal Research: Reporting of In Vivo Experiments), Guidance on the operation of the Animals (Scientific Procedures) Act 1986, and EU Directive 2010/63 for the protection of animals used for scientific purposes. The capture and handling of wolves for scientific research were approved by the relevant Italian authority, Ministero della Transizione Ecologica, with protocol n. 0070095 del 06.06.2022 for the FCNP study area, with protocol n. 0014897 del 05.07.2018, protocol n. 0123622 del 06.10.2022, and protocol n. 0070097 del 06.06.2022 for the pre-Alps and Alps study areas, with protocol n. 0173939 del 30.10.2023 for the PEC study area, and with protocol n. 0068145 del 31.05.2022 for the SRE study area.

### Author contributions

Conceptualization: MA; Data collection and curation: SC, CB, MZ, SL, DB, LC, CT, MDF, PB; Formal analysis: SC, RB; Funding acquisition and project administration: MA; Supervision: RB, MA; Writing – original draft: SC, RB, MA; Writing – review and editing: MZ, MDF, DB, NC, PB, EV.

## Acknowledgments

We would like to express our sincere gratitude to all students and volunteers who collaborated in collecting the data. The authors sincerely thank the personnel of the Castelporziano Presidential Estate, as well as the support of “Segretariato generale della Presidenza della Repubblica Italiana”, for their invaluable assistance, for providing essential logistic help throughout the study, and for having made available the necessary infrastructure and research facilities, which were instrumental in the successful completion of their research, especially during fieldwork.

## References

Apollonio M, Mattioli L, Scandura M, Mauri L, Gazzola A, Avanzinelli E (2004) Wolves in the Casentinesi Forests: Insights for wolf conservation in Italy from a protected area with a rich wild prey community. Biological Conservation 120(2):249–260.

Apollonio M, Andersen R, Putman R (2010) European Ungulates and Their Management in the 21st Century. Cambridge University Press, Cambridge, UK.

Arnemo JM, Evans AL, Ahlqvist P, Segerström P, Liberg O (2013) Evaluation of medetomidine-ketamine and atipamezole for reversible anesthesia of free-ranging gray wolves (Canis lupus). Journal of Wildlife Diseases 49(2):403–407. 10.7589/2011-12-366

Ausband DE, Mitchell MS, Stansbury CR, Stenglein JL, Waits LP (2017) Harvest and group effects on pup survival in a cooperative breeder. Proc. R. Soc. B: Biological Sciences 284(1855). 10.1098/rspb.2017.0580

Avanzinelli E, Calderola S, Valbusa F, Parricelli P, Pedrotti L, Bragalanti N, Marucco F (2017) Lo Status del lupo in Veneto. In: Marucco et al. (2017). Lo Status della popolazione di lupo sulle Alpi Italiane e Slovene 2014-2016 Relazione tecnica, Progetto LIFE 12 NAT/IT/00080 WOLFALPS – Azione A4

Avanzinelli E, Calderola S, Giombini V, Marucco F (2018) Lo Status del lupo in Veneto 2014-2018. Relazione tecnica, Progetto LIFE 12 NAT/IT/00080 WOLFALPS – Azione D1. In: Marucco et al. (2018). Lo Status della popolazione di lupo sulle Alpi Italiane e Slovene 2014-2018 Relazione tecnica, Progetto LIFE 12 NAT/IT/00080 WOLFALPS – Azione A4 e D1

Baayen RH (2008) Analyzing linguistic data. Cambridge, UK. Cambridge University Press New York.

Ballard WB, Ayres LA, Krausman PR, Reed DJ, Fancy SG (1997) Ecology of Wolves in Relation to a Migratory Caribou Herd in Northwest Alaska. Wildlife monographs: 3–47

Barr DJ, Levy R, Scheepers C, Tily HJ (2013) Random effects structure for confirmatory hypothesis testing: Keep it maximal. Journal of Memory and Language 68(3):255–278. 10.1016/j.jml.2012.11.001

Bassi E, Donaggio E, Marcon A, Scandura M, Apollonio M (2012) Trophic niche overlap and wild ungulate consumption by red fox and wolf in a mountain area in Italy. Mammalian Biology 77(5):369–376. 10.1016/j.mambio.2011.12.002

Bateman PW, Fleming PA (2012) Big city life: Carnivores in urban environments. Journal of Zoology 287(1):1–23. 10.1111/j.1469-7998.2011.00887.x

Bates D, Mächler M, Bolker B, Walker S (2015) Fitting Linear Mixed-Effects Models Using lme4. Journal of Statistical Software 67:1–48.

Blanco JC, Cortés Y (2007) Dispersal patterns, social structure and mortality of wolves living in agricultural habitats in Spain. Journal of Zoology 273(1):114–124. 10.1111/j.1469-7998.2007.00305.x

Blašković S, Gomerčić T, Topličanec I, Sindičić M(2022) Temporal overlap of human and apex predator activity on wildlife trails and forest roads. Journal of Vertebrate Biology, 71(22029):22029–1.

Blount JD, Green AM, Chynoweth M, Kittelberger KD, Hipólito D, Bojarska K, Çoban E, Kusak J, Sekercioglu CH (2024) Seasonal activity patterns and home range size of wolves in the human-dominated landscape of northeast Türkiye. Wildlife Biology e01257. 10.1002/wlb3.01257

Boitani L (2000) Action plan for the conservation of the wolves (Canis lupus) in Europe. Nature and environment, Council of Europe Publishing 113:1–86.

Boscagli G (1985) Attuale distribuzione geografica e stima numerica del lupo sul territorio italiano. Natura. Rivista di scienze naturali. Vol.76 fasc.1/4 ISSN 0369-6243.

Breitenmoser U (1998) Large predators in the Alps: The fall and rise of man’s competitors. Biological Conservation 83(3):279– 289. 10.1016/S0006-3207(97)00084-0

Brogi R, Chirichella R, Brivio F, Merli E, Bottero E, Apollonio M (2021) Capital-income breeding in wild boar: A comparison between two sexes. Scientific Reports 11(1): 4579.

Brogi R, Neirotti G, Cerri J, Lazzaroni M, Marshall-Pescini S, Mattioli L, Apollonio M (2025a) Wolves on the phone: Public calls reveal a rise in urban concerns as wolves recolonize human-dominated areas. Ambio:1-12. 10.1007/s13280-025-02264-z

Brogi R, Bongi P, Del Frate M, Sieni S, Cavallera A, Apollonio M (2025b) Intra-guild competition and ecosystem services of mammal scavengers in a new colonized wolf landscape. Behavioral Ecology and Sociobiology 79(2): 20. 10.1007/s00265-025-03565-9

Cagnolaro L, Rosso D, Spagnesi M, Venturi B (1974) Investigation on the wolf (Canis lupus) distribution in Italy and in Canton Ticino and Canton Grigioni (Switzerland). Ricerche di Biologia della Selvaggina 59:1–75.

Calabrese JM, Fleming CH, Gurarie E (2016) ctmm: An R package for analyzing animal relocation data as a continuous-time stochastic process. Methods in Ecology and Evolution 7(9):1124–1132. 10.1111/2041-210X.12559

Caniglia R, Fabbri E, Galaverni M, Milanesi P, Randi E (2014) Noninvasive sampling and genetic variability, pack structure, and dynamics in an expanding wolf population. Journal of Mammalogy 95(1):41–59.

Cavazza S, Brogi R, Apollonio M (2023) Sex-specific seasonal variations of wild boar distance traveled and home range size, Current Zoology 70(3):284–290. 10.1093/cz/zoad021

Cerri J, Brogi R, Musto C, Bassi E, Ventura G, Bianchi A, Delogu M, Scandura M, & Apollonio M (2026) Identifying and overcoming knowledge gaps in the feeding ecology of grey wolves inhabiting anthropized landscapes. Current Zoology, zoag010. 10.1093/cz/zoag010

Ciucci, P., Boitani, L., Francisci, F., Andreoli, G., 1997. Home range, activity and movements of a wolf pack in central Italy. Journal of Zoology, 243(4), 803–819. 10.1111/j.1469-7998.1997.tb01977.x

Ciucci P, Mancinelli S, Boitani L, Gallo O, Grottoli L (2020) Anthropogenic food subsidies hinder the ecological role of wolves: Insights for conservation of apex predators in human-modified landscapes. Global Ecology and Conservation 21:e00841. 10.1016/j.gecco.2019.e00841

Del Frate M, Bongi P, Tanzillo L, Russo C, Benini O, Sieni S, Scandura M, Apollonio M (2023) A Predator on the Doorstep: Kill Site Selection by a Lone Wolf in a Peri-Urban Park in a Mediterranean Area. Animals 13(3):480. 10.3390/ani13030480

Fabbri E, Miquel C, Lucchini V, Santini A, Caniglia R, Duchamp C, Weber J, Lequette B, Marucco F, Boitani L, Fumagalli L, Taberlet P, Randi E (2007) From the Apennines to the Alps: Colonization genetics of the naturally expanding Italian wolf (Canis lupus) population. Molecular Ecology 16(8):1661–1671. 10.1111/j.1365-294X.2007.03262.x

Fabbri E, Caniglia R, Kusak J, Galov A, Gomerčić T, Arbanasić H, Huber D, Randi E (2014) Genetic structure of expanding wolf (Canis lupus) populations in Italy and Croatia, and the early steps of the recolonization of the Eastern Alps. Mammalian Biology 79:138–148. 10.1016/j.mambio.2013.10.002

Ferretti F, Oliveira R, Rossa M, Belardi I, Pacini G, Mugnai S, Fattorini N, Lazzeri L (2023) Interactions between carnivore species: limited spatiotemporal partitioning between apex predator and smaller carnivores in a Mediterranean protected area. Frontiers in zoology 20(1):20.

Findo S, Chovancova B (2004) Home ranges of two wolf packs in the Slovak Carpathians. Folia Zool. 53(1):17–26.

Fleming CH, Calabrese JM, Mueller T, Olson KA, Leimgruber P, Fagan WF (2014) From fine-scale foraging to home ranges: a semivariance approach to identifying movement modes across spatiotemporal scales. The American Naturalist 183(5):E154–E167.

Fleming CH, Fagan WF, Mueller T, Olson KA, Leimgruber P, Calabrese JM (2015) Rigorous home range estimation with movement data: A new autocorrelated kernel density estimator. Ecology 96(5):1182–1188. 10.1890/14-2010.1

Fleming CH, Noonan MJ, Medici EP, Calabrese JM (2019) Overcoming the challenge of small effective sample sizes in home-range estimation. Methods in Ecology and Evolution 10(10): 1679–1689. 10.1111/2041-210X.13270

Franceschi S, Bongi P, Del Frate M, Fattorini L, Apollonio M (2023) A sampling strategy for habitat selection, mapping, and abundance estimation of deer by pellet counts. The Journal of Wildlife Management 87(2):e22345.

Franzetti B (2024) Risultati dei campionamenti delle popolazioni di ungulate presenti nella Tenuta di Castelporziano per l’anno 2024 e proposte inerenti i Piani di contenimento numeric del Cinghiale, del Daino e del Cervo per la stagione 2024-2025.

Fratianni S, Acquaotta F (2017) The Climate of Italy. In: Soldati M, Marchetti M (eds) Landscapes and Landforms of Italy. Cham: Springer International Publishing: 29–38

Fritts SH, Mech LD (1981) Dynamics, movements, and feeding ecology of a newly protected wolf population in Northwestern Minnesota. Wildlife Monographs 80:6–79.

Fuller TK (1989) Population dynamics of wolves in north-central Minneesota. Wildlife Monographs, 105:3–41.

Fuller TK, Mech LD, Cochrane JF (2003) Wolf population dynamics. In: Mech LD, Boitani L (eds) Wolves: Behaviour, Ecology and Conservation. University of Chicago Press, Chicago: 161–191.

Gese EM, Mech LD (1991) Dispersal of wolves (Canis lupus) in northeastern Minnesota, 1969-1989. Can. J. Zool. 69:2946–2955.

Greenwood PJ (1980) Mating system, philopatry and dispersal in birds and mammals. Anim. Behav. 28:1140–1162.

Guerrasio T, Brogi R, Marcon A, Apollonio M (2022) Assessing the precision of wild boar density estimations. Wildlife Society Bulletin 46(4):e1335. 10.1002/wsb.1335

Holz P, Holz RM, Barnett JE (1994) Effects of atropine on medetomidine/ketamine immobilization in the gray wolf (Canis lupus). Journal of Zoo and Wildlife Medicine 25(2):209–213.

Huxley JS (1934) A natural experiment on the territorial instinct. British Birds 27(10):270–277.

Jarausch A, Harms V, Kluth G, Reinhardt I, Nowak C (2021) How the west was won: genetic reconstruction of rapid wolf recolonization into Germany’s anthropogenic landscape. Heredity 127:92–106.

Jędrzejewski W, Schmidt K, Theuerkauf J, Jedrzejewska B, Okarma H (2001) Daily movements and territory use by radio-collared wolves (Canis lupus) in Bialowieza Primeval Forest in Poland. Can J Zool 79:1993–2004.

Jędrzejewski W, Branicki W, Veit C, Medugorac I, Pilot M, Bunevich AN, Jędrzejewska B, Schmidt K, Theuerkauf J, Okarma H, Gula R, Szymura L, Förster M (2005) Genetic diversity and relatedness within packs in an intensely hunted population of wolves Canis lupus. Acta Theriologica 50:3–22.

Jedrzejewski W, Schmidt K, Theuerkauf J, Jedrzejewska B, Kowalczyk R (2007) Territory size of wolves Canis lupus: Linking local (Białowieża Primeval Forest, Poland) and Holarctic-scale patterns. Ecography 30(1):66–76. 10.1111/j.0906-7590.2007.04826.x

Jimenez MD, Bangs EE, Boyd DK, Smith DW, Becker SA, Ausband DE, Woodruff SP, Bradley EH, Holyan J, Laudon K (2017) Wolf disperesal in the Rocky Mountains, Western United States: 1993-2008. Journal of Wildlife Management 81:581–592. 10.1002/jwmg.21238

Johnson I, Brinkman T, Lake B, Brown C (2017). Winter hunting behavior and habitat selection of wolves in a low-density prey system. Wildlife Biology 1:1–9. 10.2981/wlb.00290

Karamanlidis AA, De Gabriel Hernando M, Georgiadi L., Kusak, J., 2016. Activity, movement, home range and habitat use of an adult gray wolf in a Mediterranean landscape of northern Greece. Mammalia, 81(1). 10.1515/mammalia-2015-0091

Kojola I, Aspi J, Hakala A, Heikkinen S, Ilmoni C, Ronkainen S (2006) Dispersal in an expanding wolf population in Finland. Journal of Mammalogy 87(2):281–286. 10.1644/05-MAMM-A-061R2.1

Kolenosky GB, Johnston DH (1967) Radio-Tracking Timber Wolves in Ontario. American Zoologist 7(2):289–303. 10.1093/icb/7.2.289

Kusak J, Skrbinšek AM, Huber D (2005) Home ranges, movements, and activity of wolves (Canis lupus) in the Dalmatian part of Dinarids, Croatia. European Journal of Wildlife Research 51(4):254–262. 10.1007/s10344-005-0111-2

La Morgia V, Marucco F, Aragno P, Salvatori V, Gervasi V, De Angelis D, Fabbri E, Caniglia R, Velli E, Avanzinelli E, Boiani MV, Genovesi P (2022) Stima della distribuzione e consistenza del lupo a scala nazionale 2020/2021. Relazione tecnica realizzata nell’ambito della convenzione ISPRA-Ministero della Transizione Ecologica “Attività di monitoraggio nazionale nell’ambito del Piano di Azione del lupo”.

Mancinelli S, Boitani L, Ciucci P (2018) Determinants of home range size and space use patterns in a protected wolf (Canis lupus) population in the central Apennines, Italy. Canadian Journal of Zoology 96(8):828–838. 10.1139/cjz-2017-0210

Mancinelli S, Ciucci P (2018) Beyond home: Preliminary data on wolf extraterritorial forays and dispersal in Central Italy. Mammalian Biology 93:51–55. 10.1016/j.mambio.2018.08.003

Marucco F, Pilgrim KL, Avanzinelli E, Schwarts MK, Rossi L (2022) Wolf dispersal patterns in the Italian Alps and implications for wildlife diseases spreading. Animals 12(10):1260. 10.3390/ani12101260

Mattioli L, Canu A, Passilongo D, Scandura M, Apollonio M (2018) Estimation of pack density in grey wolf (Canis lupus) by applying spatially explicit capture-recapture models to camera trap data supported by genetic monitoring. Frontiers in zoology 15:1–15. 10.1186/s12983-018-0281-x

Mattisson J, Sand H, Wabakken P, Gervasi V, Liberg O, Linnell JDC, Rauset GR, Pedersen HC (2013) Home range size variation in a recovering wolf population: Evaluating the effect of environmental, demographic, and social factors. Oecologia 173(3):813– 825. 10.1007/s00442-013-2668-x

Mech LD, Meier TJ, Burch JW, Adams LG (1995) Patterns of prey selection by wolves in Denaly national park, Alaska. In Ecology and conservation of wolves in a charging world. Proceedings of the second North American symposium on wolves: Occasional Publication 35, Edmonton, AB, pp 231–243.

Mech LD, Boitani L (2003) Wolf social ecology. In Mech LD, Boitani L (eds) Wolves: Behaviour, Ecology and Conservation. University of Chicago Press, Chicago, pp 1–34

Merli E, Mattioli L, Bassi E, Bongi P, Berzi D, Ciuti F, Luccarini S, Morimando F, Viviani V, Caniglia R, Galaverni M, Fabbri E, Scandura M, Apollonio M (2023) Estimating Wolf Population Size and Dynamics by Field Monitoring and Demographic Models: Implications for Management and Conservation. Animals 13(11):1735. 10.3390/ani13111735

Messier F (1985) Social organization, spatial distribution, and population density of wolves in relation to moose density. Canadian Journal of Zoology 63(5):1068–1077. 10.1139/z85-160

Mohr C (1947) Table of equivalent populations of North American small mammals. American Midland Naturalist 37:223-249 Morales-González A, Fernández-Gil A, Quevedo M, Revilla E (2021) Patterns and determinants of dispersal in grey wolves (Canis lupus). Biological Reviews 97(2):466–480. 10.1111/brv.12807

Mu H, Li X, Wen Y, Huang J, Du P, Su W, Miao S, Geng M (2022) A global record of annual terrestrial Human Footprint dataset from 2000 to 2018. Sci Data 9:176. 10.1038/s41597-022-01284-8

Musiani M, Okarma H, Jedrzejewski W (1998) Speed and actual distances travelled by radiocollared wolves in Bialowieza Primeval Forest (Poland). Acta Theriologica 43(4):409–416.

Mysłajek RW, Tracz M, Tracz M, Tomczak P, Szewczyk M, Niedźwiecka N, Nowak S (2018) Spatial organization in wolves Canis lupus recolonizing north-west Poland: Large territories at low population density. Mammalian Biology 92:37–44. 10.1016/j.mambio.2018.01.006

Noonan MJ, Fleming CH, Akre TS, Drescher-Lehman J, Gurarie E, Harrison A.-L, Kays R, Calabrese JM (2019) Scale-insensitive estimation of speed and distance traveled from animal tracking data. Movement Ecology 7(35). 10.1186/s40462-019-0177-1

Okarma H., Jędrzejewski W, Schmidt K, Śnieżko S, Bunevich AN, Jędrzejewska B (1998) Home Ranges of Wolves in Białowieża Primeval Forest, Poland, Compared with Other Eurasian Populations. Journal of Mammalogy 79(3):842–852. 10.2307/1383092

“Organizzazione e realizzazione del conteggio del cervo al bramito nel Parco Nazionale Delle Foreste Casentinesi, Monte Falterona E Campigna_Relazione finale -Anno 2023”.

Pacheco C, Rio-Maior H, Nakamura M, Álvares F, Godinho R (2024) Relatedness-based mate choice and female philopatry: Inbreeding trends of wolf packs in a human-dominated landscape. Heredity 132:211–220. 10.1038/s41437-024-00676-3

Paquet PC, Wierzchowski J, Callaghan C (1996) Effects of human activity on gray wolves in the Bow River valley, Banff National Park, Alberta. in: Green J, Pacas C, Cornwell L, Bayley S (eds). Ecological outlooks project. A cumulative effects assessment and futures outlook of the Banff bow valley. prepared for the Banff Bow valley study. Department of Canadian Heritage, Ottawa, on. Chapter 7, pp 74–120.

Pereira HM, Navarro LM (2015) Rewilding European Landscapes. Springer Nature. 10.1007/978-3-319-12039-3

Peterson RO, Woolington JD, Bailey TN (1984) Wolves of the Kenai peninsula, Alaska. Wildlife Monographs 88:3–52. https://www.jstor.org/stable/3830728

Pletscher DH, Ream RR, Boyd DK, Fairchild MW, Kunkel KE (1997) Population dynamics of a recolonizing wolf population. The Journal of wildlife management 61(2):459–465. 10.2307/3802604.

Ražen N, Brugnoli A, Castagna C, Groff C, Kaczensky P, Kljun F, Knauer F, Kos I, Krofel M, Luštrik R, Majić A, Rauer G, Righetti D, Potočnik H (2016) Long-distance dispersal connects Dinaric-Balkan and Alpine grey wolf (Canis lupus) populations. European Journal of Wildlife Research 62(1):137–142. 10.1007/s10344-015-0971-z

Rich LN, Mitchell MS, Gude JA, Sime CA (2012) Anthropogenic mortality, intraspecific competition, and prey availability influence territory sizes of wolves in Montana. Journal of Mammalogy 93(3):722–731. 10.1644/11-MAMM-A-079.2

Roffler GH, Gregovich DP (2018) Wolf space use during denning season on Prince of Wales Island, Alaska. Wildlife Biology 2018(1):1–11. 10.2981/wlb.00468

Sanz-Pérez A, Ordiz A, Sand H, Swenson JE, Wabakken P, Wikenros C, Zimmermann B, Akesson M, Milleret C (2018) No place like home? A test of the natal habitat-biased dispersal hypothesis in Scandinavian wolves. Royal Society Open Science 5(12):181379. 10.1098/rsos.181379

Scandura M, Apollonio M, Mattioli L (2001) Recent recovery of the Italian wolf population: a genetic investigation using microsatellites. Mammalian Biology 66:321–331.

Schielzeth H, Forstmeier W (2009) Conclusions beyond support: Overconfident estimates in mixed models. Behavioral Ecology 20(2):416–420. 10.1093/beheco/arn145

Schielzeth H (2010) Simple means to improve the interpretability of regression coefficients. Methods in Ecology and Evolution 1(2):103–113. 10.1111/j.2041-210X.2010.00012.x

Stolwijk AM, Straatman H, Zielhuis GA (1999) Studying seasonality by using sine and cosine functions in regression analysis. Journal of Epidemiology and Community Health 53(4):235–238. 10.1136/jech.53.4.235

Tinbergen N (1957) The Functions of Territory. Bird Study 4(1):14–27.

Vorel A, Kadlec I, Toulec T, Selimovic A, Horníček A, Vojtěch O, Mokrý J, Pavlačík L, Arnold W, Cornils J, Kutal M, Dula M, Žák L, Barták V (2024) Home range and habitat selection of wolves recolonising central European human-dominated landscapes. Wildlife Biology e01245. 10.1002/wlb3.01245

Zanni M, Brogi R, Merli E, Apollonio M (2023) The wolf and the city: Insights on wolves’ conservation in the anthropocene. Animal Conservation 26(6):766–780. 10.1111/acv.12858

Zub K, Theuerkauf J, Jędrzejewski W, Jędrzejewska B, Schmidt K, Kowalczyk R (2003) Wolf pack territory marking in the Białowieża Primeval Forest (Poland). Behaviour, 635–648. https://www.jstor.org/stable/4536049

Zuur AF, Ieno EN, Walker N, Saveliev AA, Smith GM (2009) Mixed effects models and extensions in ecology with R. Springer New York. 10.1007/978-0-387-87458-6

