## Supplementary material for "Extreme spatial plasticity in Italian wolves: ecological determinants of range size and movement patterns across heterogeneous landscapes": S1

**Table S1** Wild ungulate densities and total biomass across the five study areas and within the home ranges of the 13 GPS-tracked wolf packs in Italy

| Study area | Pack | Ungulate species | Density (heads/km^2^) | Individual body mass (kg) | Biomass density (kg/km^2^) | Total prey biomass availability (kg/km^2^) |
| --- | --- | --- | --- | --- | --- | --- |
| FCNP | FCNP1; FCNP2; FCNP3 | Red deer | 7.25 | 100 | 725 | 1,449 |
|  |  | Roe deer | 1.86 | 20 | 37 |  |
|  |  | Fallow deer | 5.23 | 50 | 262 |  |
|  |  | Wild boar | 14.18 | 30 | 425 |  |
| SRE | SRE1; SRE2 | Fallow deer | 40.1 | 50 | 2,005 | 2,167 |
|  |  | Wild boar | 5.4 | 30 | 162 |  |
| PEC | PEC | Red deer | 0.20 | 100 | 20 | 1,605 |
|  |  | Roe deer | 9.9 | 20 | 198 |  |
|  |  | Fallow deer | 6.8 | 50 | 340 |  |
|  |  | Wild boar | 34.9 | 30 | 1,047 |  |
| Pre-Alps | Asiago Plateau | Red deer | 0.8 | 100 | 80 | 210 |
|  |  | Roe deer | 1.79 | 20 | 36 |  |
|  |  | Chamois | 1.33 | 25 | 33 |  |
|  |  | Mouflon | 2.03 | 30 | 61 |  |
|  | Grappa1 | Red deer | 0.92 | 100 | 92 | 256 |
|  |  | Roe deer | 4.66 | 20 | 93 |  |
|  |  | Chamois | 2.33 | 25 | 58 |  |
|  |  | Mouflon | 0.44 | 30 | 13 |  |
|  | Grappa Monfenera | Red deer | 1.04 | 100 | 104 | 210 |
|  |  | Roe deer | 3.18 | 20 | 64 |  |
|  |  | Chamois | 1.05 | 25 | 26 |  |
|  |  | Mouflon | 0.54 | 30 | 16 |  |
|  | Grappa Piave | Red deer | 1.6 | 100 | 160 | 286 |
|  |  | Roe deer | 4.67 | 20 | 93 |  |
|  |  | Chamois | 0.46 | 25 | 12 |  |
|  |  | Mouflon | 0.69 | 30 | 21 |  |
| Alps | DBNP1; DBNP2 | Red deer | 4.79 | 100 | 479 | 808 |
|  |  | Chamois | 13.14 | 25 | 329 |  |
|  | Dubiea | Red deer | 2.91 | 100 | 291 | 467 |
|  |  | Roe deer | 2.66 | 20 | 53 |  |
|  |  | Chamois | 4.2 | 25 | 105 |  |
|  |  | Mouflon | 0.61 | 30 | 18 |  |

FCNP: Foreste Casentinesi National Park; SRE: San Rossore Estate; PEC: Presidential Estate of Castelporziano; DBPN: Dolomiti Bellunesi National Park; See section 2.2 for more details about the methods used for the population density estimation, the average individual body mass considered, and the calculation of total prey biomass availability.

**Table S2** Likelihood ratio tests comparing the full model with reduced models explaining the variation of wolf monthly range size.

| Comparison | Reduced model | X^2^ | df | P |
| --- | --- | --- | --- | --- |
| Full vs Null | HR ~ 1 \| id | 168.50 | 12 | < 0.0001 |
| Full vs area-reduced | HR ~ HF + sex : social status + sex + social status + month + social status : month \| id | 12.15 | 4 | 0.02 |
| Full vs sex*status interaction-reduced | HR ~ HF + area + sex + social status + month + social status : month \| id | 3.96 | 1 | 0.046 |
| Full vs month*status interaction-reduced | HR ~ HF + area + sex:social status + sex + social status + month \| id | 1.71 | 2 | 0.42 |

Null: intercept-only model; HR: monthly range size; HF: Human Footprint Index; X^2^: likelihood ratio test statistic; df: degrees of freedom; P: p-value

**Table S3** Likelihood ratio tests comparing the full model with reduced models explaining the variation of wolf monthly distance travelled.

| Comparison | Reduced model | X^2^ | df | P |
| --- | --- | --- | --- | --- |
| Full vs Null | DT ~ 1 \| id | 32.20 | 12 | < 0.01 |
| Full vs area-reduced | DT ~ HF + sex : social status + sex + social status + month + social status : month \| id | 18.28 | 4 | < 0.01 |
| Full vs sex*status interaction-reduced | DT ~ HF + area + sex + social status + month + social status : month \| id | 0.31 | 1 | 0.57 |
| Full vs month*status interaction-reduced | DT ~ HF + area + sex:social status + sex + social status + month \| id | 1.63 | 2 | 0.44 |

Null: intercept-only model; DT: monthly distance travelled; HF: Human Footprint Index; X^2^: likelihood ratio test statistic; df: degrees of freedom; P: p-value

**Table S4** GPS-tracked wolves’ seasonal ranges estimated by means of Autocorrelation Kernel Density Estimation (AKDE) and Minimum Convex Polygon (MCP); and seasonal distance travelled estimated by means of Continuous-Time Speed and Distance (CTSD) and Straight-Line Displacement (SLD)

| ID | Area | Sex | Status | Season | AKDE hrs (km^2^) 95% | AKDE hrs CI | MCP hrs (km^2^) 95% | CTSD dt (km/day) | CTSD dt CI | SLD dt (km/day) |
| --- | --- | --- | --- | --- | --- | --- | --- | --- | --- | --- |
| WF2001 | Pre-Alps | f | R | win | 158.70 | 131.59-188.31 | 141.6 | 31.65 | 31.21-32.10 | 7.50 |
| WF2002 | Pre-Alps | f | F | sum | 420.71 | 303.06-557.35 | 330.9 | 29.82 | 29.30-30.33 | 7.12 |
| WM2101 | Pre-Alps | m | R | sum | 91.08 | 81.13-101.59 | 97.98 | 36.27 | 35.73-36.81 | 6.46 |
| WF2101 | Pre-Alps | f | R | sum | 80.92 | 70.00-92.63 | 60.05 | 32.08 | 31.31-32.86 | 4.88 |
| WF2202 | Alps | f | R | win | 355.17 | 254.20-  472.75 | 226.7 | 27.26 | 26.85-27.67 | 7.42 |
|  |  |  |  | sum | 133.82 | 114.05-155.15 | 136 | 29.42 | 28.92-29.93 | 6.88 |
|  |  |  |  | win | 125.71 | 102.40-151.39 | 106.5 | 23.35 | 23.01-23.69 | 6.98 |
| WM2201 | FCNP | m | R | win | 18.28 | 16.82-19.80 | 20.48 | 24.25 | 23.68-24.84 | 4.52 |
|  |  |  |  | sum | 22.25 | 20.37-24.21 | 20.88 | 22.78 | 22.34-23.23 | 4.54 |
| WF2203 | FCNP | f | R | sum | 13.50 | 12.47-14.57 | 11.66 | 56.25 | 50.57-62.08 | 2.96 |
| WM2303 | SRE | m | R | sum | 52.29 | 48.22-56.52 | 48.45 | 37.20 | 36.73-37.66 | 8.75 |
| WM2304 | SRE | m | F | sum | 242.42 | 177.40-317.44 | 185.8 | 21.94 | 21.41-22.48 | 5.09 |
| WM2305 | FCNP | m | R | win | 58.51 | 52.13-65.25 | 54.54 | 27.94 | 27.63-28.25 | 6.90 |
|  |  |  |  | sum | 93.86 | 78.55-110.51 | 76.71 | 27.43 | 26.85-28.02 | 5.74 |
| WF2401 | SRE | f | R | sum | 47.19 | 42.70-51.90 | 41.75 | 56.68 | 56.00-57.35 | 6.41 |
|  |  |  |  | win | 63.29 | 57.07-69.82 | 58.42 | 45.96 | 45.11-46.82 | 9.11 |
| WM2401 | SRE | m | R | sum | 44.67 | 40.92-48.59 | 37.86 | 50.84 | 49.97-51.70 | 5.85 |
|  |  |  |  | win | 92.68 | 82.43-103.53 | 78.23 | 45.33 | 44.78-45.88 | 8.94 |
| WM2402 | PEC | m | R | sum | 67.10 | 59.78-74.83 | 63.87 | 32.10 | 31.59-32.63 | 7.85 |
| WF2402 | Alps | f | F | win | 710.63 | 490.30-971.16 | 633.5 | 26.04 | 25.70-26.39 | 7.91 |
| WF2403 | Alps | f | R | win | 87.93 | 69.19-108.88 | 70.2 | 20.06 | 19.79-20.34 | 4.70 |

FCNP: Foreste Casentinesi National Park; SRE: San Rossore Estate; PEC: Presidential Estate of Castelporziano; CI: 95% Confidence Intervals; hrs: home range size; dt: distance travelled; f: female; m: male; R: pack-member individual; F: floater individual; win: winter; sum: summer.

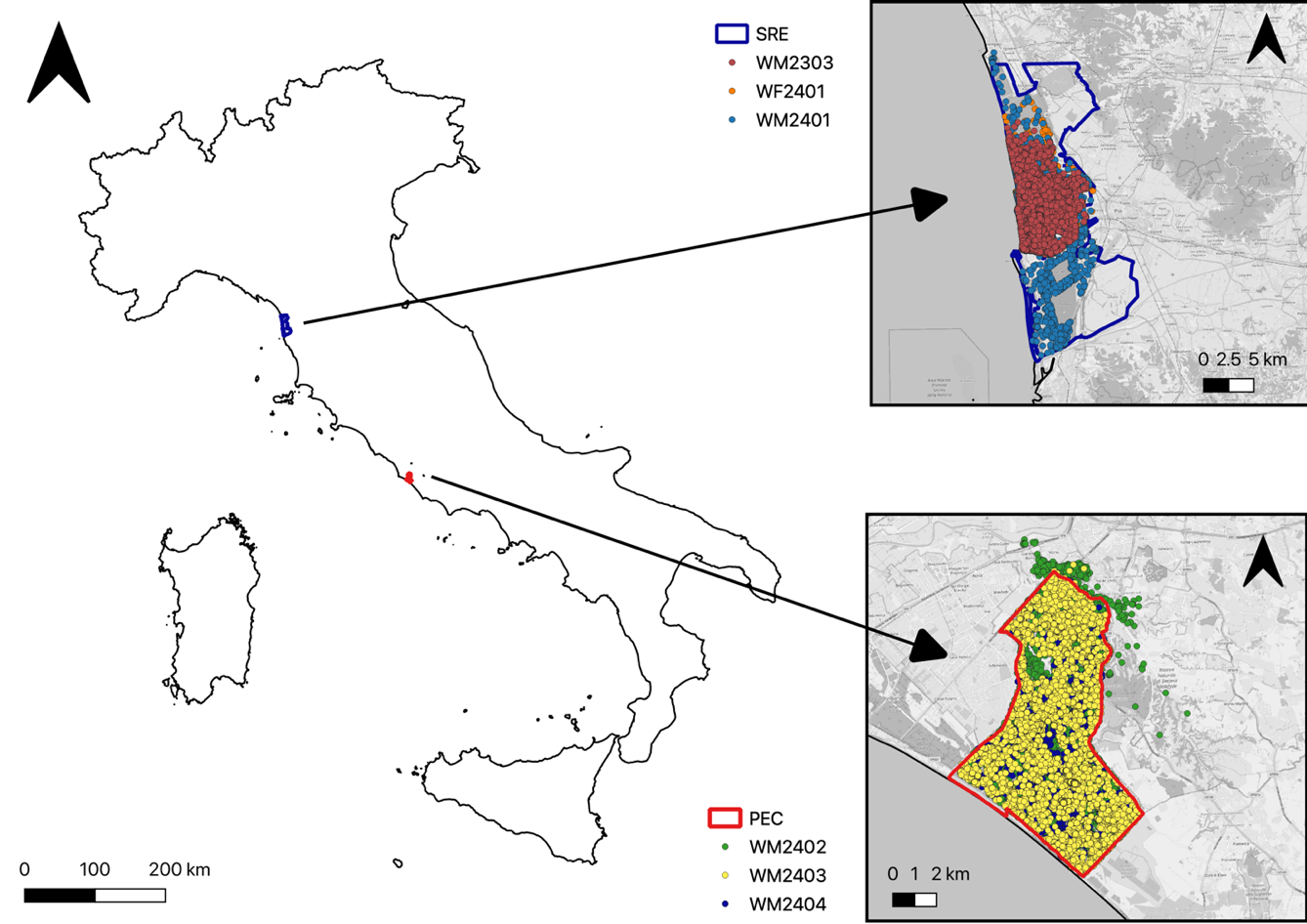

**Fig. S4** Spatial distribution of wolf packs inhabiting the San Rossore Estate (SRE) and the Presidential Estate of Castelporziano (PEC). GPS locations show that pack activity was concentrated almost entirely within the boundaries of these partially fenced, protected areas.

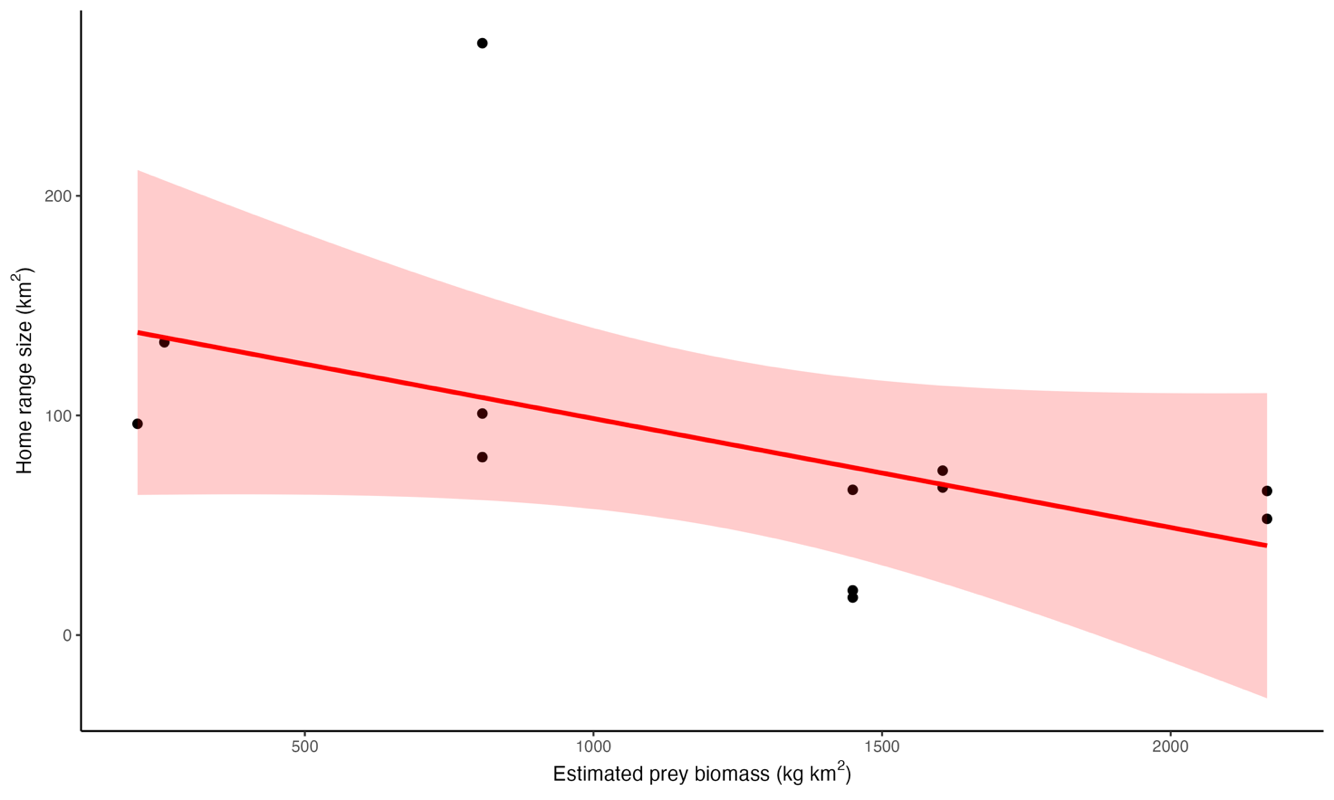

**Fig. S5** Relationship between annual resident wolf home range size (km^2^) and estimated prey biomass (kg/km^2^). Each point represents paired annual values of home range size and prey biomass; the solid line shows the linear regression and the shaded area indicates the 95% confidence interval.

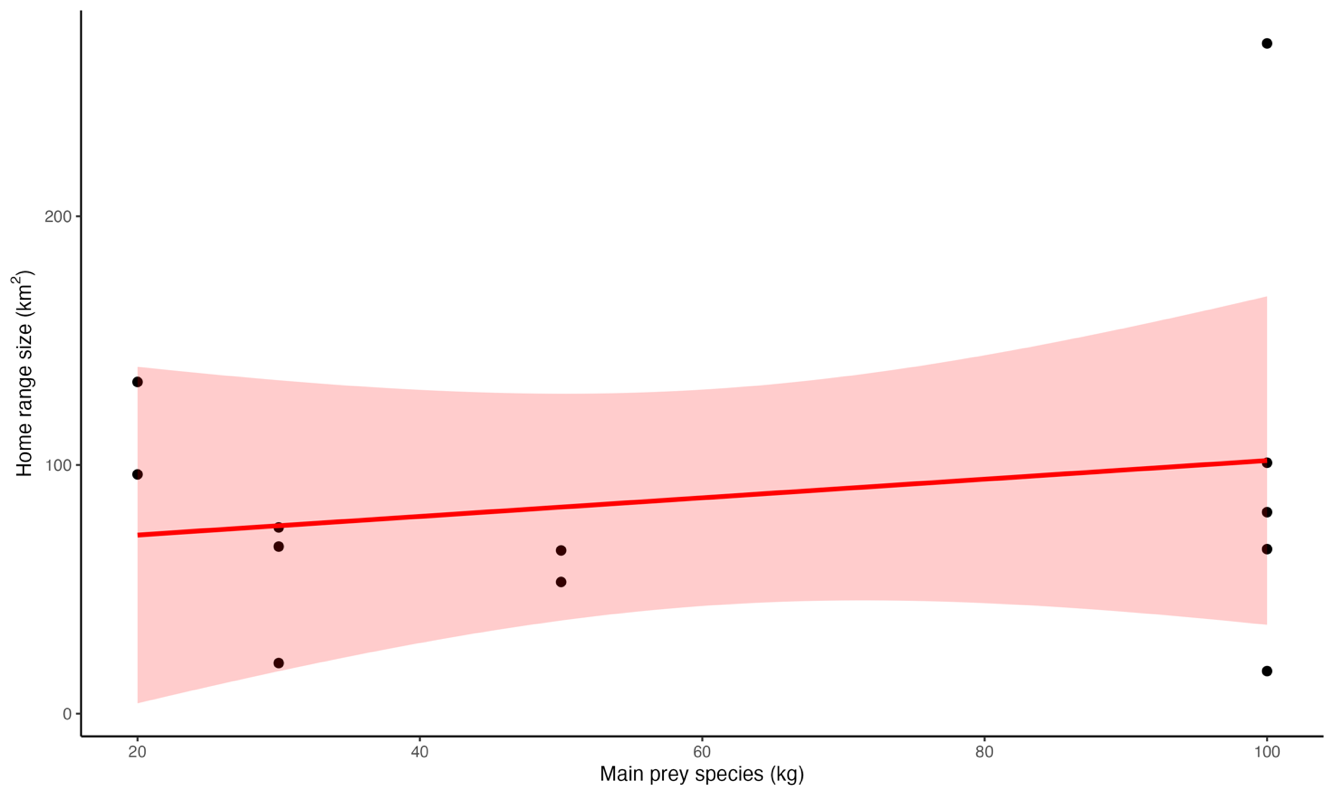

**Fig. S6** Relationship between annual resident wolf home range size (km^2^) and the mean body mass of the main prey species (kg). Each point represents paired annual values of home range size and prey body mass; the solid line shows the linear regression and the shaded area indicates the 95% confidence interval.
