## Supplementary material for "Extreme spatial plasticity in Italian wolves: ecological determinants of range size and movement patterns across heterogeneous landscapes": S2

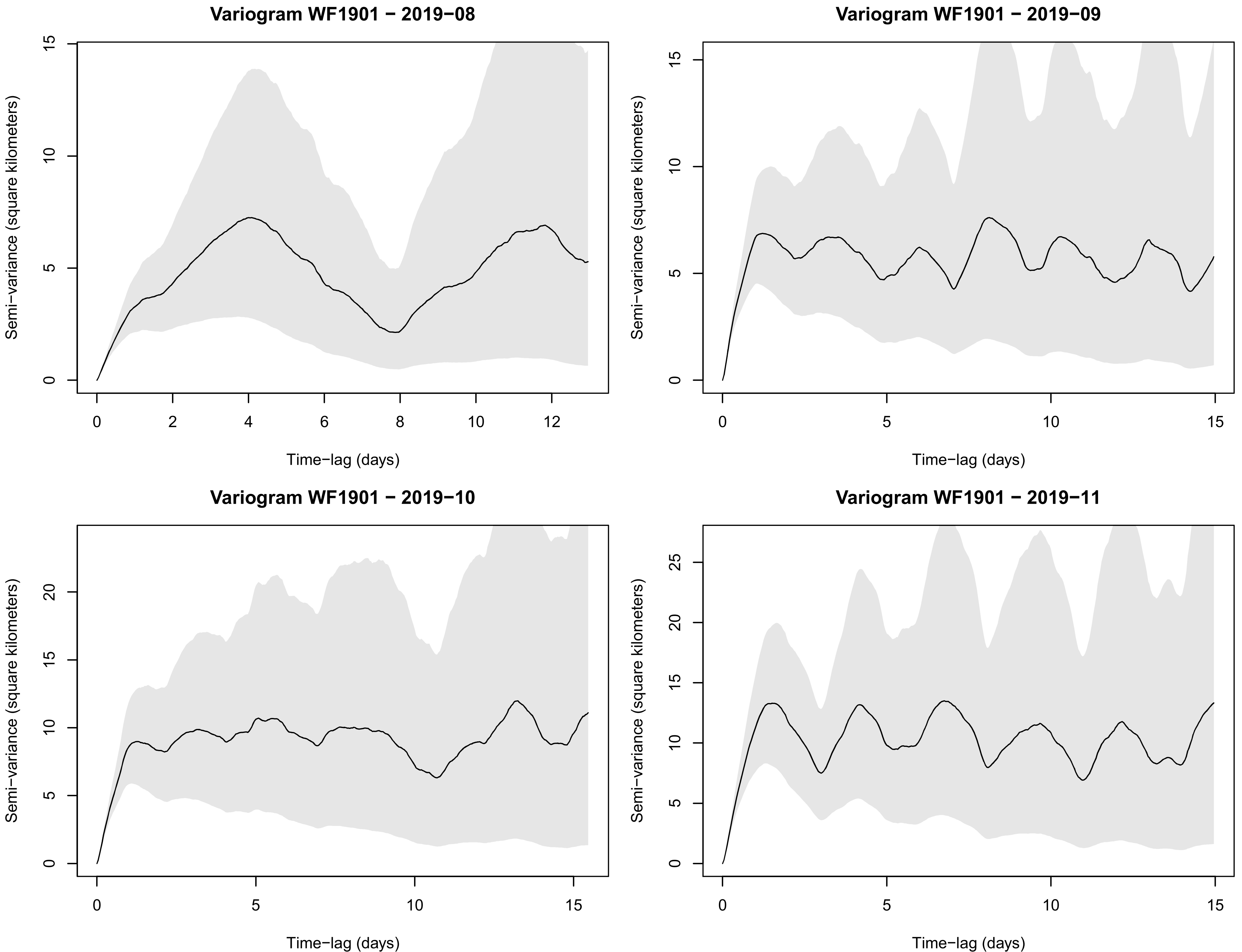
Variogram WF1901 − 2019−08
Semi−variance (square kilometers)
Time−lag (days)
0
5
10
15
0
2
4
6
8
10
12
Variogram WF1901 − 2019−09
Semi−variance (square kilometers)
Time−lag (days)
0
5
10
15
0
5
10
15
Variogram WF1901 − 2019−10
Semi−variance (square kilometers)
Time−lag (days)
0
5
10
15
20
0
5
10
15
Variogram WF1901 − 2019−11
Semi−variance (square kilometers)
Time−lag (days)
0
5
10
15
20
25
0
5
10
15

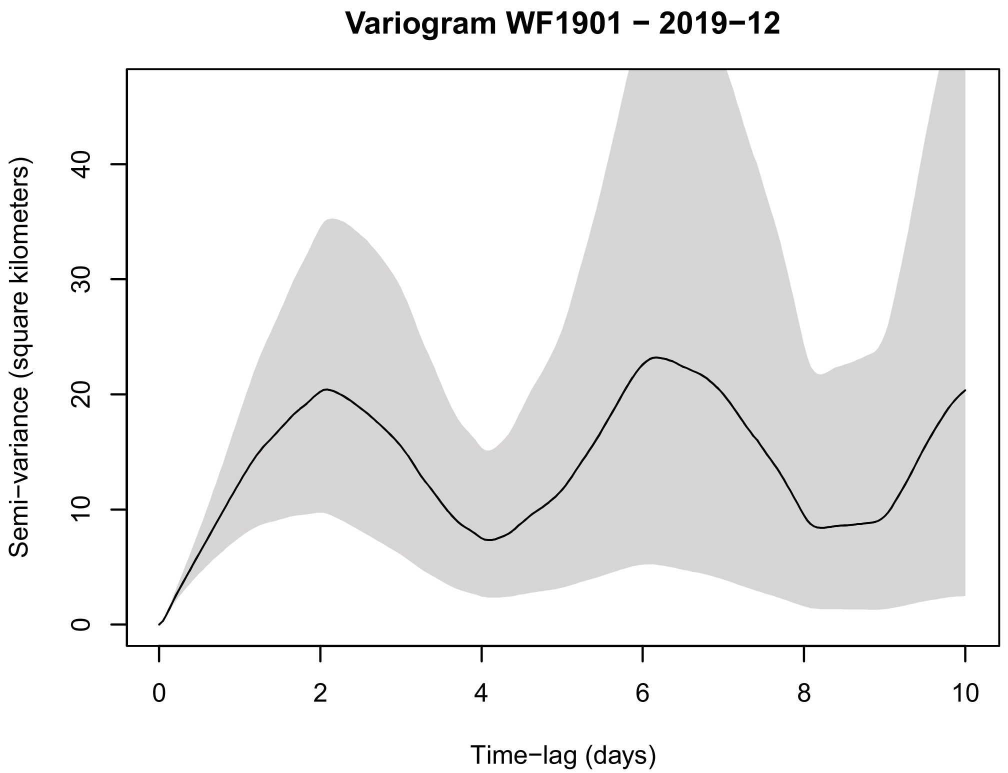
Variogram WF1901 − 2019−12
Semi−variance (square kilometers)
Time−lag (days)

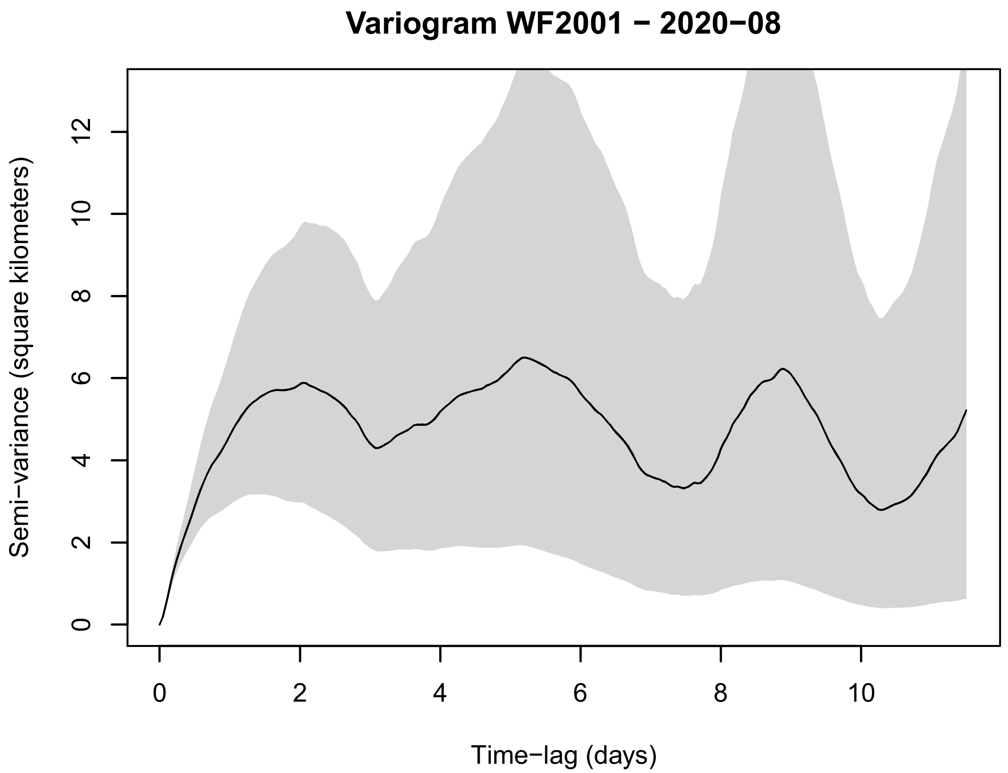
Variogram WF2001 − 2020−08
Semi−variance (square kilometers)
Time−lag (days)

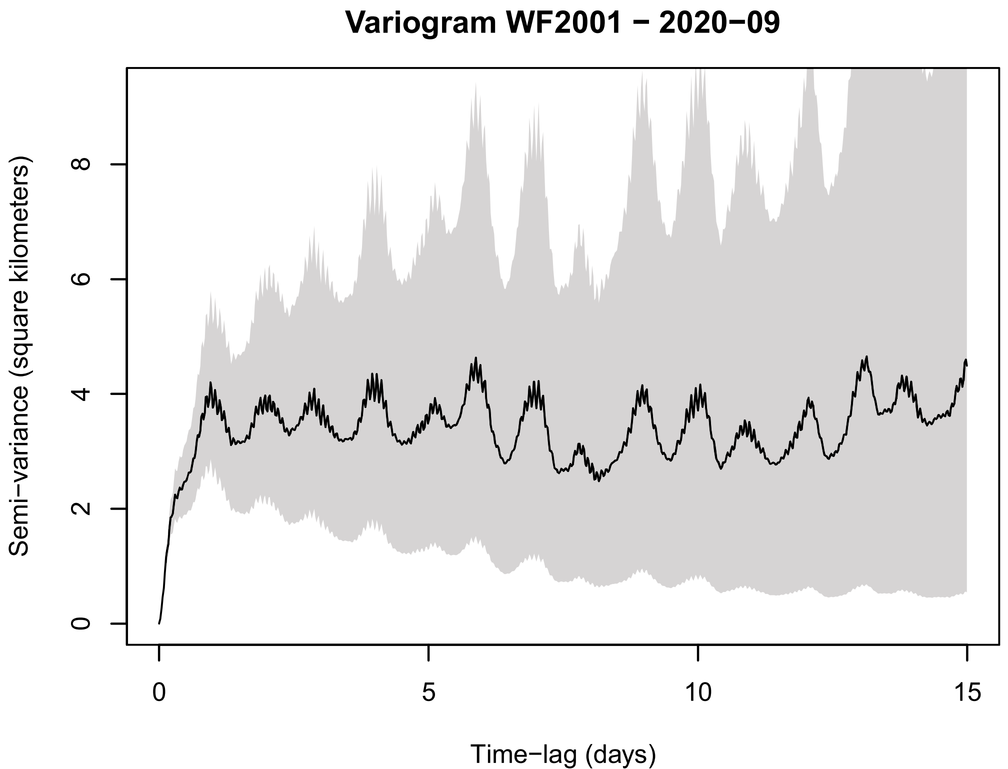
Variogram WF2001 − 2020−09
Semi−variance (square kilometers)
Time−lag (days)

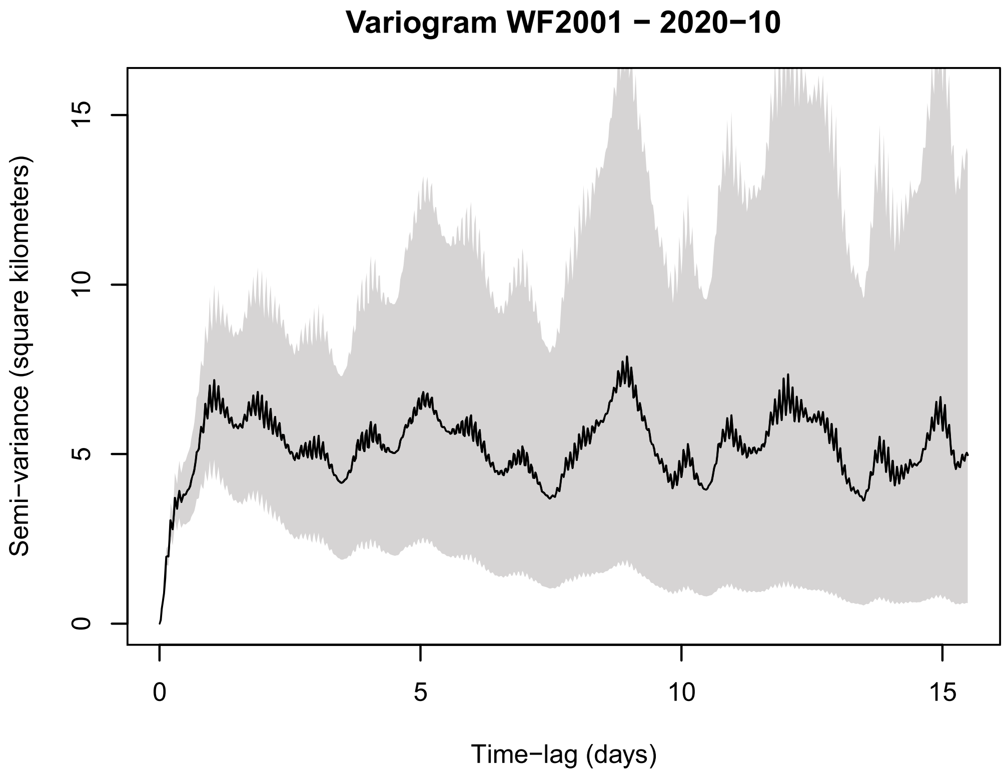
Variogram WF2001 − 2020−10
Semi−variance (square kilometers)
Time−lag (days)

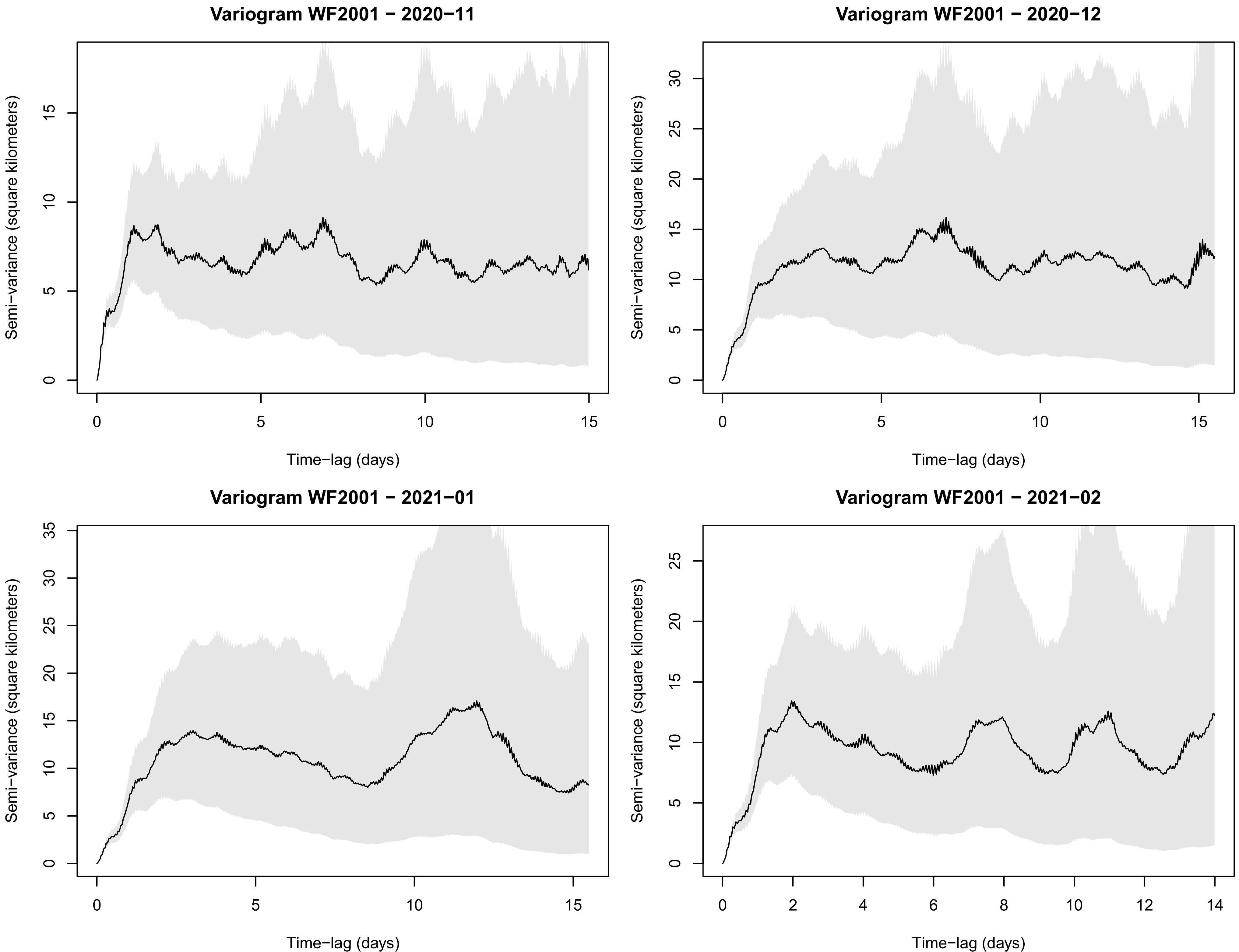

Variogram WF2001 − 2020−11
Variogram WF2001 − 2020−12
Variogram WF2001 − 2021−01
Variogram WF2001 − 2021−02
Semi−variance (square kilometers)
Time−lag (days)

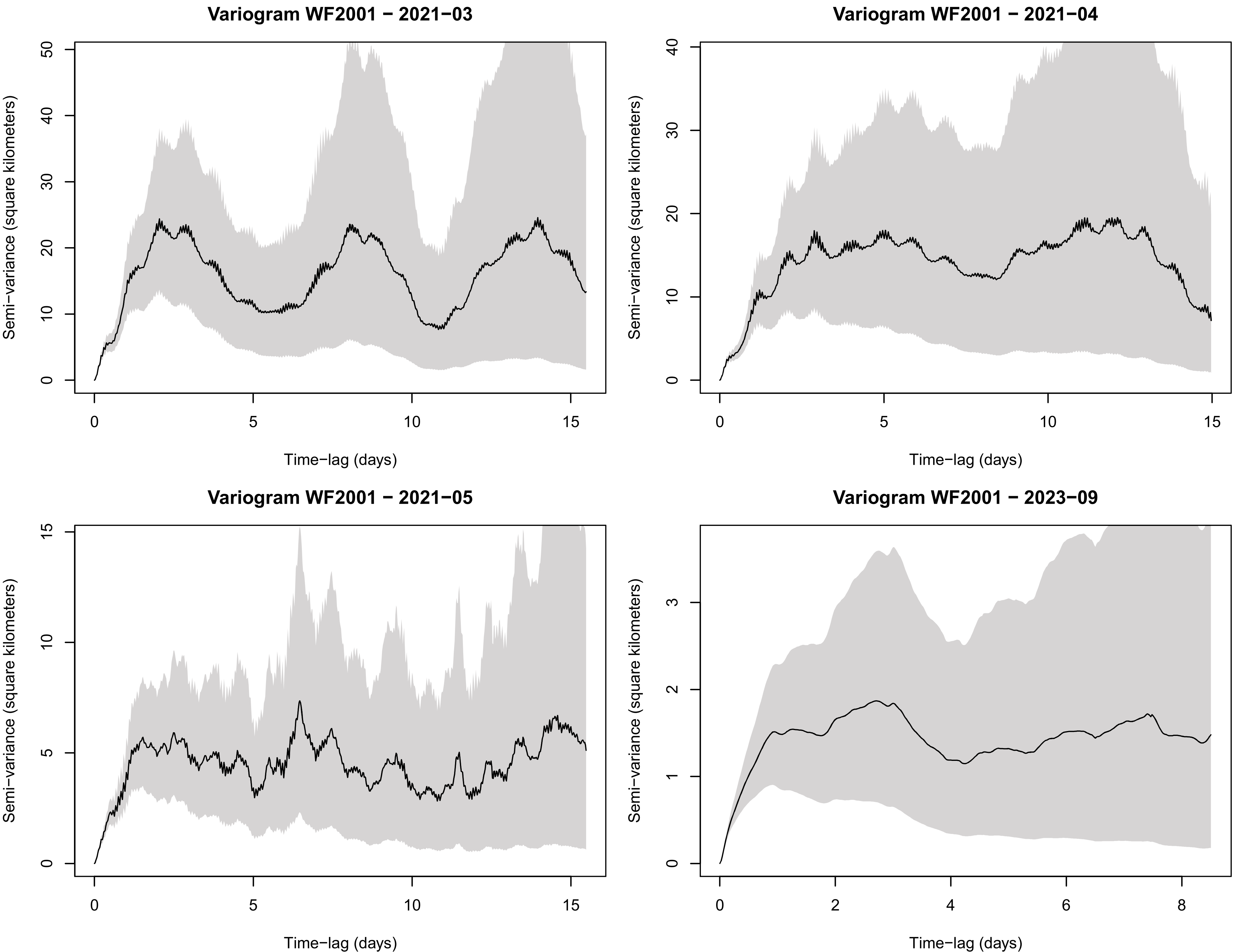
Variogram WF2001 − 2021−03
Variogram WF2001 − 2021−04
Variogram WF2001 − 2021−05
Variogram WF2001 − 2023−09
Semi−variance (square kilometers)
Time−lag (days)

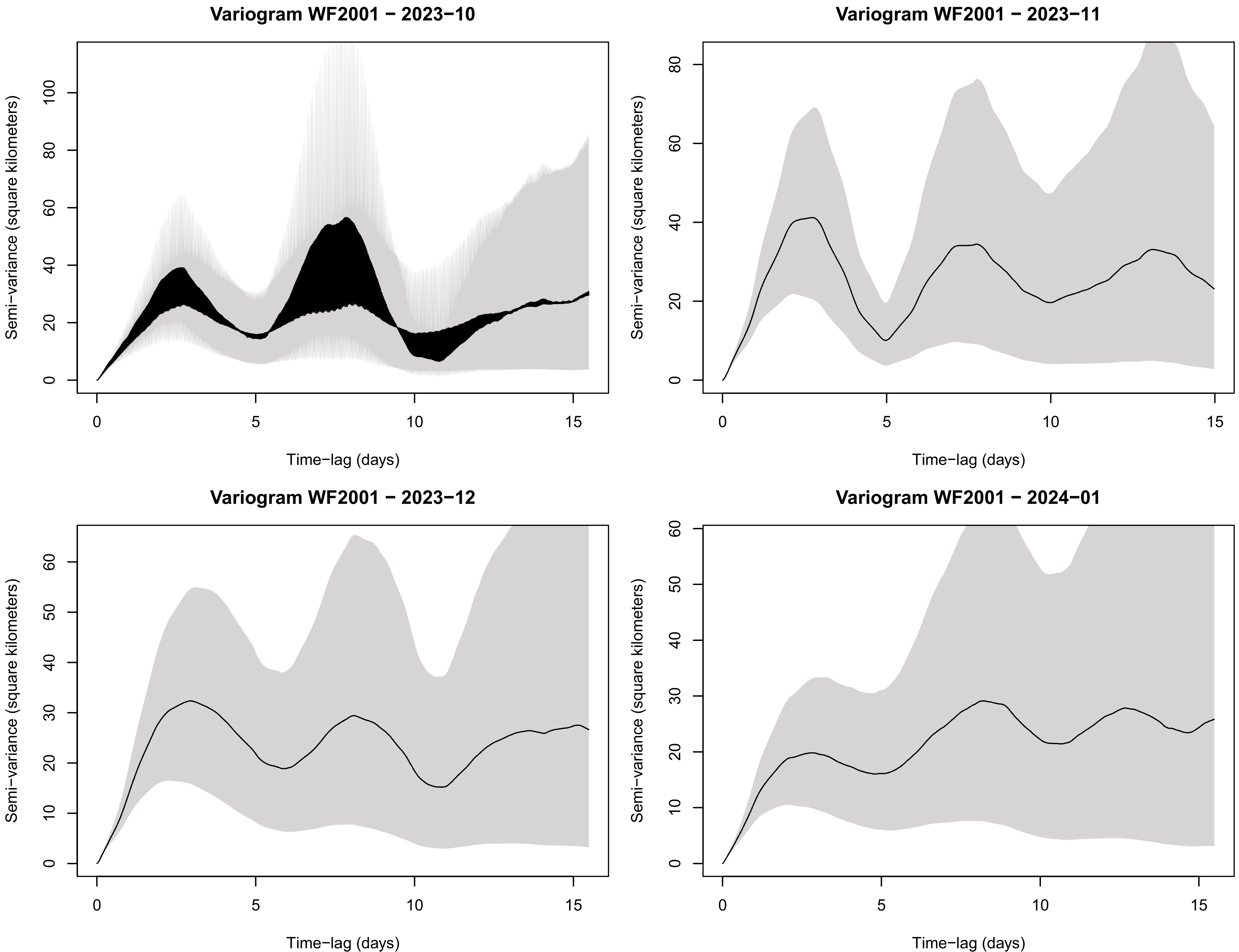

Variogram WF2001 − 2023−10
Variogram WF2001 − 2023−11
Variogram WF2001 − 2023−12
Variogram WF2001 − 2024−01
Semi−variance (square kilometers)
Time−lag (days)

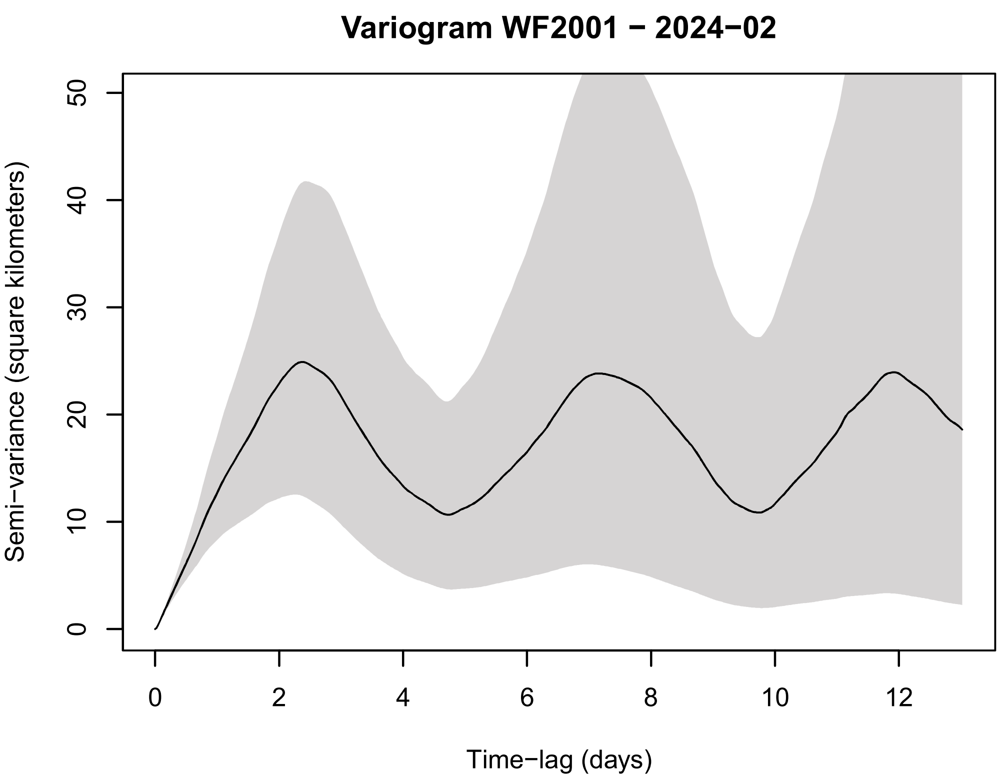
Variogram WF2001 − 2024−02
Semi−variance (square kilometers)
0
10
20
30
40
50
0
2
4
6
8
10
12
Time−lag (days)

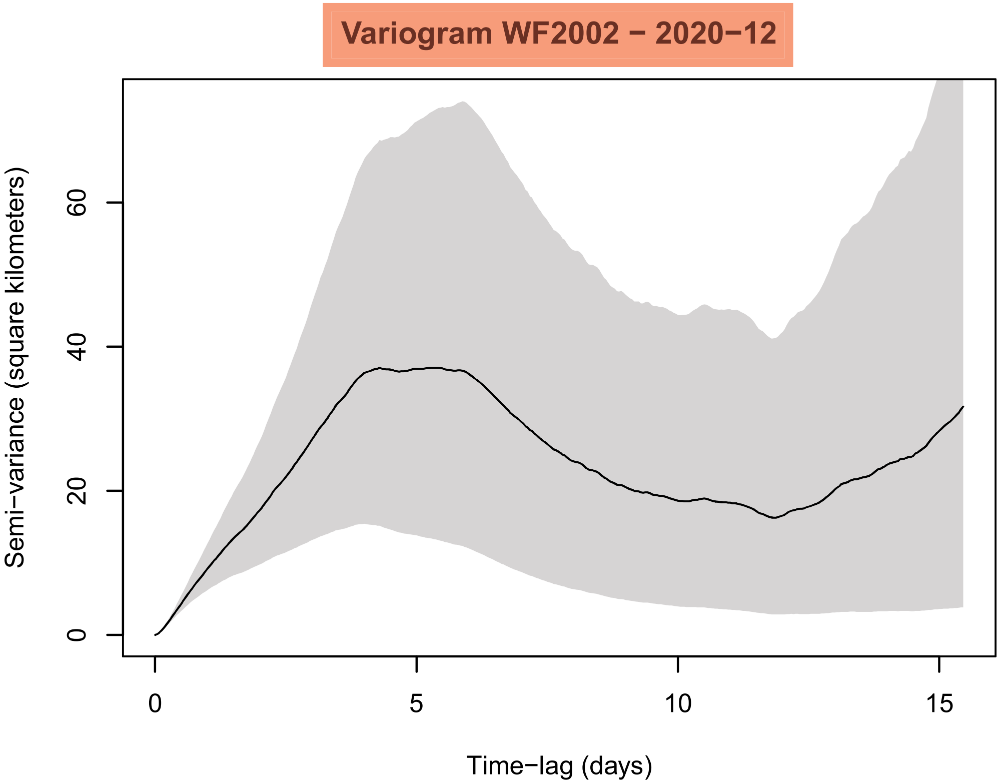
Variogram WF2002 − 2020−12
Semi−variance (square kilometers)
0
20
40
60
0
5
10
15
Time−lag (days)

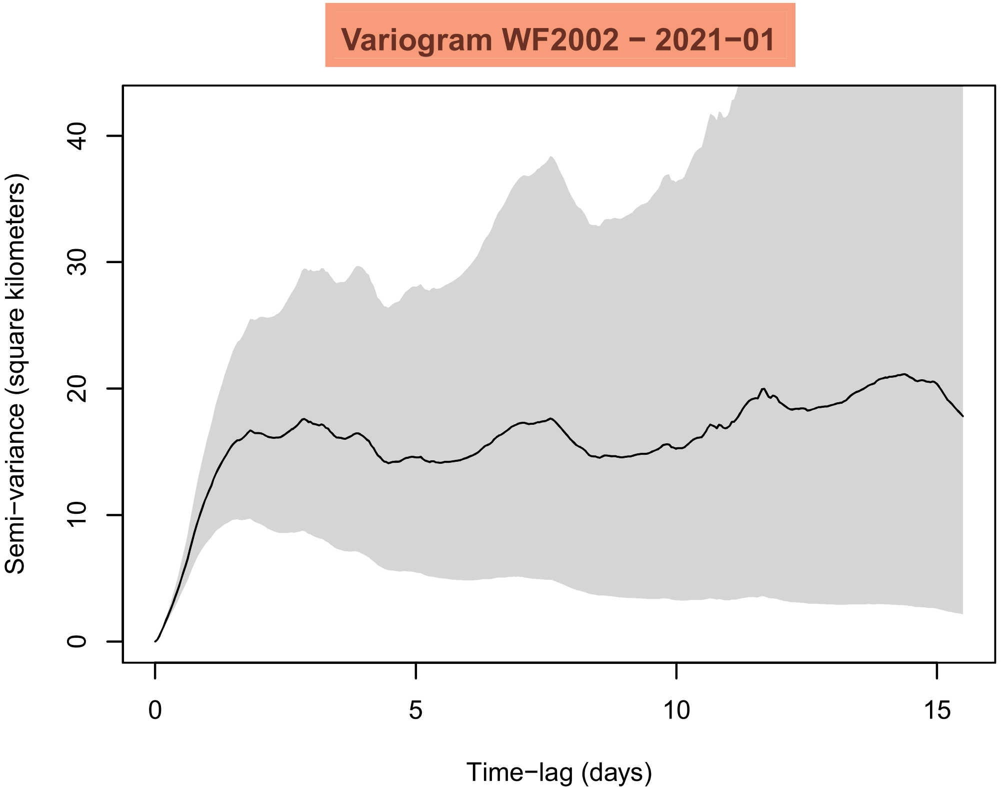
Variogram WF2002 − 2021−01
Semi−variance (square kilometers)
0
10
20
30
40
0
5
10
15
Time−lag (days)

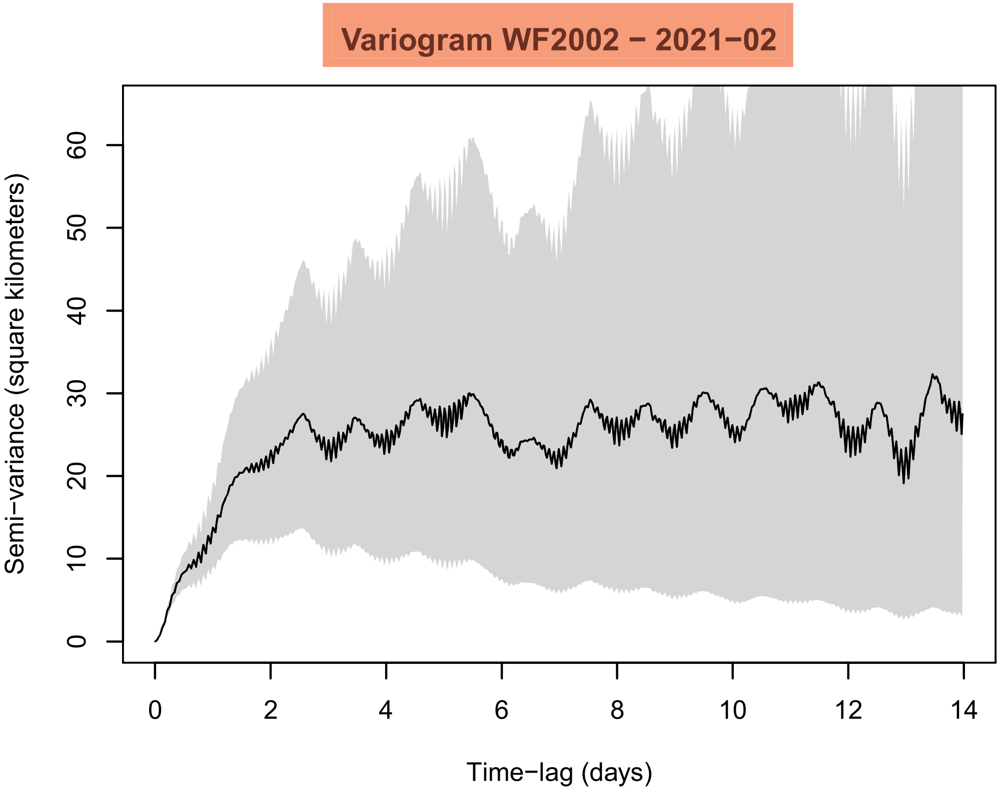
Variogram WF2002 − 2021−02
Semi−variance (square kilometers)
0
10
20
30
40
50
60
0
2
4
6
8
10
12
14
Time−lag (days)

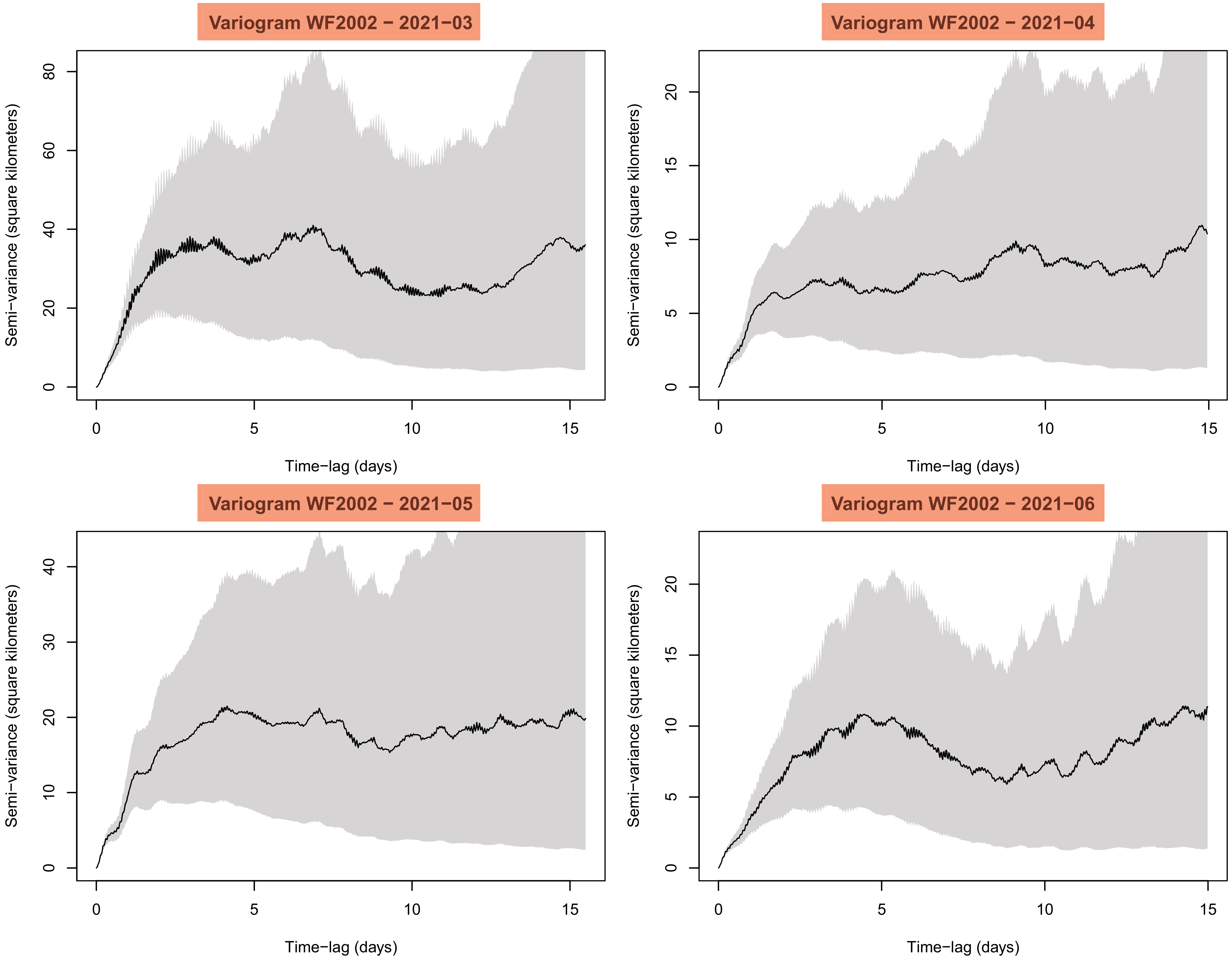
Variogram WF2002 − 2021−03
Variogram WF2002 − 2021−04
Variogram WF2002 − 2021−05
Variogram WF2002 − 2021−06
Semi−variance (square kilometers)
Time−lag (days)
0
5
10
15
0
20
40
60
80
0
5
10
15
20
0
10
20
30
40

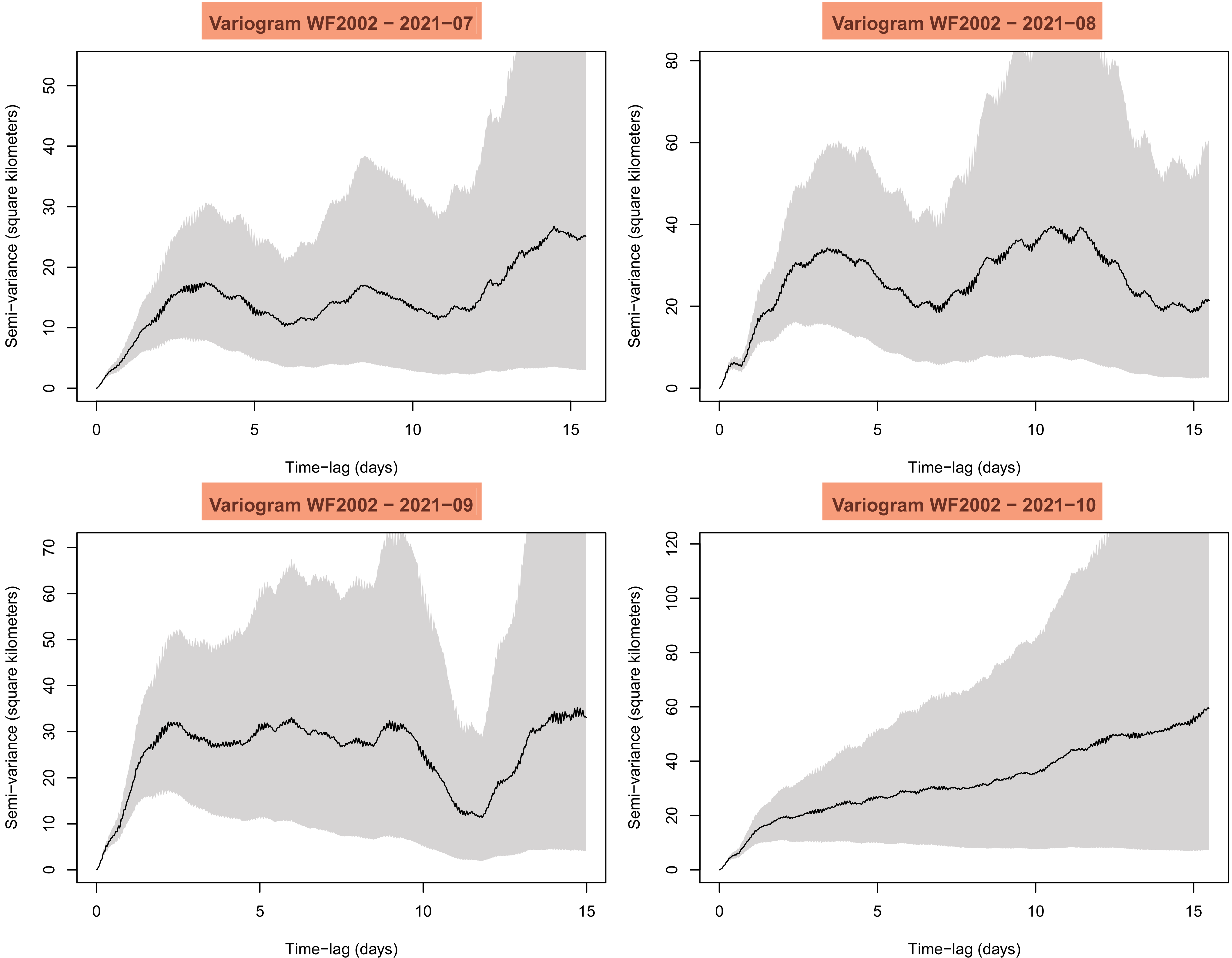
Variogram WF2002 − 2021−07
Variogram WF2002 − 2021−08
Variogram WF2002 − 2021−09
Variogram WF2002 − 2021−10
Semi−variance (square kilometers)
Time−lag (days)
0
5
10
15
20
30
40
50
60
70
80
100
120

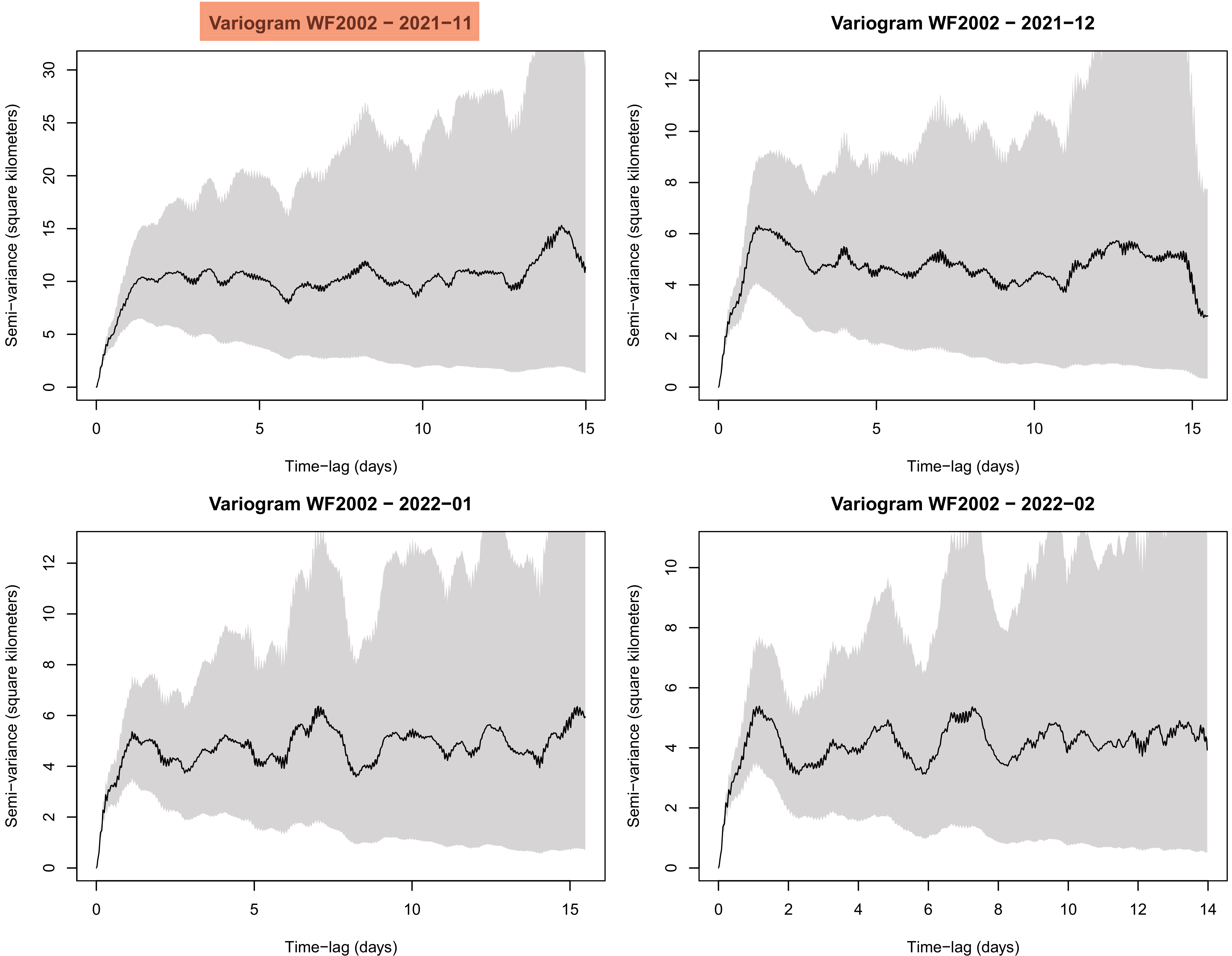

Variogram WF2002 − 2021−11
Variogram WF2002 − 2021−12
Variogram WF2002 − 2022−01
Variogram WF2002 − 2022−02
Semi−variance (square kilometers)
Time−lag (days)

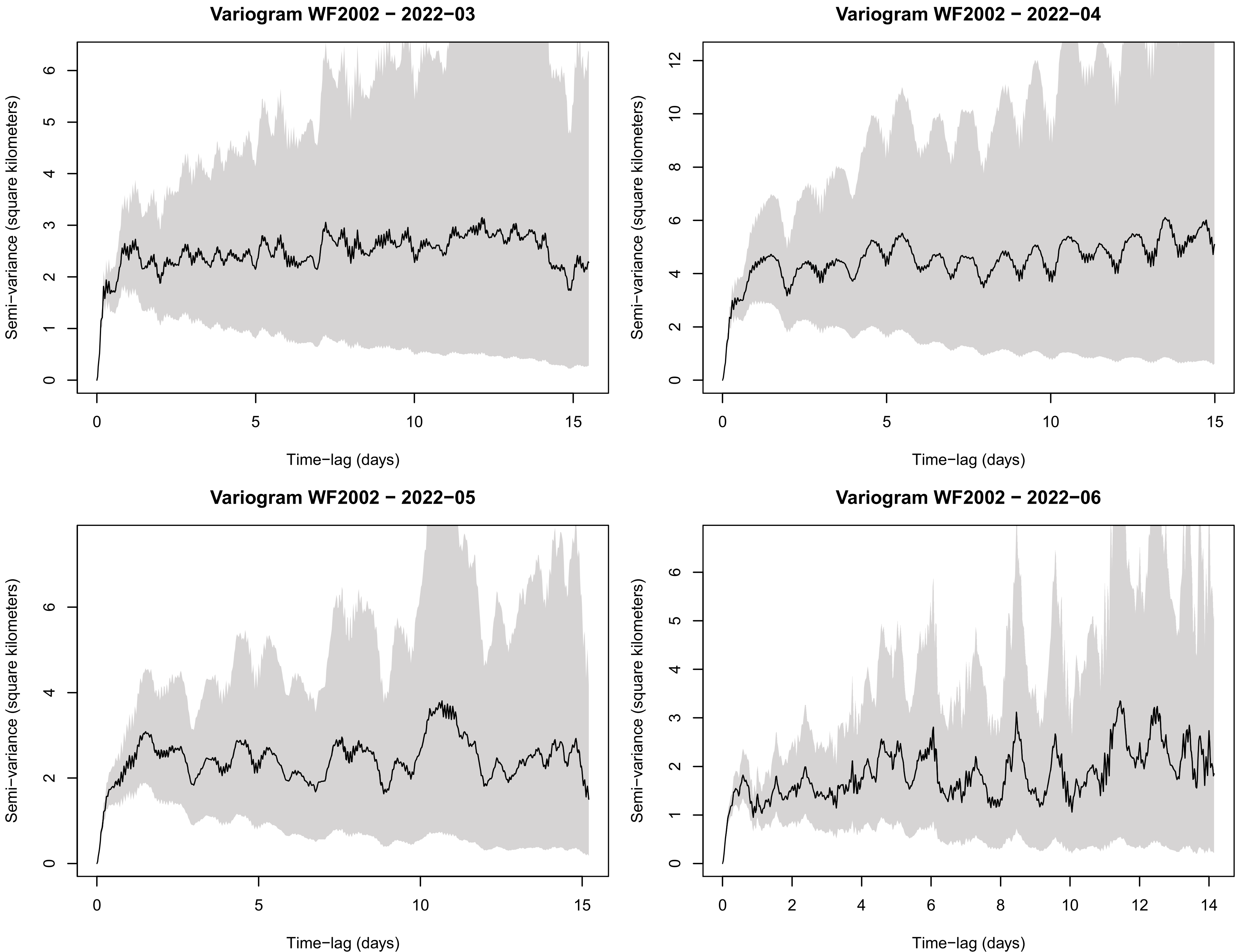
Variogram WF2002 − 2022−03
Variogram WF2002 − 2022−04
Variogram WF2002 − 2022−05
Variogram WF2002 − 2022−06
Semi−variance (square kilometers)
Time−lag (days)

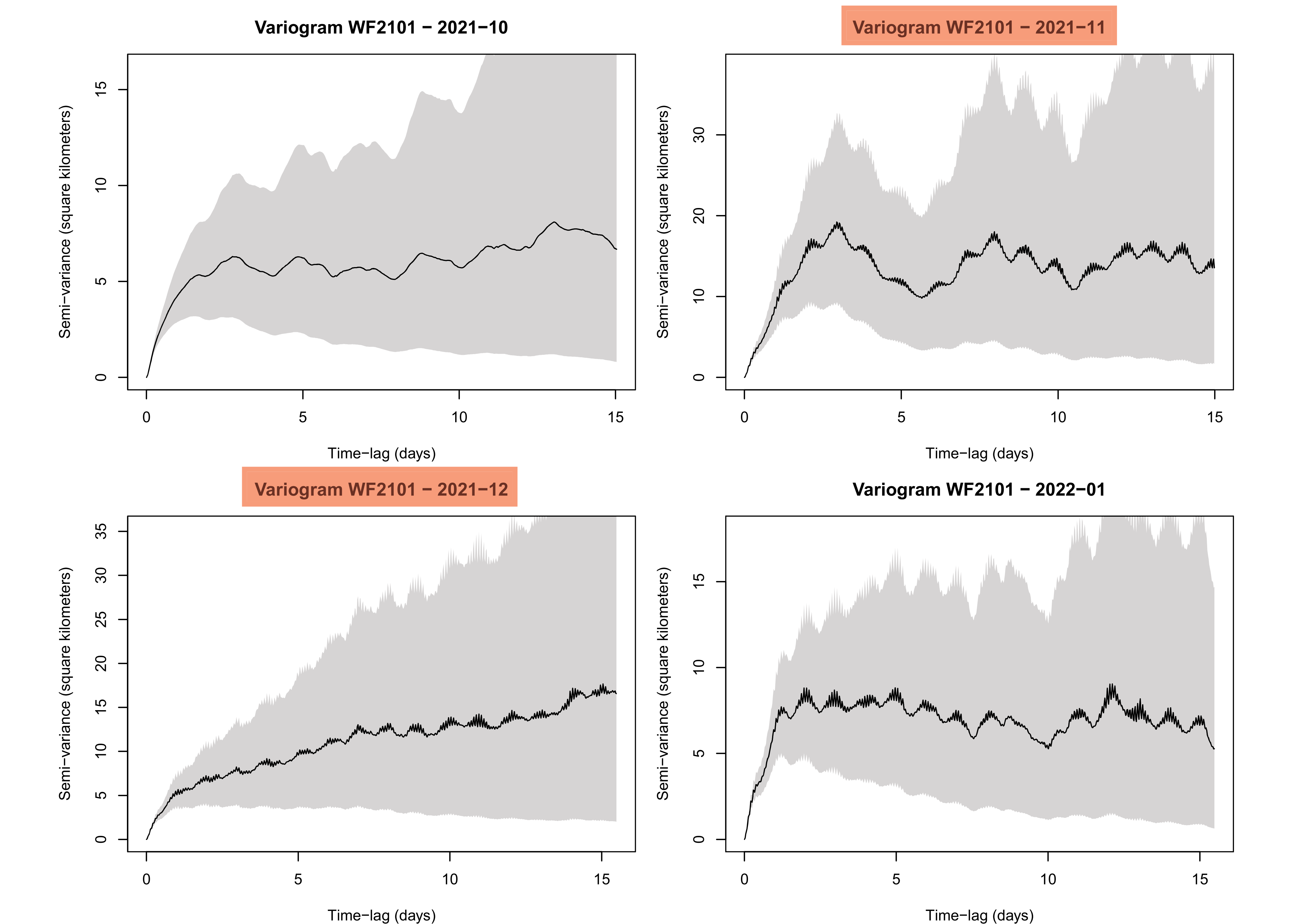

Variogram WF2101 − 2021−10
Variogram WF2101 − 2021−11
Variogram WF2101 − 2021−12
Variogram WF2101 − 2022−01
Semi−variance (square kilometers)
Time−lag (days)

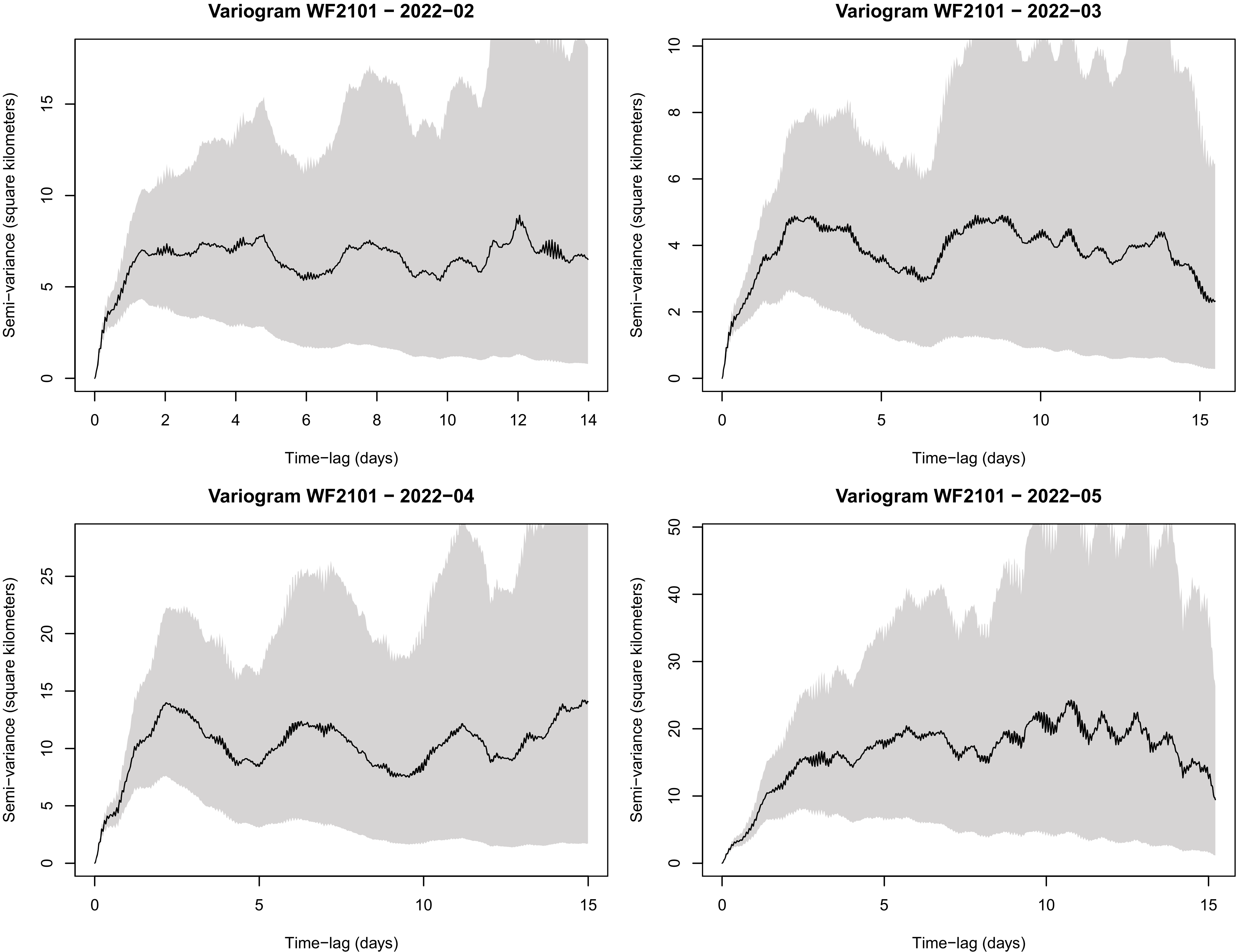
Variogram WF2101 − 2022−02
Semi−variance (square kilometers)
Time−lag (days)
0
5
10
15
0
2
4
6
8
10
12
14
Variogram WF2101 − 2022−03
Semi−variance (square kilometers)
Time−lag (days)
0
2
4
6
8
10
0
5
10
15
Variogram WF2101 − 2022−04
Semi−variance (square kilometers)
Time−lag (days)
0
5
10
15
20
25
0
5
10
15
Variogram WF2101 − 2022−05
Semi−variance (square kilometers)
Time−lag (days)
0
10
20
30
40
50
0
5
10
15

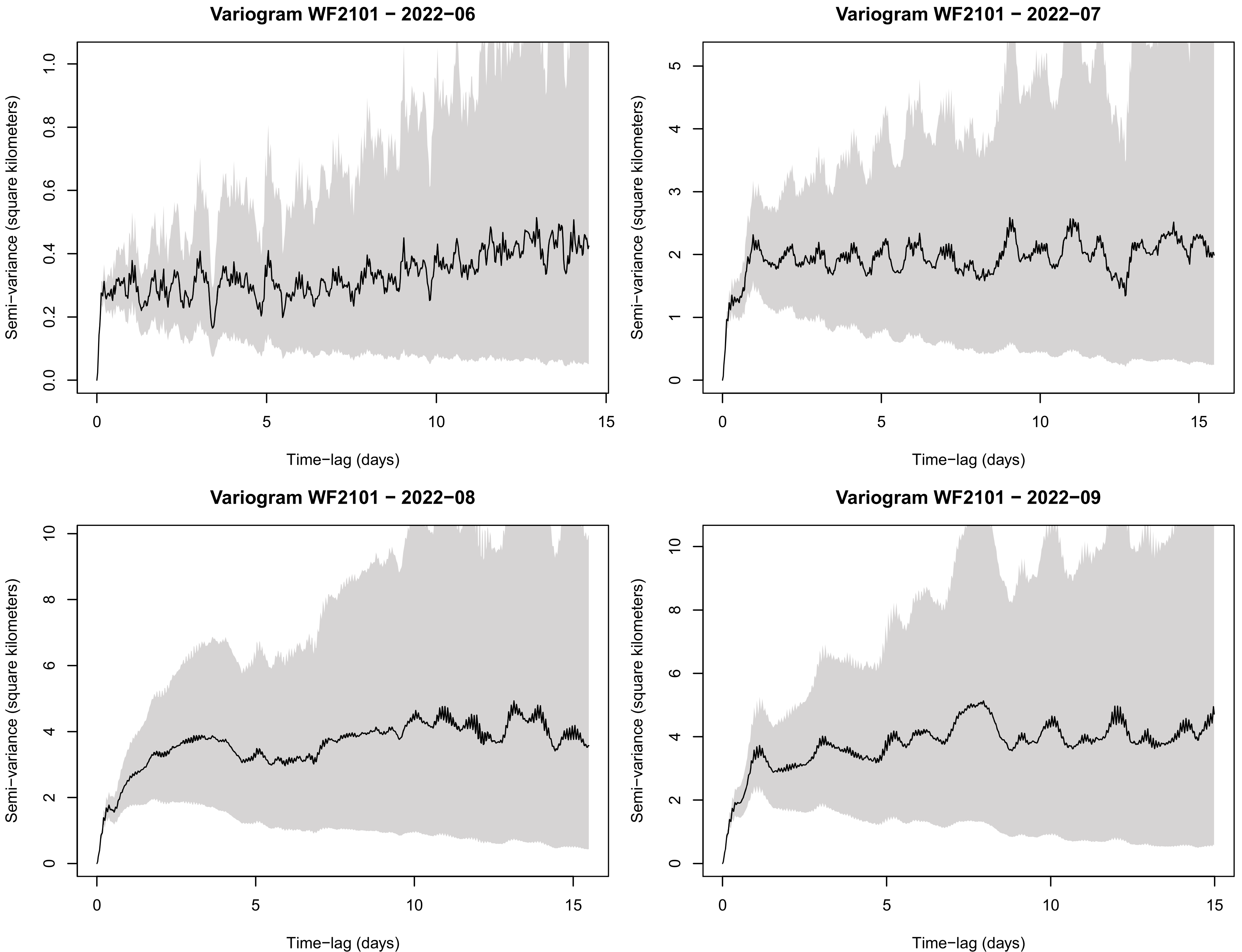
Variogram WF2101 − 2022−06
Variogram WF2101 − 2022−07
Variogram WF2101 − 2022−08
Variogram WF2101 − 2022−09
Semi−variance (square kilometers)
Time−lag (days)
0.0
0.2
0.4
0.6
0.8
1.0
0
1
2
3
4
5
6
8
10
15

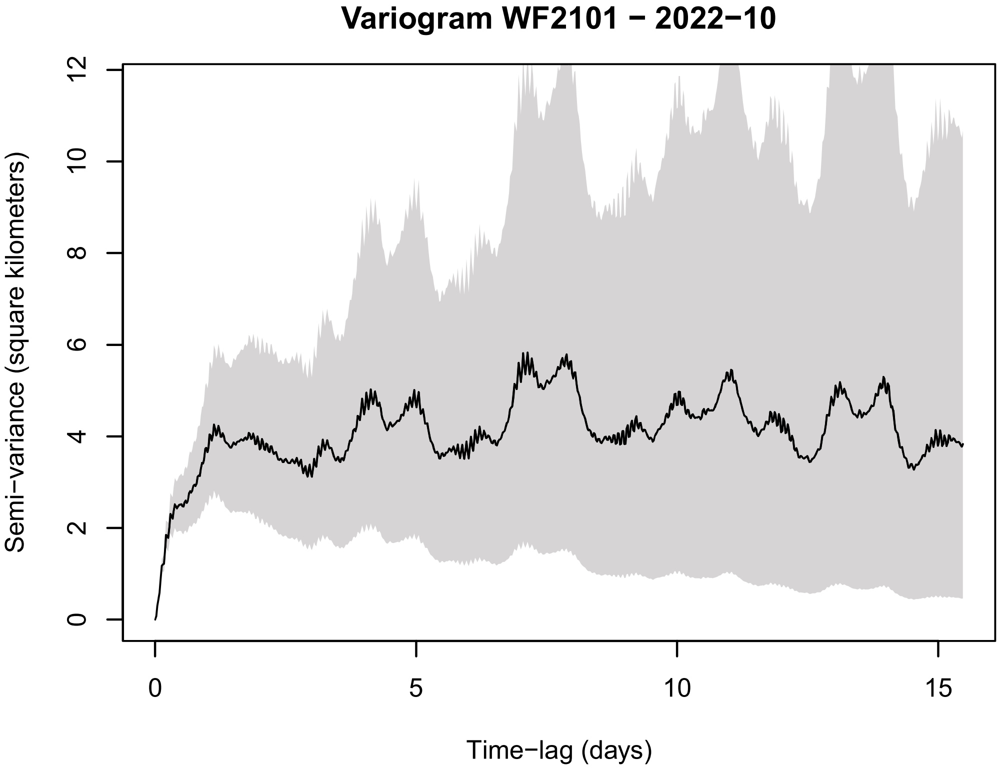
Variogram WF2101 − 2022−10
Semi−variance (square kilometers)
Time−lag (days)

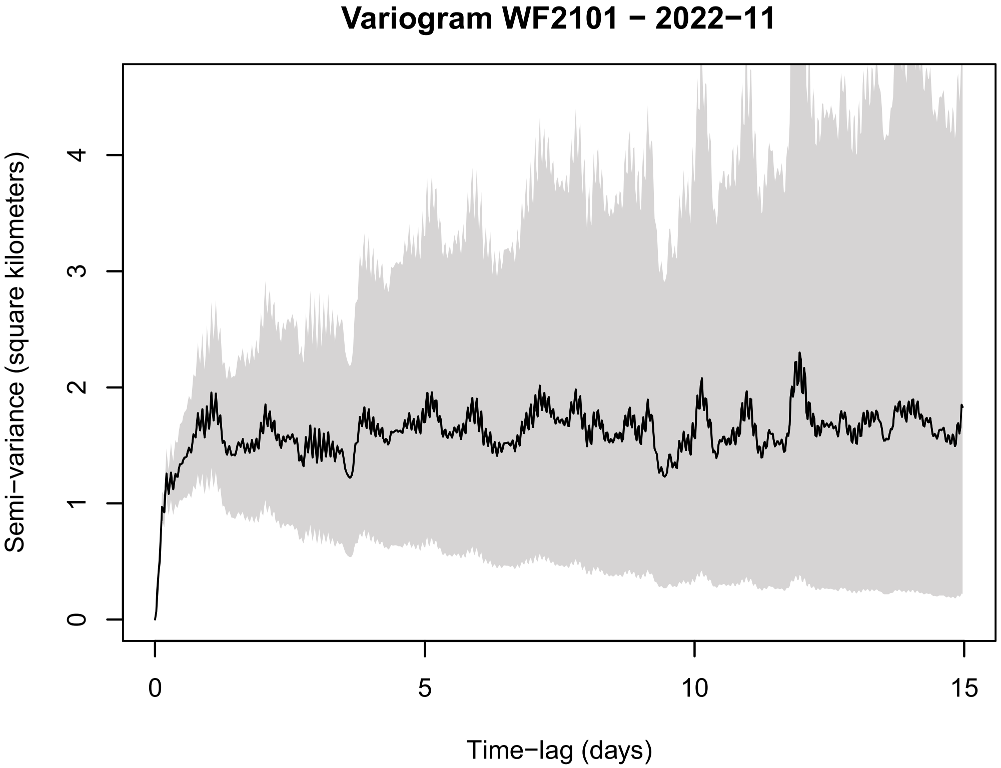
Variogram WF2101 − 2022−11
Semi−variance (square kilometers)
Time−lag (days)

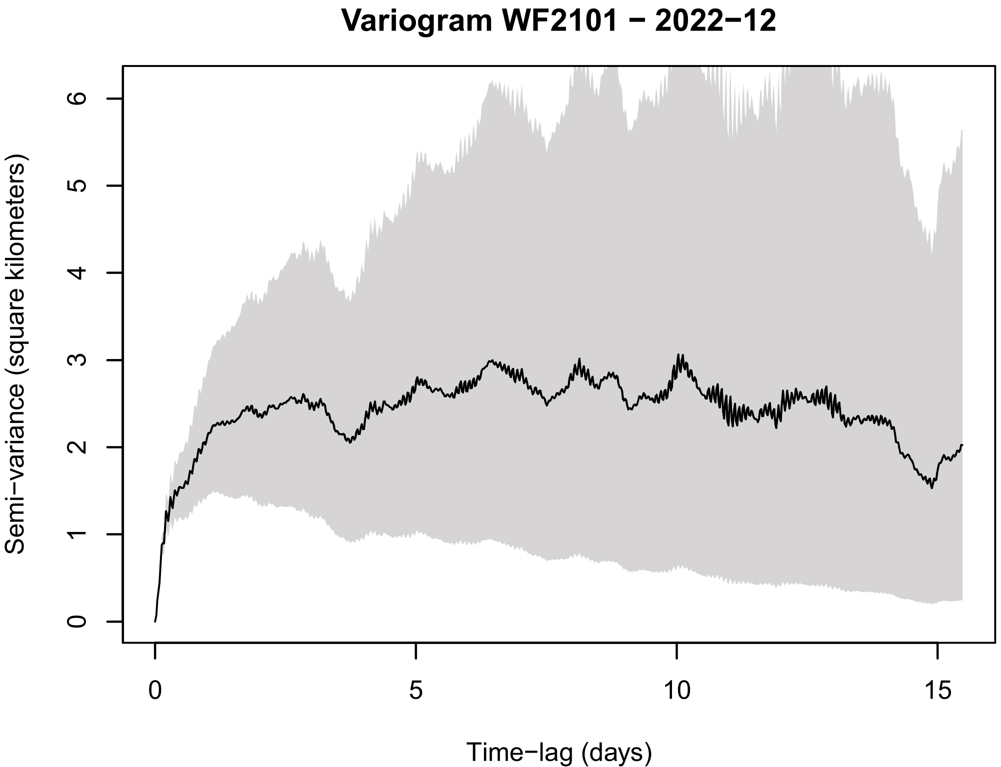
Variogram WF2101 − 2022−12
Semi−variance (square kilometers)
Time−lag (days)

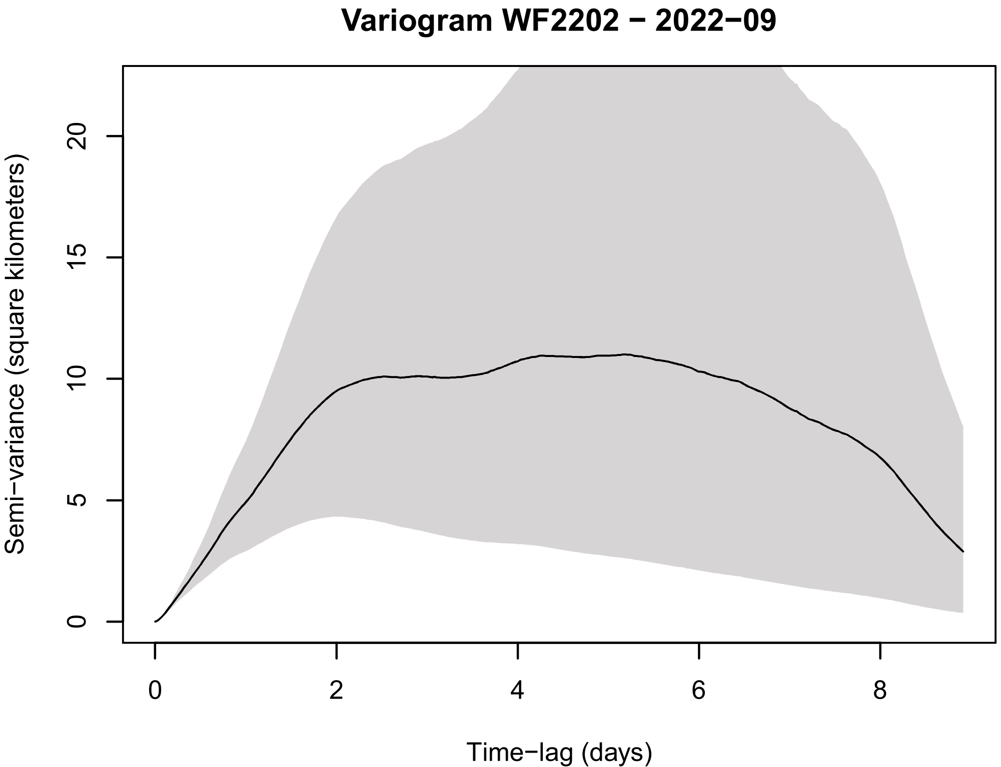
Variogram WF2202 − 2022−09
Semi−variance (square kilometers)
Time−lag (days)

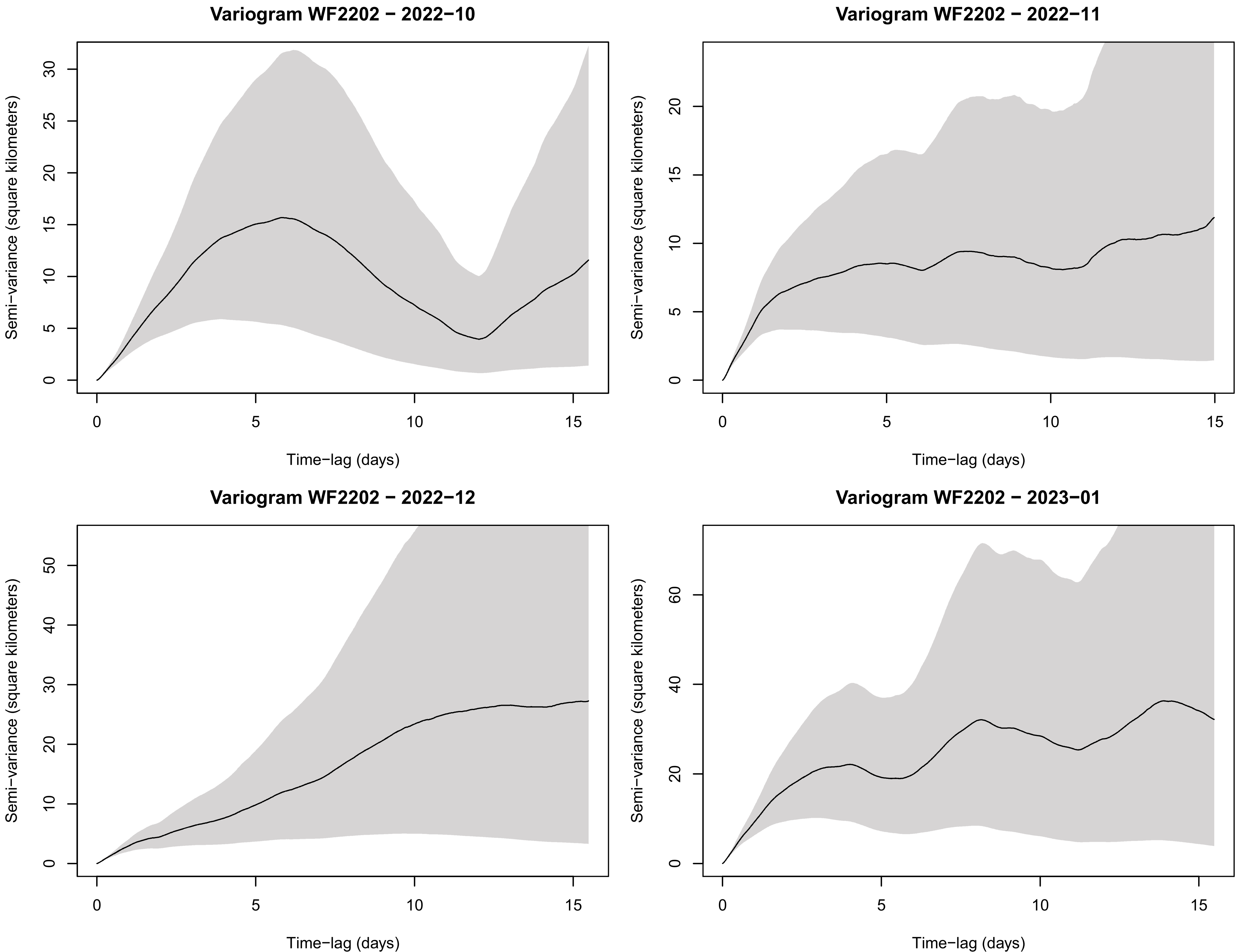

Variogram WF2202 − 2022−10
Variogram WF2202 − 2022−11
Variogram WF2202 − 2022−12
Variogram WF2202 − 2023−01
Semi−variance (square kilometers)
Time−lag (days)

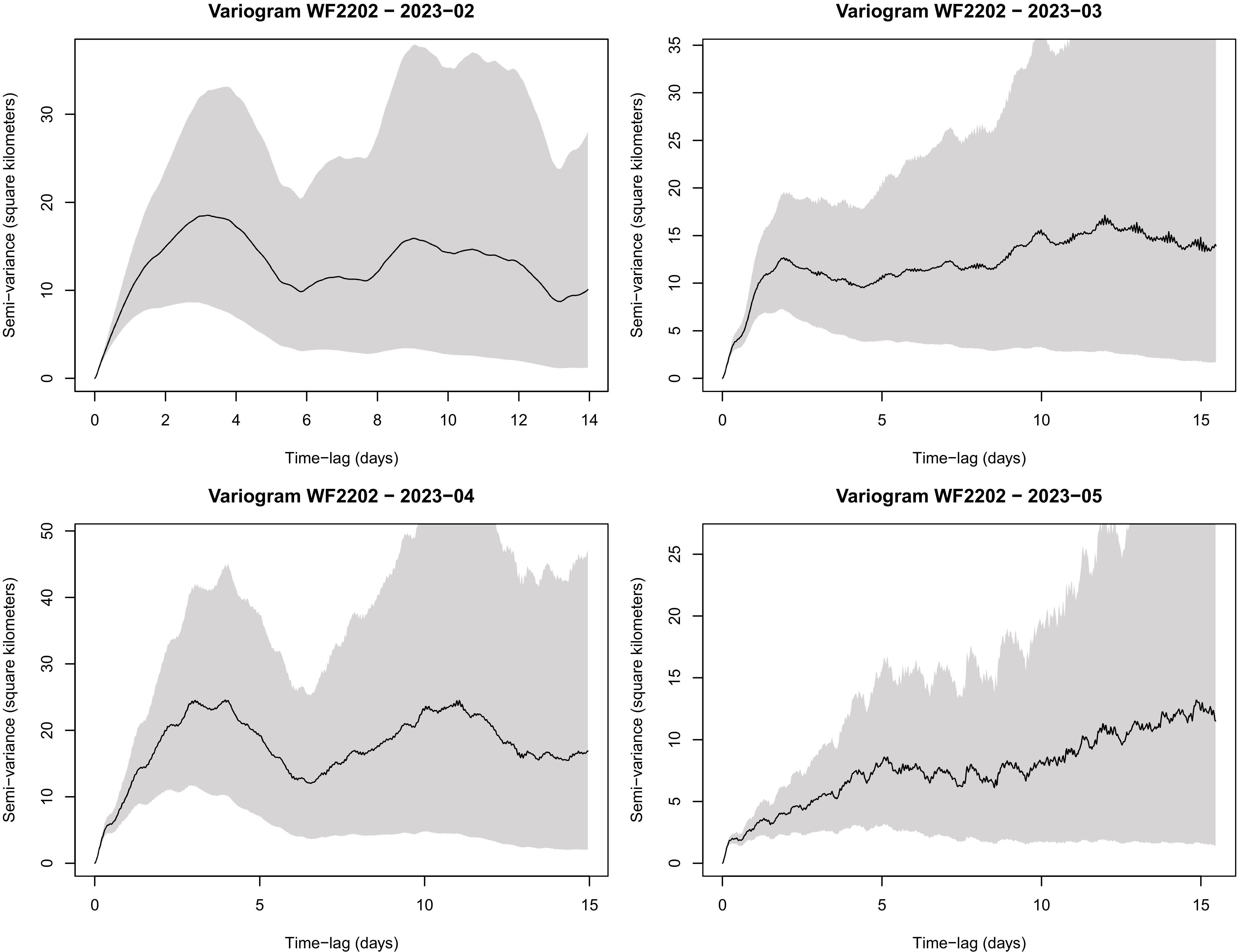
Variogram WF2202 − 2023−02
Semi−variance (square kilometers)
Time−lag (days)
Variogram WF2202 − 2023−03
Semi−variance (square kilometers)
Time−lag (days)
Variogram WF2202 − 2023−04
Semi−variance (square kilometers)
Time−lag (days)
Variogram WF2202 − 2023−05
Semi−variance (square kilometers)
Time−lag (days)

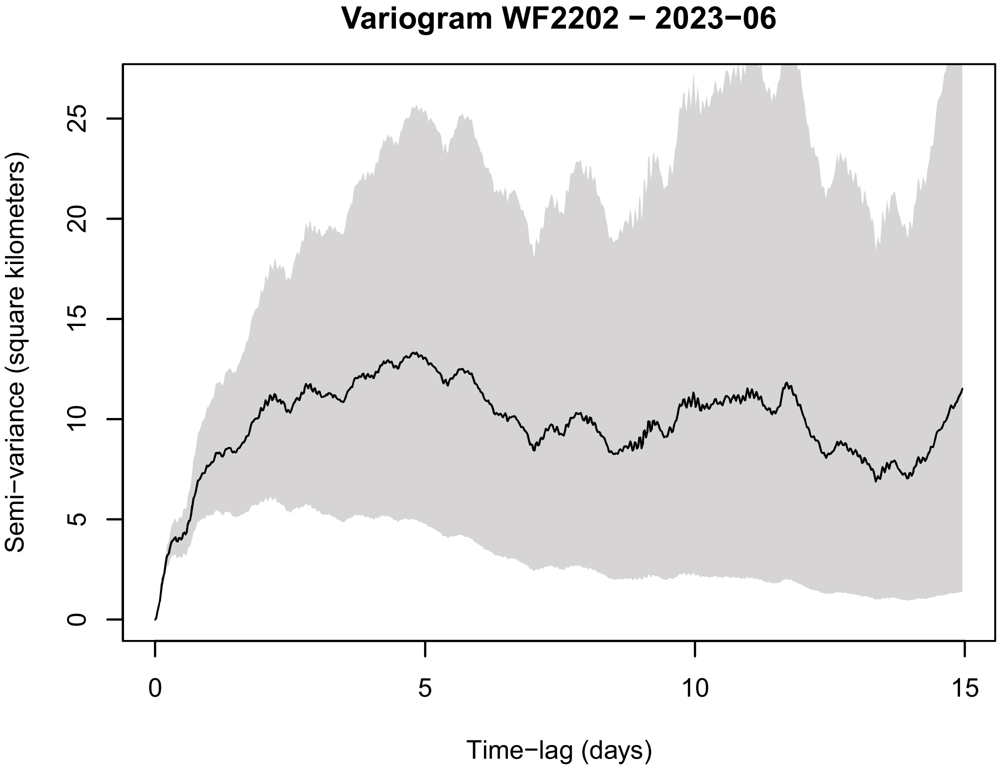
Variogram WF2202 − 2023−06
Semi−variance (square kilometers)
0
5
10
15
20
25
Time−lag (days)
0
5
10
15

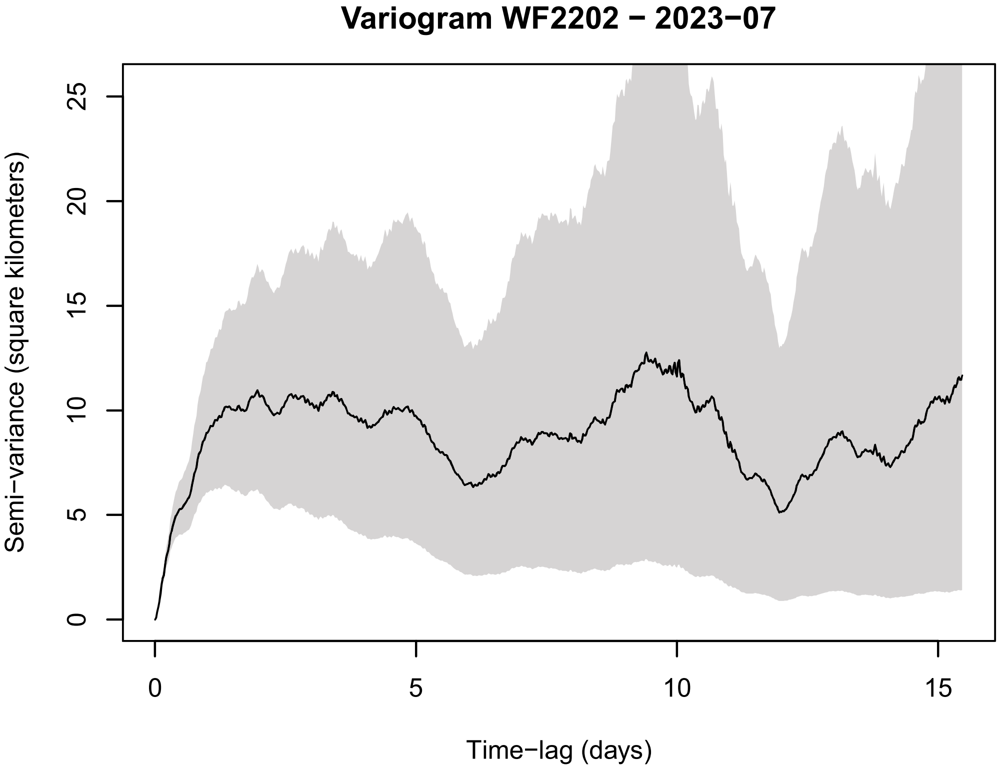
Variogram WF2202 − 2023−07
Semi−variance (square kilometers)
0
5
10
15
20
25
Time−lag (days)
0
5
10
15

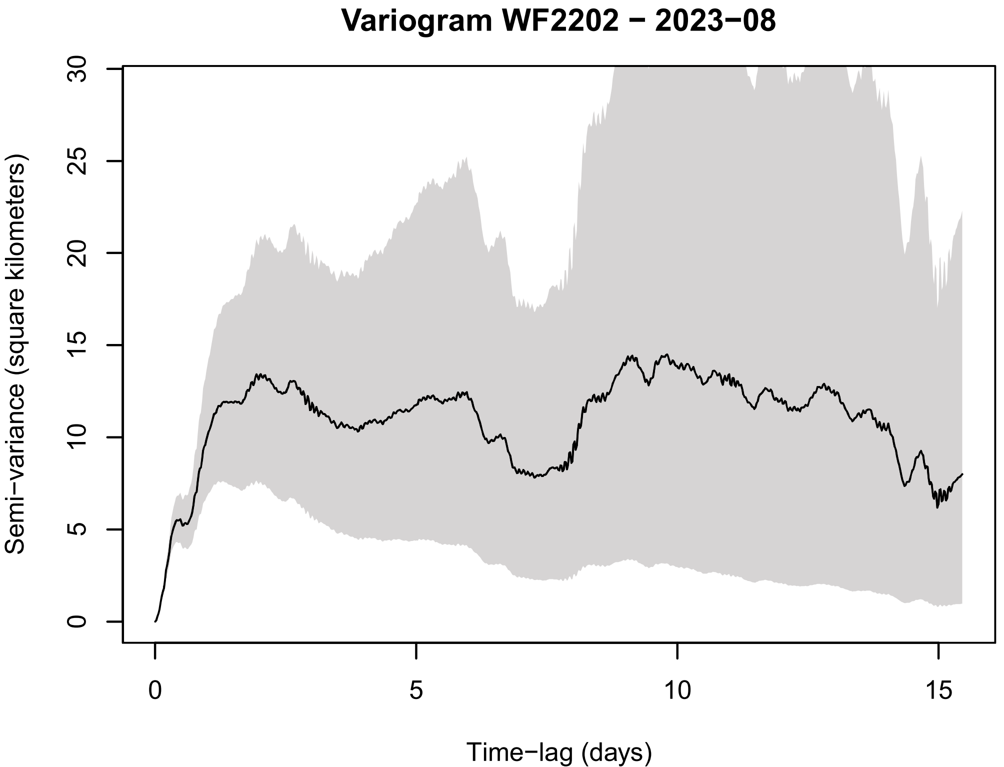
Variogram WF2202 − 2023−08
Semi−variance (square kilometers)
0
5
10
15
20
25
30
Time−lag (days)
0
5
10
15

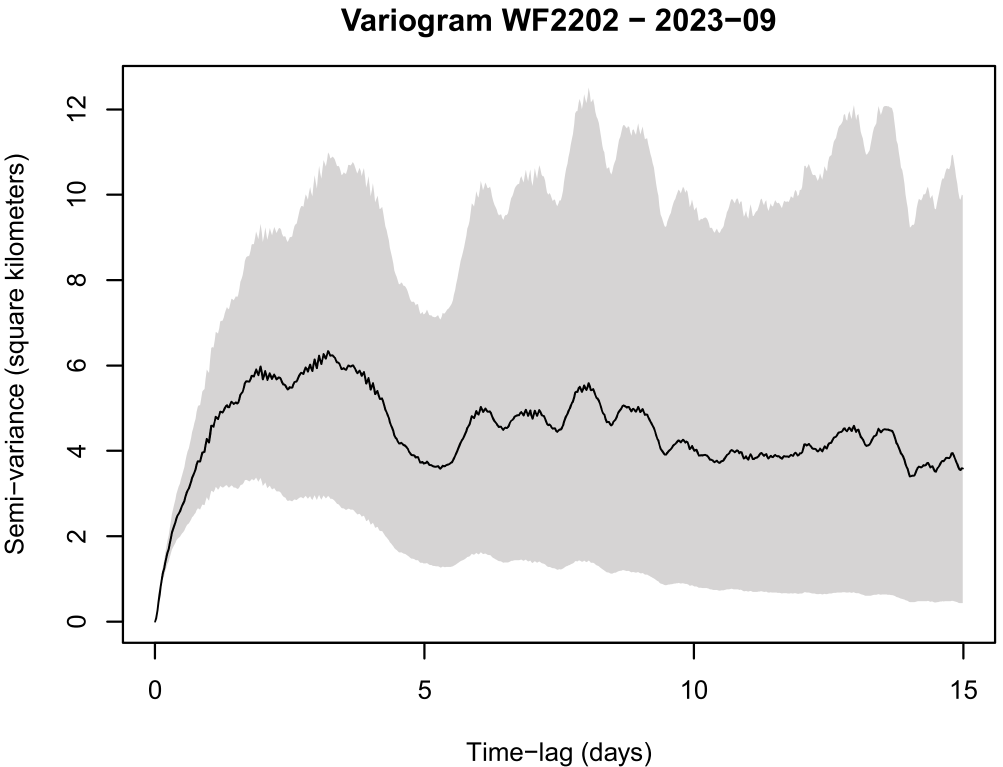
Variogram WF2202 − 2023−09
Semi−variance (square kilometers)
0
2
4
6
8
10
12
Time−lag (days)
0
5
10
15

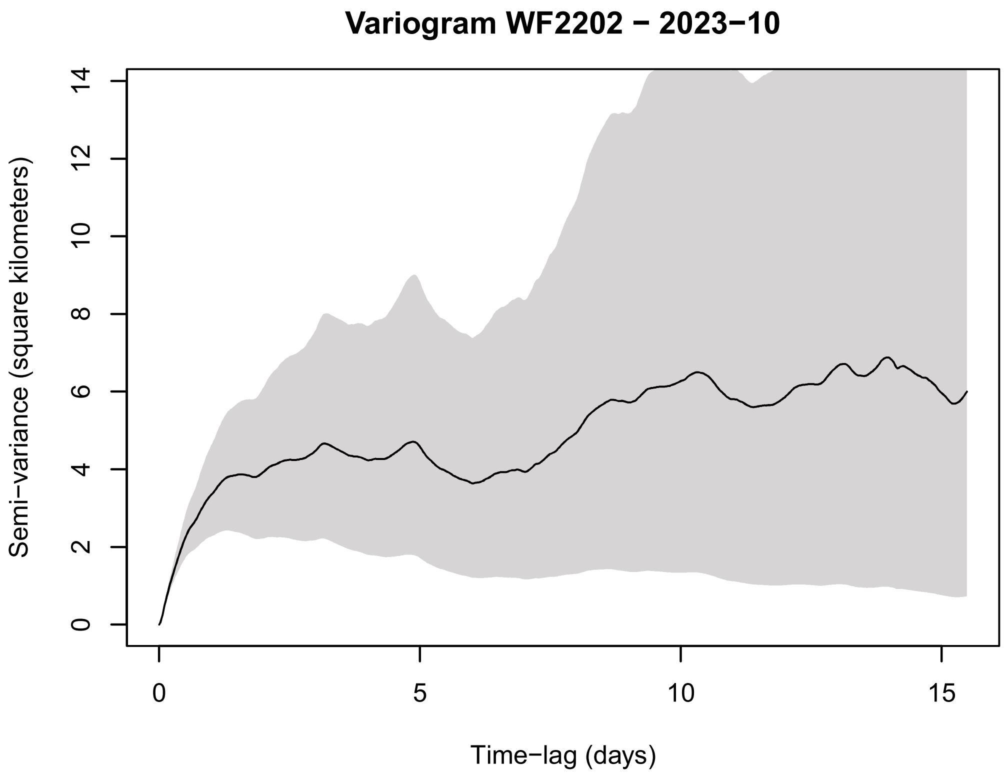
Variogram WF2202 − 2023−10
Semi−variance (square kilometers)
0
2
4
6
8
10
12
14
0
5
10
15
Time−lag (days)

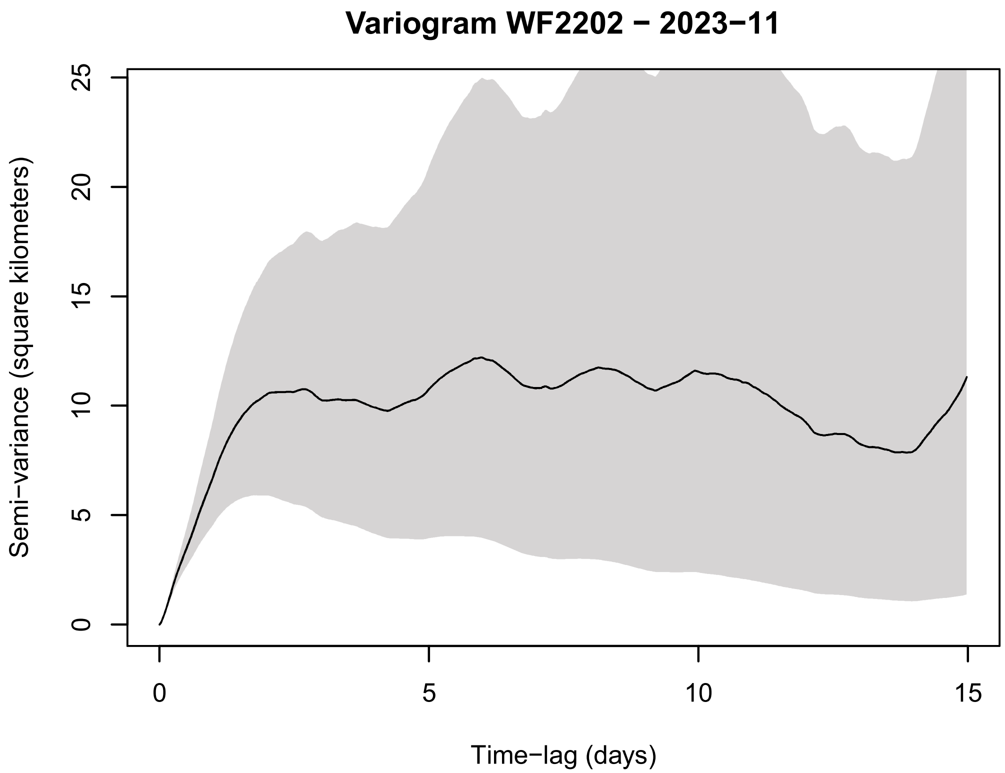
Variogram WF2202 − 2023−11
Semi−variance (square kilometers)
0
5
10
15
20
25
0
5
10
15
Time−lag (days)

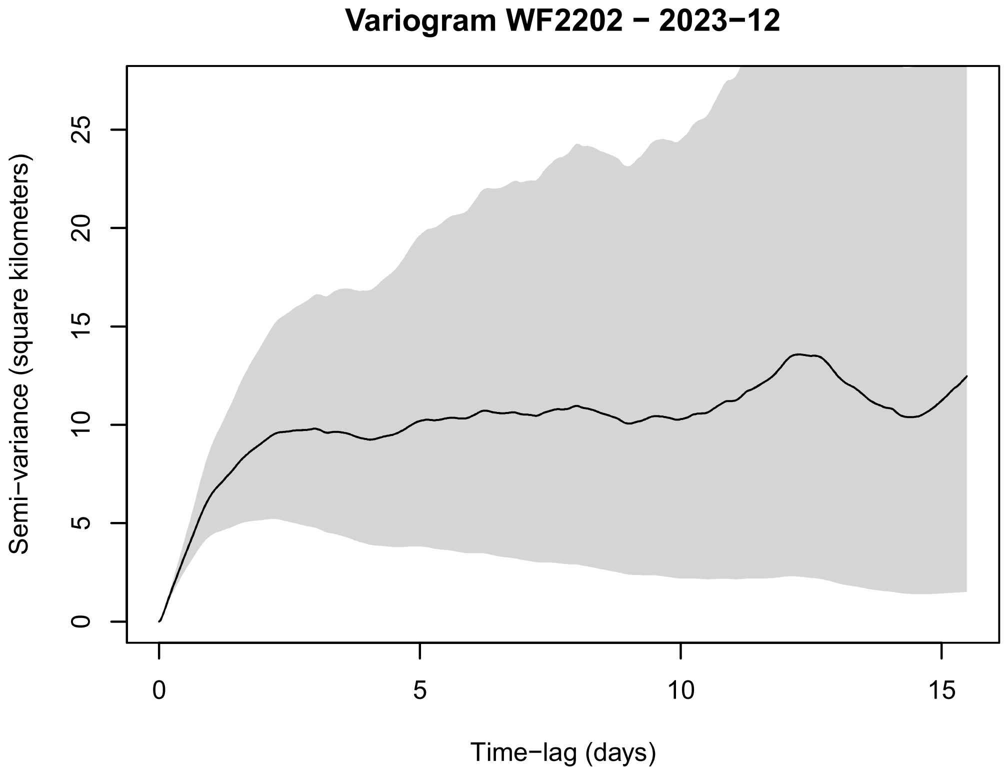
Variogram WF2202 − 2023−12
Semi−variance (square kilometers)
0
5
10
15
20
25
0
5
10
15
Time−lag (days)

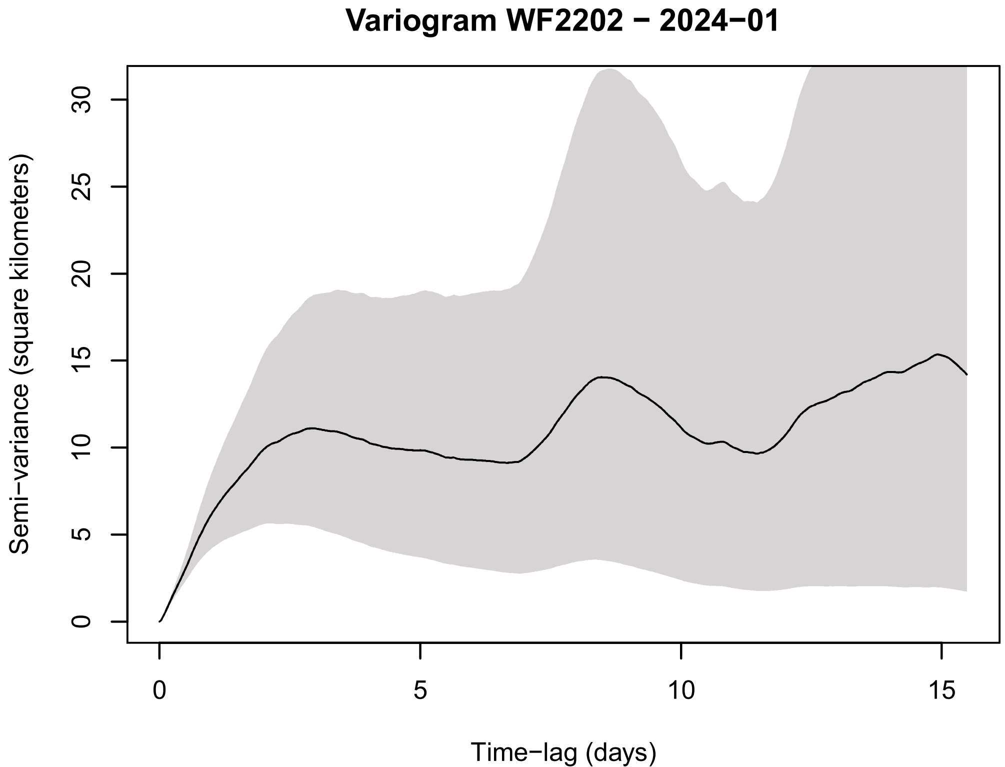
Variogram WF2202 − 2024−01
Semi−variance (square kilometers)
0
5
10
15
20
25
30
0
5
10
15
Time−lag (days)

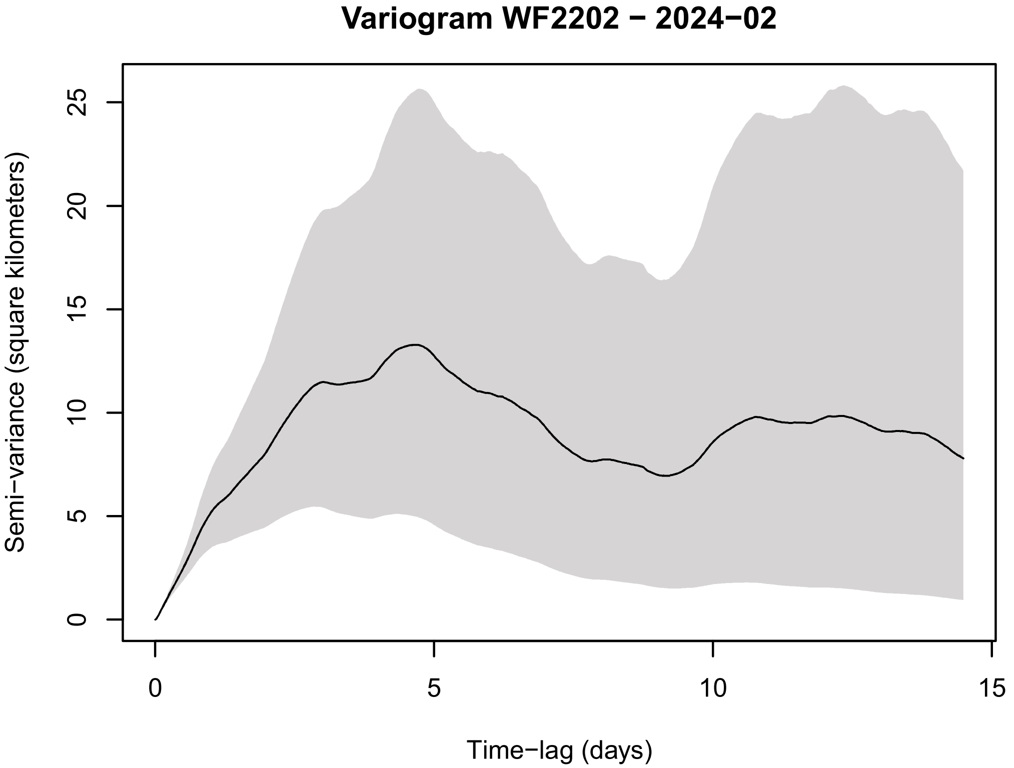
Variogram WF2202 − 2024−02
Semi−variance (square kilometers)
Time−lag (days)
0
5
10
15
20
25

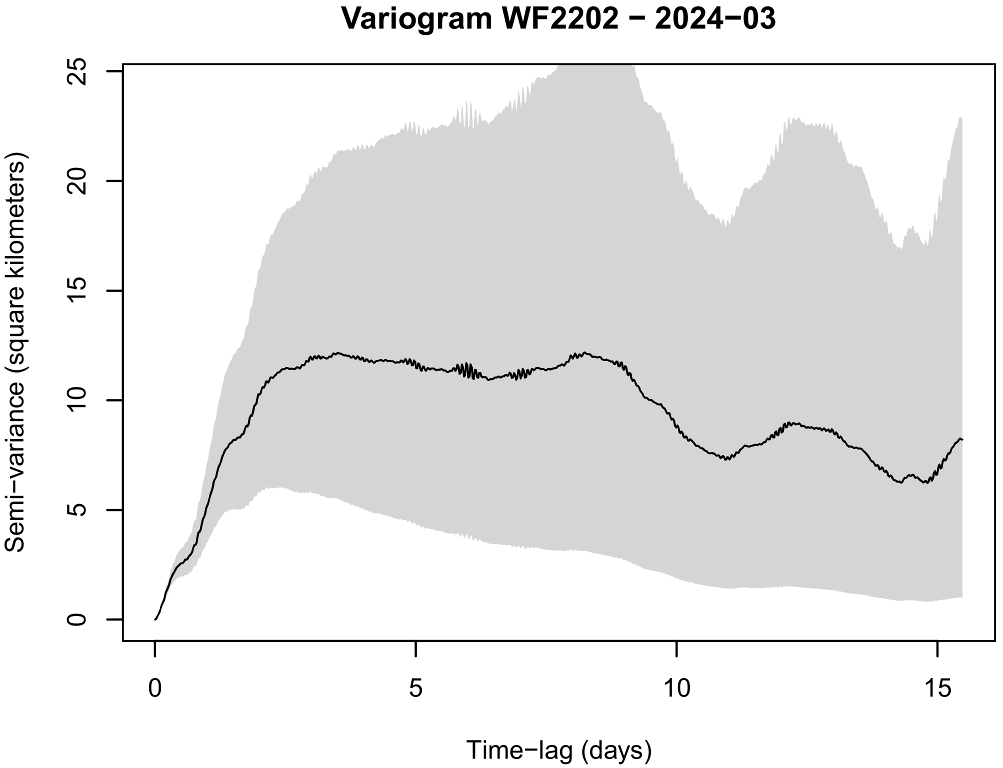
Variogram WF2202 − 2024−03
Semi−variance (square kilometers)
Time−lag (days)
0
5
10
15
20
25

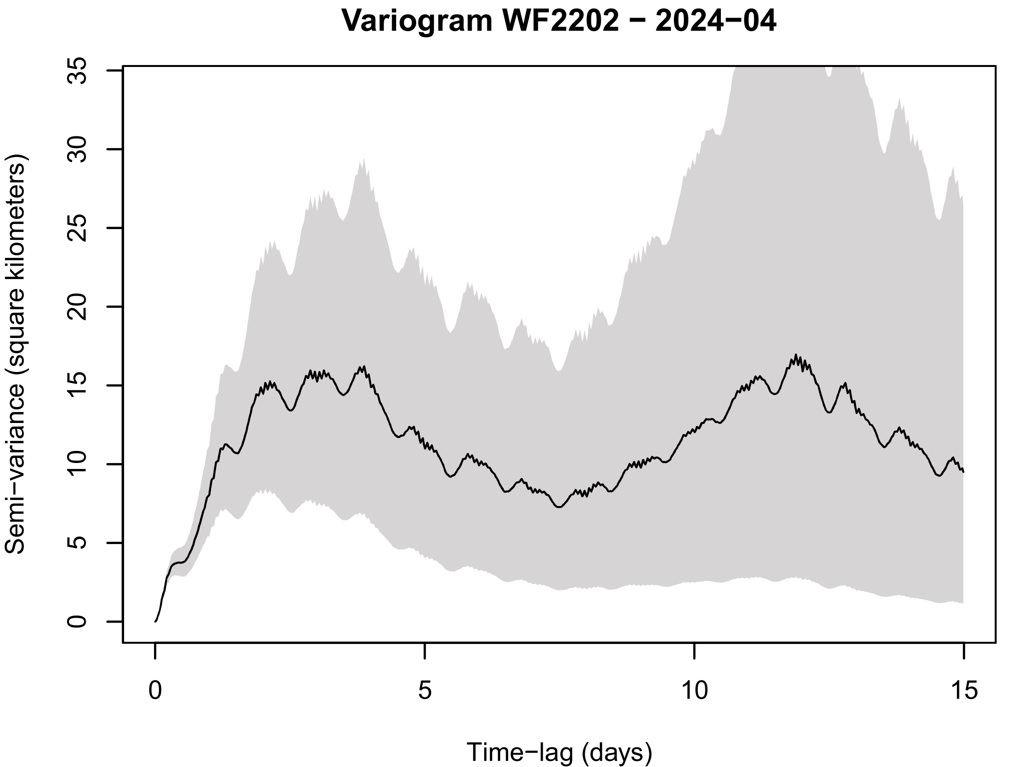
Variogram WF2202 − 2024−04
Semi−variance (square kilometers)
Time−lag (days)
0
5
10
15
20
25
30
35

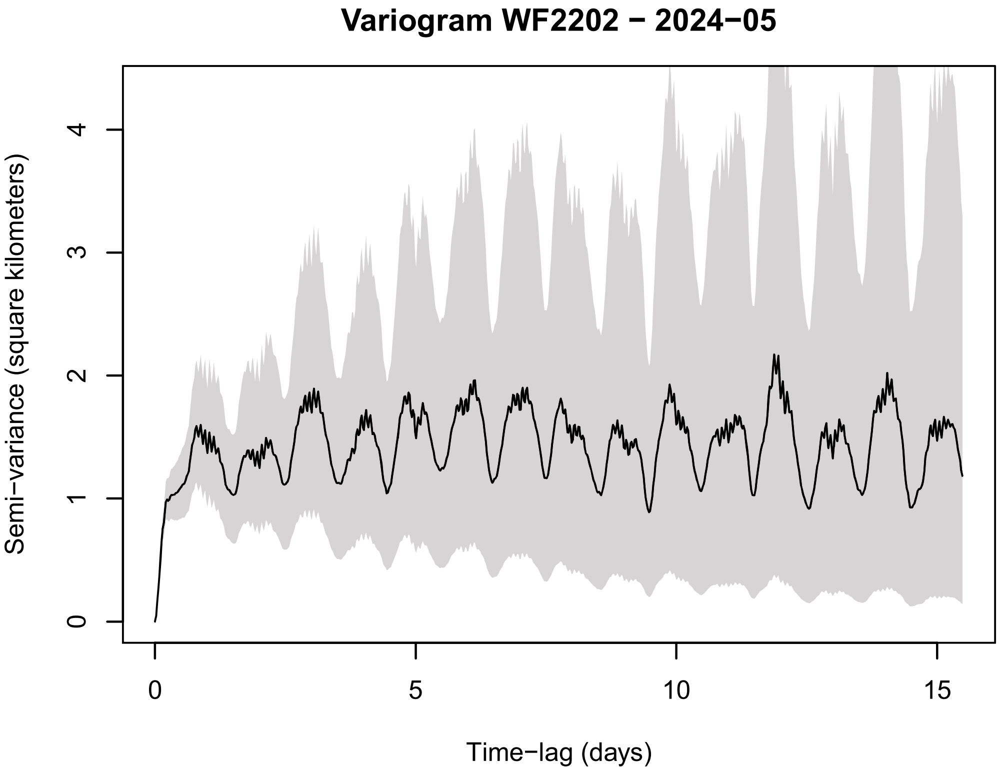
Variogram WF2202 − 2024−05
Semi−variance (square kilometers)
Time−lag (days)
0
1
2
3
4
5
10
15

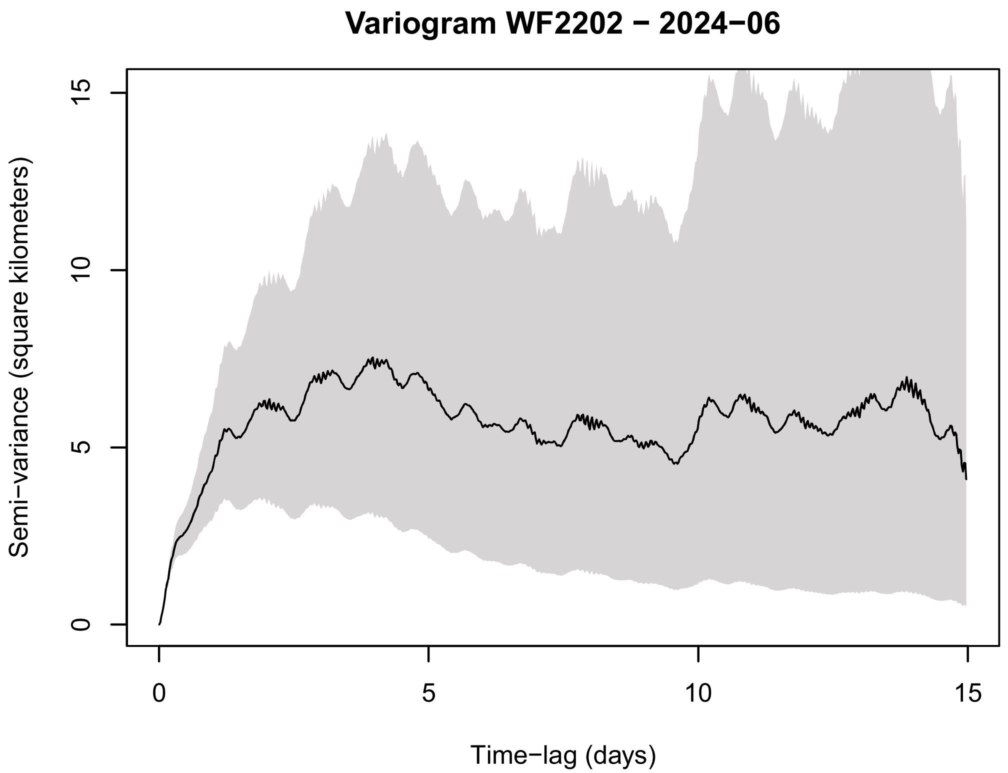
Variogram WF2202 − 2024−06
Semi−variance (square kilometers)
Time−lag (days)
0
5
10
15

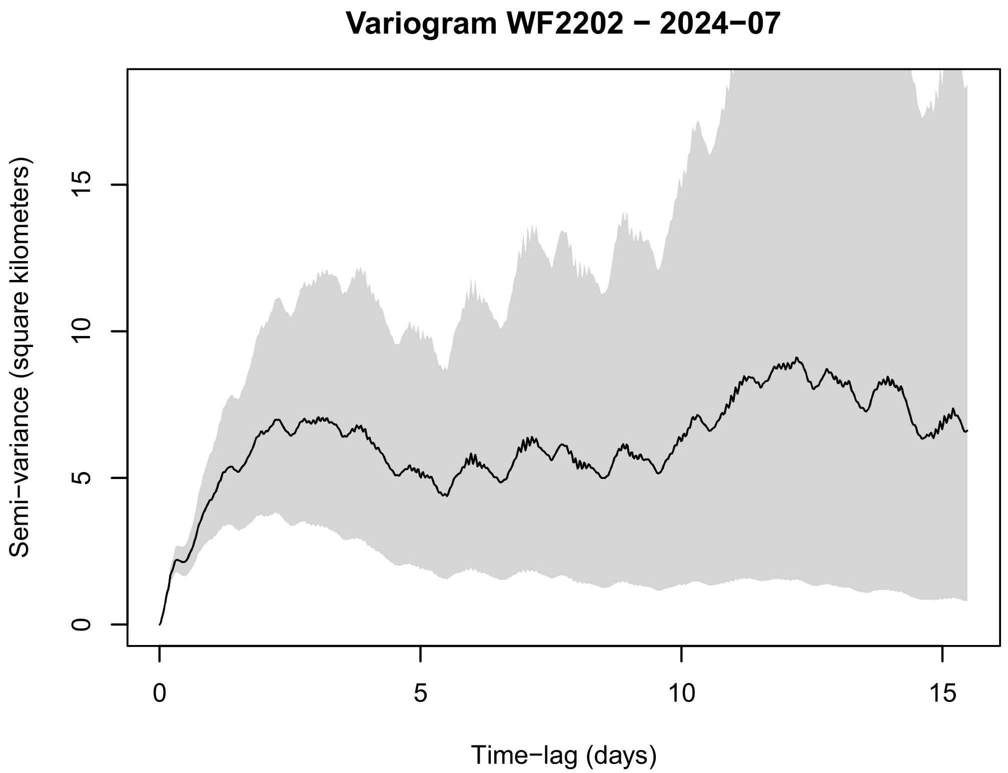
Variogram WF2202 − 2024−07
Semi−variance (square kilometers)
Time−lag (days)
0
5
10
15

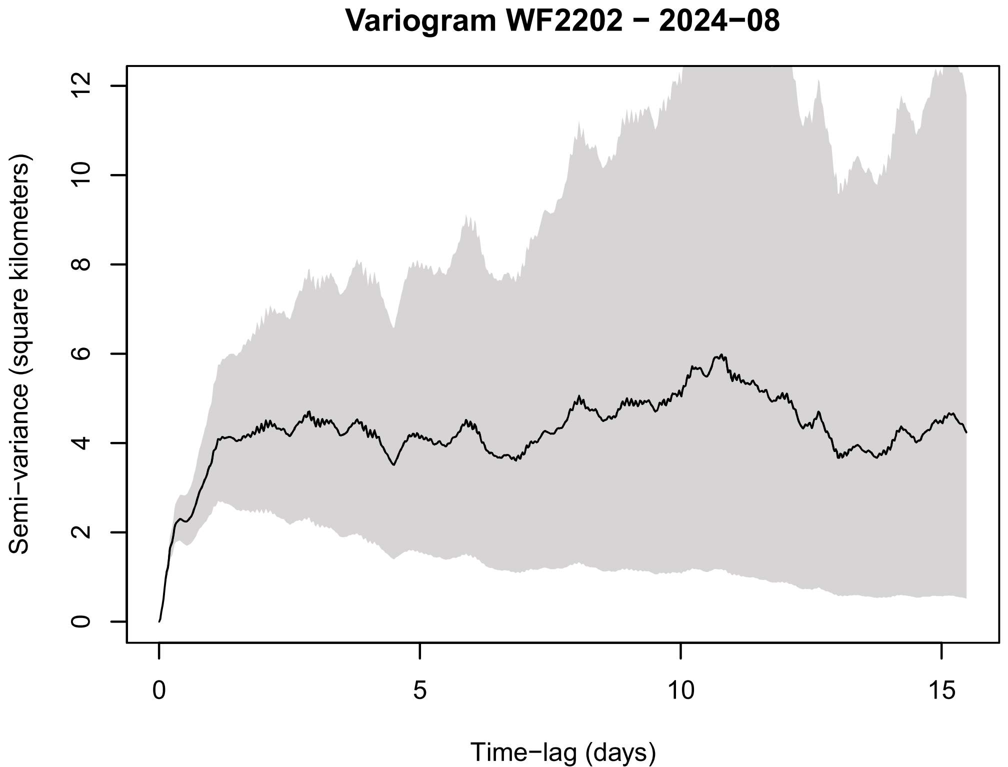
Variogram WF2202 − 2024−08
Semi−variance (square kilometers)
Time−lag (days)
0
2
4
6
8
10
12
0
5
10
15

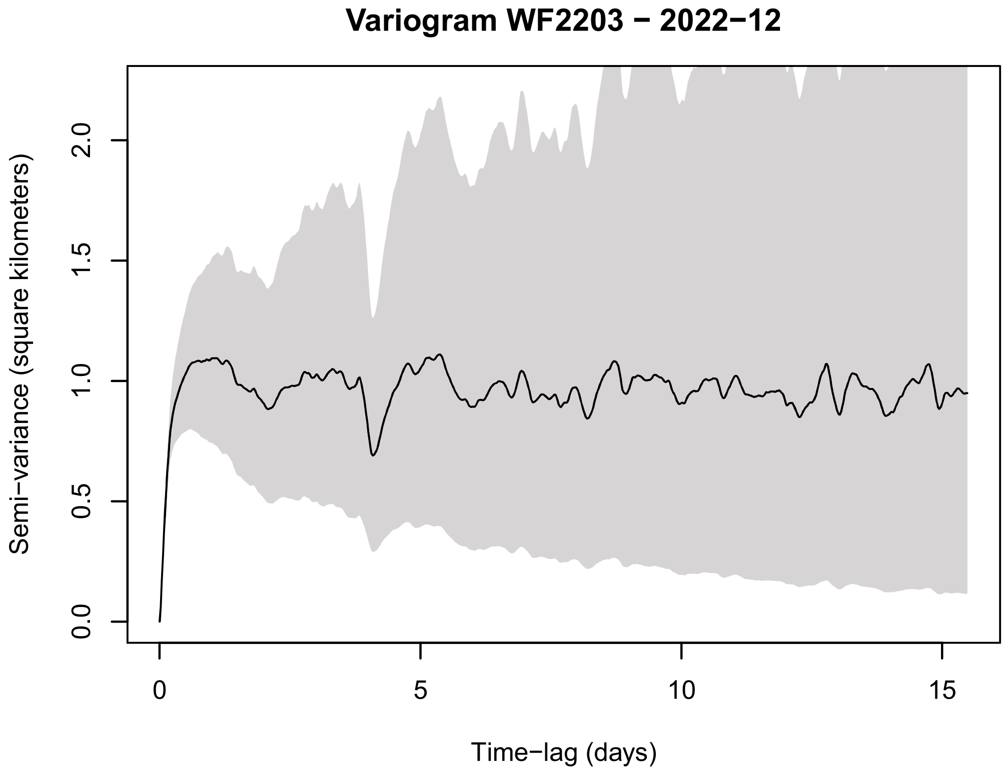
Variogram WF2203 − 2022−12
Semi−variance (square kilometers)
Time−lag (days)
0.0
0.5
1.0
1.5
2.0
0
5
10
15

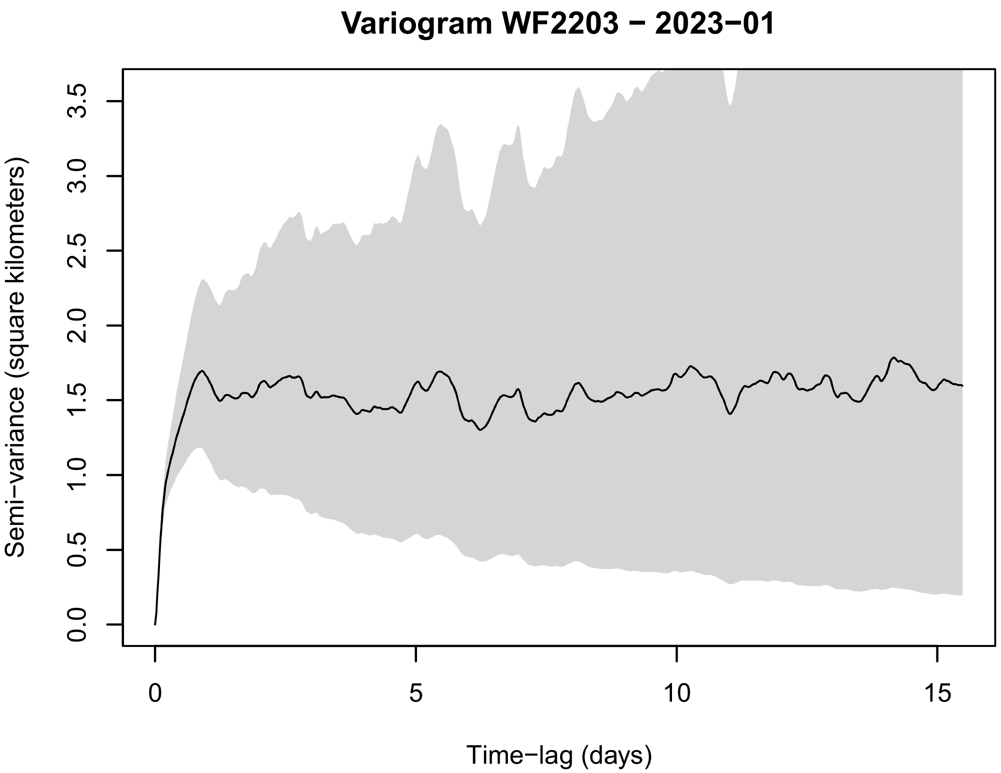

Variogram WF2203 − 2023−01
Semi−variance (square kilometers)
0.0
0.5
1.0
1.5
2.0
2.5
3.0
3.5
0
5
10
15
Time−lag (days)


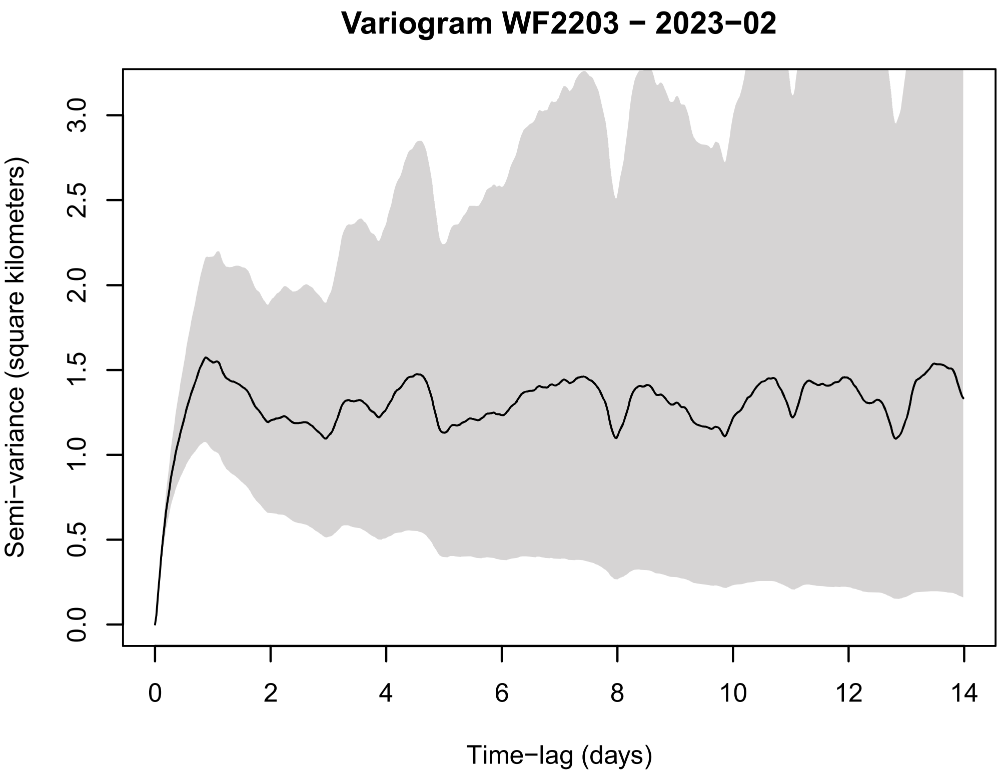

Variogram WF2203 − 2023−02
Semi−variance (square kilometers)
0.0
0.5
1.0
1.5
2.0
2.5
3.0
0
2
4
6
8
10
12
14
Time−lag (days)


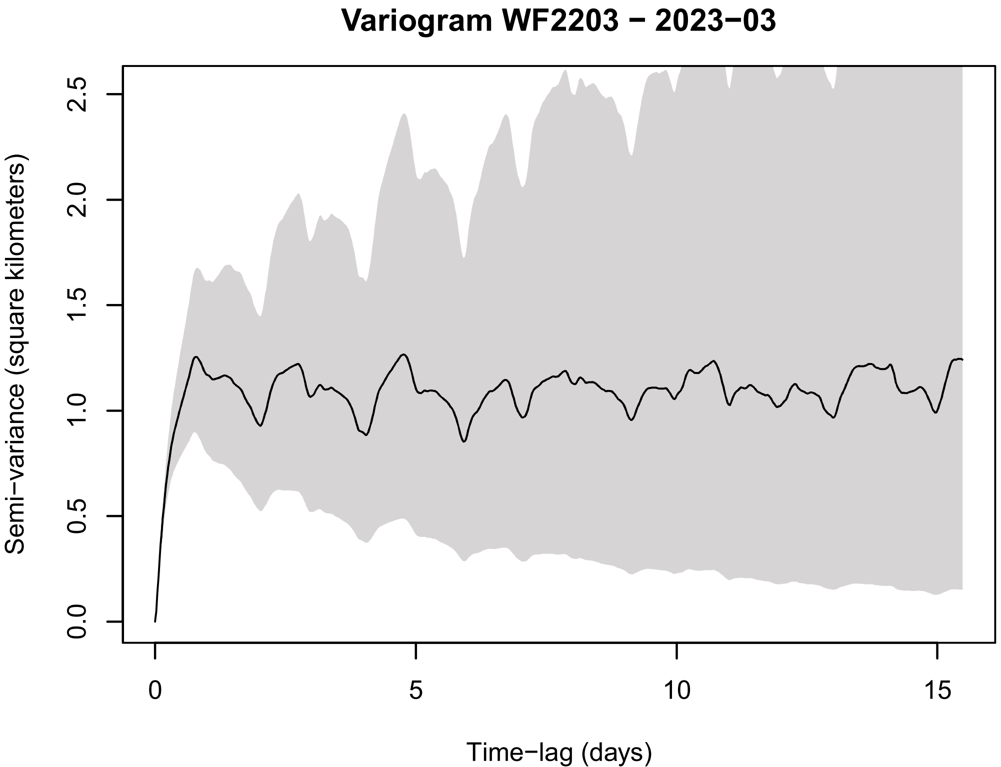

Variogram WF2203 − 2023−03
Semi−variance (square kilometers)
0.0
0.5
1.0
1.5
2.0
2.5
0
5
10
15
Time−lag (days)


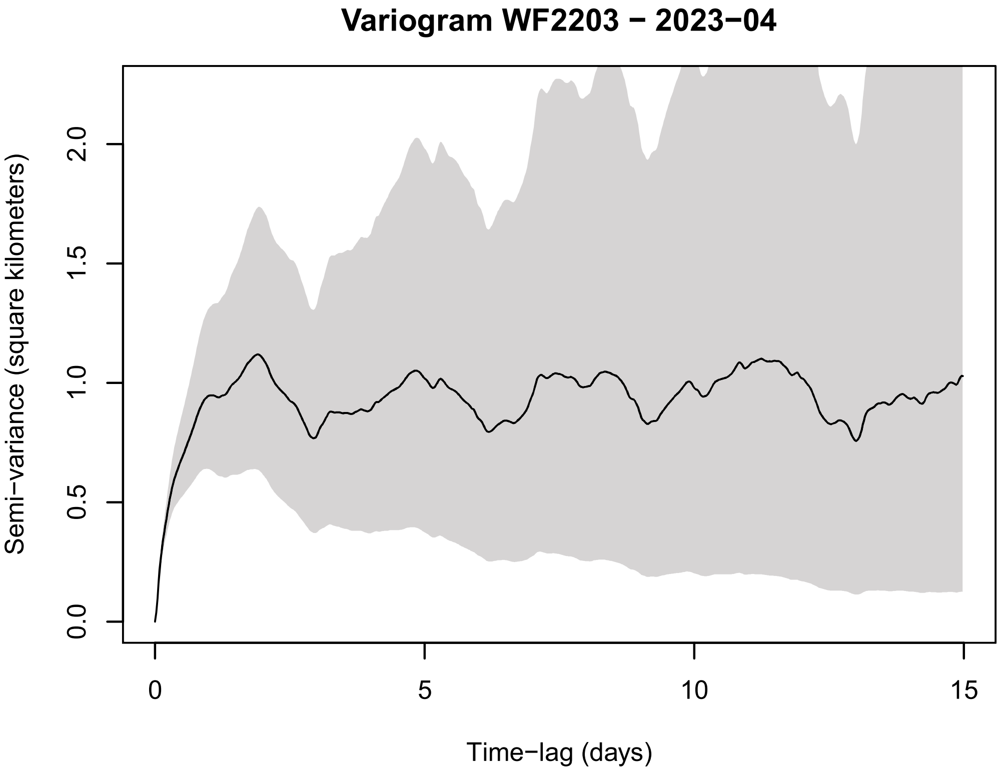

Variogram WF2203 − 2023−04
Semi−variance (square kilometers)
0.0
0.5
1.0
1.5
2.0
0
5
10
15
Time−lag (days)

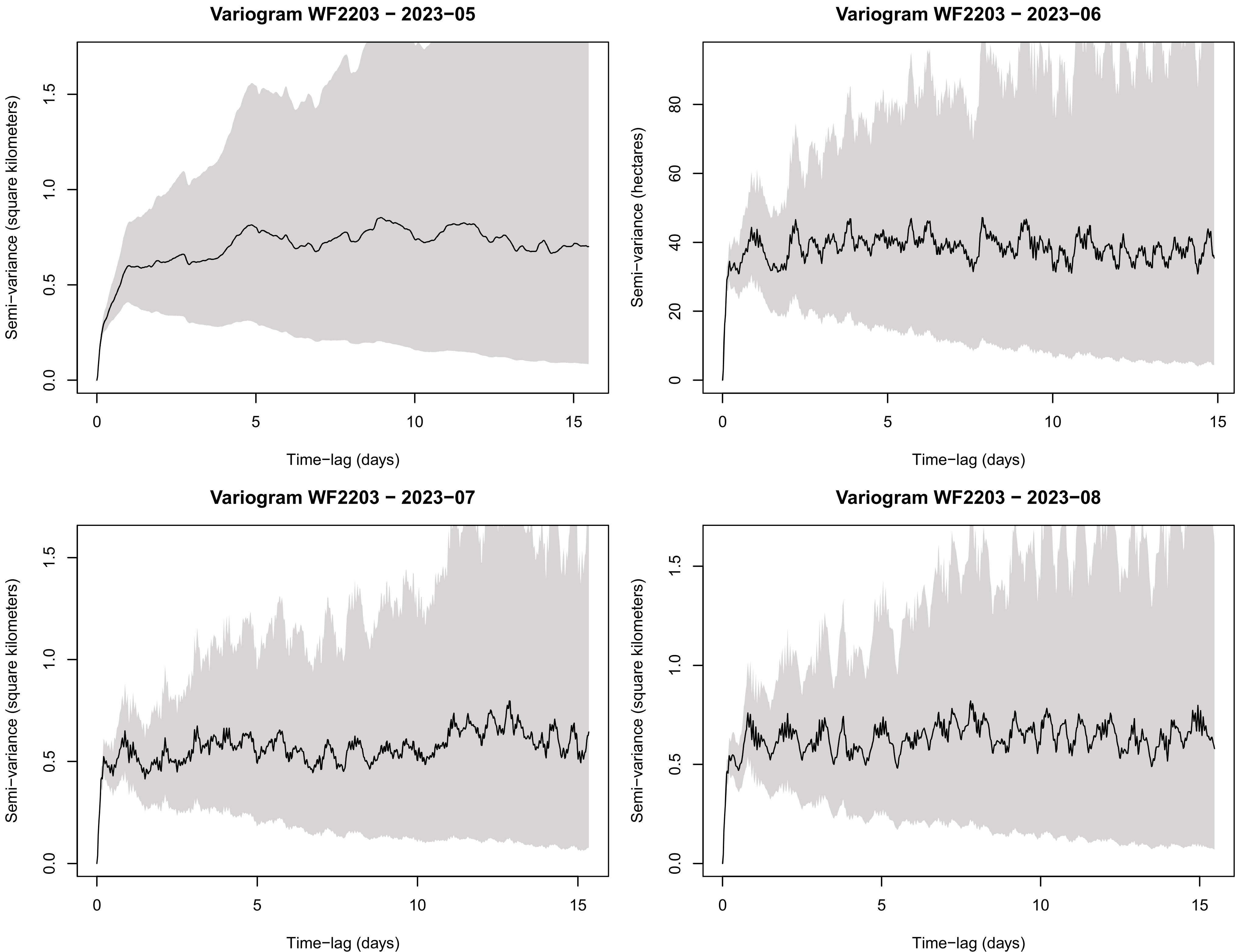

Variogram WF2203 − 2023−05
Semi−variance (square kilometers)
Time−lag (days)
Variogram WF2203 − 2023−06
Semi−variance (hectares)
Time−lag (days)
Variogram WF2203 − 2023−07
Semi−variance (square kilometers)
Time−lag (days)
Variogram WF2203 − 2023−08
Semi−variance (square kilometers)
Time−lag (days)

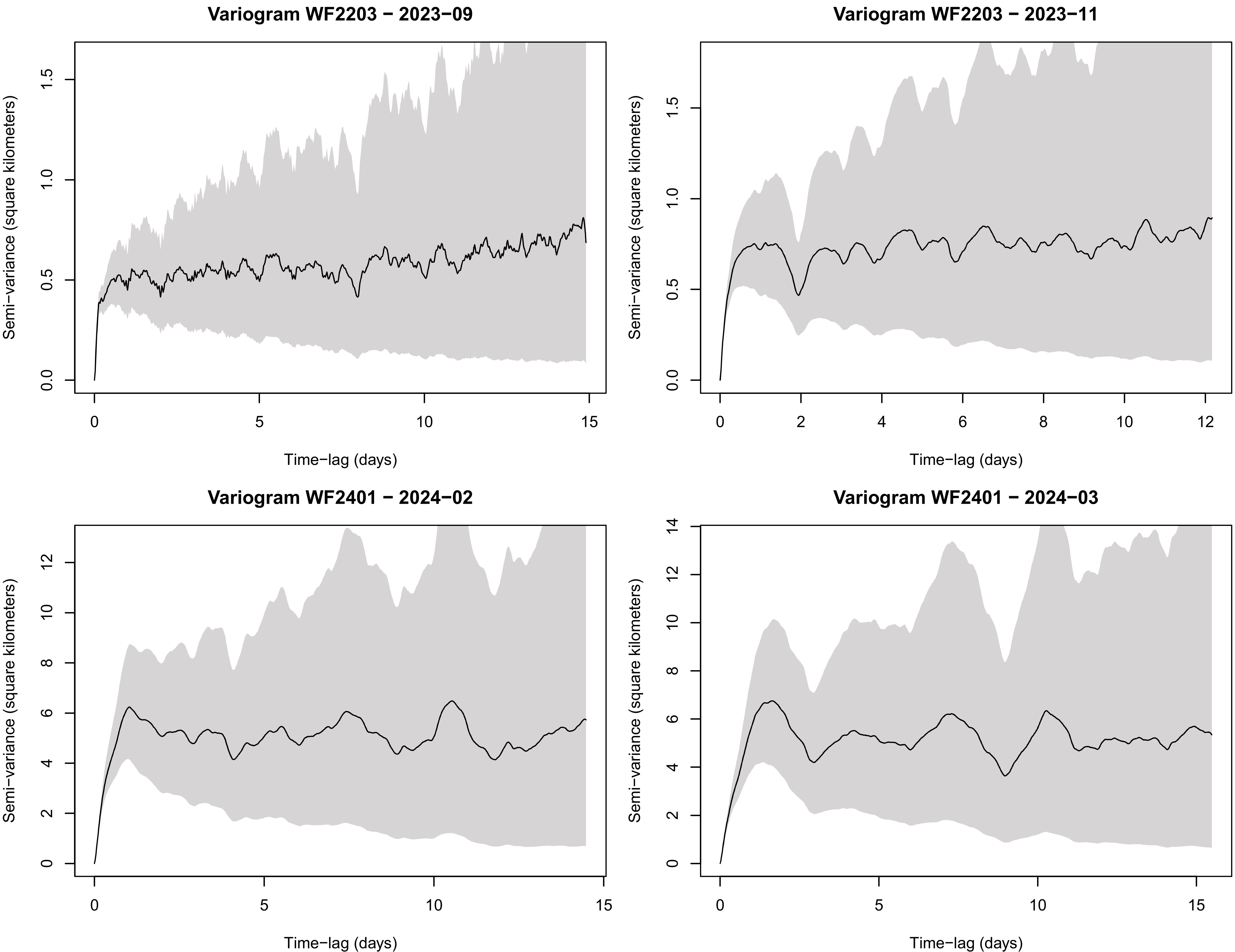

Variogram WF2203 − 2023−09
Variogram WF2203 − 2023−11
Variogram WF2401 − 2024−02
Variogram WF2401 − 2024−03
Semi−variance (square kilometers)
Time−lag (days)
0.0
0.5
1.0
1.5
0
2
4
6
8
10
12
14
5
15

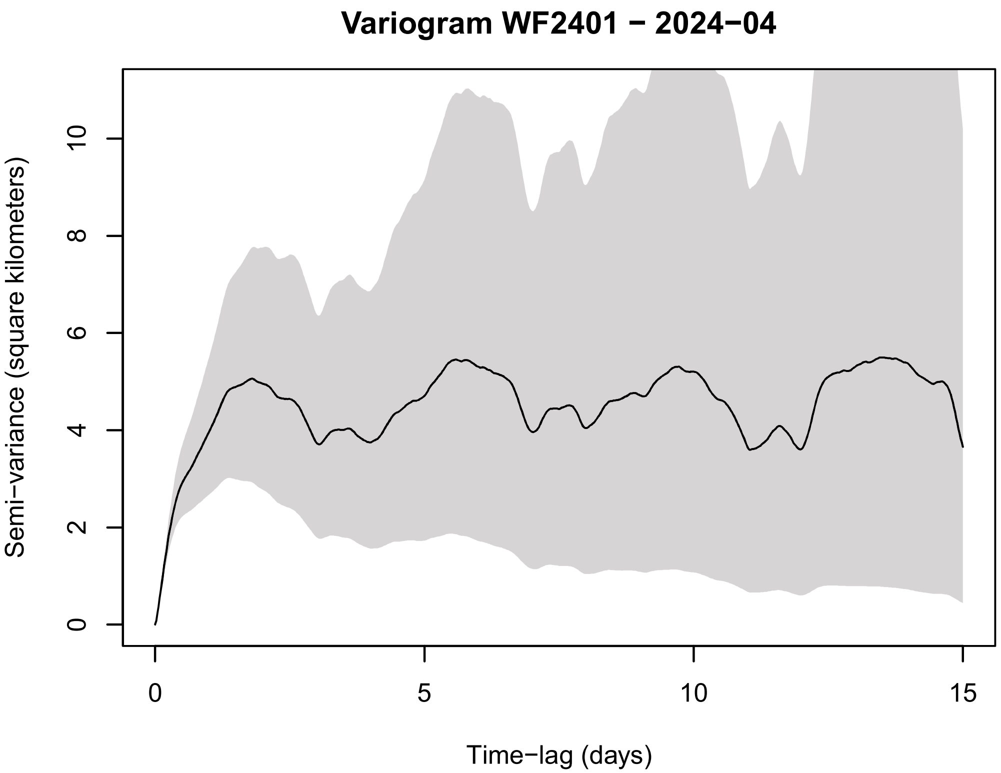
Variogram WF2401 − 2024−04
Semi−variance (square kilometers)
0
2
4
6
8
10
Time−lag (days)
0
5
10
15

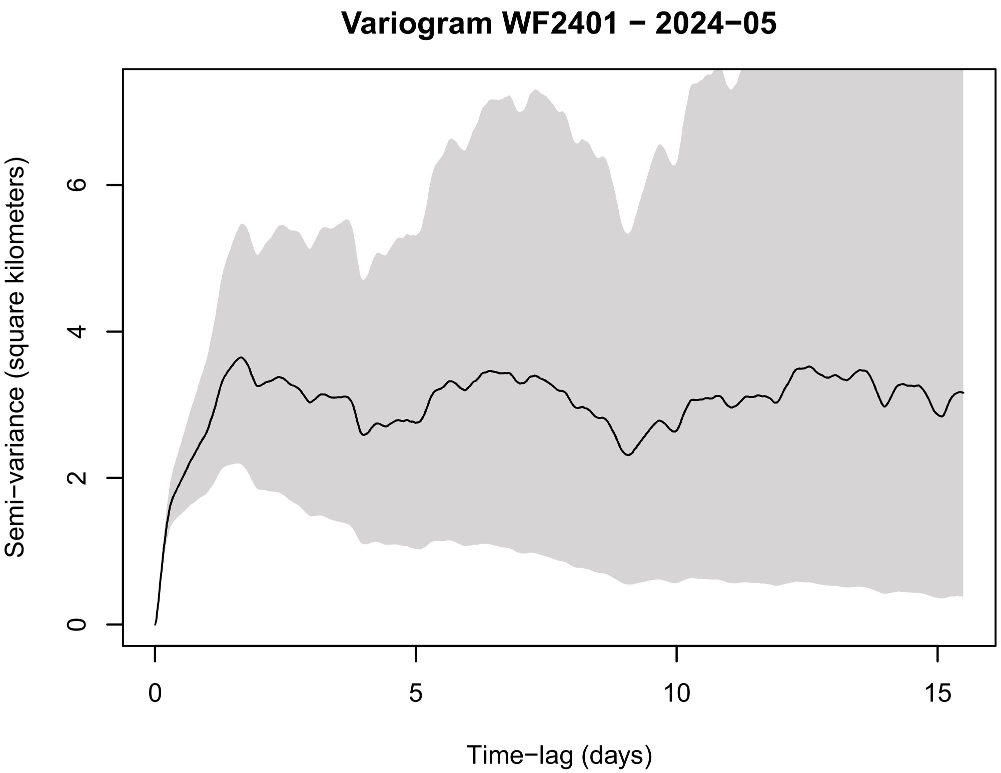
Variogram WF2401 − 2024−05
Semi−variance (square kilometers)
0
2
4
6
Time−lag (days)
0
5
10
15

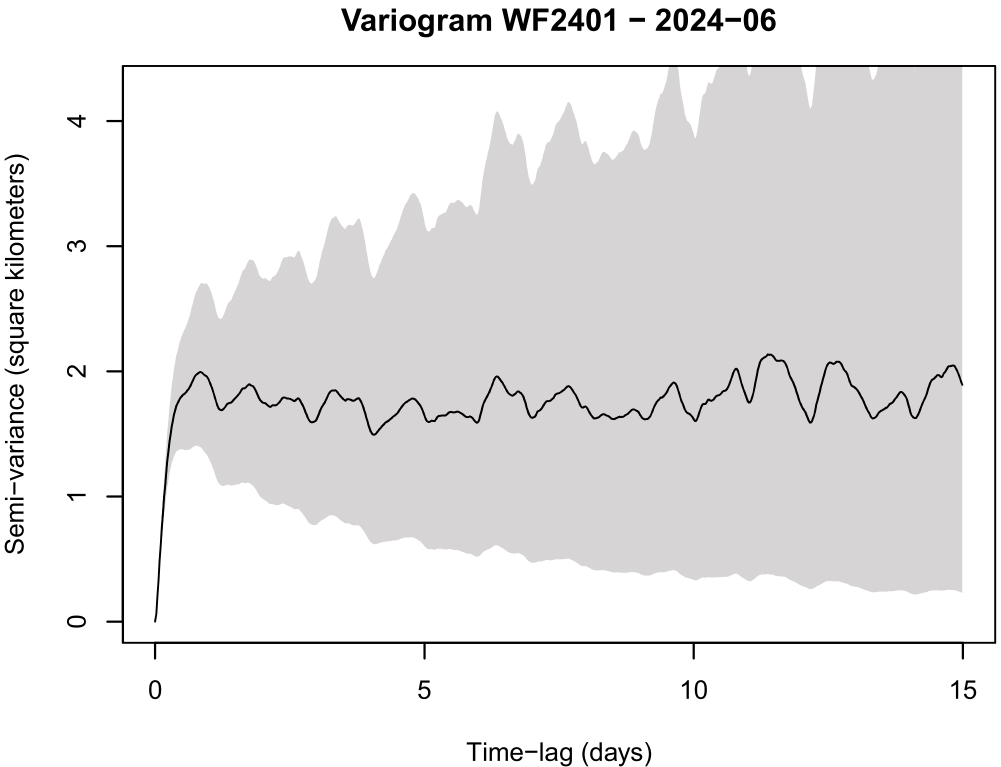
Variogram WF2401 − 2024−06
Semi−variance (square kilometers)
0
1
2
3
4
Time−lag (days)
0
5
10
15

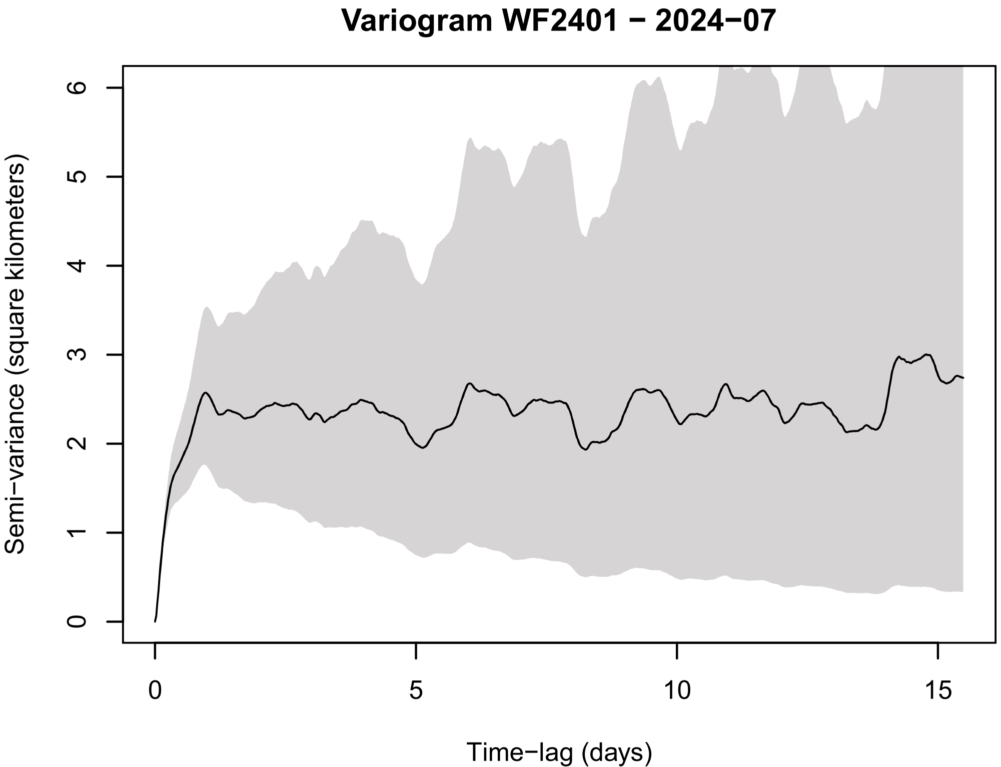
Variogram WF2401 − 2024−07
Semi−variance (square kilometers)
0
1
2
3
4
5
6
Time−lag (days)
0
5
10
15

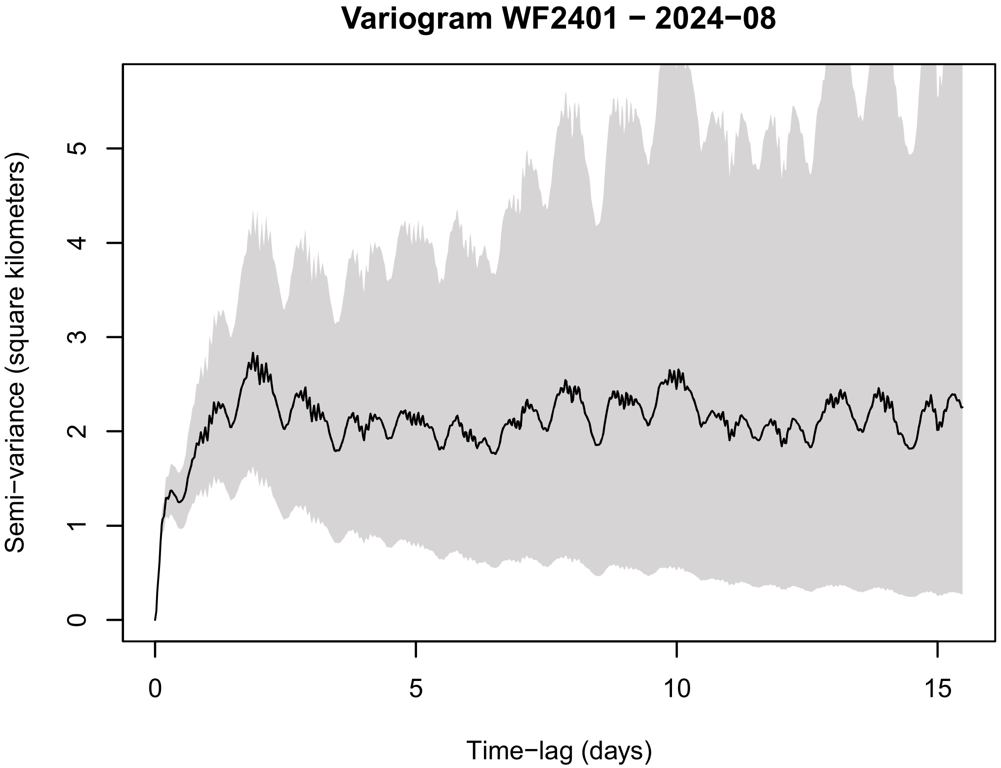
Variogram WF2401 − 2024−08
Semi−variance (square kilometers)
Time−lag (days)
0
1
2
3
4
5
0
5
10
15

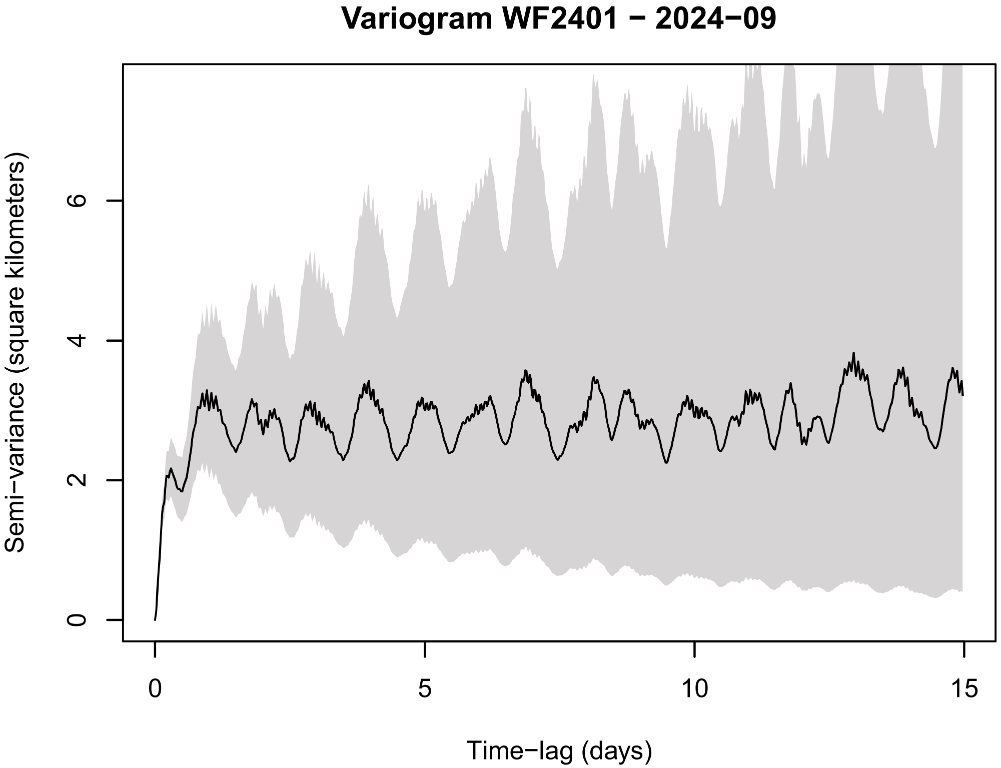
Variogram WF2401 − 2024−09
Semi−variance (square kilometers)
Time−lag (days)
0
2
4
6
0
5
10
15

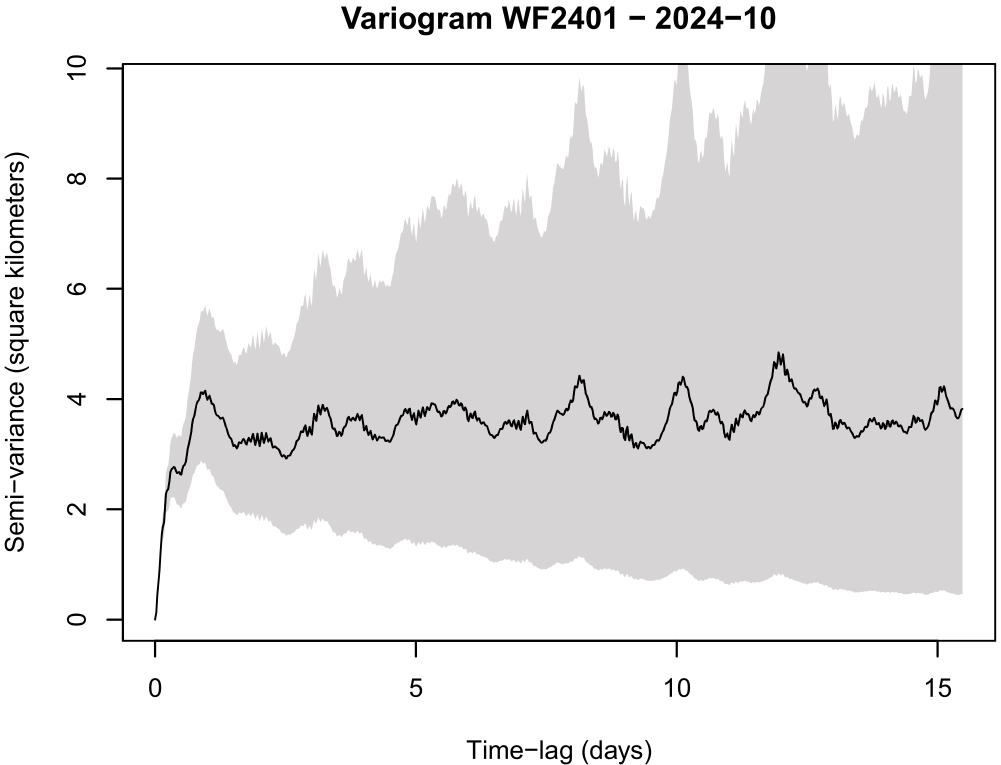
Variogram WF2401 − 2024−10
Semi−variance (square kilometers)
Time−lag (days)
0
2
4
6
8
10
0
5
10
15

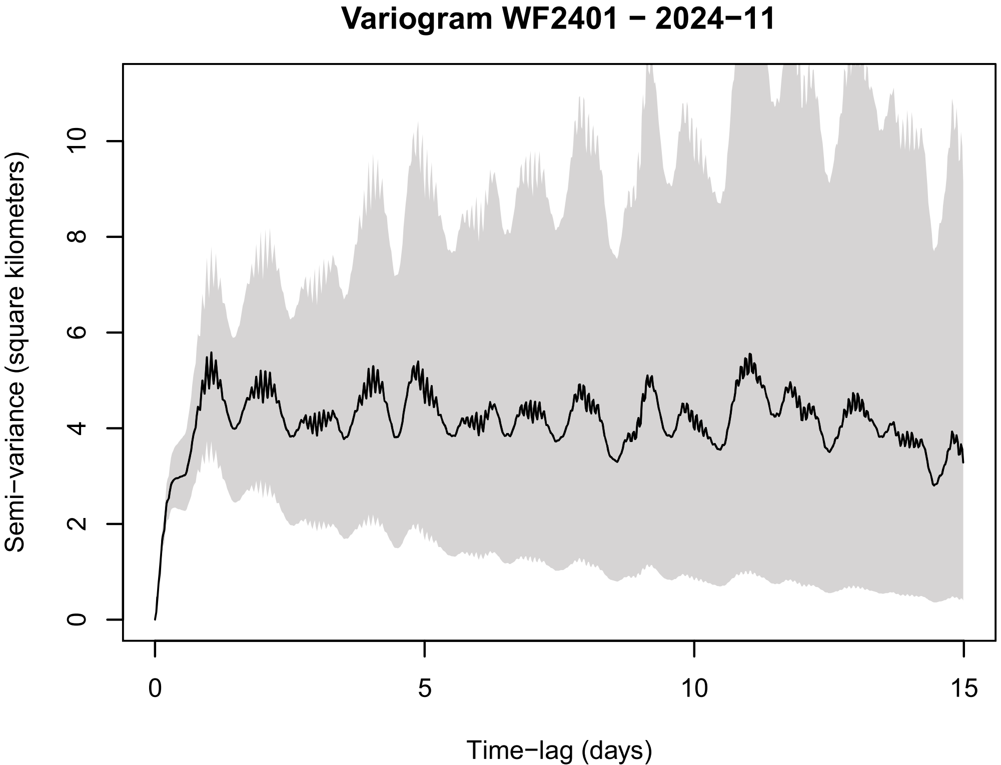
Variogram WF2401 − 2024−11
Semi−variance (square kilometers)
Time−lag (days)
0
2
4
6
8
10
0
5
10
15

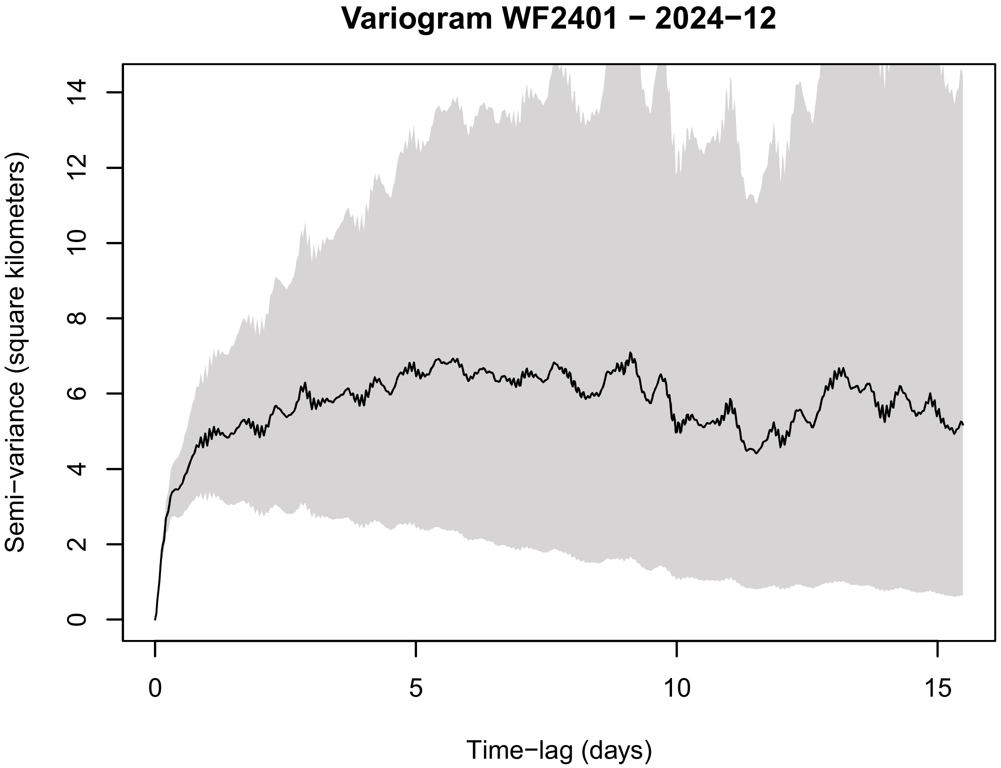
Variogram WF2401 − 2024−12
Semi−variance (square kilometers)
Time−lag (days)
0
2
4
6
8
10
12
14
0
5
10
15

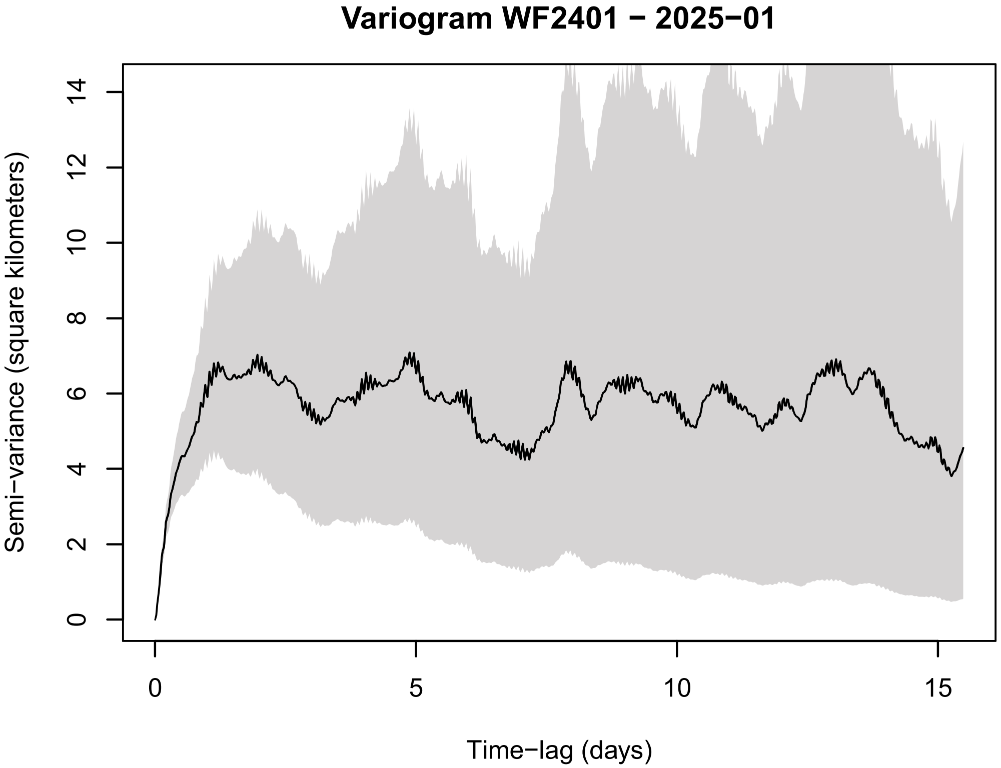
Variogram WF2401 − 2025−01
Semi−variance (square kilometers)
Time−lag (days)
0
2
4
6
8
10
12
14
0
5
10
15

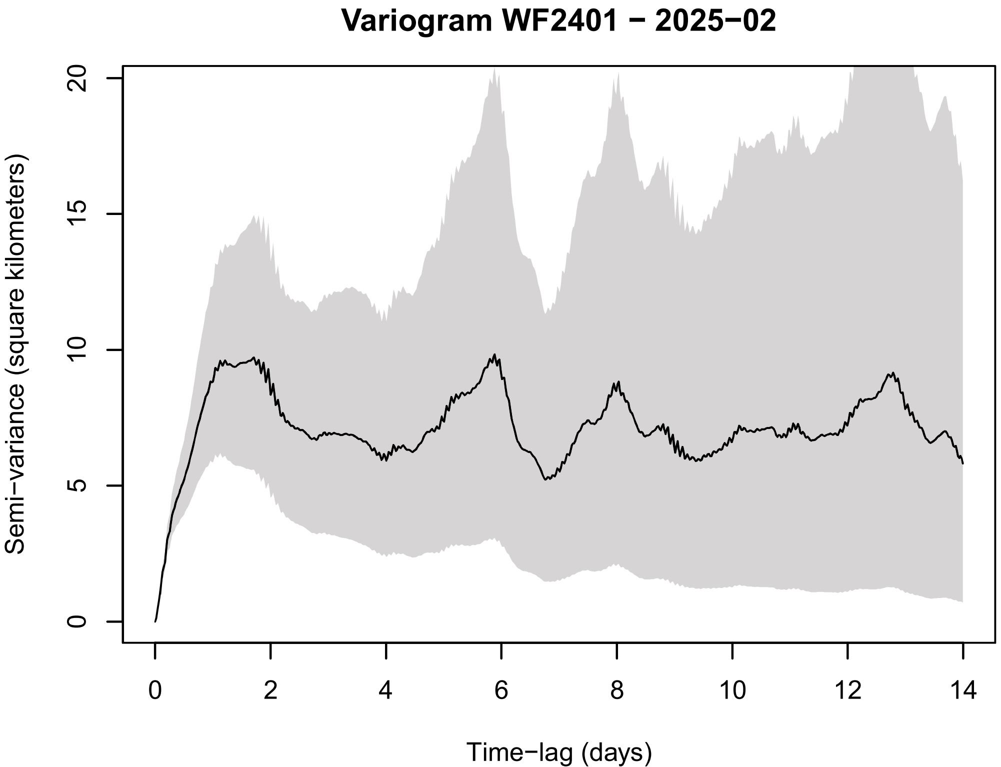
Variogram WF2401 − 2025−02
Semi−variance (square kilometers)
Time−lag (days)
0
5
10
15
20
0
2
4
6
8
10
12
14

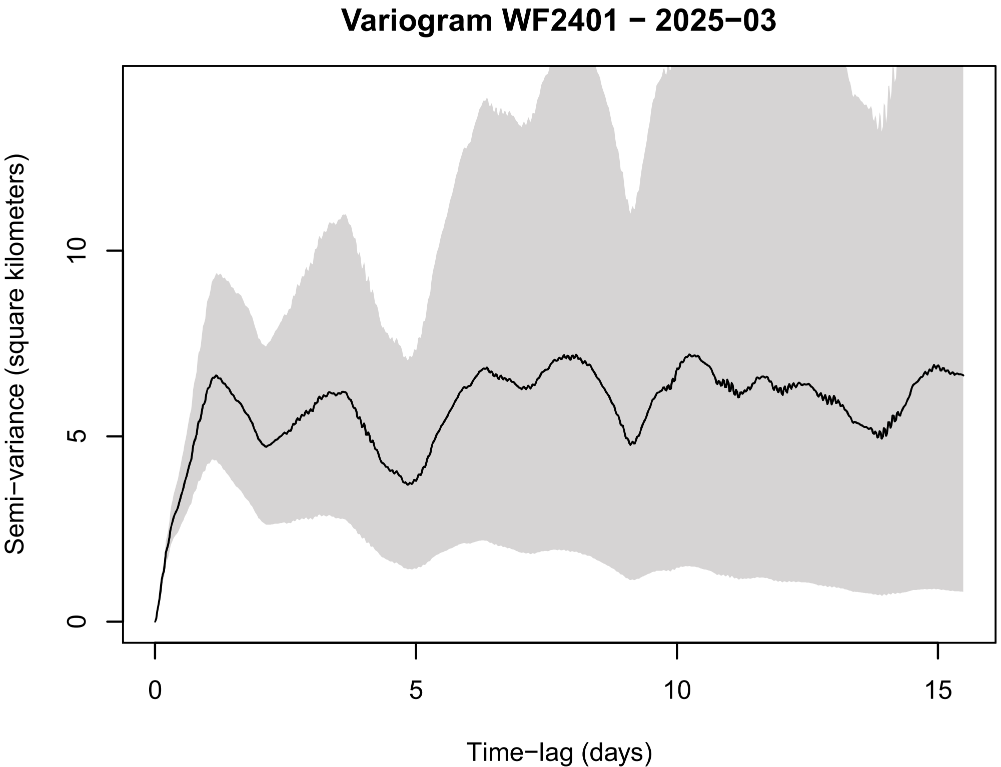
Variogram WF2401 − 2025−03
Semi−variance (square kilometers)
Time−lag (days)
0
5
10
0
5
10
15

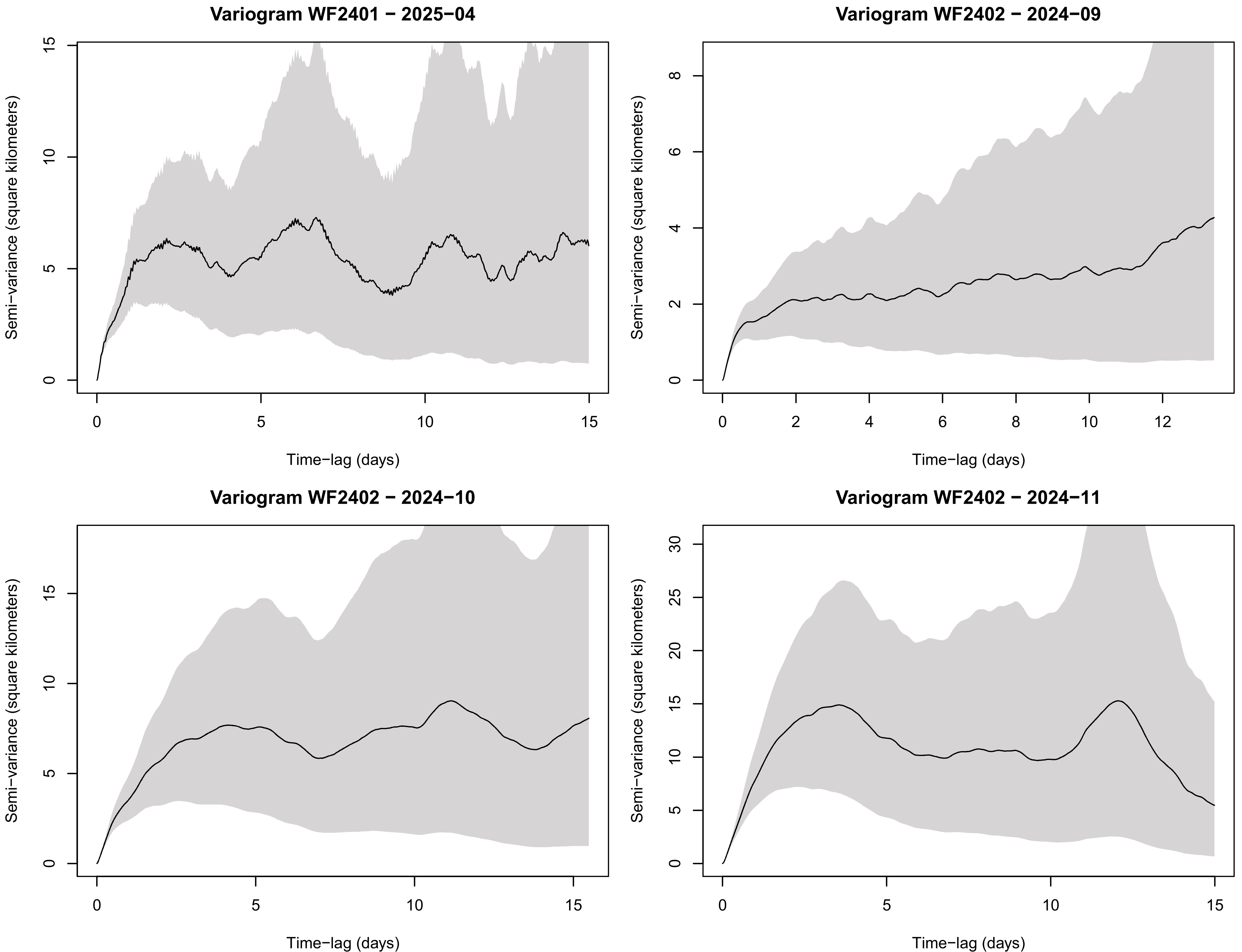
Variogram WF2401 − 2025−04
Variogram WF2402 − 2024−09
Variogram WF2402 − 2024−10
Variogram WF2402 − 2024−11
Semi−variance (square kilometers)
Time−lag (days)

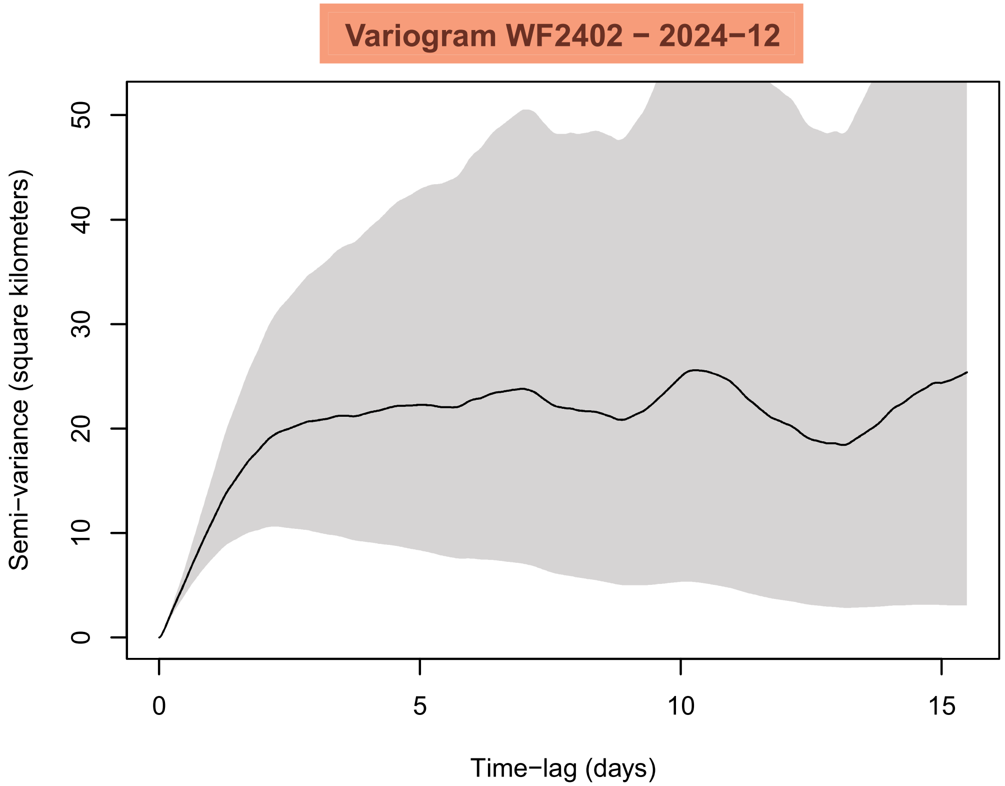
Variogram WF2402 − 2024−12
Semi−variance (square kilometers)
0
10
20
30
40
50
0
5
10
15
Time−lag (days)

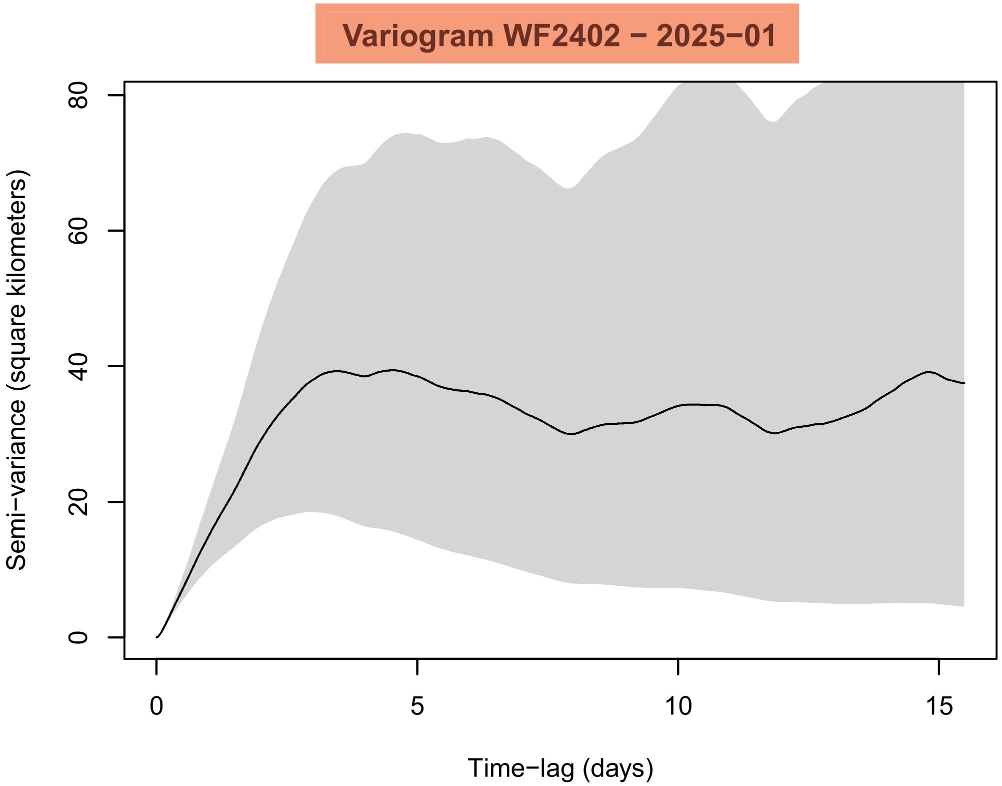
Variogram WF2402 − 2025−01
Semi−variance (square kilometers)
0
20
40
60
80
0
5
10
15
Time−lag (days)

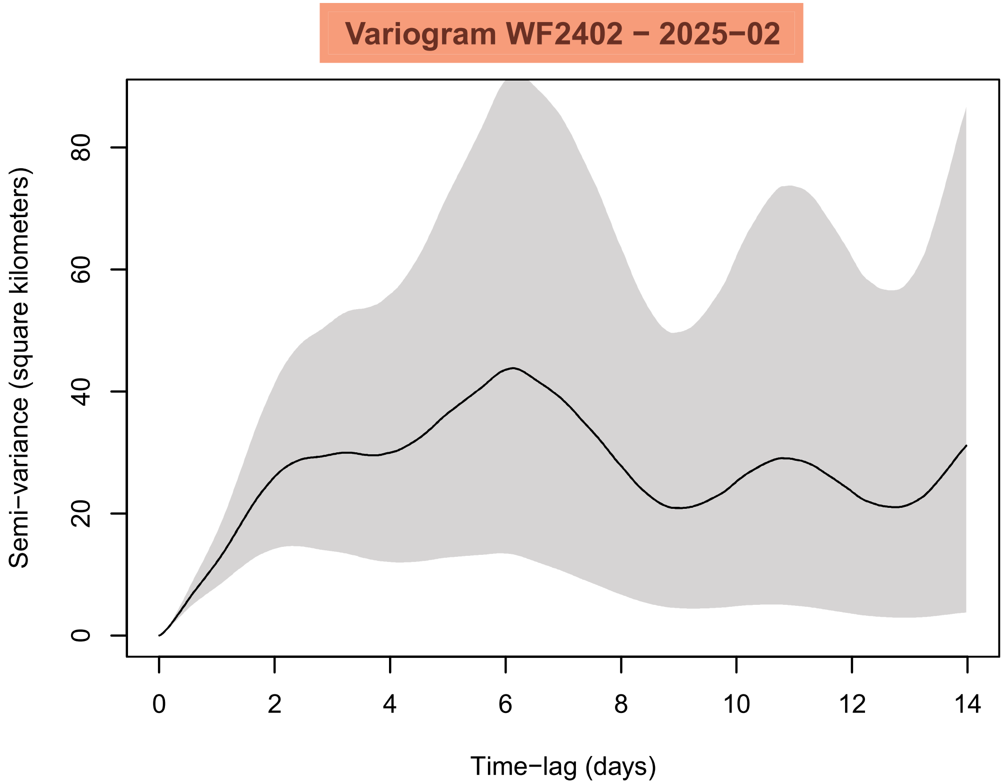
Variogram WF2402 − 2025−02
Semi−variance (square kilometers)
0
20
40
60
80
0
2
4
6
8
10
12
14
Time−lag (days)

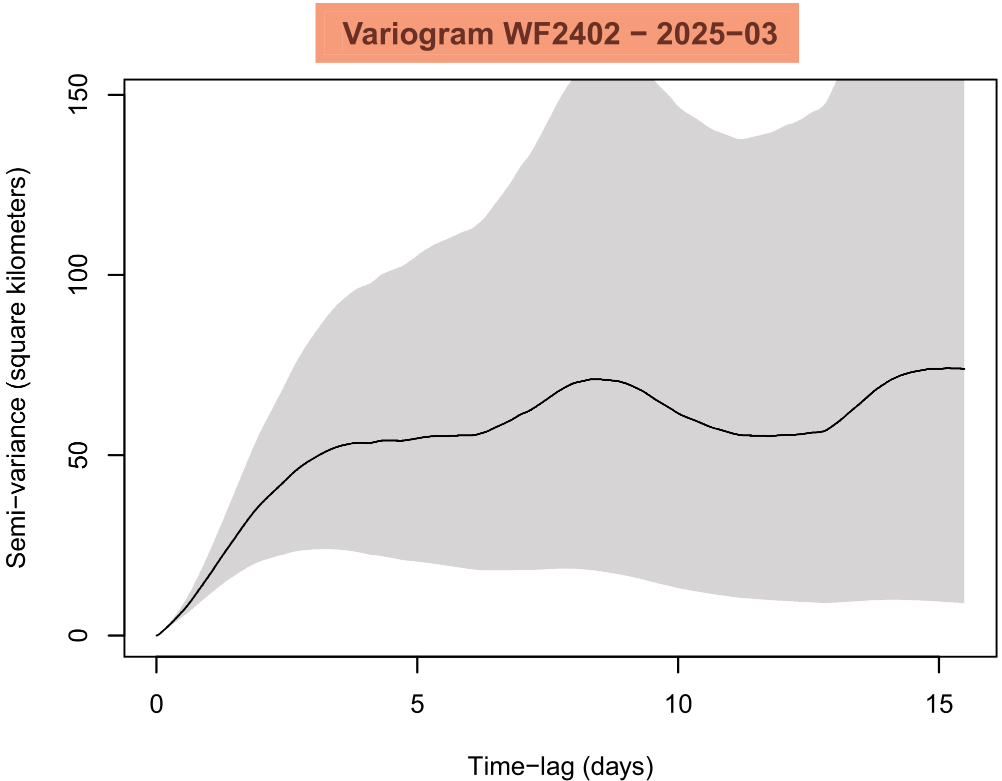
Variogram WF2402 − 2025−03
Semi−variance (square kilometers)
0
50
100
150
0
5
10
15
Time−lag (days)

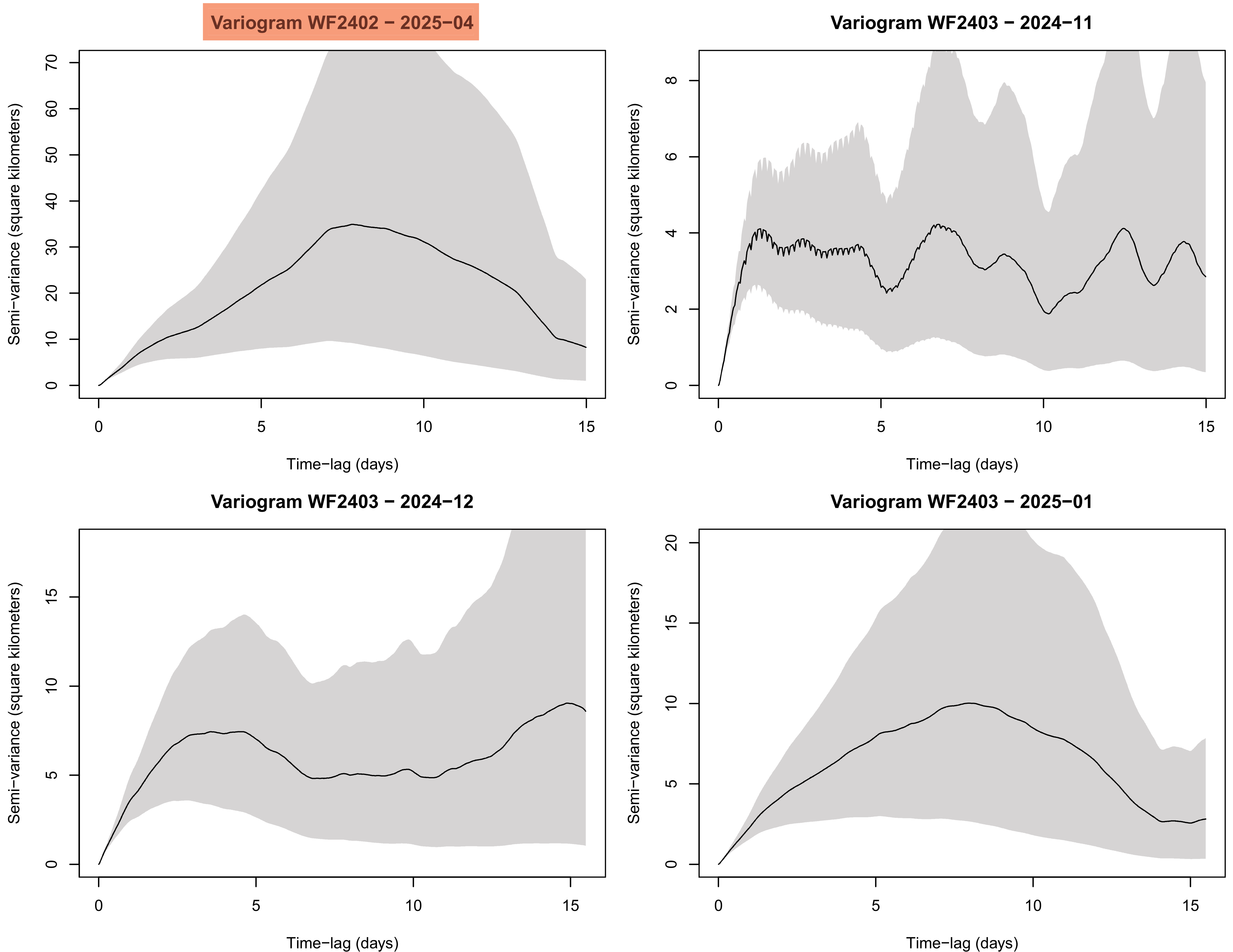

Variogram WF2402 − 2025−04
Semi−variance (square kilometers)
Time−lag (days)
Variogram WF2403 − 2024−11
Semi−variance (square kilometers)
Time−lag (days)
Variogram WF2403 − 2024−12
Semi−variance (square kilometers)
Time−lag (days)
Variogram WF2403 − 2025−01
Semi−variance (square kilometers)
Time−lag (days)

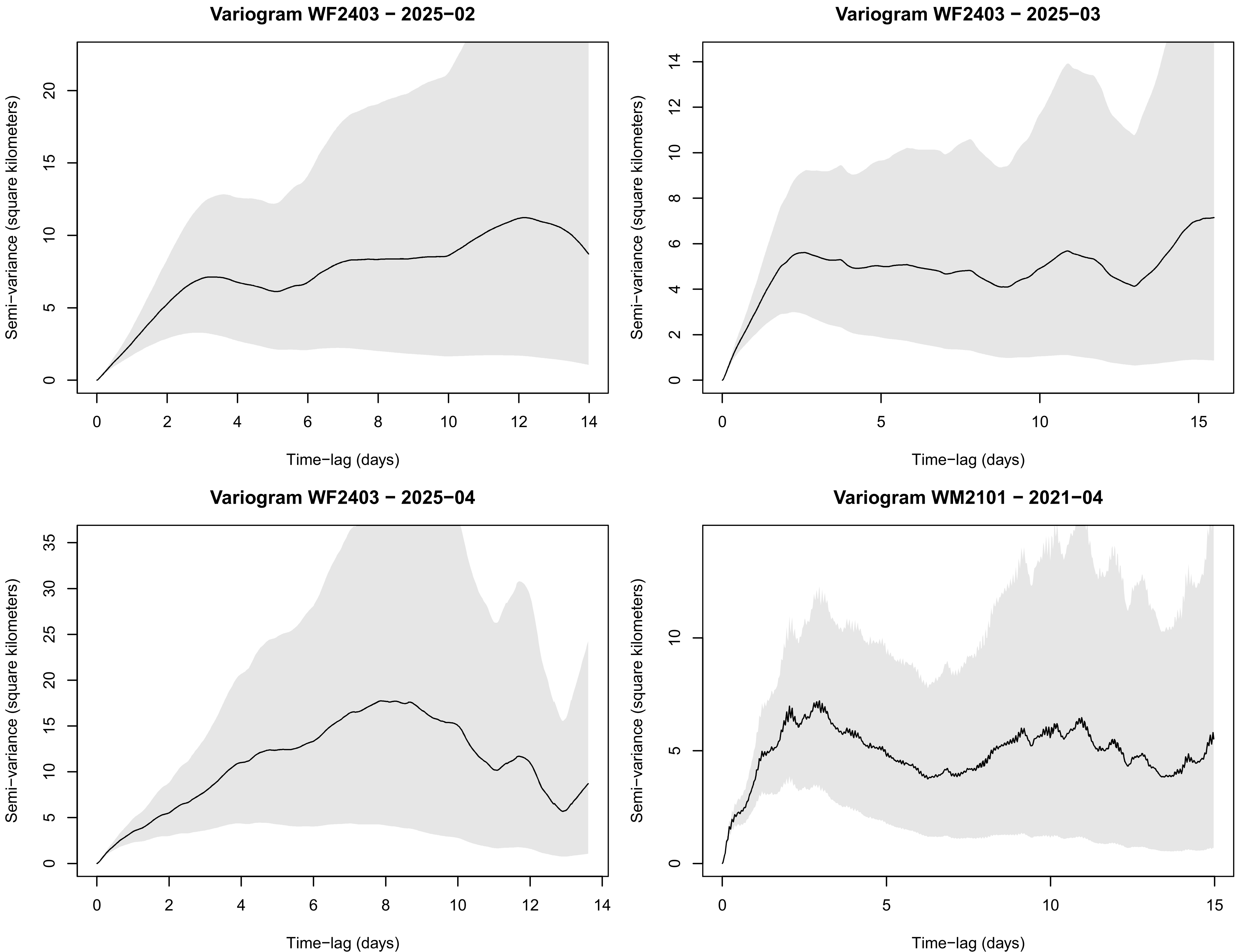
Variogram WF2403 − 2025−02
Semi−variance (square kilometers)
0
5
10
15
20
0
2
4
6
8
10
12
14
Time−lag (days)
Variogram WF2403 − 2025−03
Semi−variance (square kilometers)
0
2
4
6
8
10
12
14
0
5
10
15
Time−lag (days)
Variogram WF2403 − 2025−04
Semi−variance (square kilometers)
0
5
10
15
20
25
30
35
0
2
4
6
8
10
12
14
Time−lag (days)
Variogram WM2101 − 2021−04
Semi−variance (square kilometers)
0
5
10
0
5
10
15
Time−lag (days)

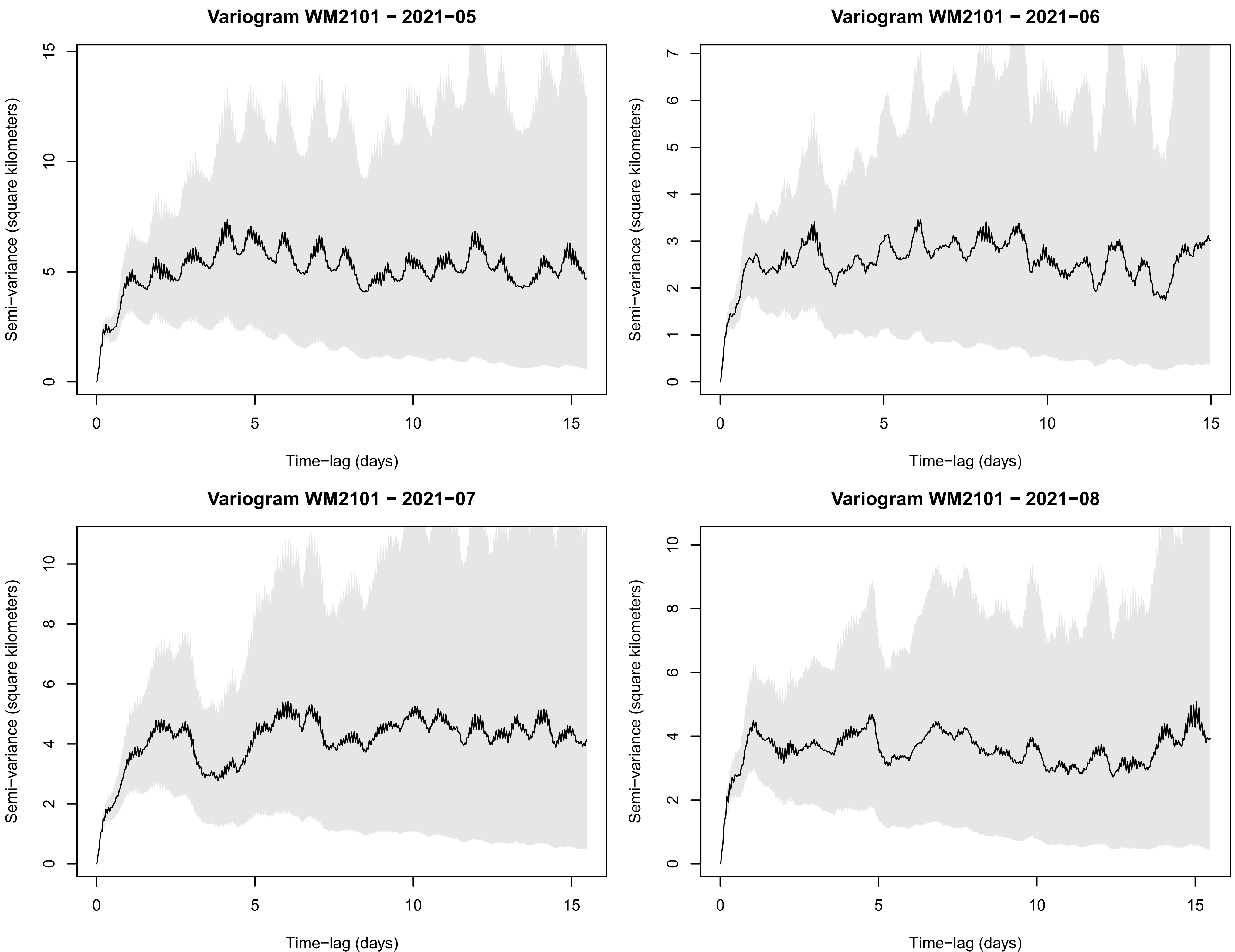
Variogram WM2101 − 2021−05
Semi−variance (square kilometers)
Time−lag (days)
Variogram WM2101 − 2021−06
Semi−variance (square kilometers)
Time−lag (days)
Variogram WM2101 − 2021−07
Semi−variance (square kilometers)
Time−lag (days)
Variogram WM2101 − 2021−08
Semi−variance (square kilometers)
Time−lag (days)

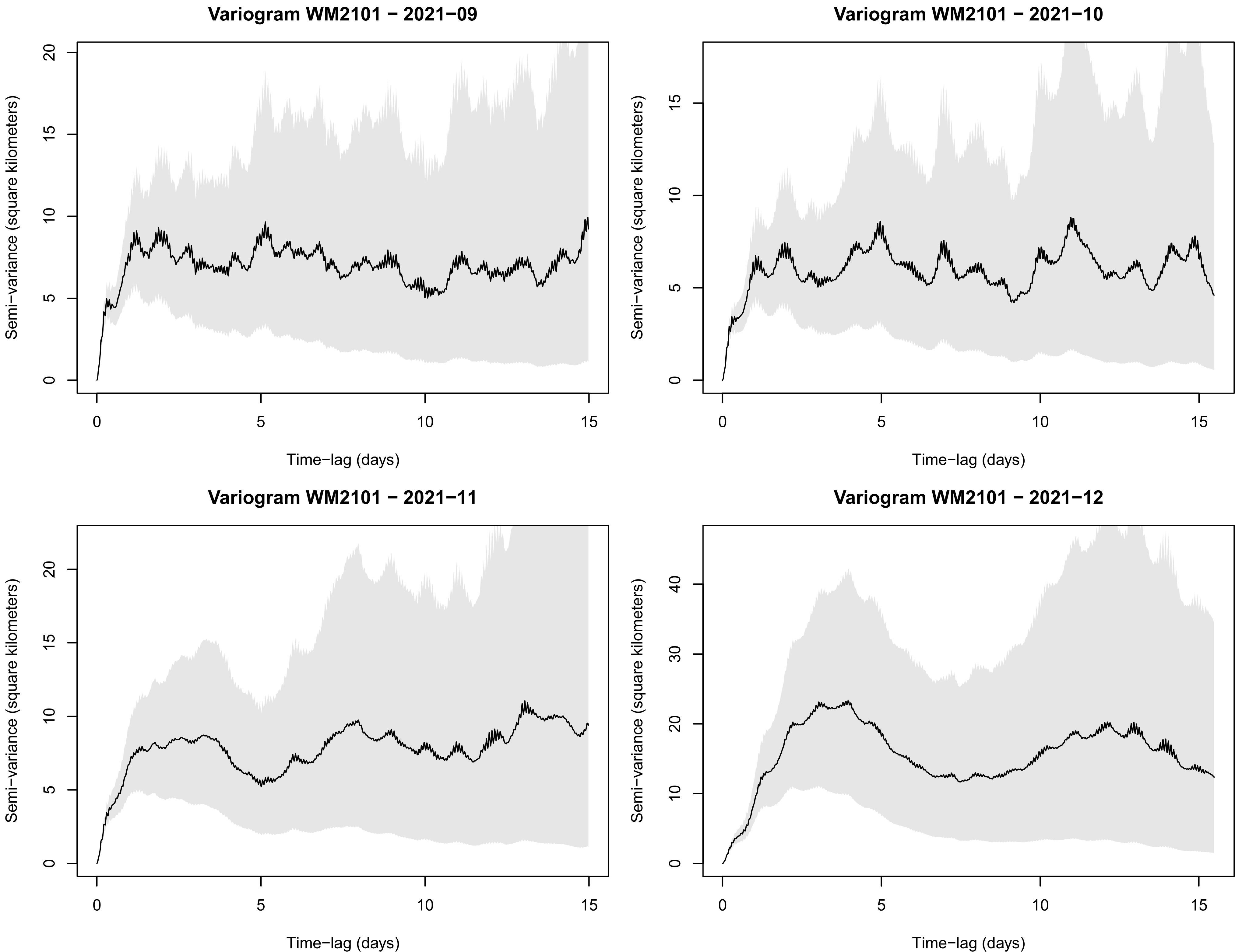

Variogram WM2101 − 2021−09
Variogram WM2101 − 2021−10
Variogram WM2101 − 2021−11
Variogram WM2101 − 2021−12
Semi−variance (square kilometers)
Time−lag (days)

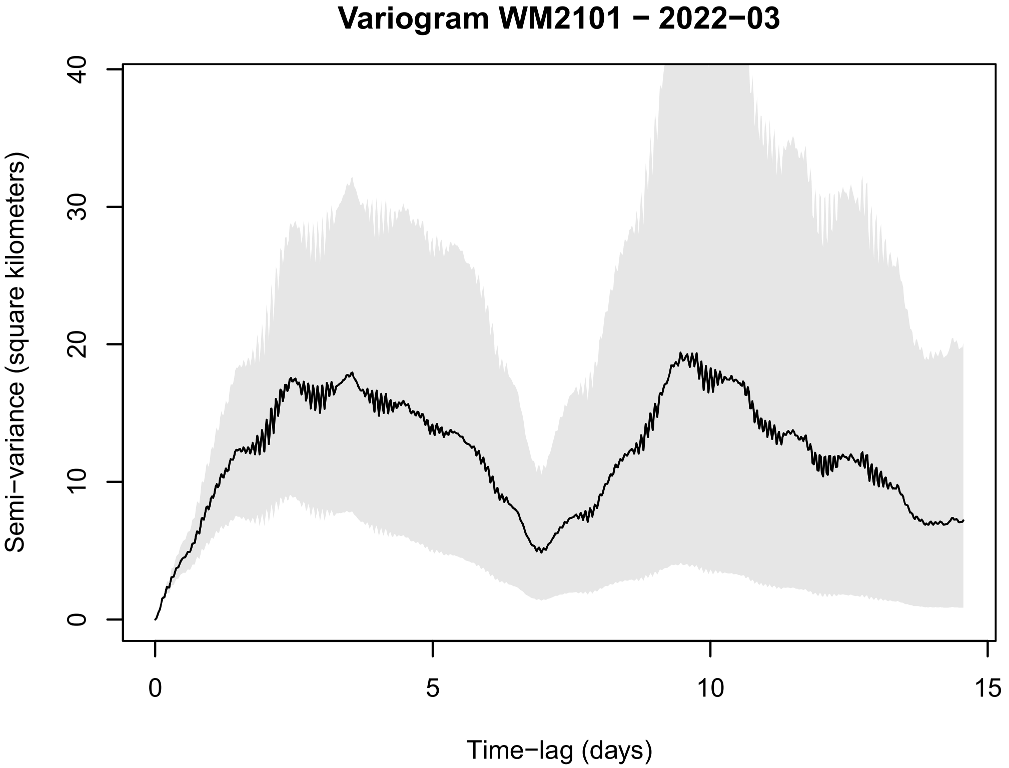
Variogram WM2101 − 2022−03
Semi−variance (square kilometers)
0
10
20
30
40
0
5
10
15
Time−lag (days)
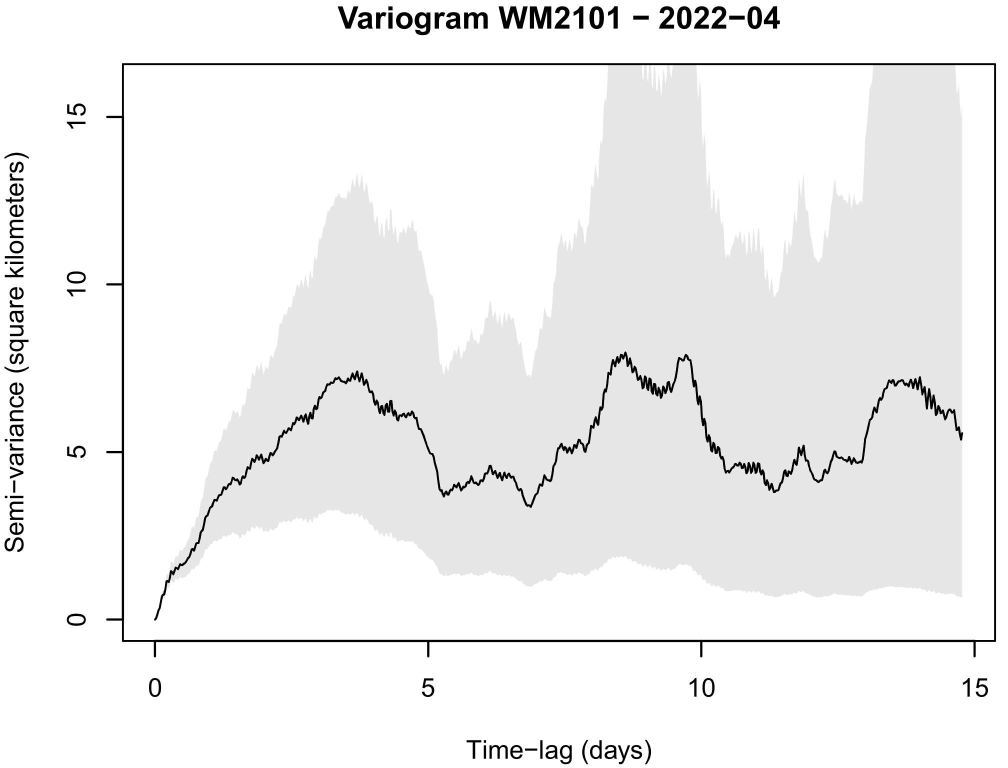
Variogram WM2101 − 2022−04
Semi−variance (square kilometers)
0
5
10
15
0
5
10
15
Time−lag (days)
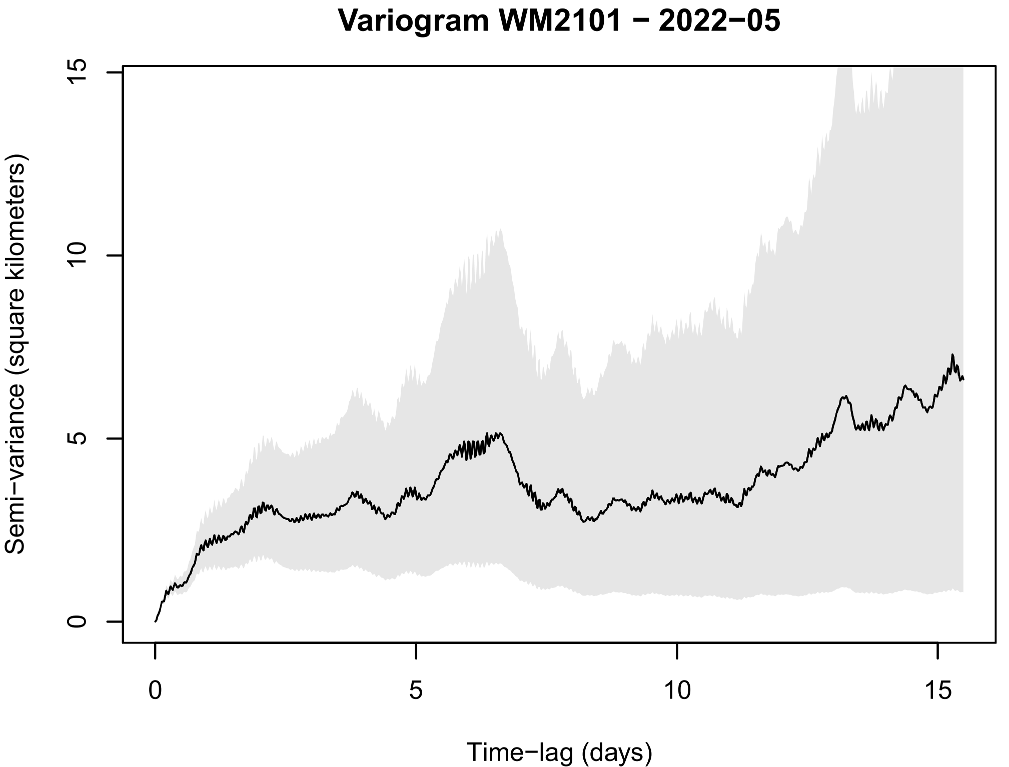
Variogram WM2101 − 2022−05
Semi−variance (square kilometers)
0
5
10
15
0
5
10
15
Time−lag (days)
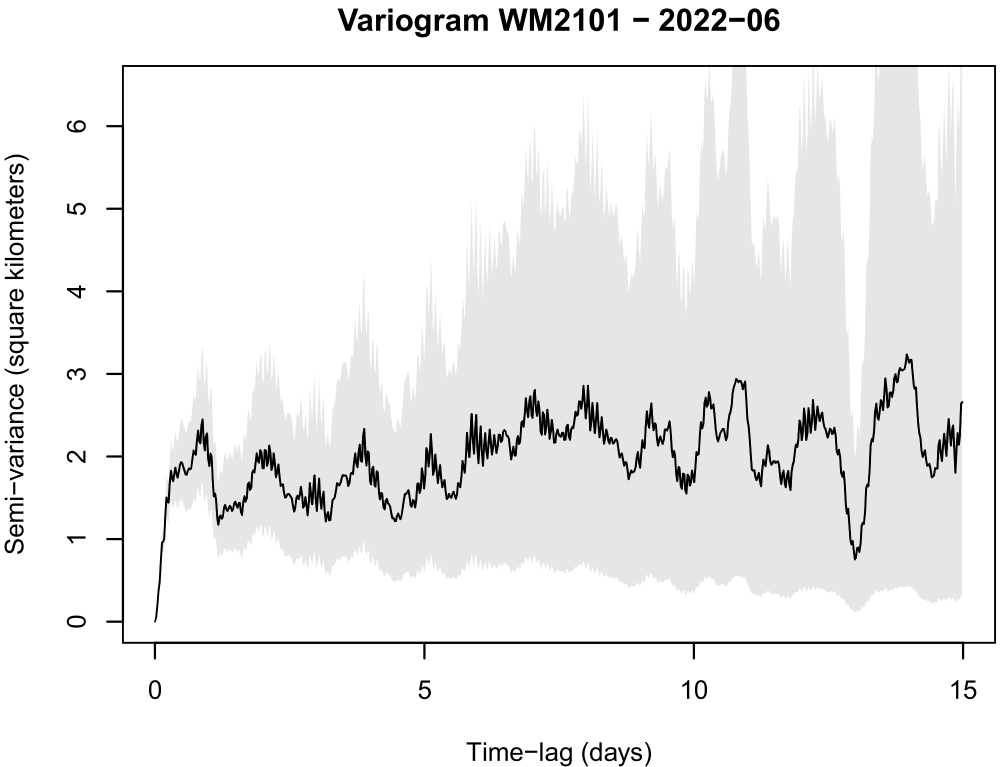
Variogram WM2101 − 2022−06
Semi−variance (square kilometers)
0
1
2
3
4
5
6
0
5
10
15
Time−lag (days)

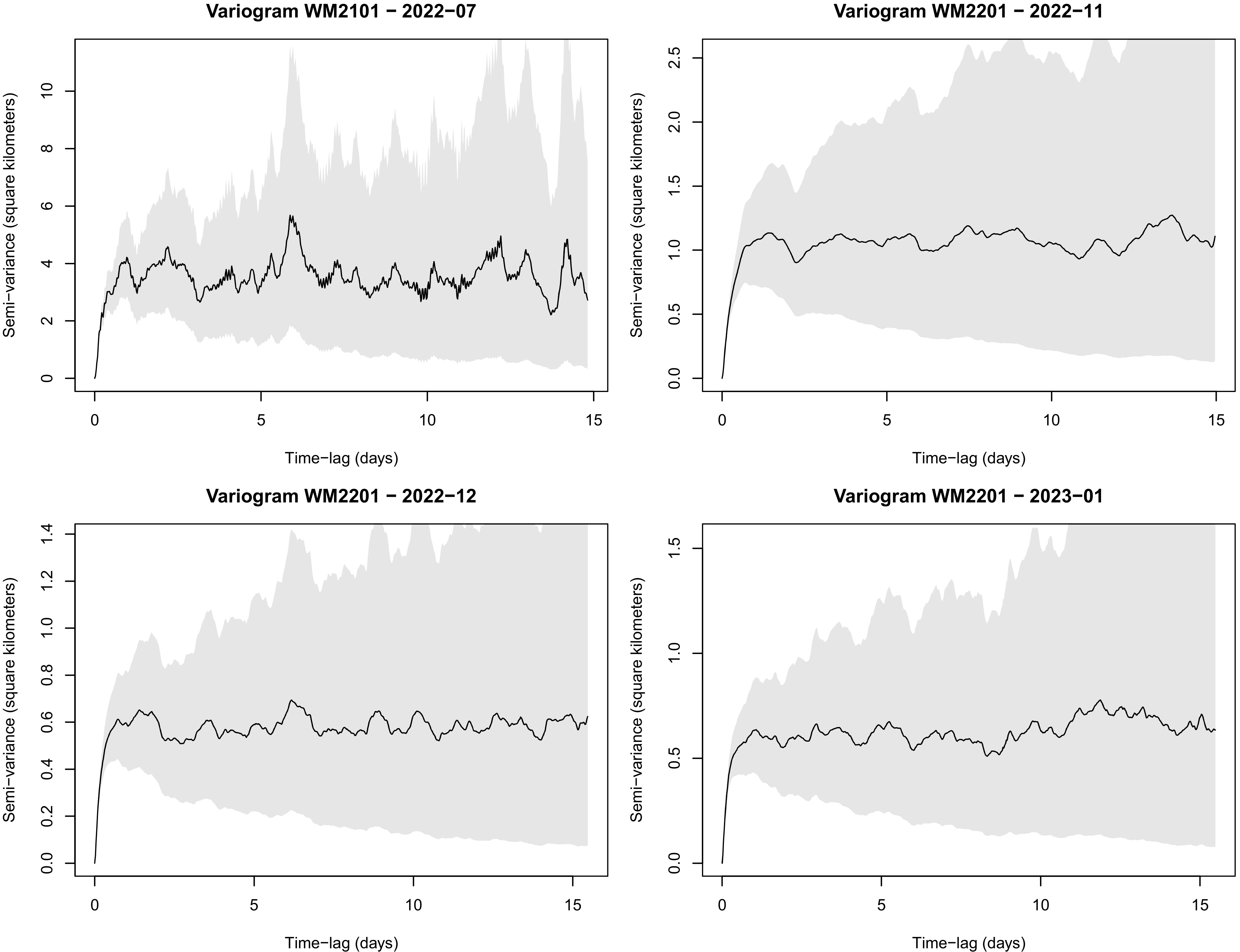
Variogram WM2101 − 2022−07
Semi−variance (square kilometers)
Time−lag (days)
Variogram WM2201 − 2022−11
Semi−variance (square kilometers)
Time−lag (days)
Variogram WM2201 − 2022−12
Semi−variance (square kilometers)
Time−lag (days)
Variogram WM2201 − 2023−01
Semi−variance (square kilometers)
Time−lag (days)

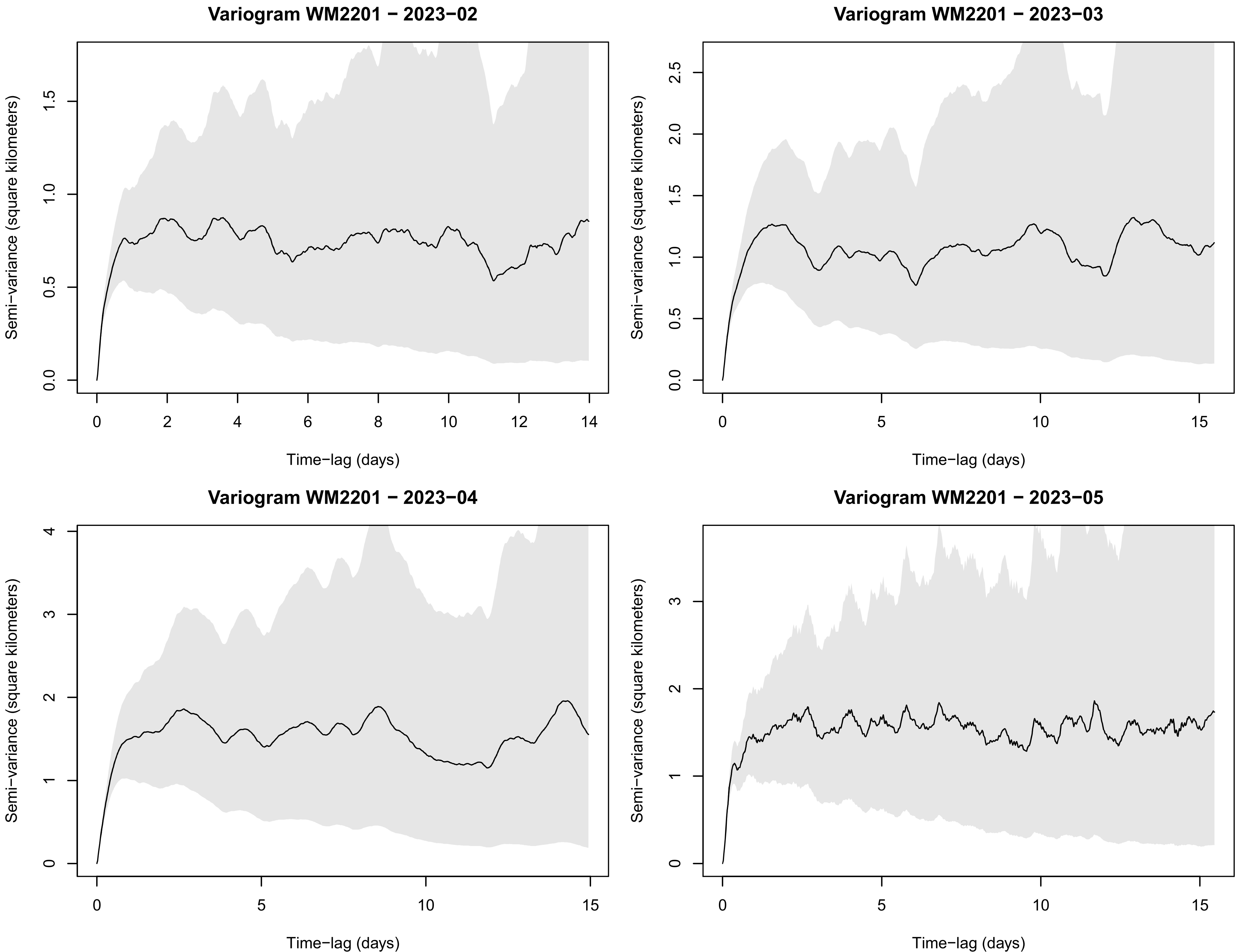
Variogram WM2201 − 2023−02
Semi−variance (square kilometers)
Time−lag (days)
Variogram WM2201 − 2023−03
Semi−variance (square kilometers)
Time−lag (days)
Variogram WM2201 − 2023−04
Semi−variance (square kilometers)
Time−lag (days)
Variogram WM2201 − 2023−05
Semi−variance (square kilometers)
Time−lag (days)

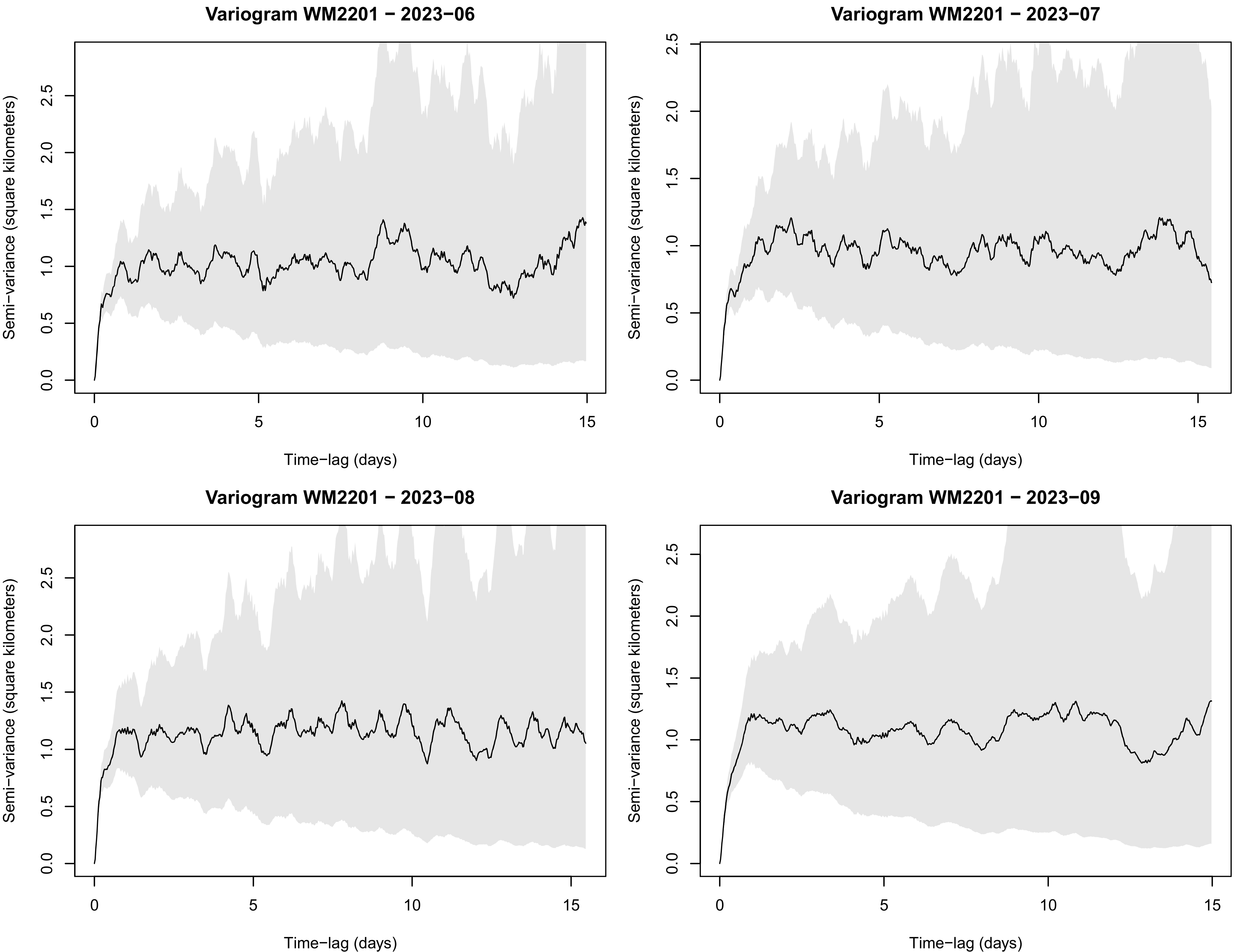
Variogram WM2201 − 2023−06
Variogram WM2201 − 2023−07
Variogram WM2201 − 2023−08
Variogram WM2201 − 2023−09
Semi−variance (square kilometers)
Time−lag (days)
0.0
0.5
1.0
1.5
2.0
2.5
0
5
10
15

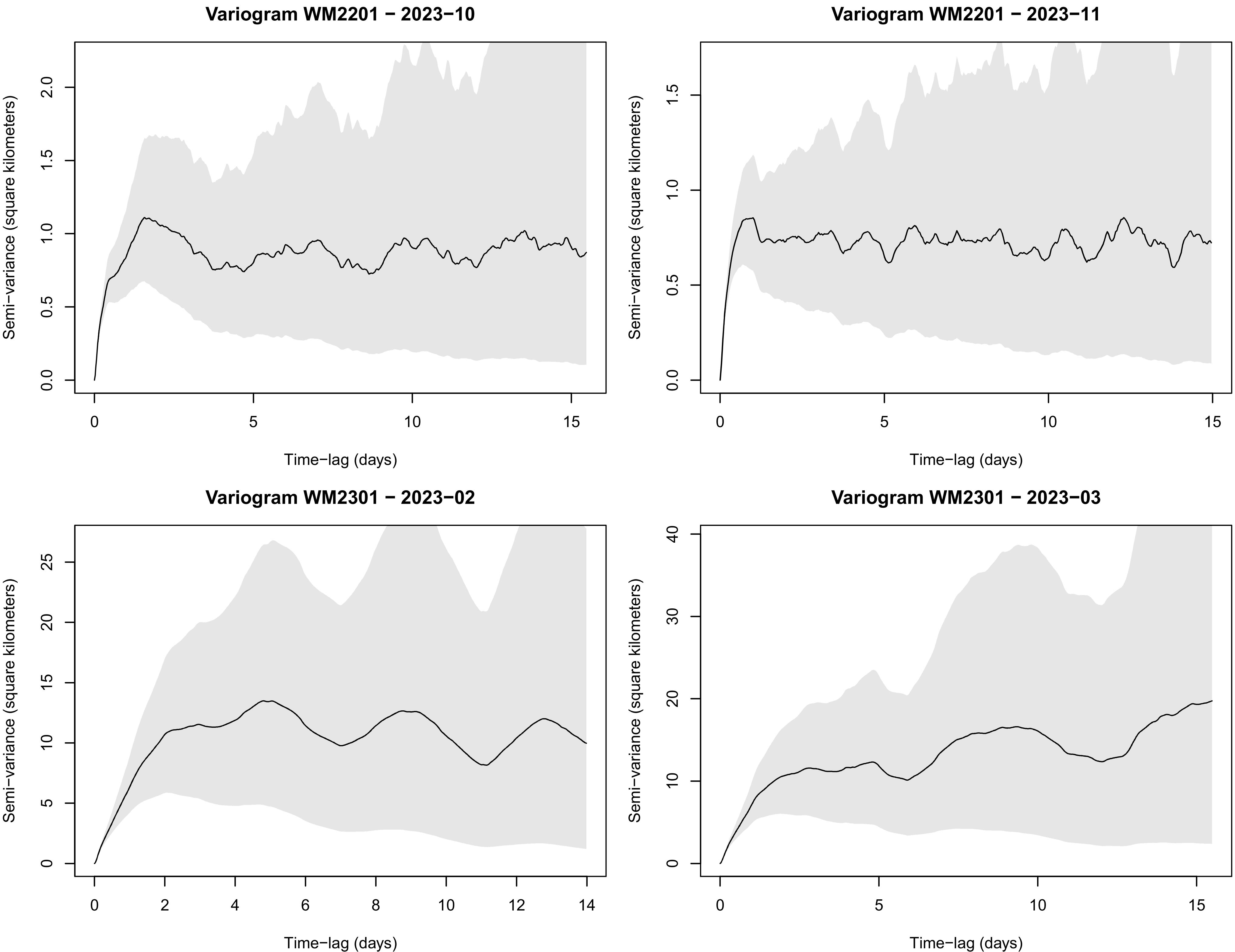

Variogram WM2201 − 2023−10
Variogram WM2201 − 2023−11
Variogram WM2301 − 2023−02
Variogram WM2301 − 2023−03
Semi−variance (square kilometers)
Time−lag (days)

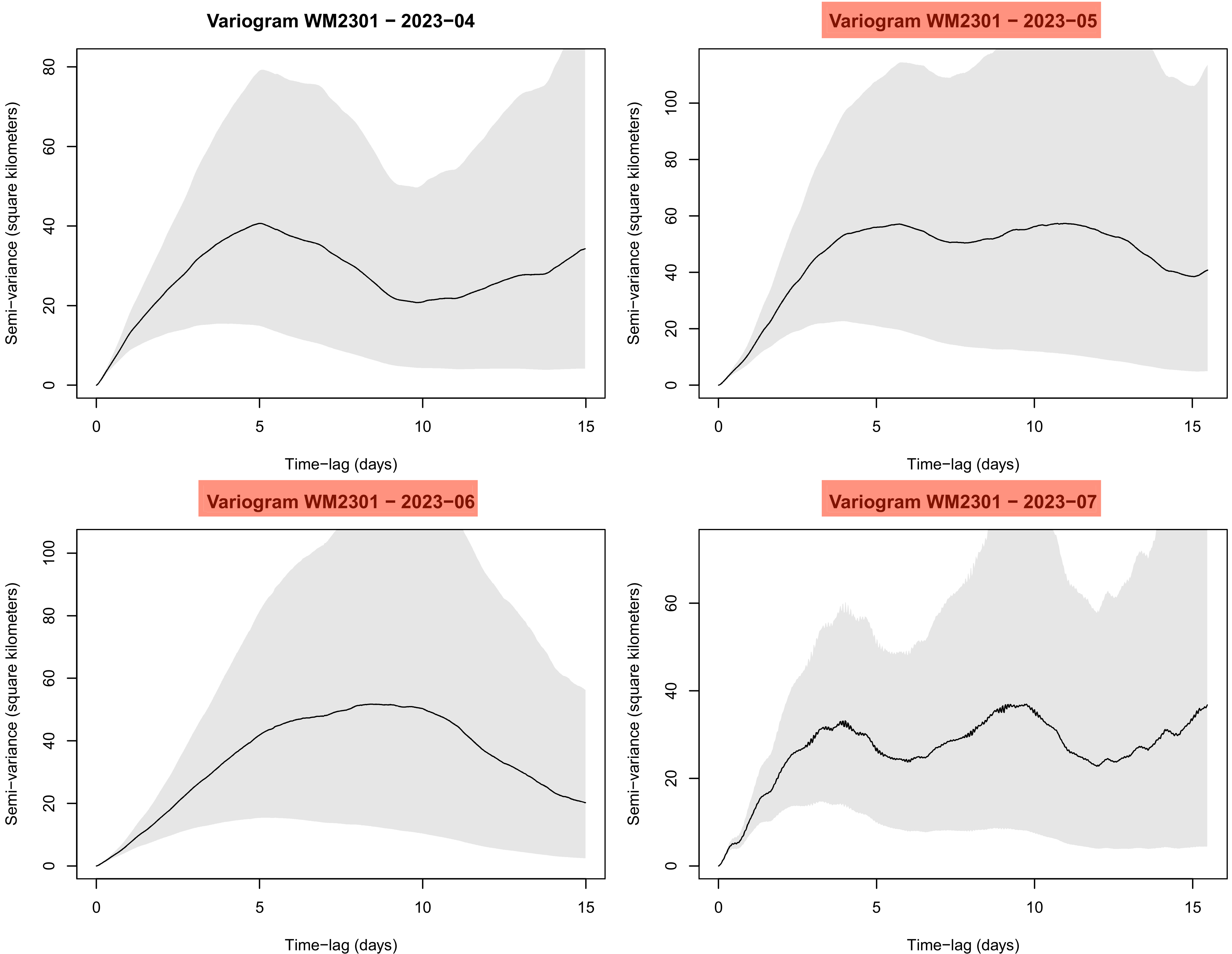
Variogram WM2301 − 2023−04
Variogram WM2301 − 2023−05
Variogram WM2301 − 2023−06
Variogram WM2301 − 2023−07
Semi−variance (square kilometers)
Time−lag (days)
0
5
10
15
0
20
40
60
80
100

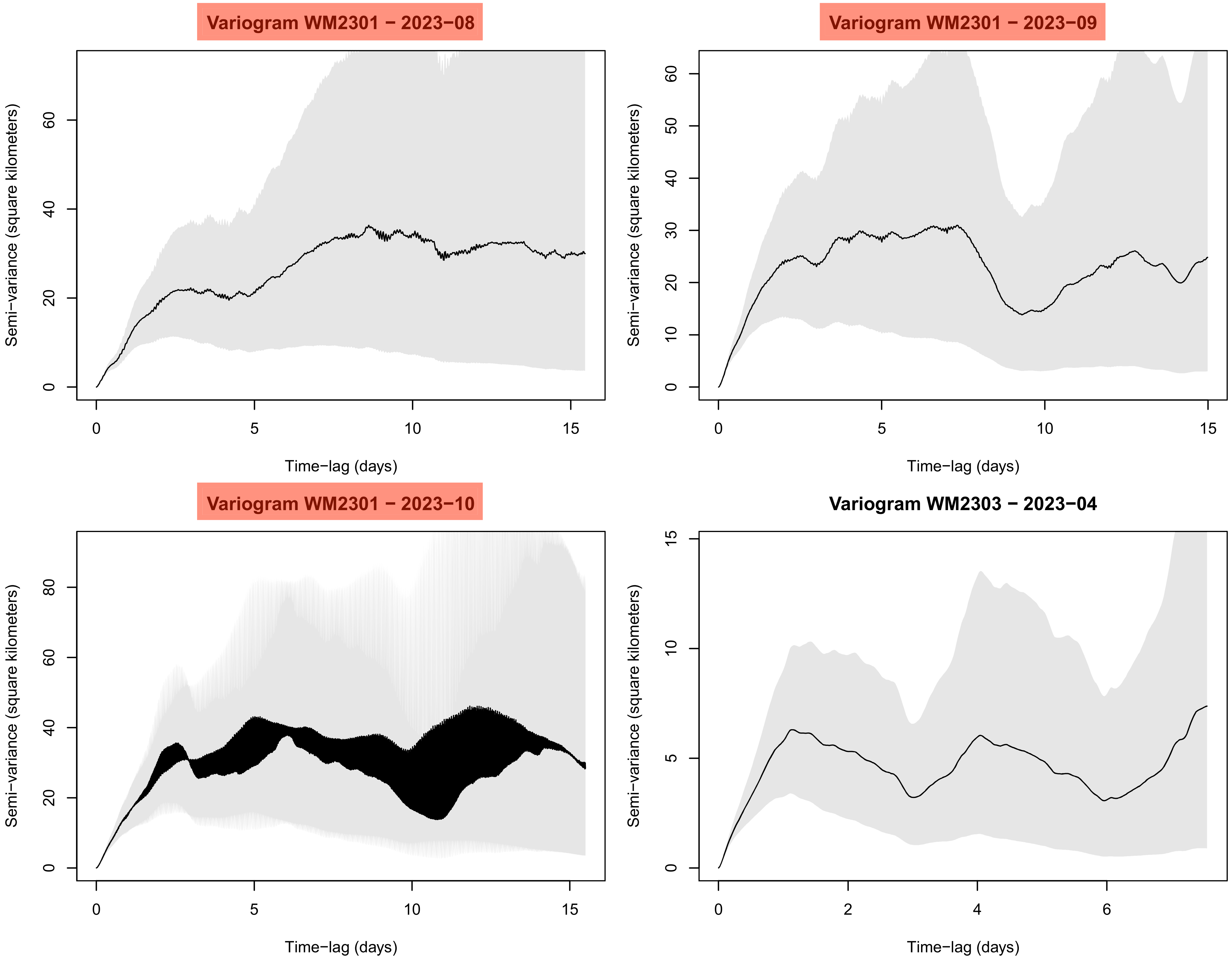
Variogram WM2301 − 2023−08
Semi−variance (square kilometers)
Time−lag (days)
Variogram WM2301 − 2023−09
Semi−variance (square kilometers)
Time−lag (days)
Variogram WM2301 − 2023−10
Semi−variance (square kilometers)
Time−lag (days)
Variogram WM2303 − 2023−04
Semi−variance (square kilometers)
Time−lag (days)

Variogram WM2303 − 2023−05

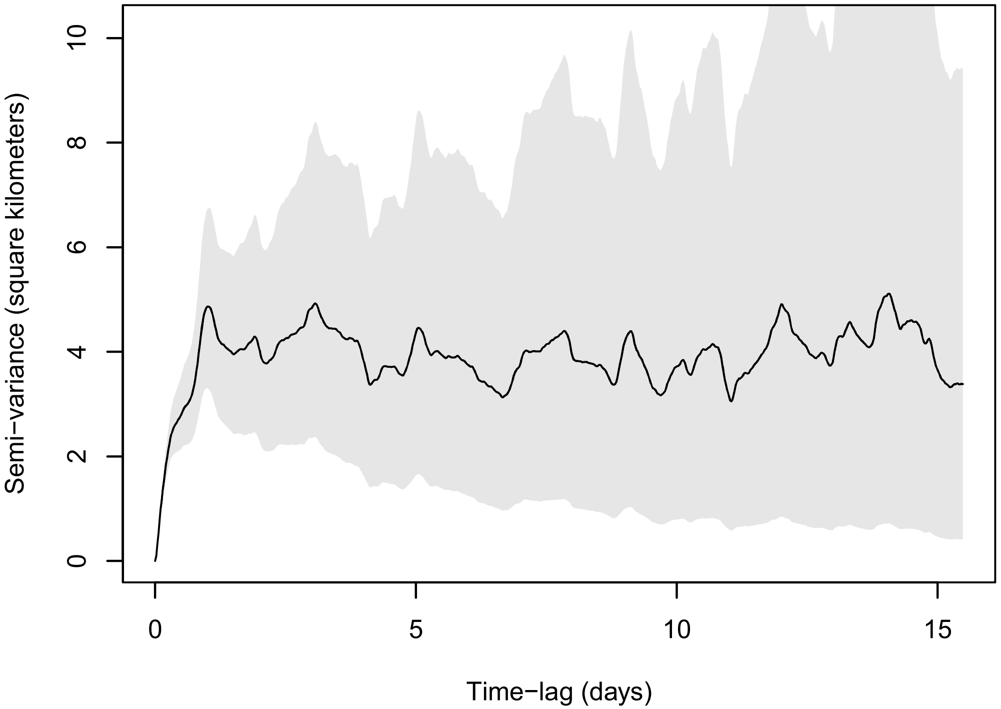


Variogram WM2303 − 2023−06

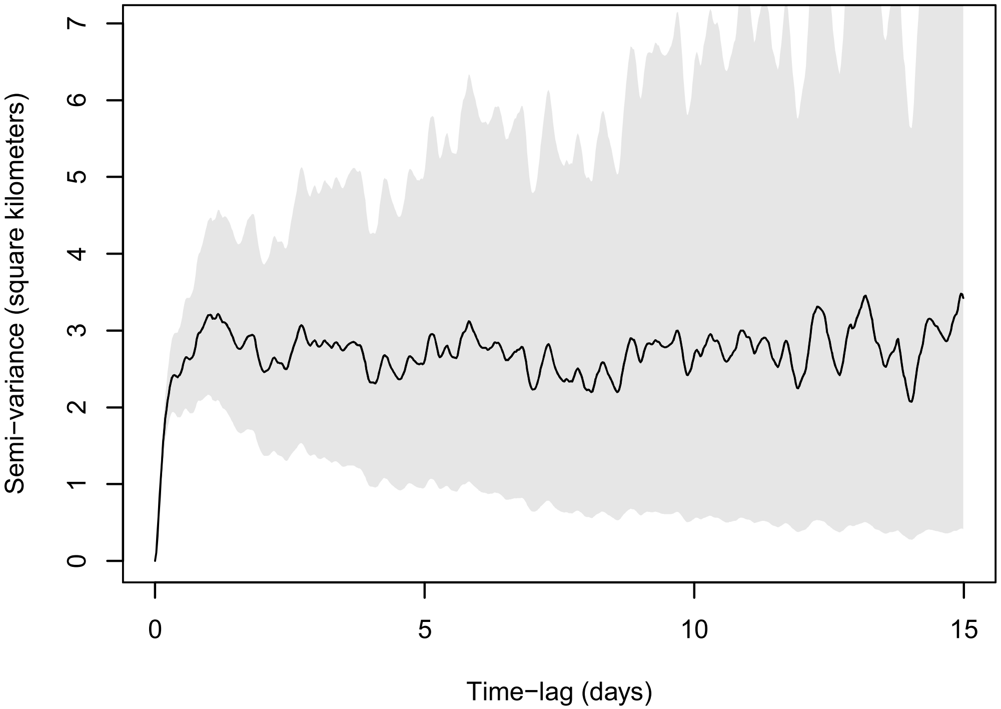


Variogram WM2303 − 2023−07

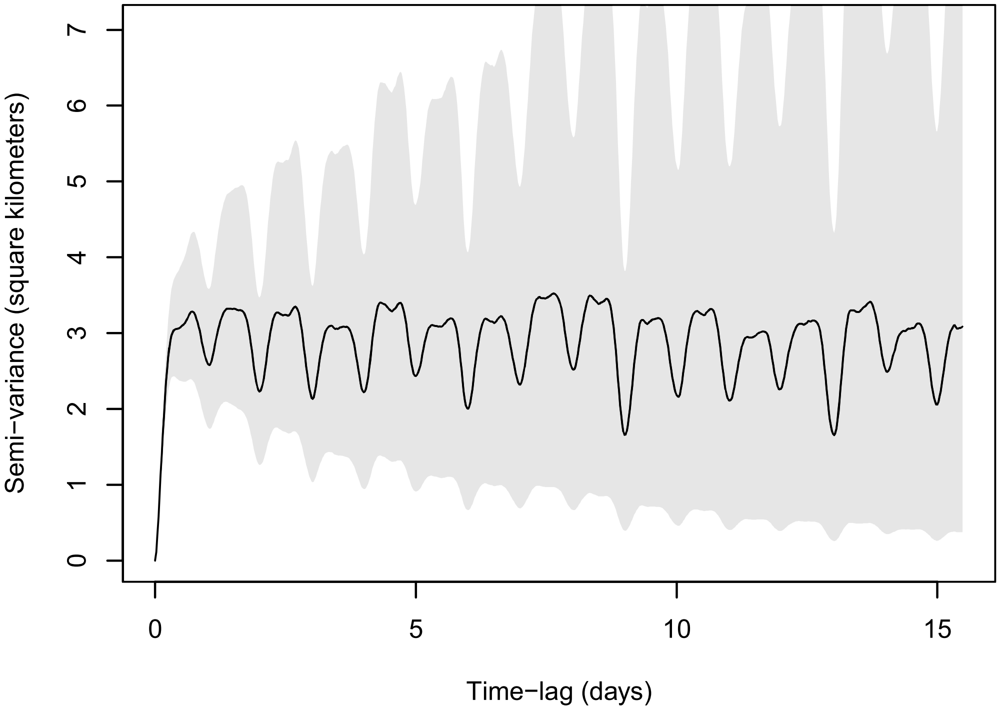


Variogram WM2303 − 2023−08

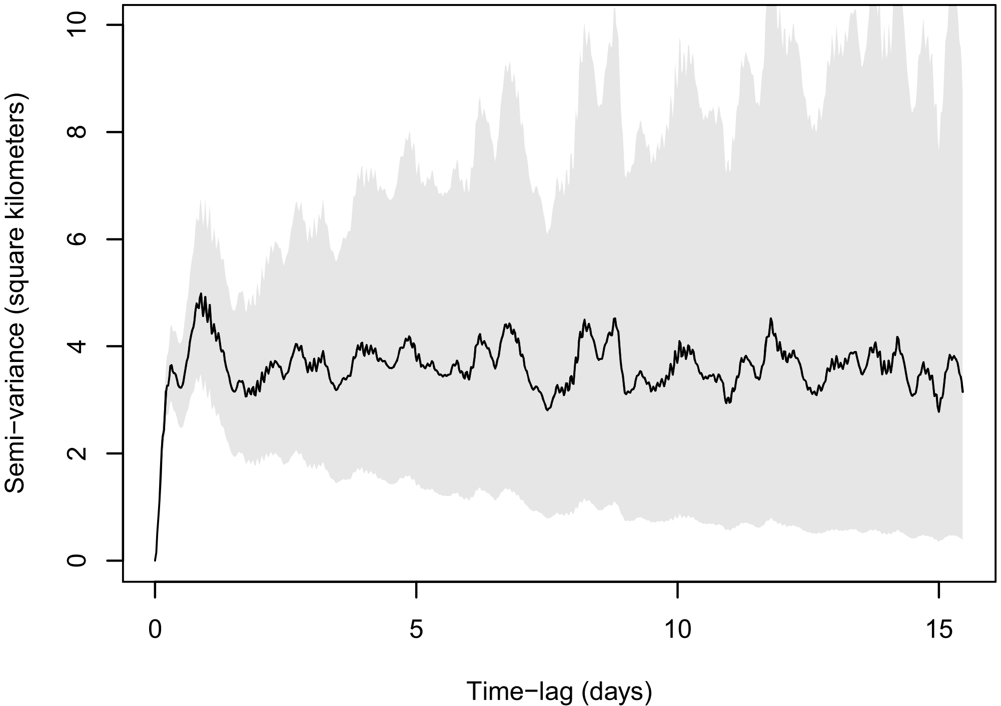

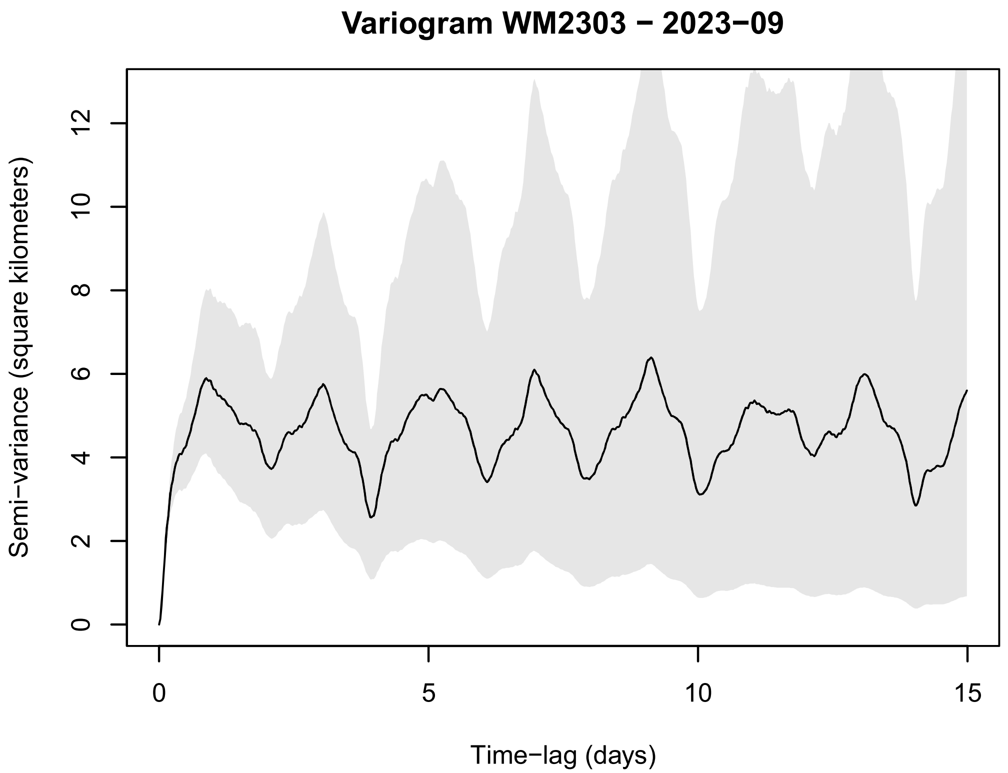
Variogram WM2303 − 2023−09
Semi−variance (square kilometers)
0
2
4
6
8
10
12
Time−lag (days)
0
5
10
15

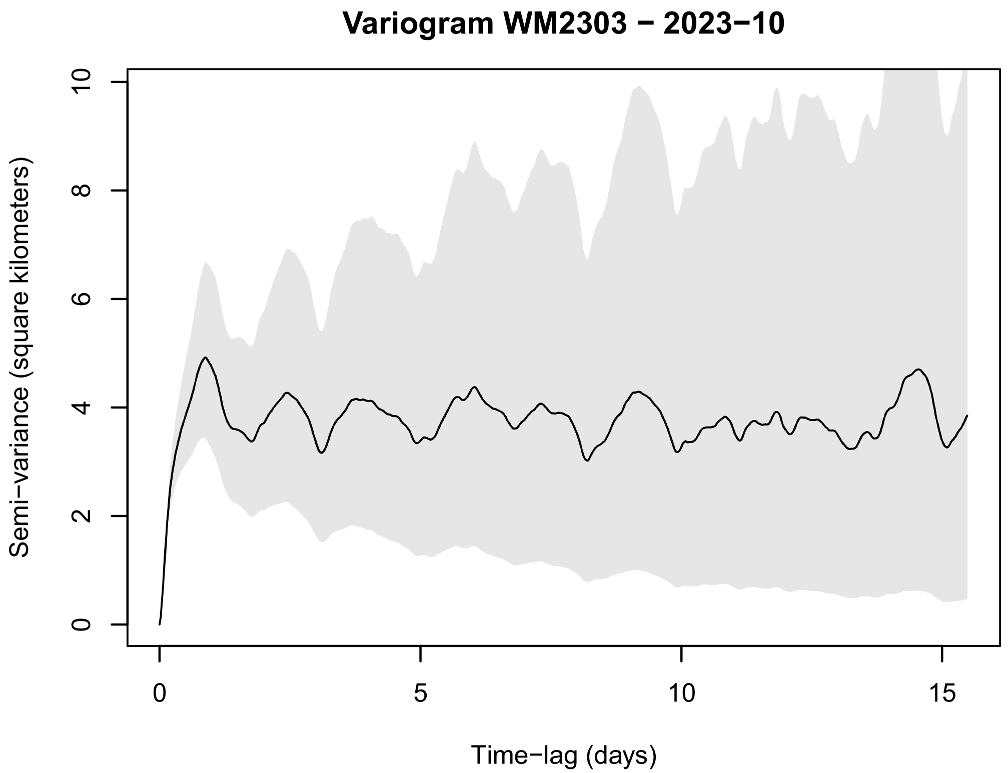
Variogram WM2303 − 2023−10
Semi−variance (square kilometers)
0
2
4
6
8
10
Time−lag (days)
0
5
10
15

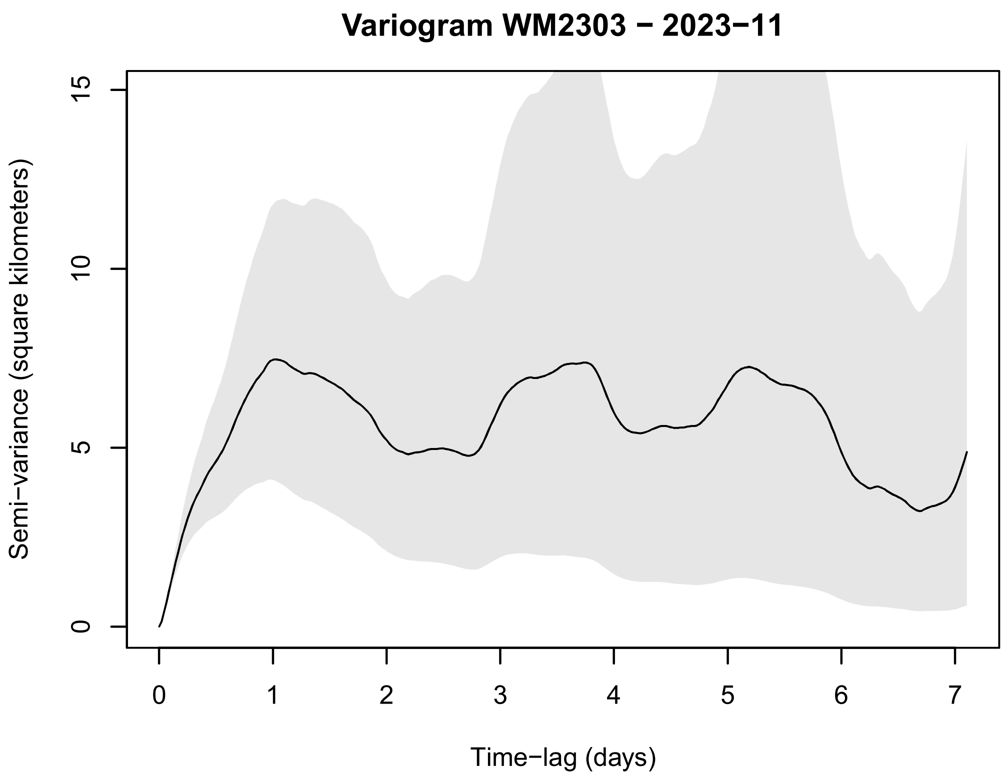
Variogram WM2303 − 2023−11
Semi−variance (square kilometers)
0
5
10
15
Time−lag (days)
0
1
2
3
4
5
6
7

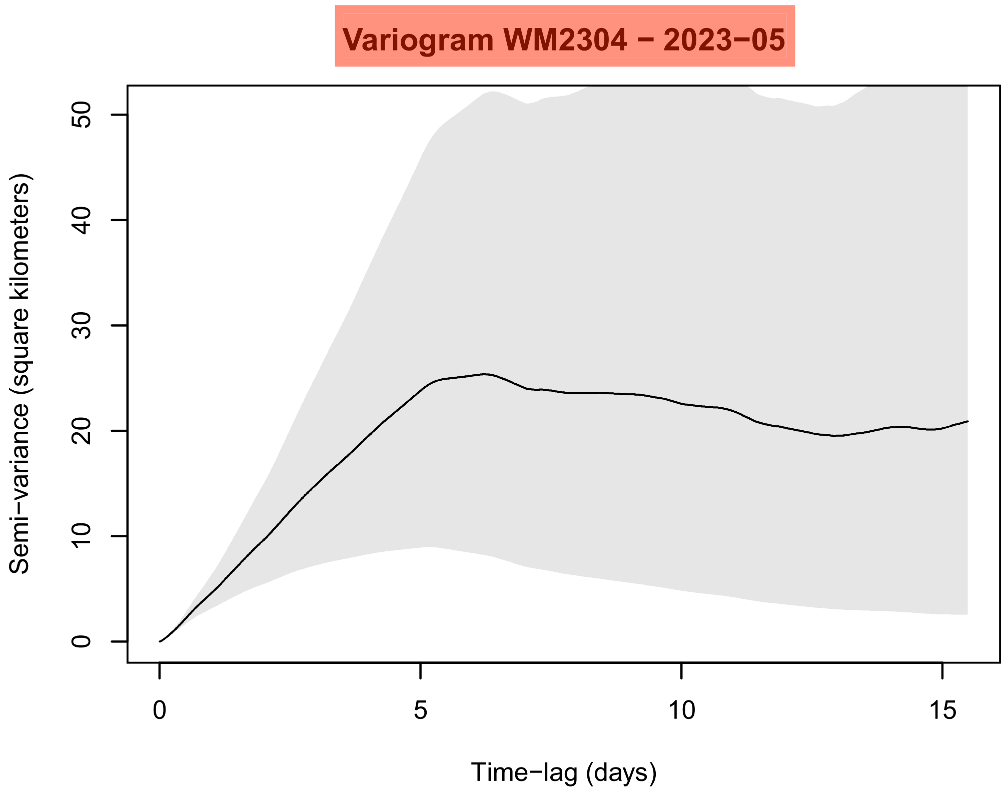
Variogram WM2304 − 2023−05
Semi−variance (square kilometers)
0
10
20
30
40
50
Time−lag (days)
0
5
10
15

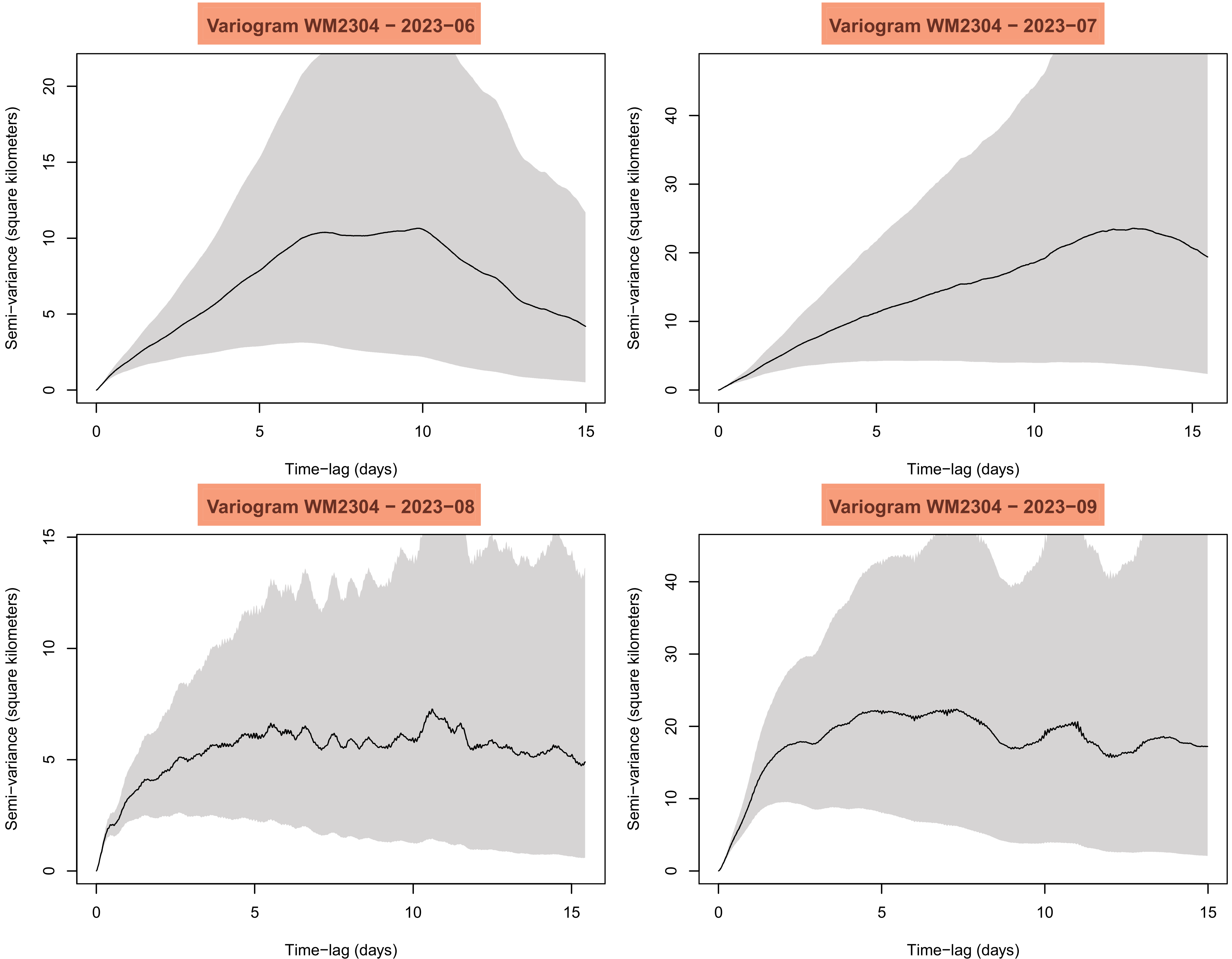

Variogram WM2304 − 2023−06
Variogram WM2304 − 2023−07
Variogram WM2304 − 2023−08
Variogram WM2304 − 2023−09
Semi−variance (square kilometers)
Time−lag (days)
0
5
10
15
20
30
40

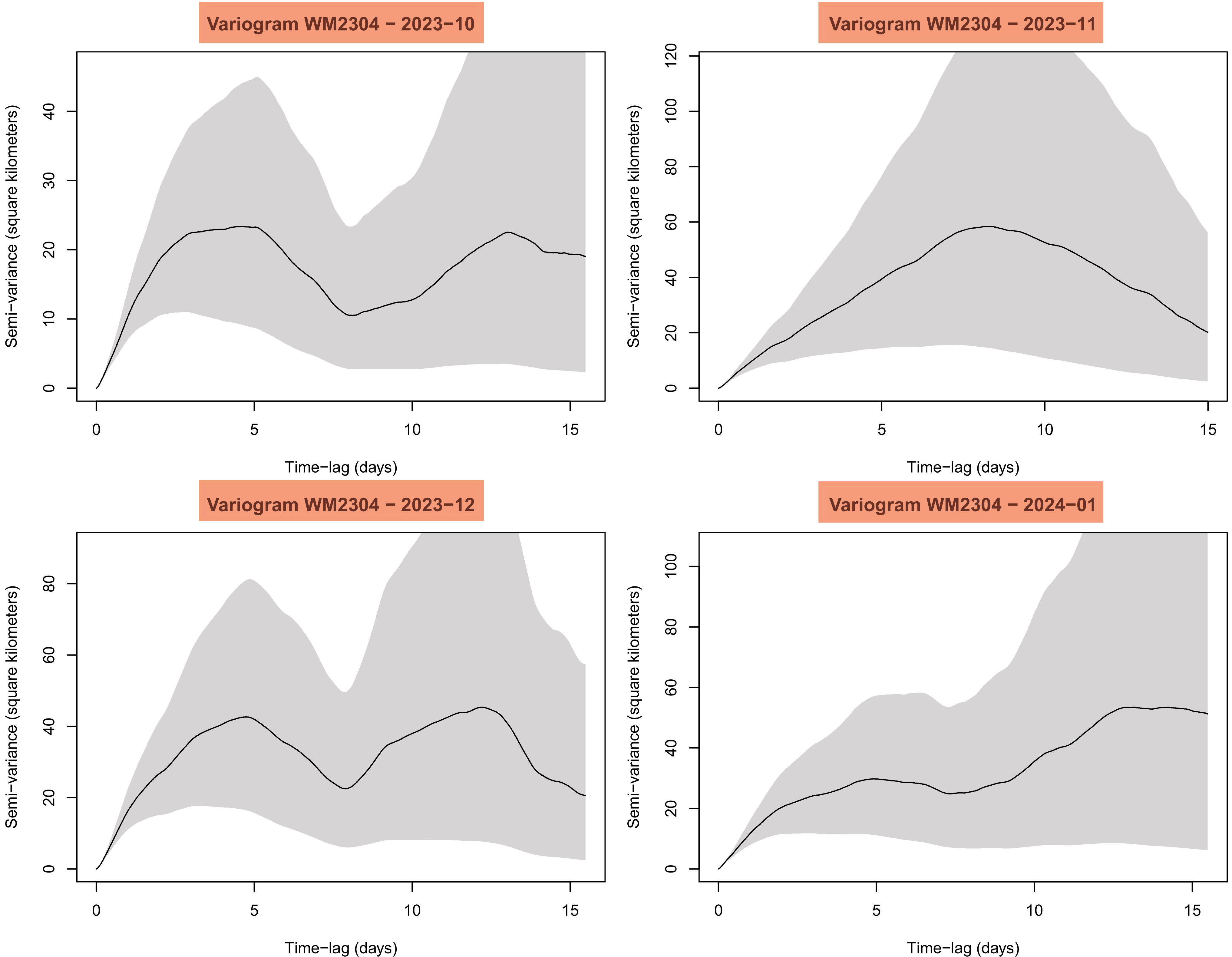
Variogram WM2304 − 2023−10
Semi−variance (square kilometers)
Time−lag (days)
Variogram WM2304 − 2023−11
Semi−variance (square kilometers)
Time−lag (days)
Variogram WM2304 − 2023−12
Semi−variance (square kilometers)
Time−lag (days)
Variogram WM2304 − 2024−01
Semi−variance (square kilometers)
Time−lag (days)

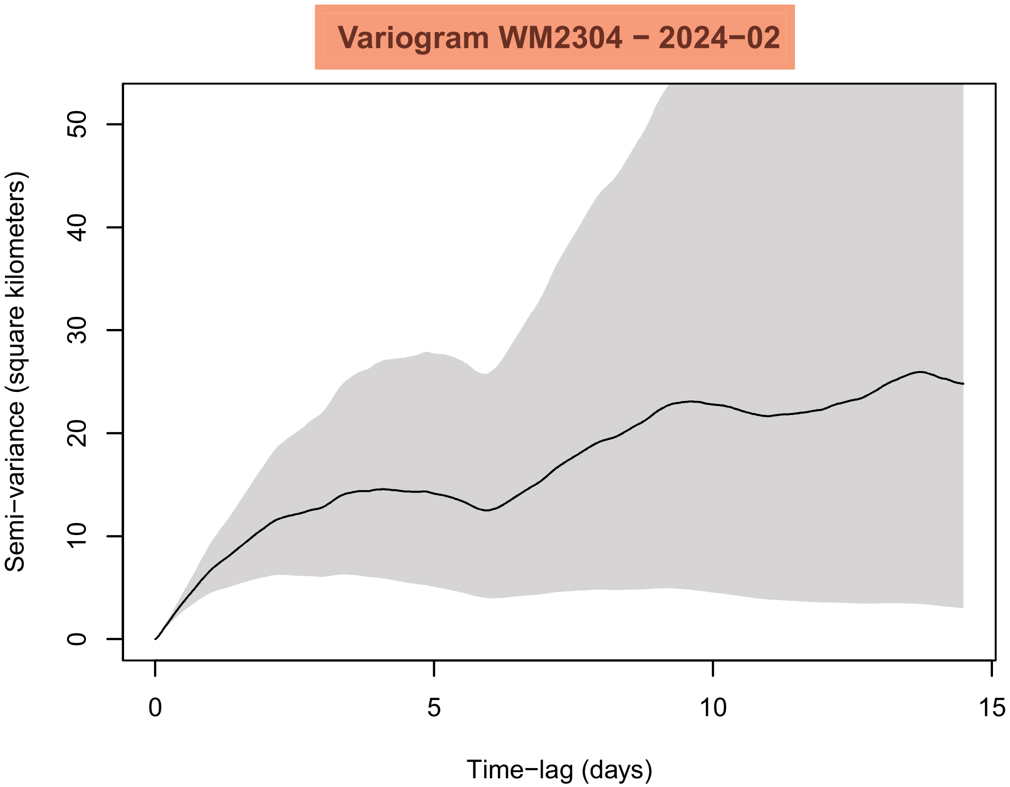

Variogram WM2304 − 2024−02
Semi−variance (square kilometers)
Time−lag (days)
0
10
20
30
40
50
0
5
10
15


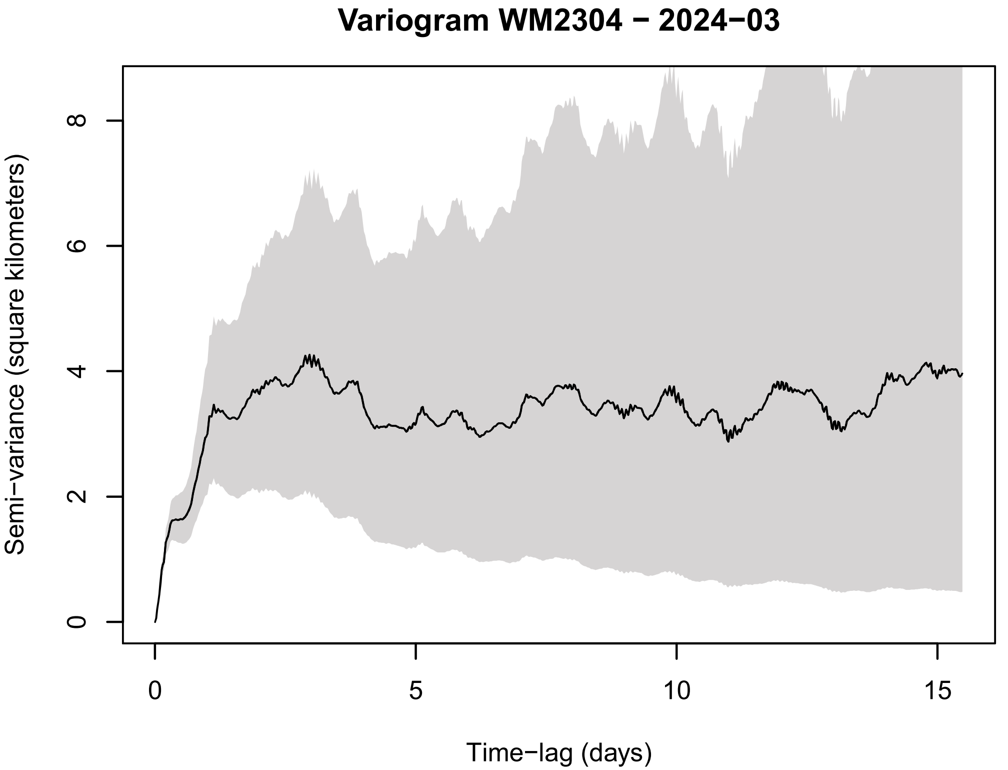

Variogram WM2304 − 2024−03
Semi−variance (square kilometers)
Time−lag (days)
0
2
4
6
8
0
5
10
15


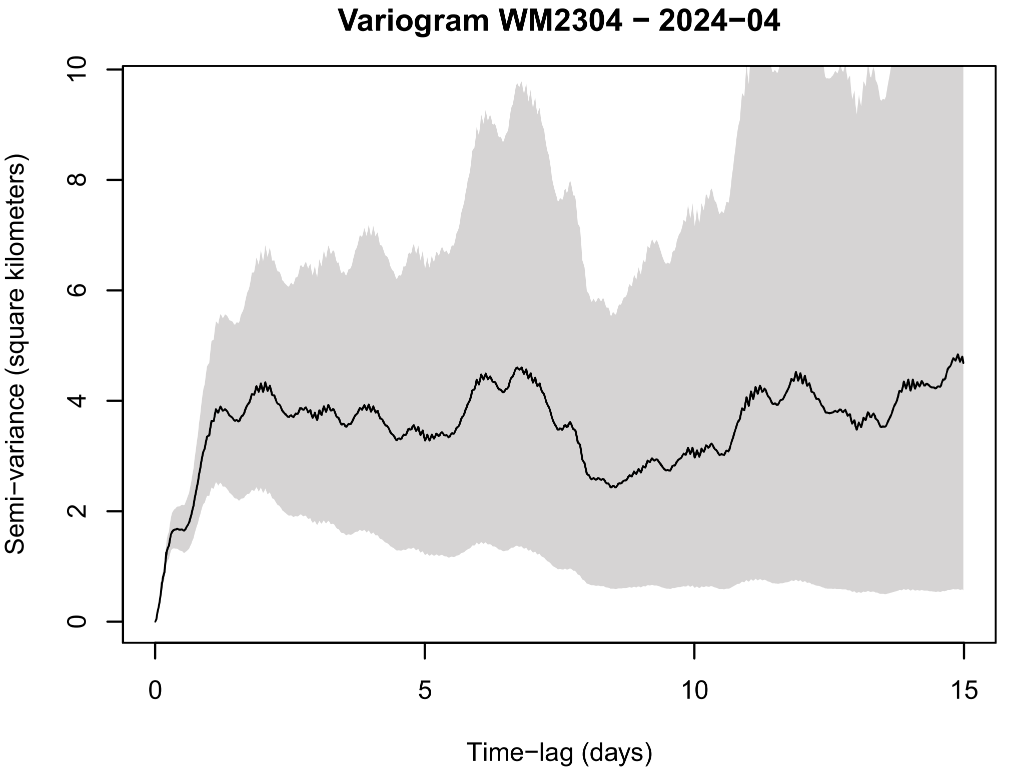

Variogram WM2304 − 2024−04
Semi−variance (square kilometers)
Time−lag (days)
0
2
4
6
8
10
0
5
10
15


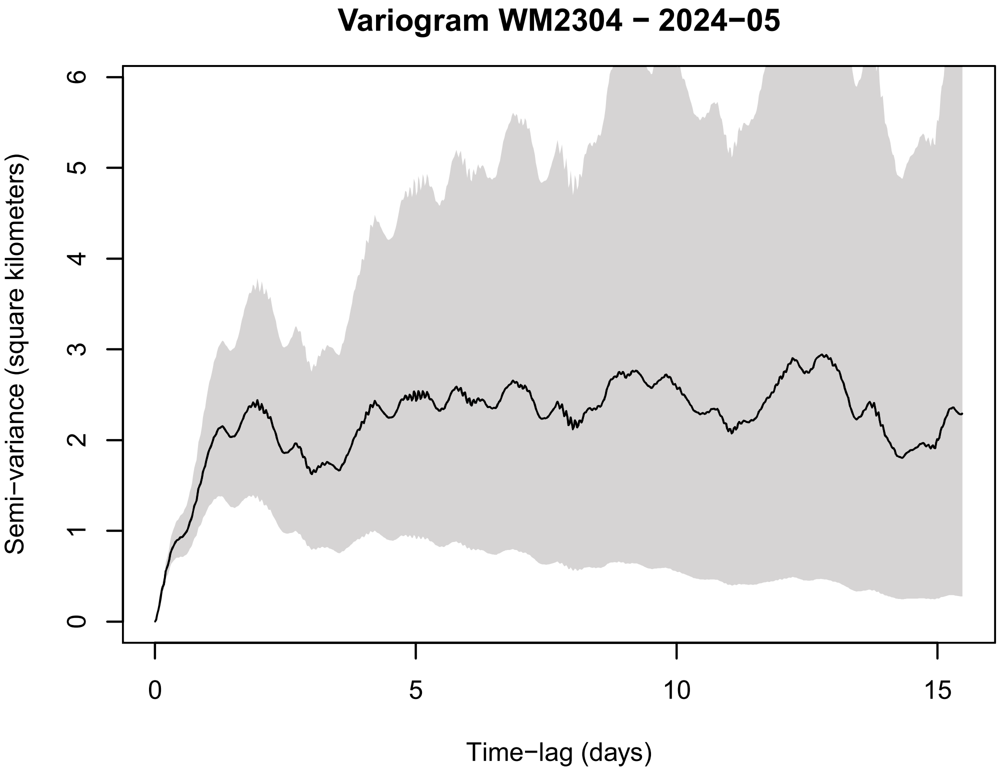

Variogram WM2304 − 2024−05
Semi−variance (square kilometers)
Time−lag (days)
0
1
2
3
4
5
6
0
5
10
15

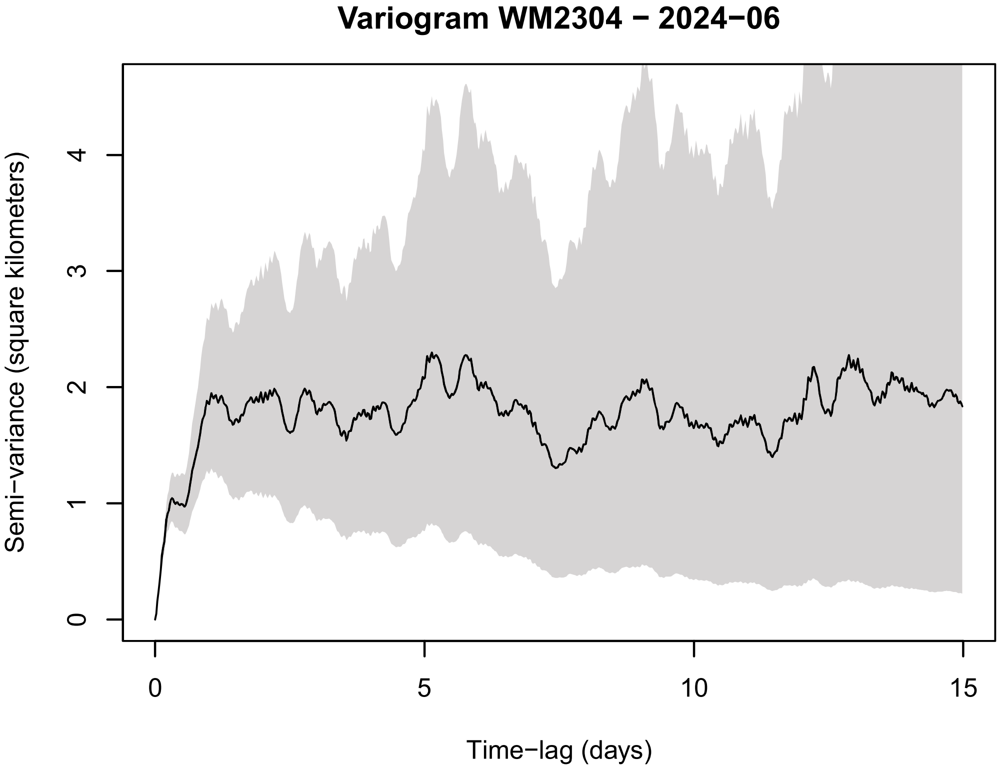

Variogram WM2304 − 2024−06
Semi−variance (square kilometers)
0
1
2
3
4
Time−lag (days)
0
5
10
15


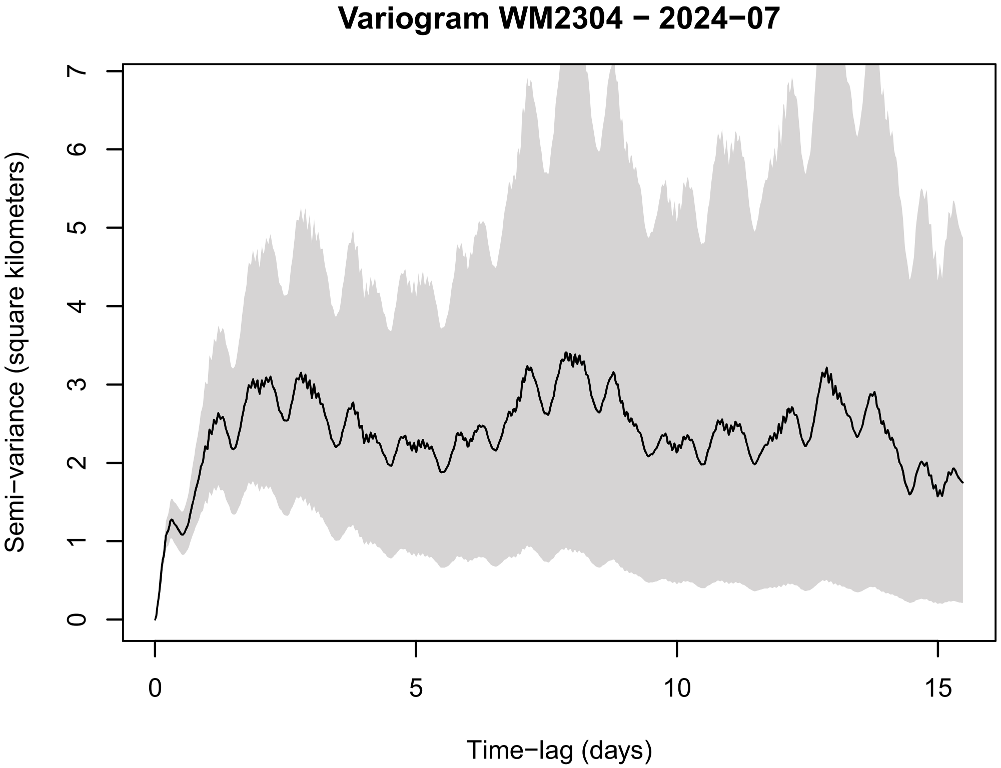

Variogram WM2304 − 2024−07
Semi−variance (square kilometers)
0
1
2
3
4
5
6
7
Time−lag (days)
0
5
10
15


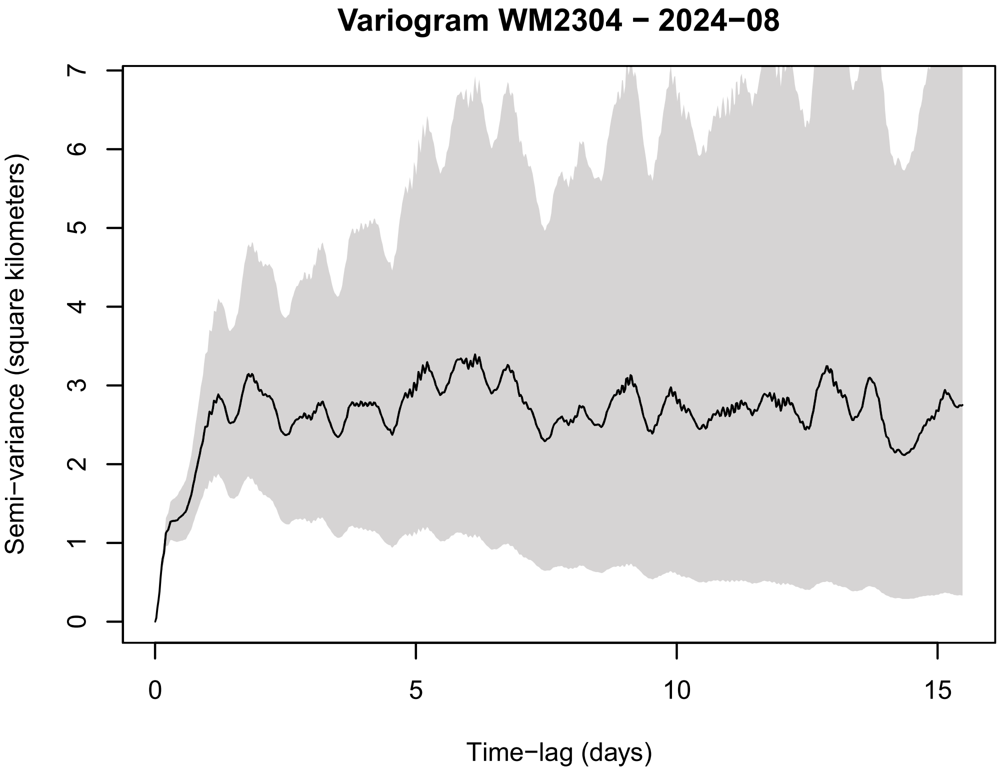

Variogram WM2304 − 2024−08
Semi−variance (square kilometers)
0
1
2
3
4
5
6
7
Time−lag (days)
0
5
10
15


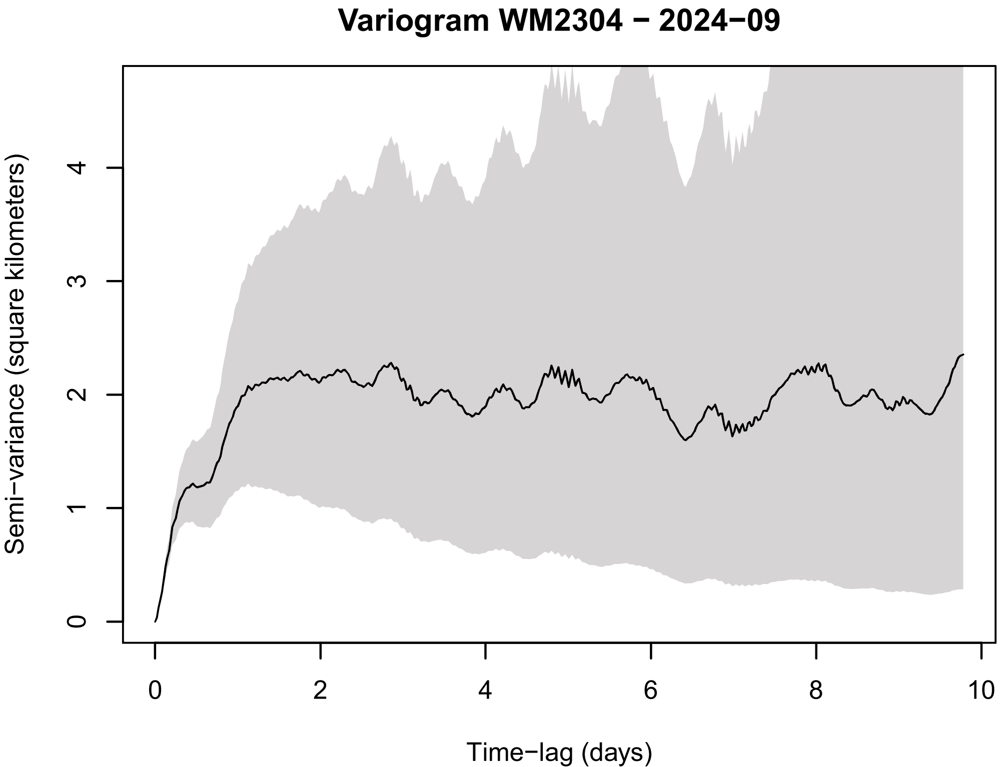

Variogram WM2304 − 2024−09
Semi−variance (square kilometers)
0
1
2
3
4
Time−lag (days)
0
2
4
6
8
10

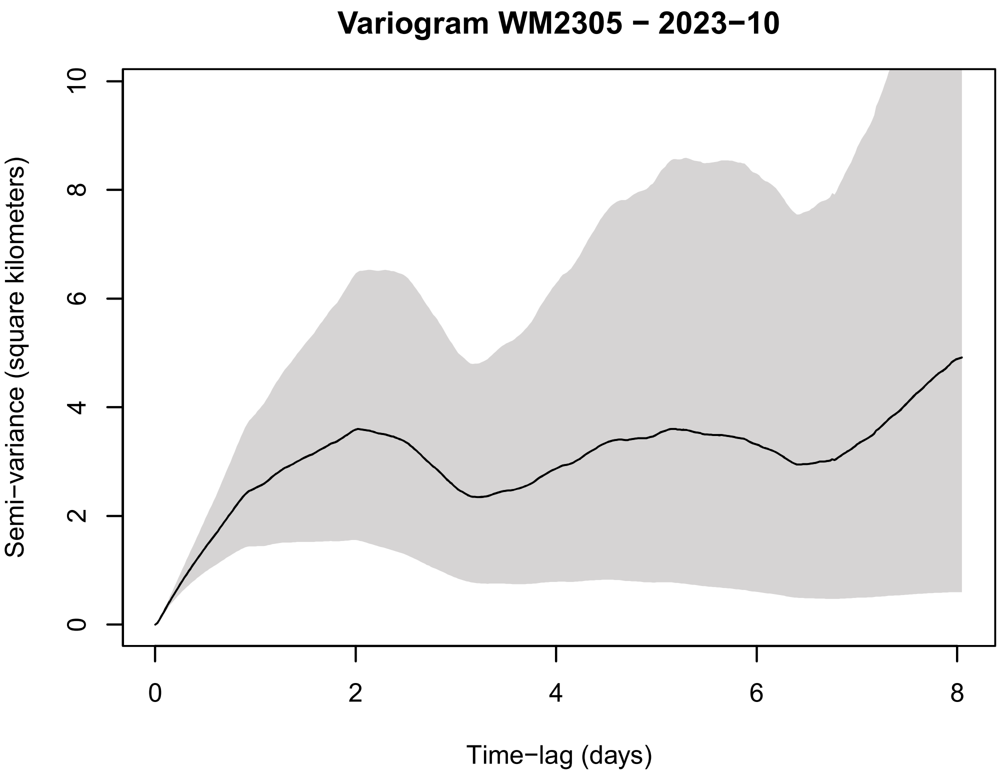
Variogram WM2305 − 2023−10
Semi−variance (square kilometers)
Time−lag (days)

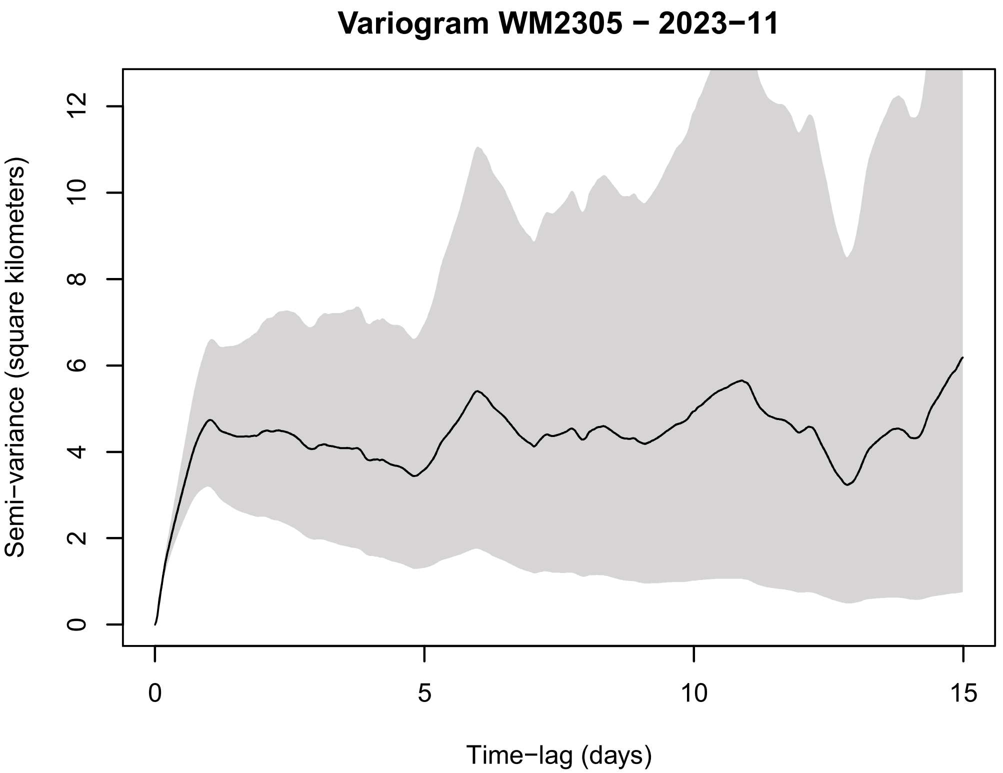
Variogram WM2305 − 2023−11
Semi−variance (square kilometers)
Time−lag (days)

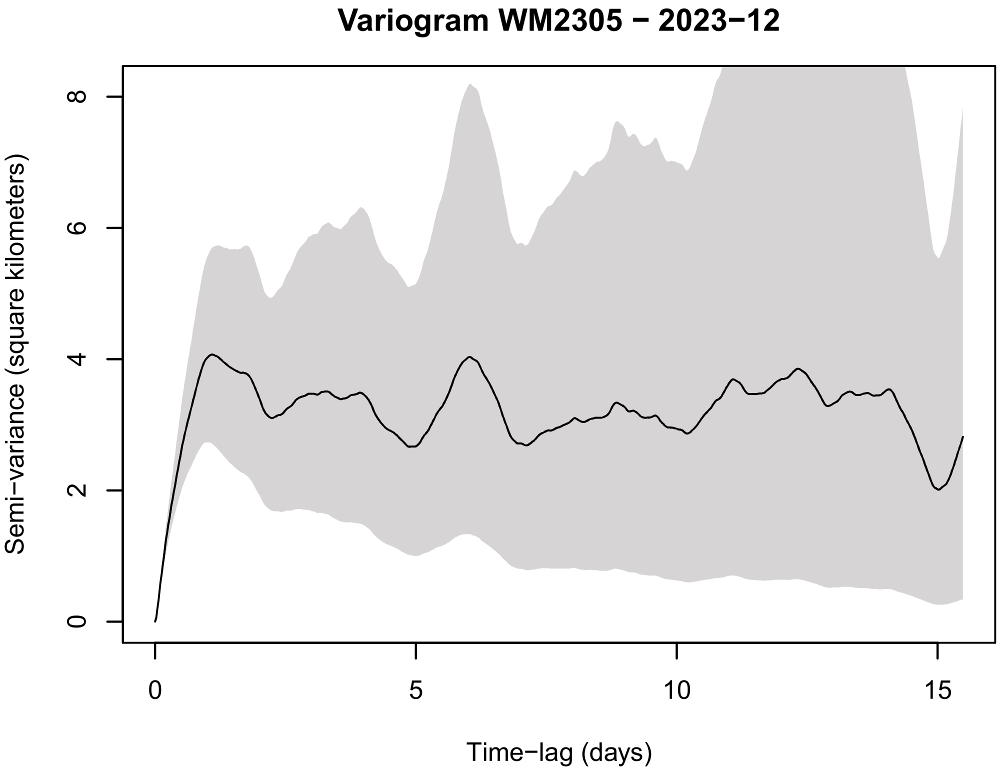
Variogram WM2305 − 2023−12
Semi−variance (square kilometers)
Time−lag (days)

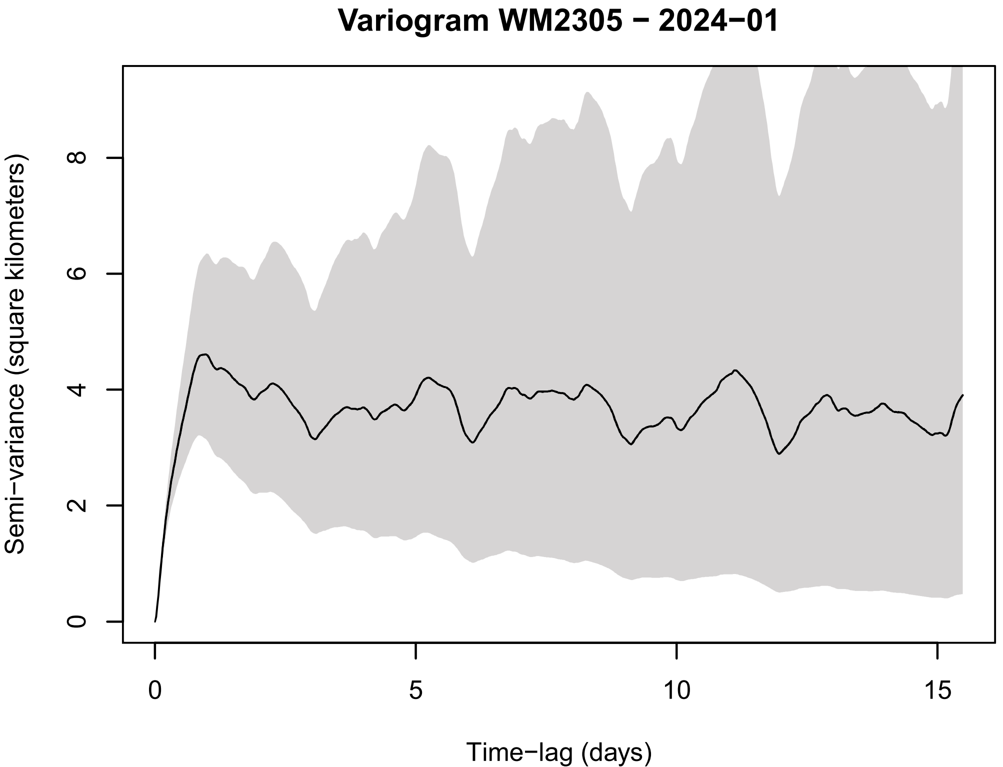
Variogram WM2305 − 2024−01
Semi−variance (square kilometers)
Time−lag (days)

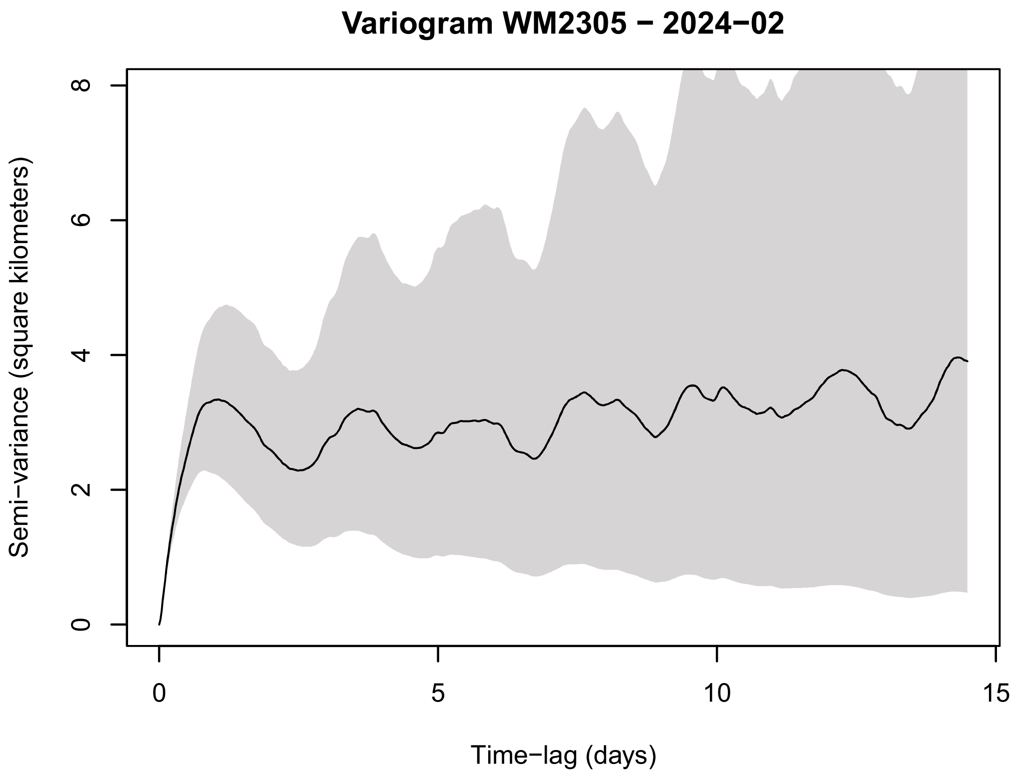
Variogram WM2305 − 2024−02
Semi−variance (square kilometers)
0
2
4
6
8
Time−lag (days)
0
5
10
15

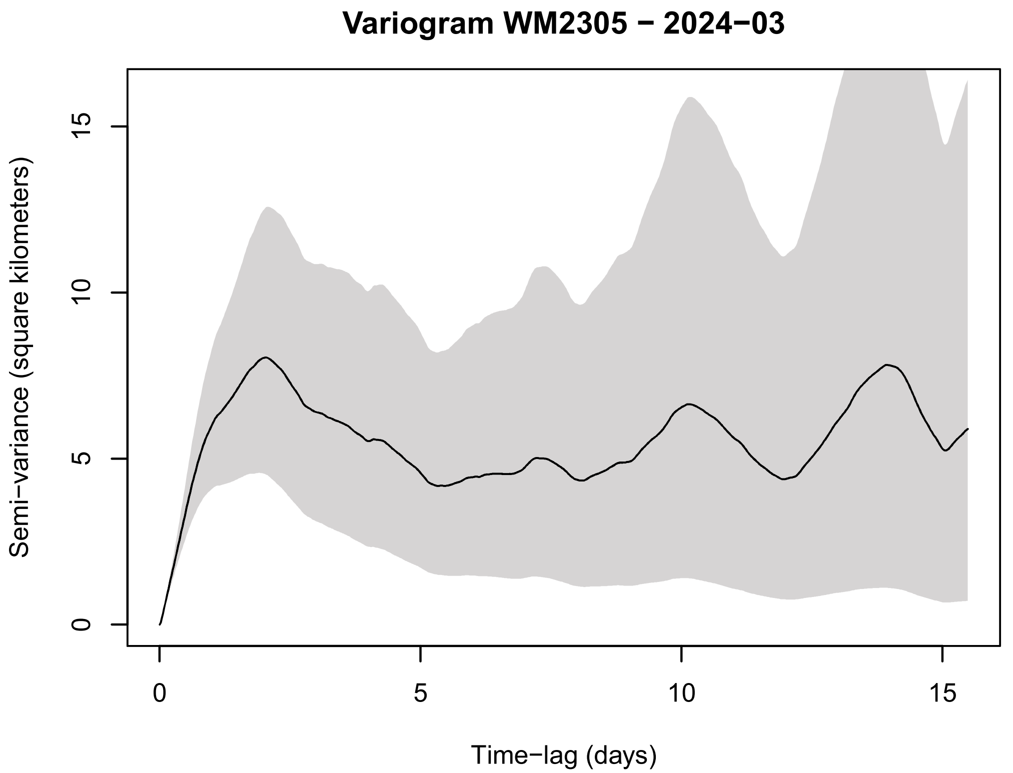
Variogram WM2305 − 2024−03
Semi−variance (square kilometers)
0
5
10
15
Time−lag (days)
0
5
10
15

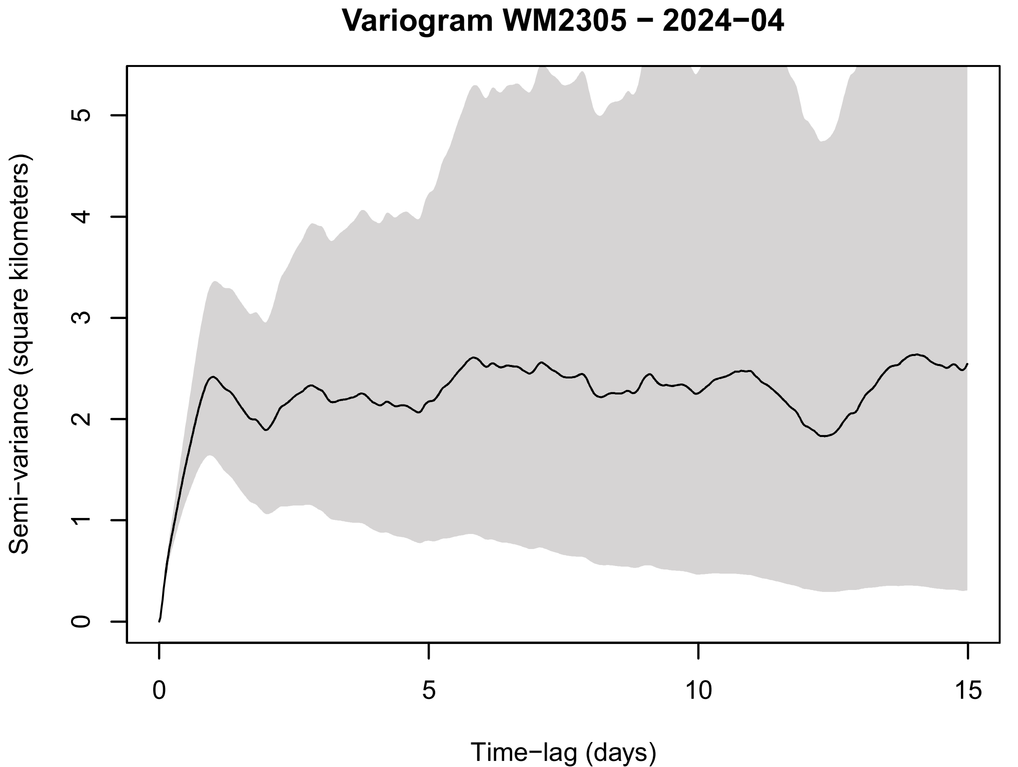
Variogram WM2305 − 2024−04
Semi−variance (square kilometers)
0
1
2
3
4
5
Time−lag (days)
0
5
10
15

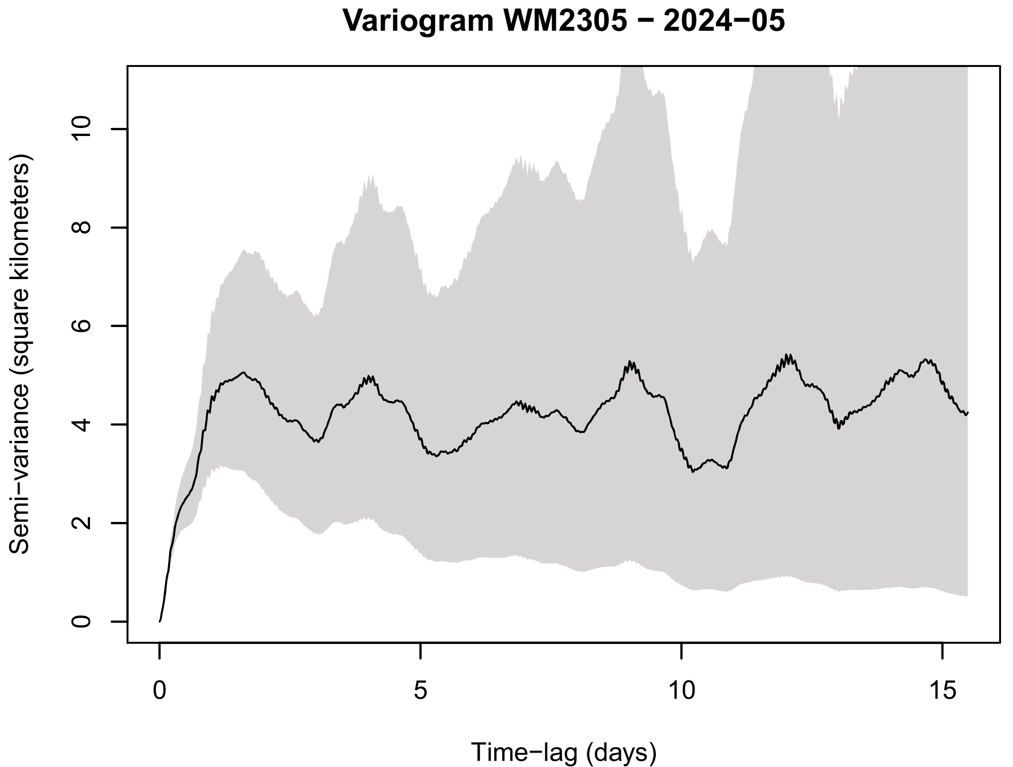
Variogram WM2305 − 2024−05
Semi−variance (square kilometers)
0
2
4
6
8
10
Time−lag (days)
0
5
10
15

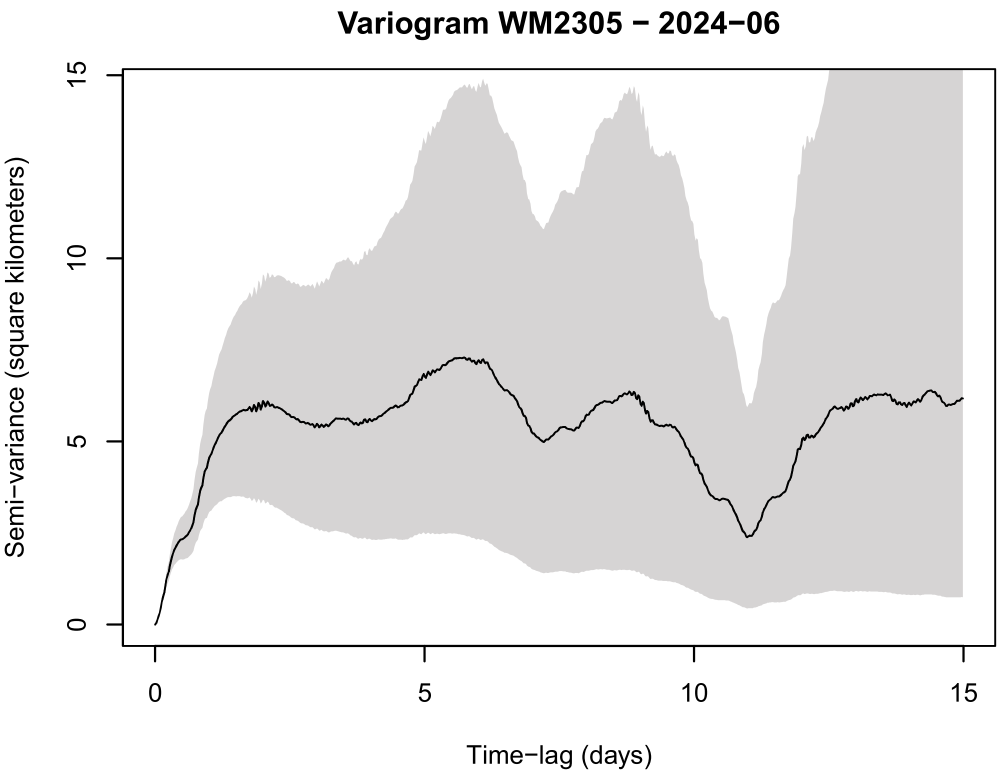
Variogram WM2305 − 2024−06
Semi−variance (square kilometers)
0
5
10
15
Time−lag (days)
0
5
10
15

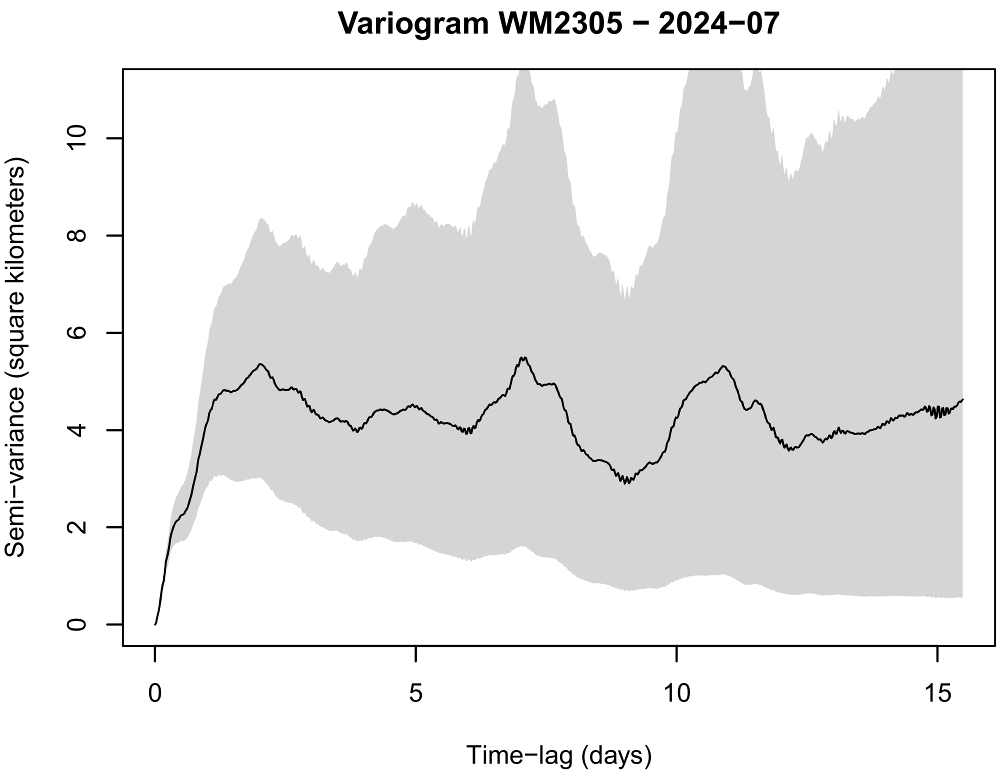
Variogram WM2305 − 2024−07
Semi−variance (square kilometers)
0
2
4
6
8
10
Time−lag (days)
0
5
10
15

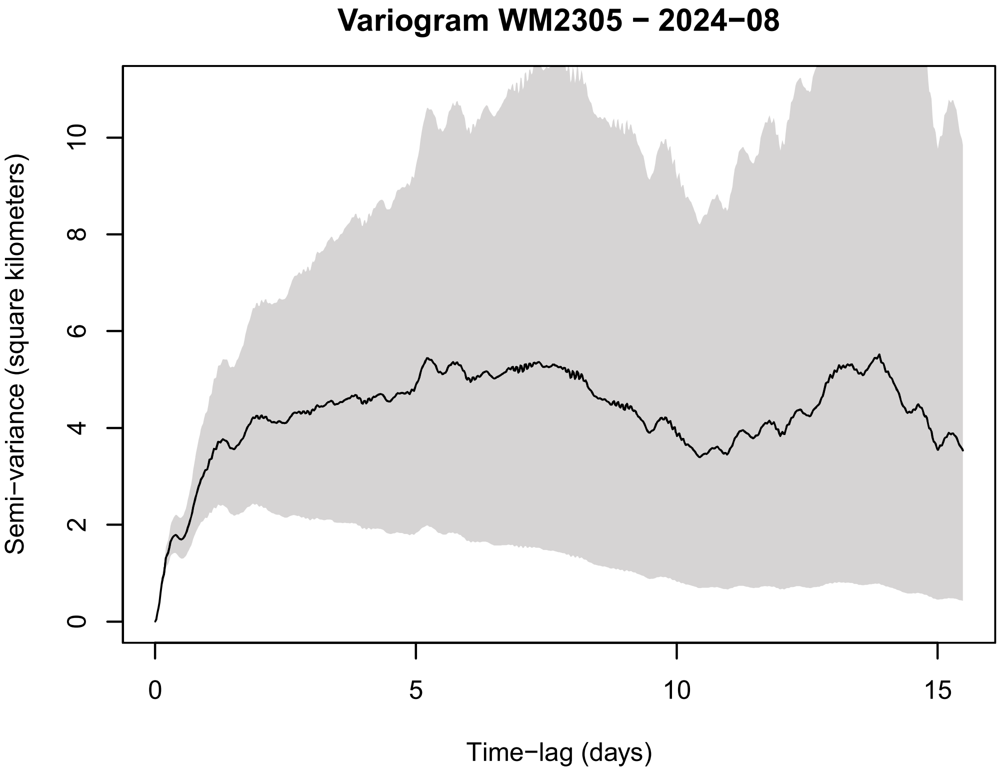
Variogram WM2305 − 2024−08
Semi−variance (square kilometers)
0
2
4
6
8
10
Time−lag (days)
0
5
10
15

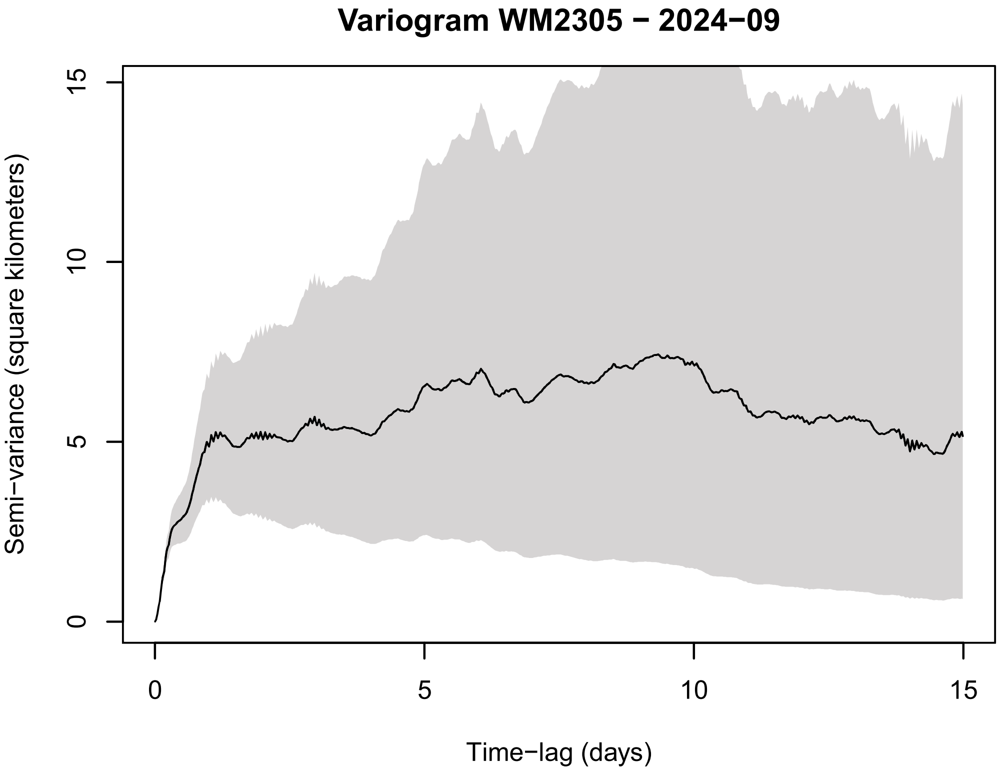
Variogram WM2305 − 2024−09
Semi−variance (square kilometers)
0
5
10
15
Time−lag (days)
0
5
10
15

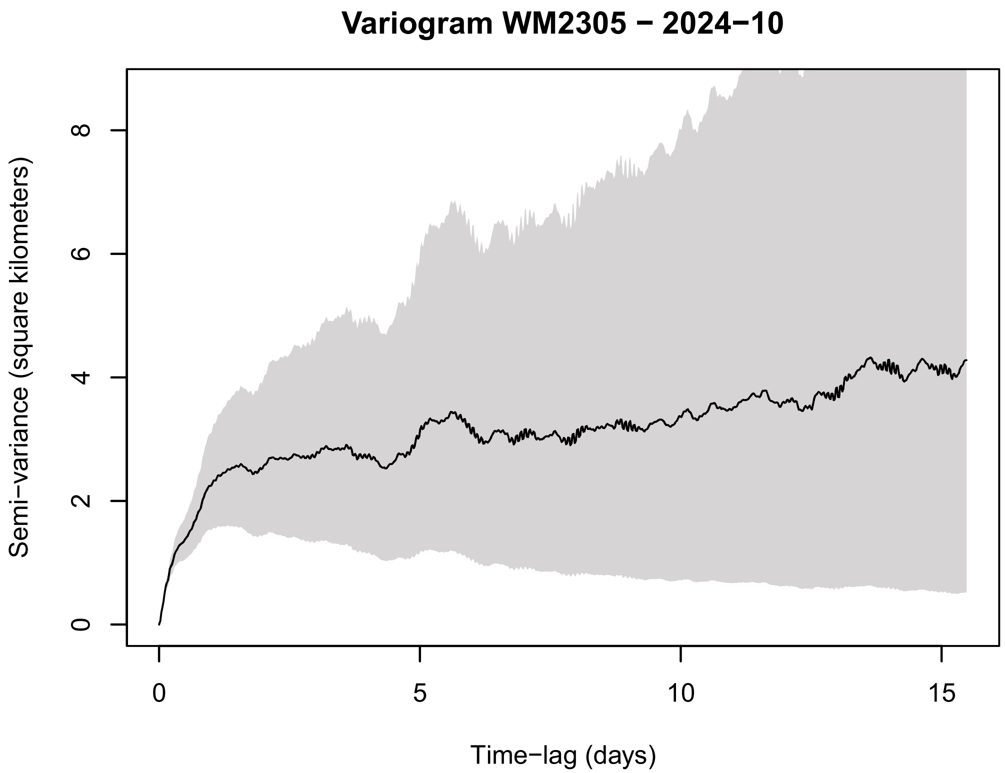
Variogram WM2305 − 2024−10
Semi−variance (square kilometers)
0
2
4
6
8
Time−lag (days)
0
5
10
15

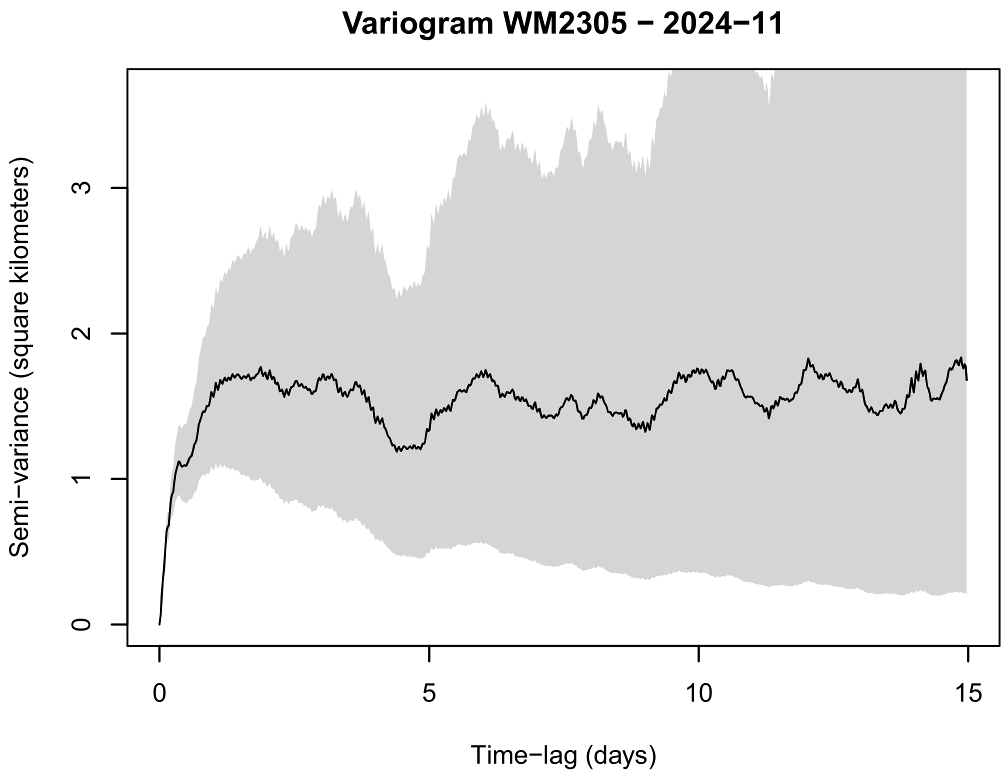
Variogram WM2305 − 2024−11
Semi−variance (square kilometers)
0
1
2
3
Time−lag (days)
0
5
10
15

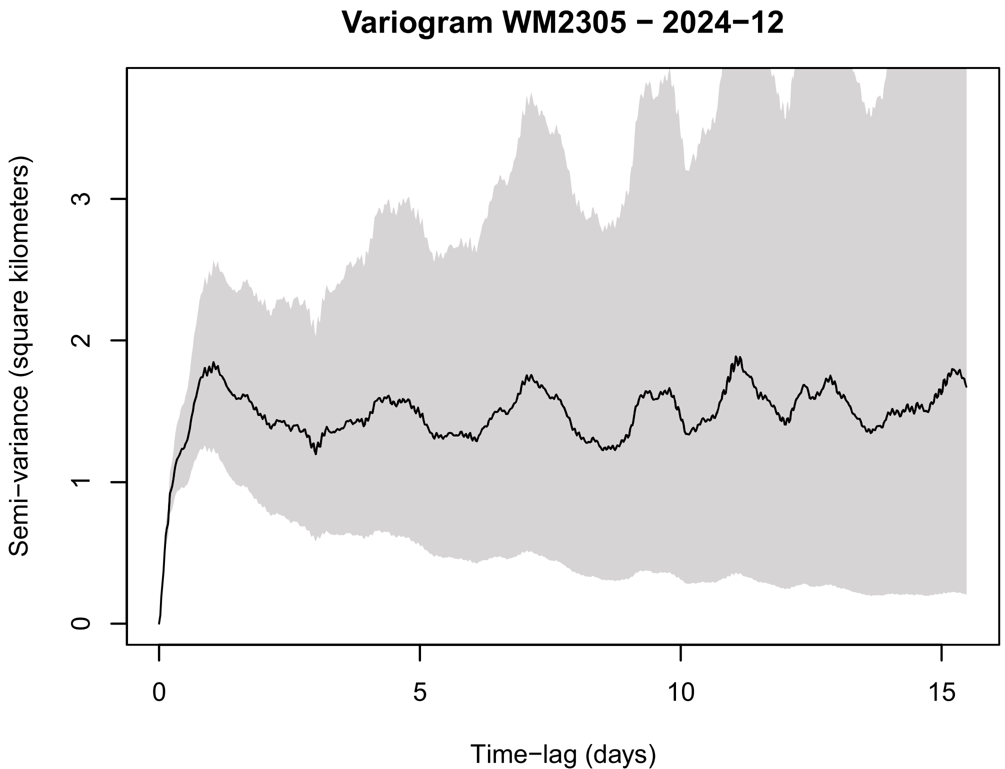
Variogram WM2305 − 2024−12
Semi−variance (square kilometers)
0
1
2
3
Time−lag (days)
0
5
10
15

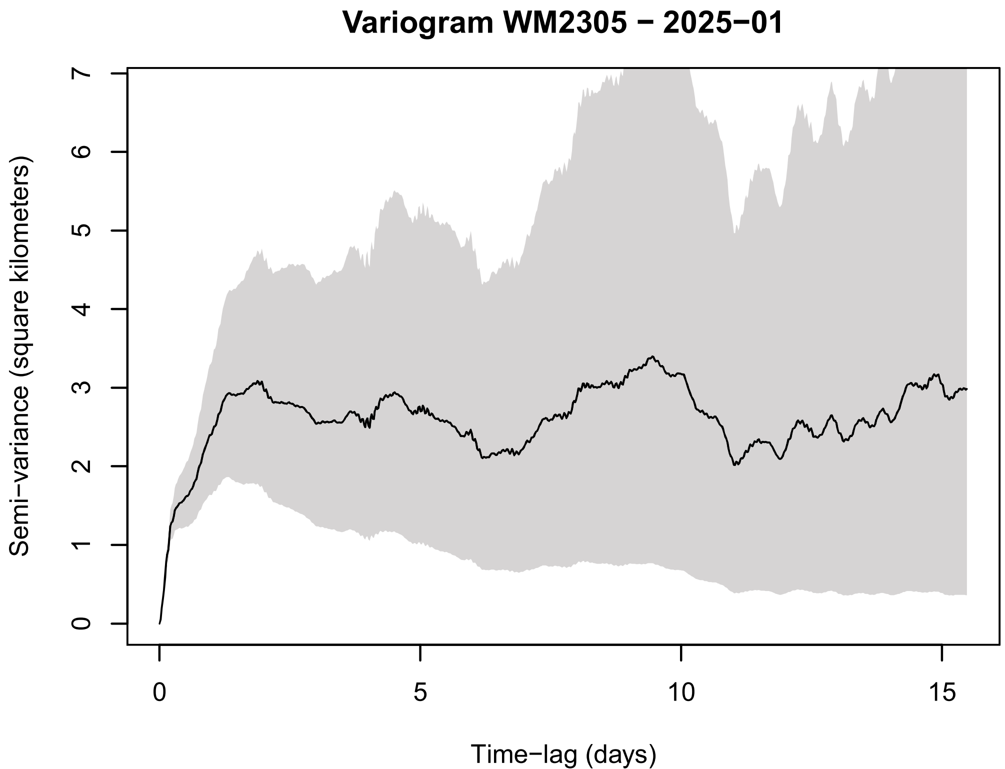
Variogram WM2305 − 2025−01
Semi−variance (square kilometers)
0
1
2
3
4
5
6
7
Time−lag (days)
0
5
10
15

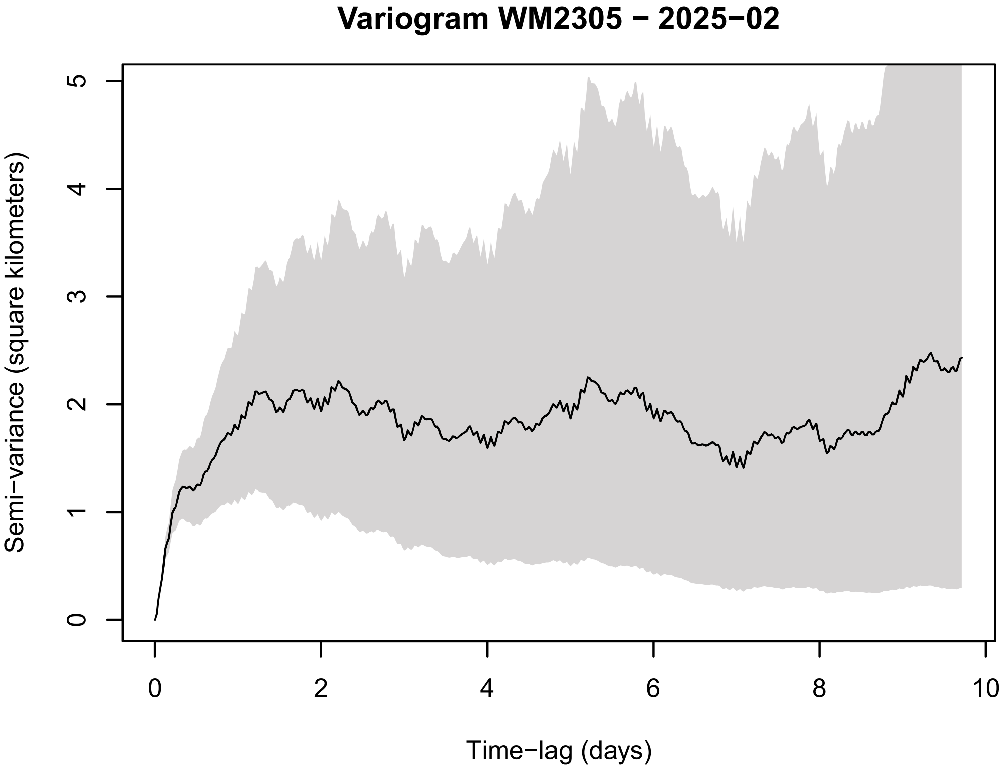
Variogram WM2305 − 2025−02
Semi−variance (square kilometers)
Time−lag (days)
0
1
2
3
4
5
0
2
4
6
8
10

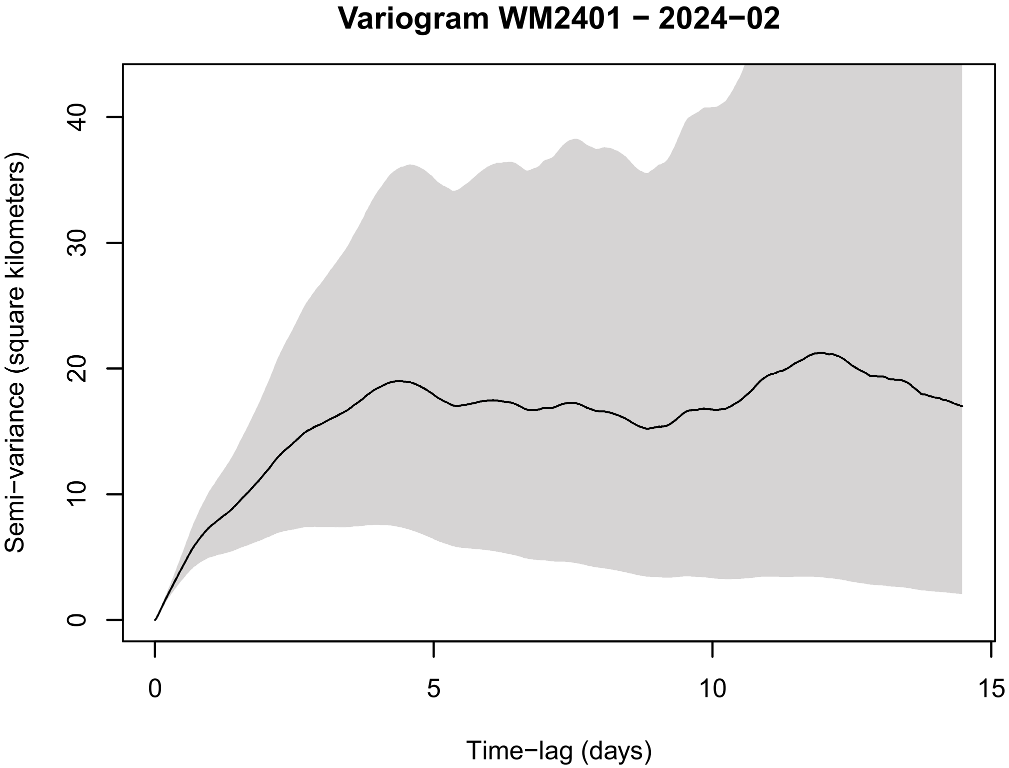
Variogram WM2401 − 2024−02
Semi−variance (square kilometers)
Time−lag (days)
0
10
20
30
40
0
5
10
15

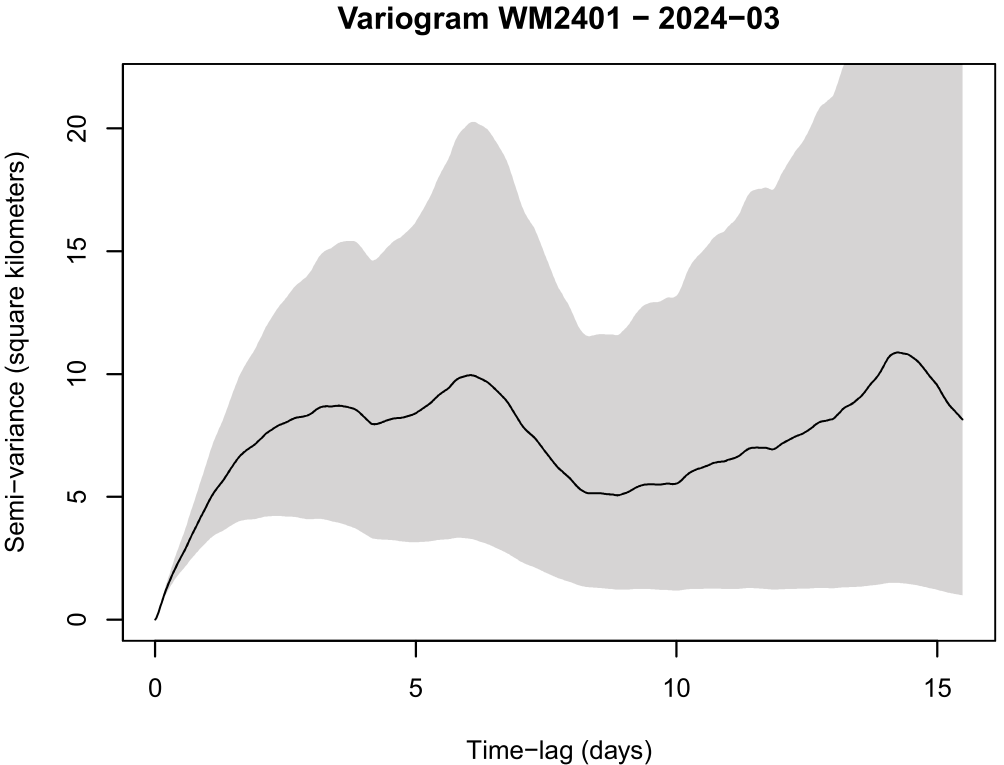
Variogram WM2401 − 2024−03
Semi−variance (square kilometers)
Time−lag (days)
0
5
10
15
20
0
5
10
15

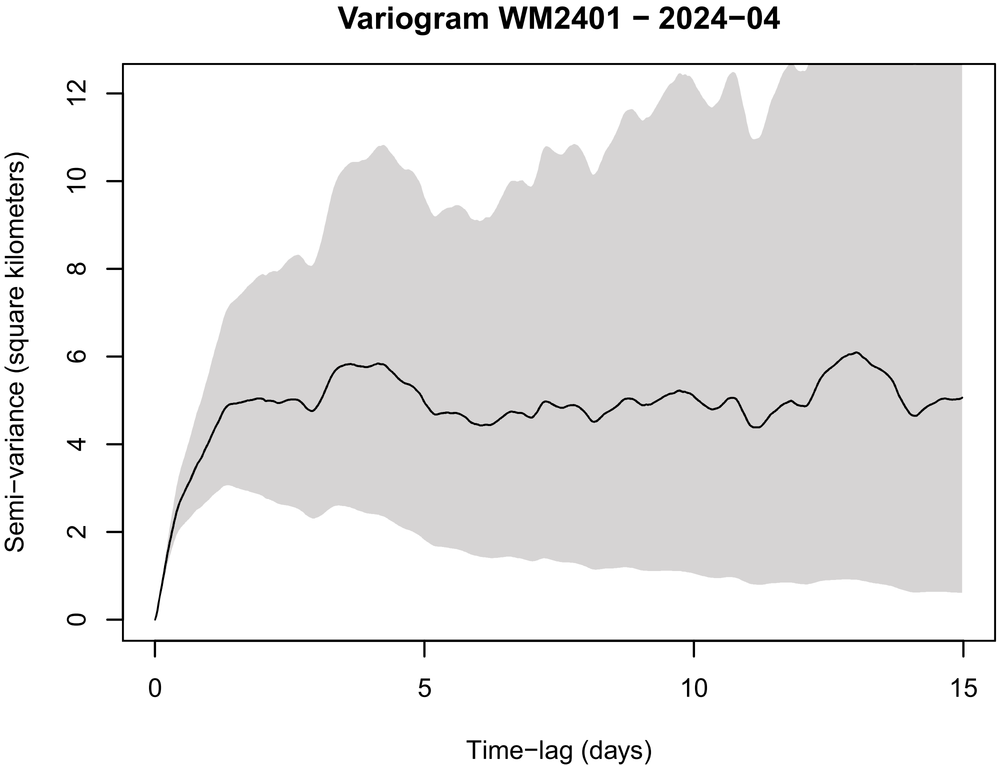
Variogram WM2401 − 2024−04
Semi−variance (square kilometers)
Time−lag (days)
0
2
4
6
8
10
12
0
5
10
15

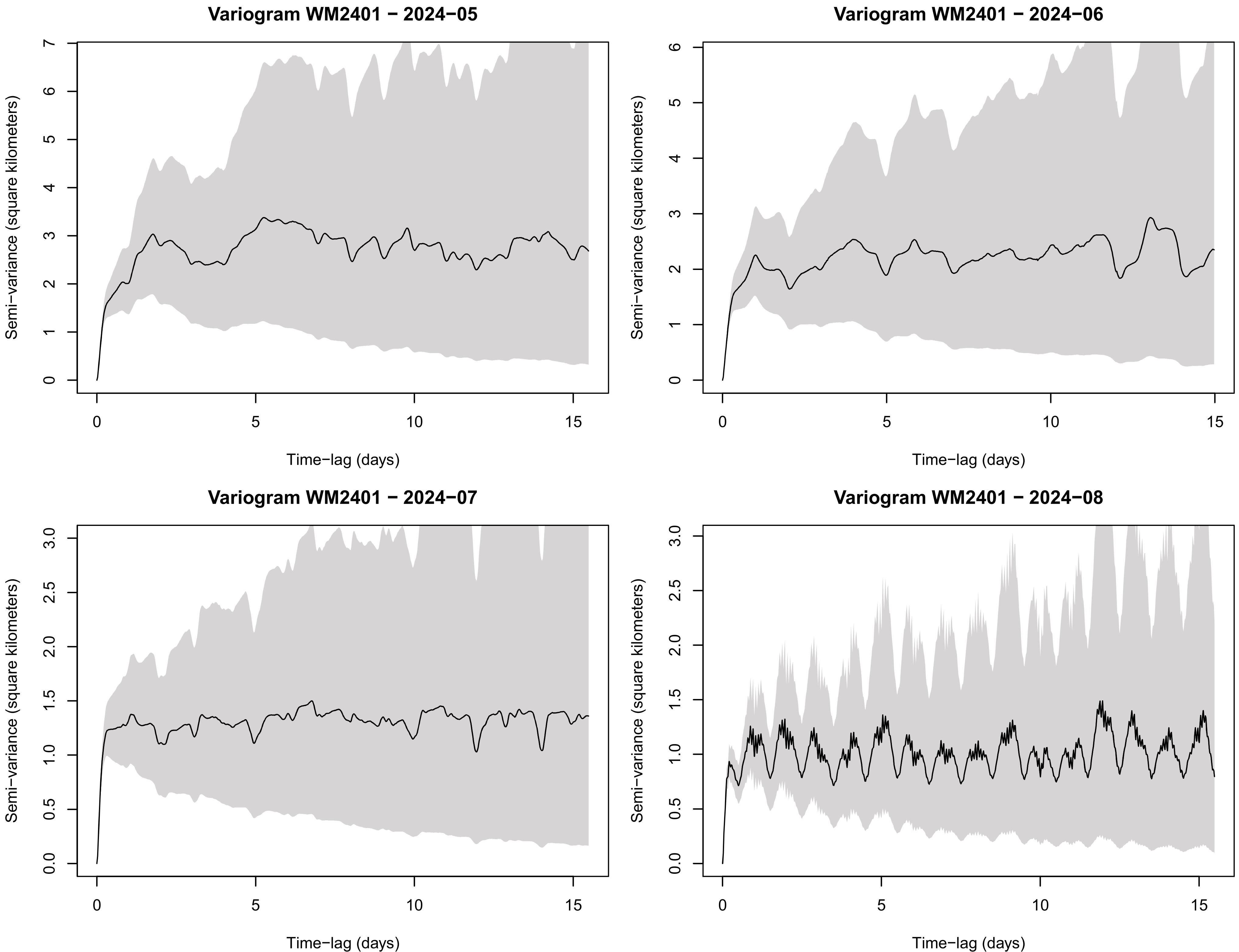

Variogram WM2401 − 2024−05
Variogram WM2401 − 2024−06
Variogram WM2401 − 2024−07
Variogram WM2401 − 2024−08
Semi−variance (square kilometers)
Time−lag (days)

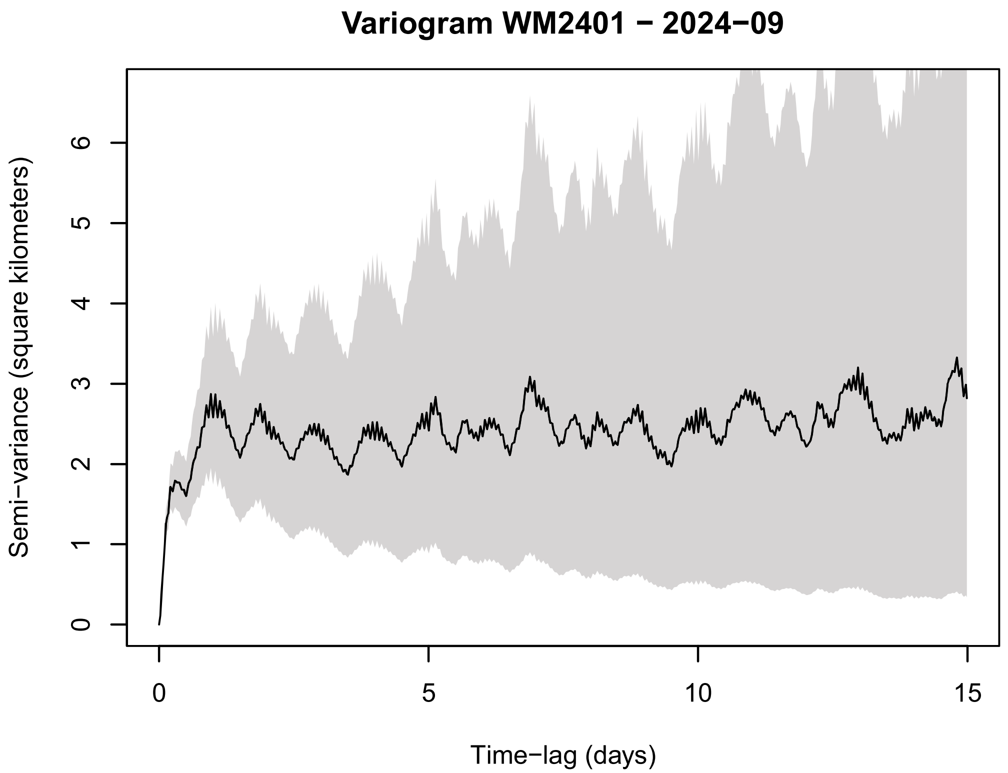
Variogram WM2401 − 2024−09
Semi−variance (square kilometers)
Time−lag (days)

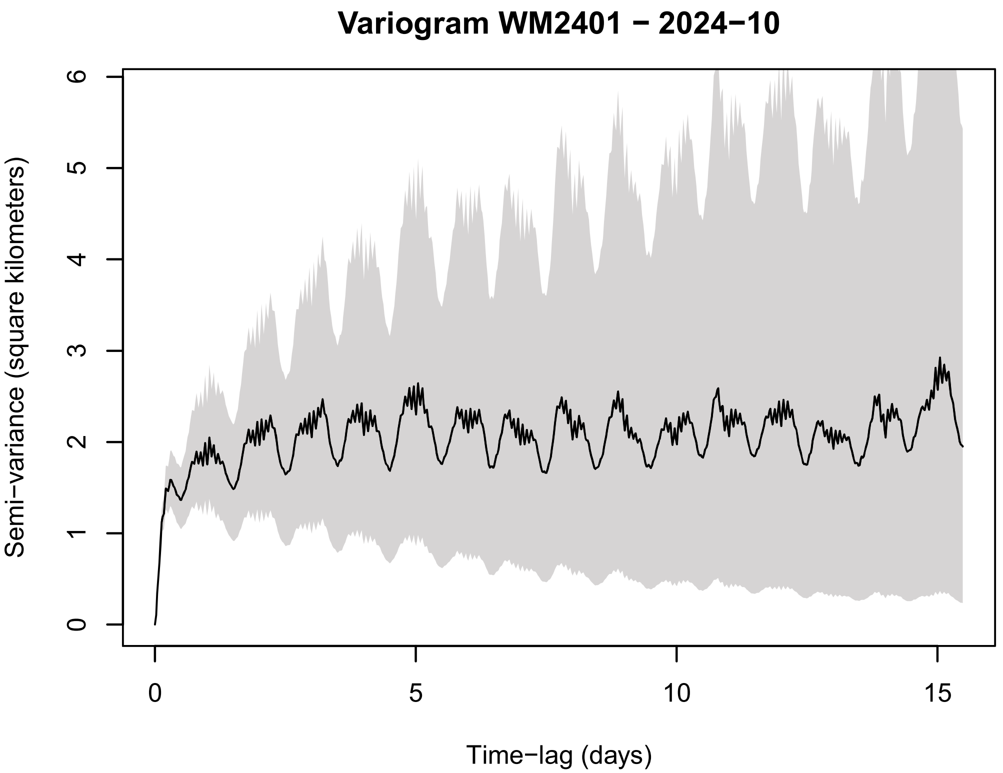
Variogram WM2401 − 2024−10
Semi−variance (square kilometers)
Time−lag (days)

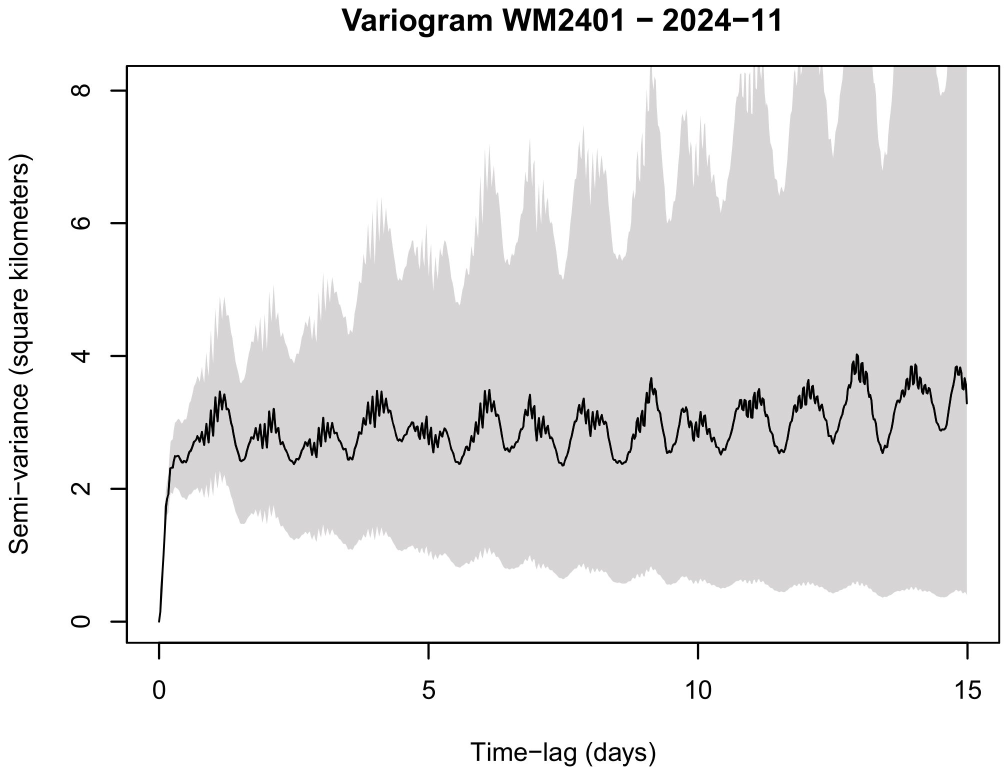
Variogram WM2401 − 2024−11
Semi−variance (square kilometers)
Time−lag (days)

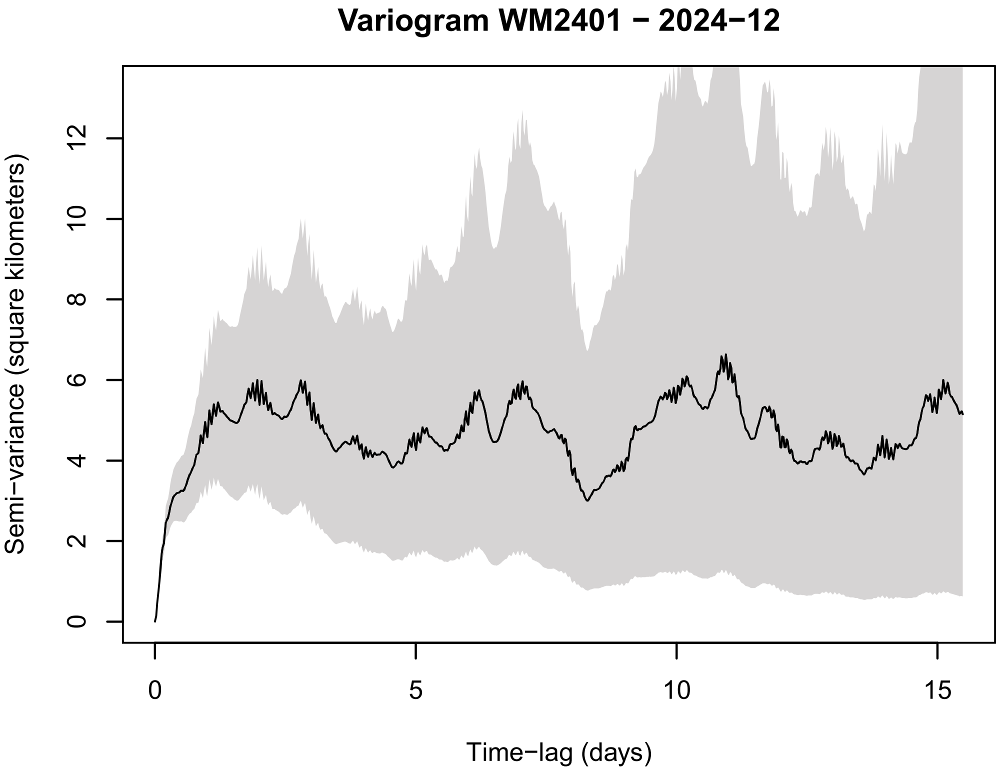
Variogram WM2401 − 2024−12
Semi−variance (square kilometers)
Time−lag (days)

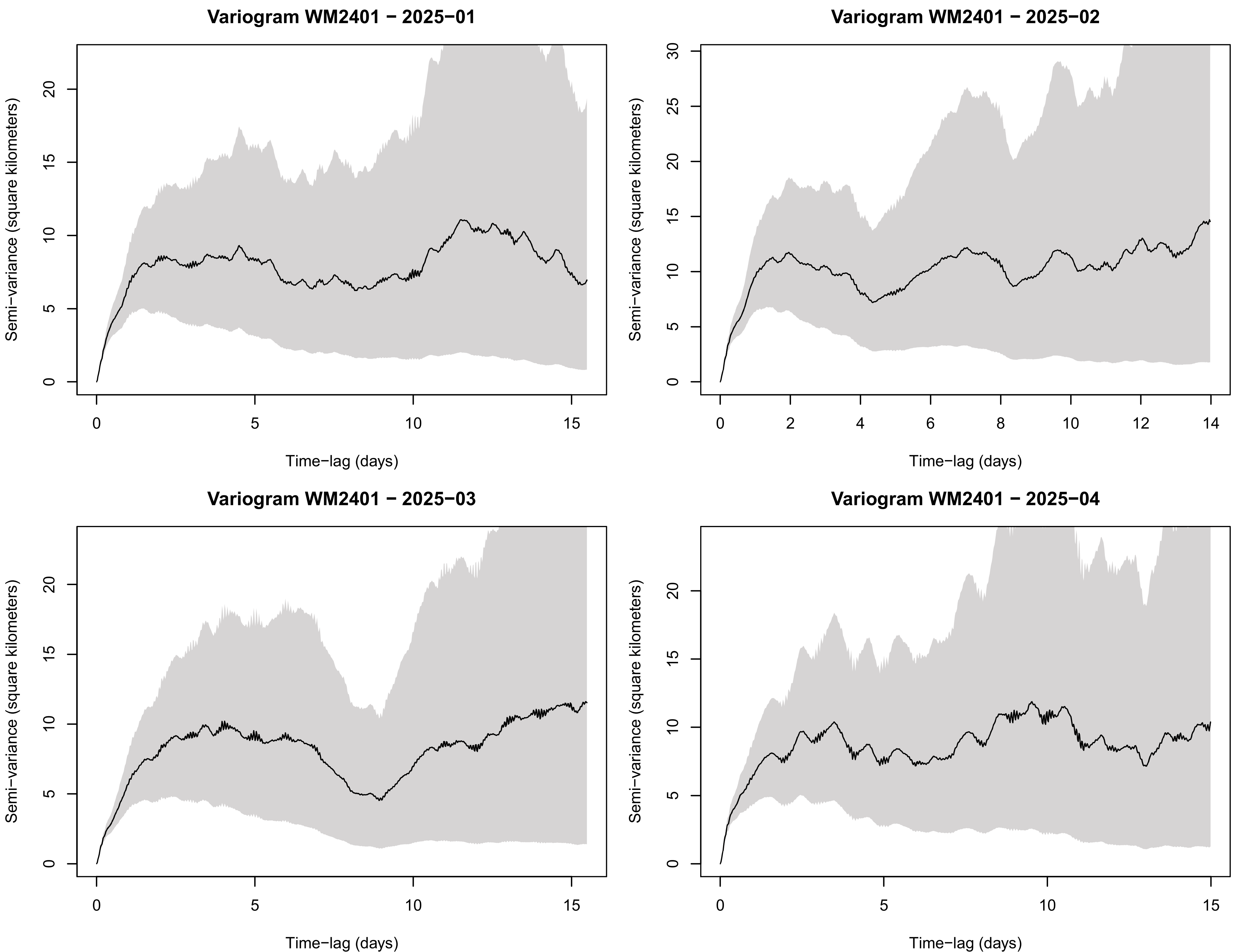
Variogram WM2401 − 2025−01
Variogram WM2401 − 2025−02
Variogram WM2401 − 2025−03
Variogram WM2401 − 2025−04
Semi−variance (square kilometers)
Time−lag (days)

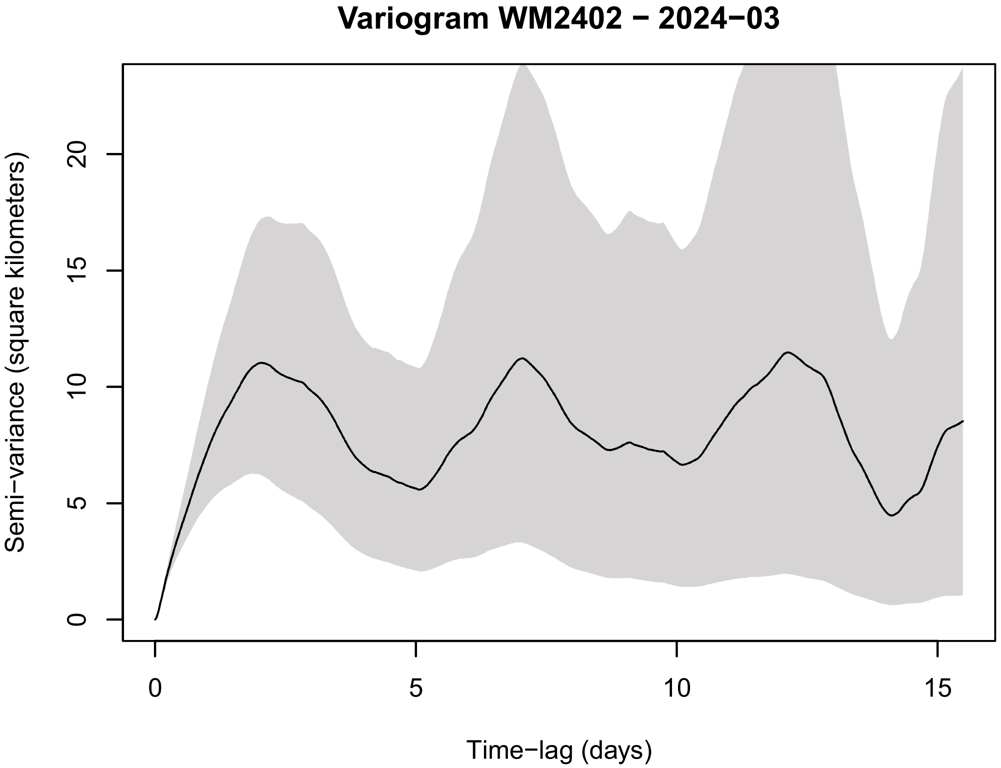
Variogram WM2402 − 2024−03
Semi−variance (square kilometers)
Time−lag (days)
0
5
10
15
20

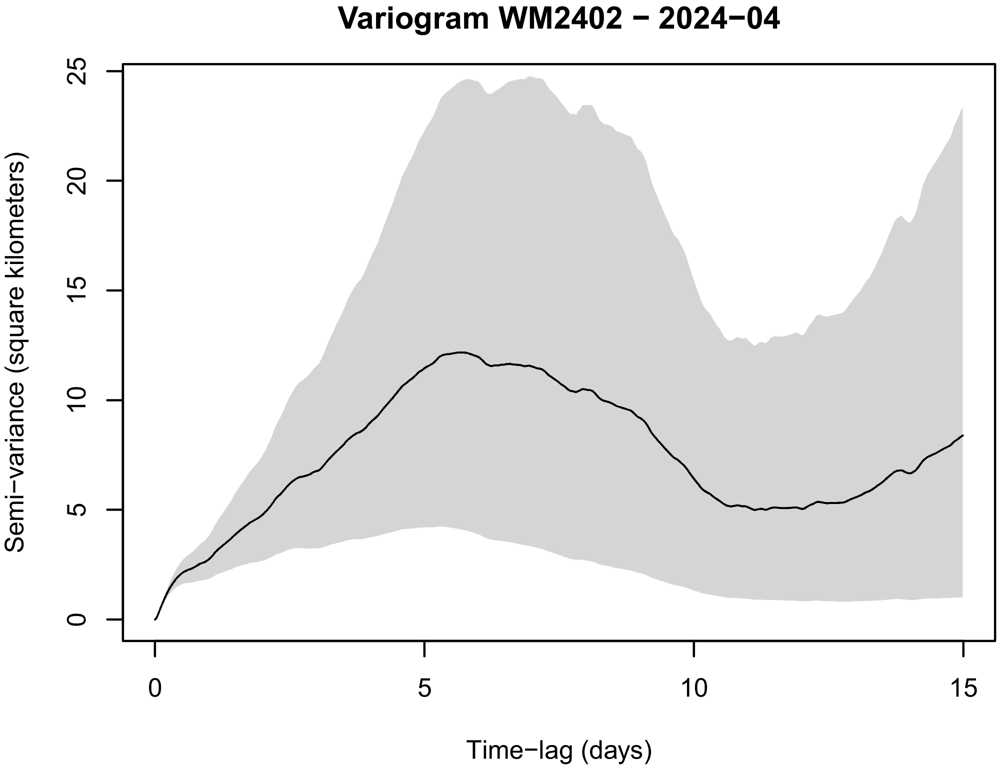
Variogram WM2402 − 2024−04
Semi−variance (square kilometers)
Time−lag (days)
0
5
10
15
20
25

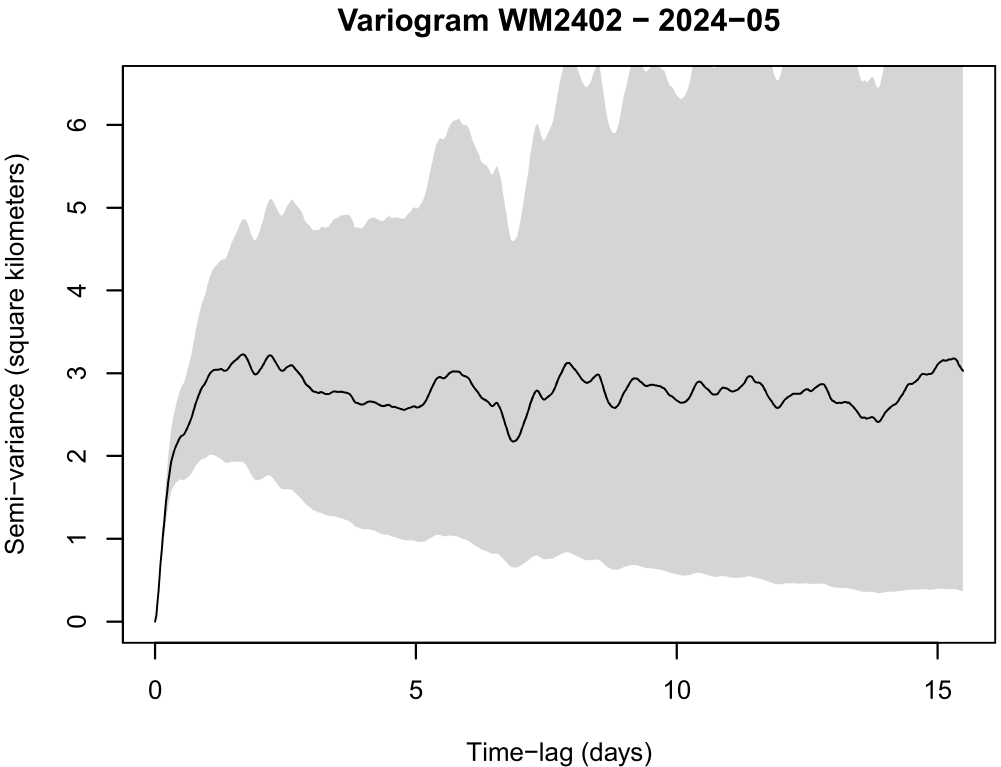
Variogram WM2402 − 2024−05
Semi−variance (square kilometers)
Time−lag (days)
0
1
2
3
4
5
6
10
15

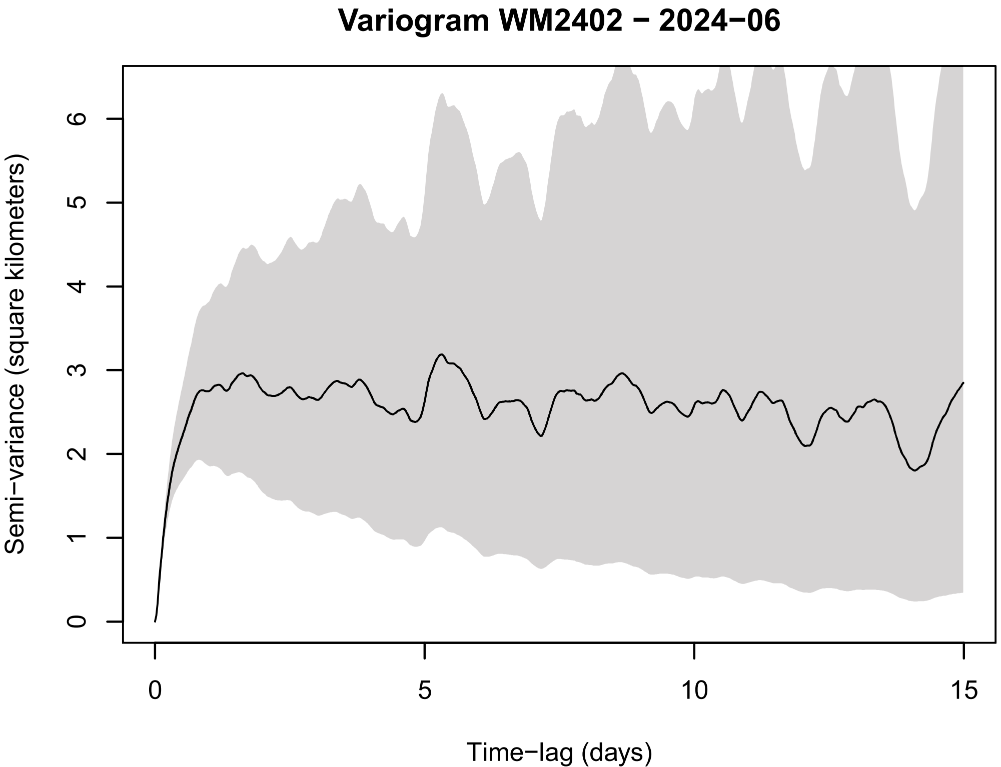
Variogram WM2402 − 2024−06
Semi−variance (square kilometers)
Time−lag (days)
0
1
2
3
4
5
6
10
15

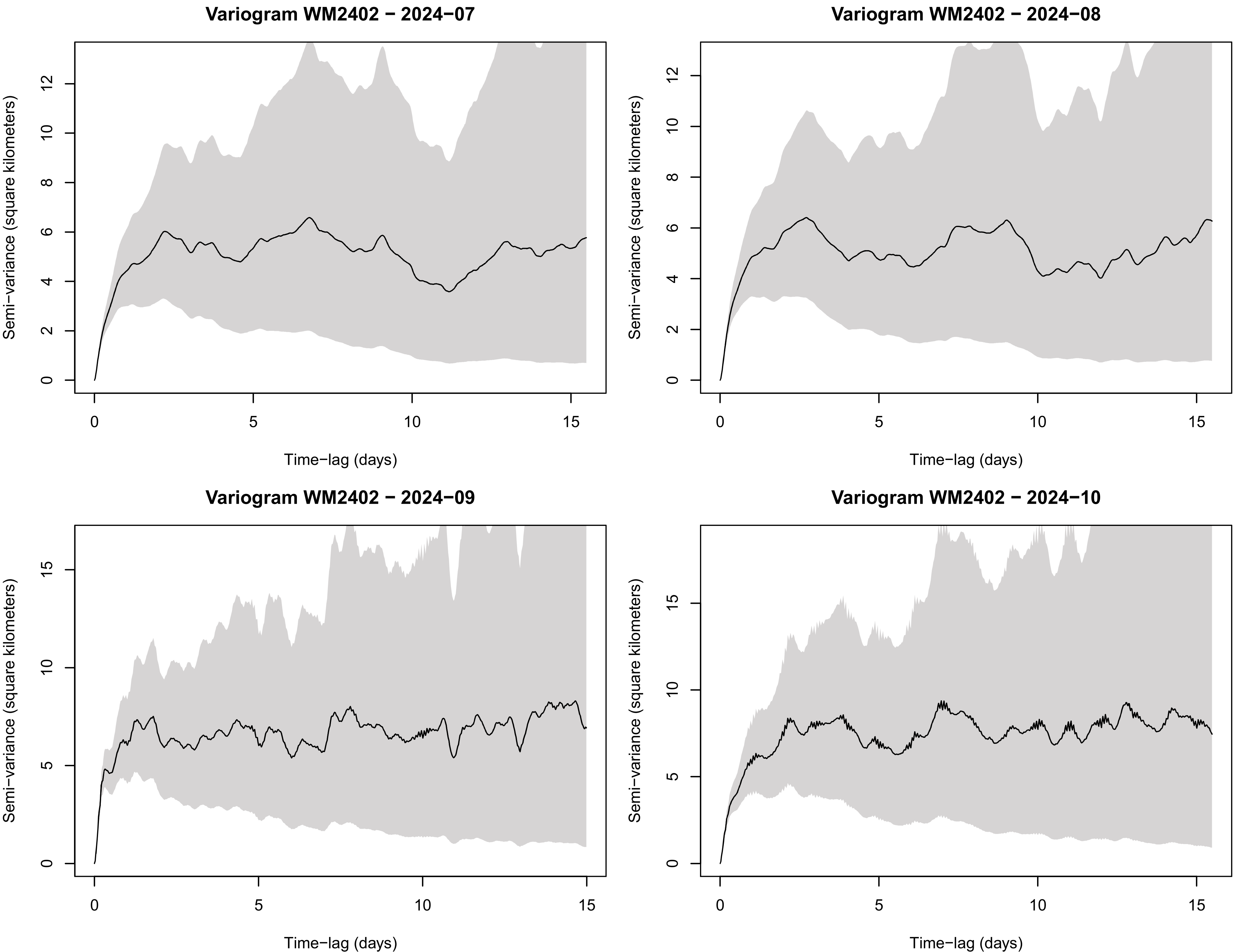
Variogram WM2402 − 2024−07
Variogram WM2402 − 2024−08
Variogram WM2402 − 2024−09
Variogram WM2402 − 2024−10
Semi−variance (square kilometers)
Time−lag (days)
0
5
10
15
2
4
6
8
12

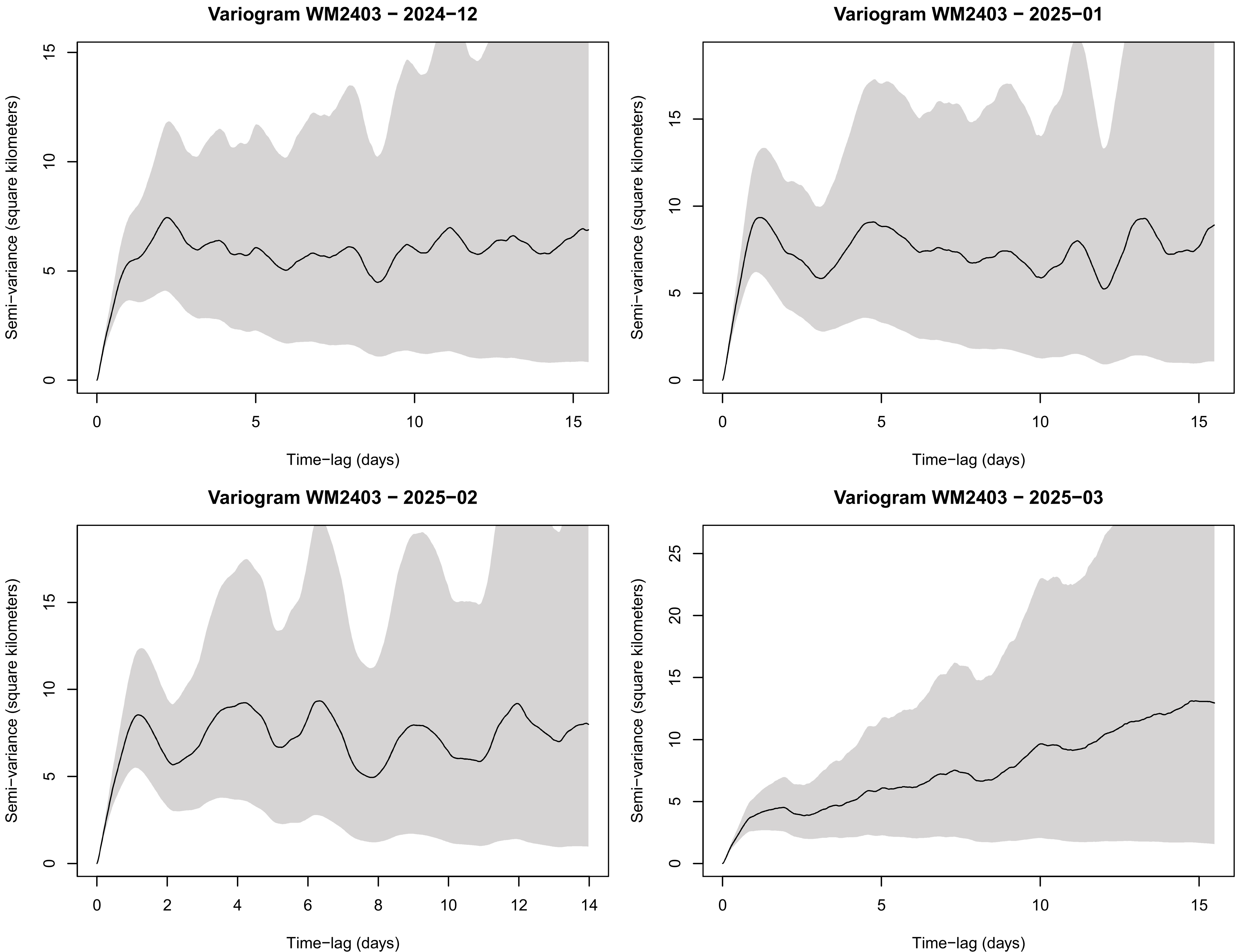

Variogram WM2403 − 2024−12
Variogram WM2403 − 2025−01
Variogram WM2403 − 2025−02
Variogram WM2403 − 2025−03
Semi−variance (square kilometers)
Time−lag (days)
0
5
10
15
20
25
0
2
4
6
8
10
12
14

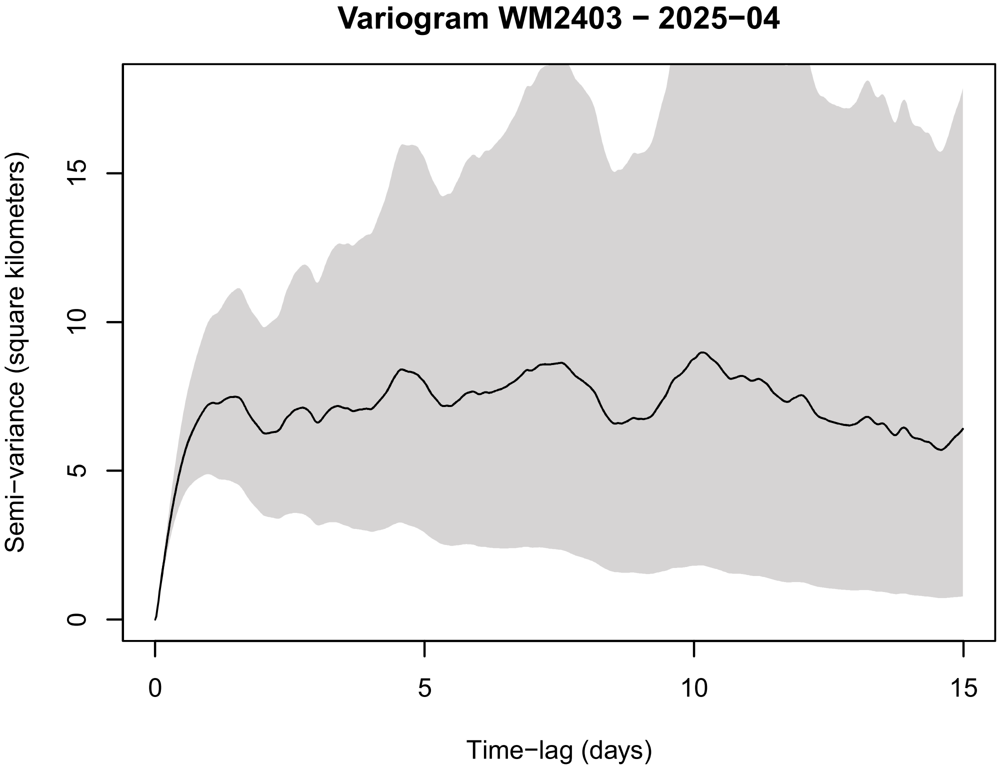
Variogram WM2403 − 2025−04
Semi−variance (square kilometers)
0
5
10
15
Time−lag (days)
0
5
10
15

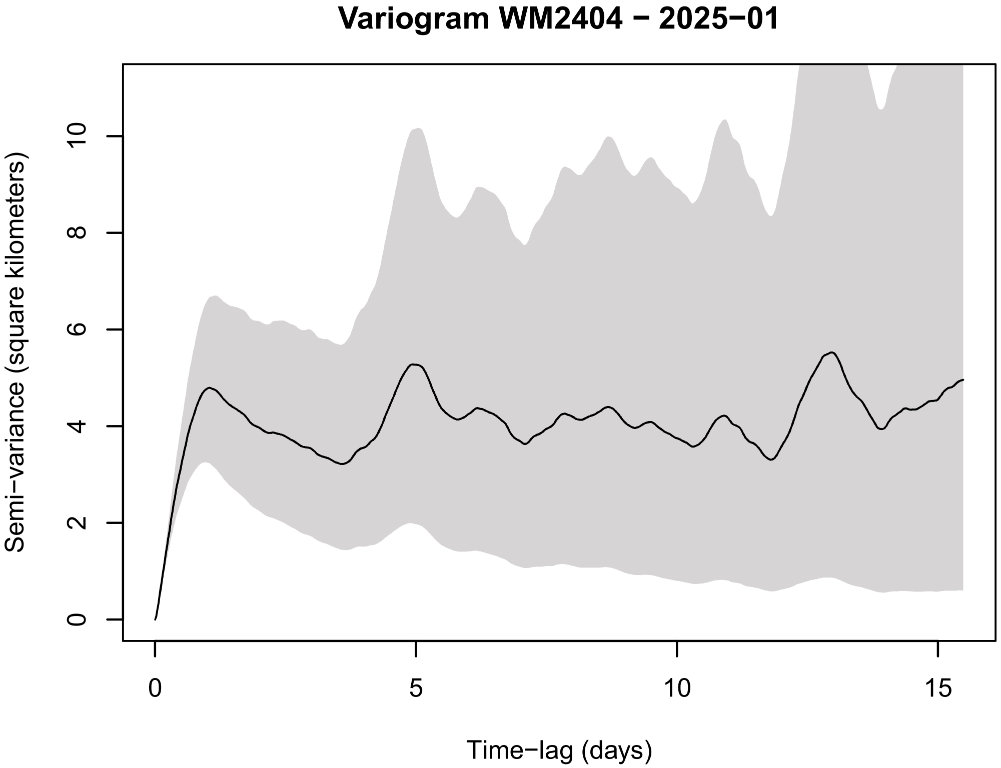
Variogram WM2404 − 2025−01
Semi−variance (square kilometers)
0
2
4
6
8
10
Time−lag (days)
0
5
10
15

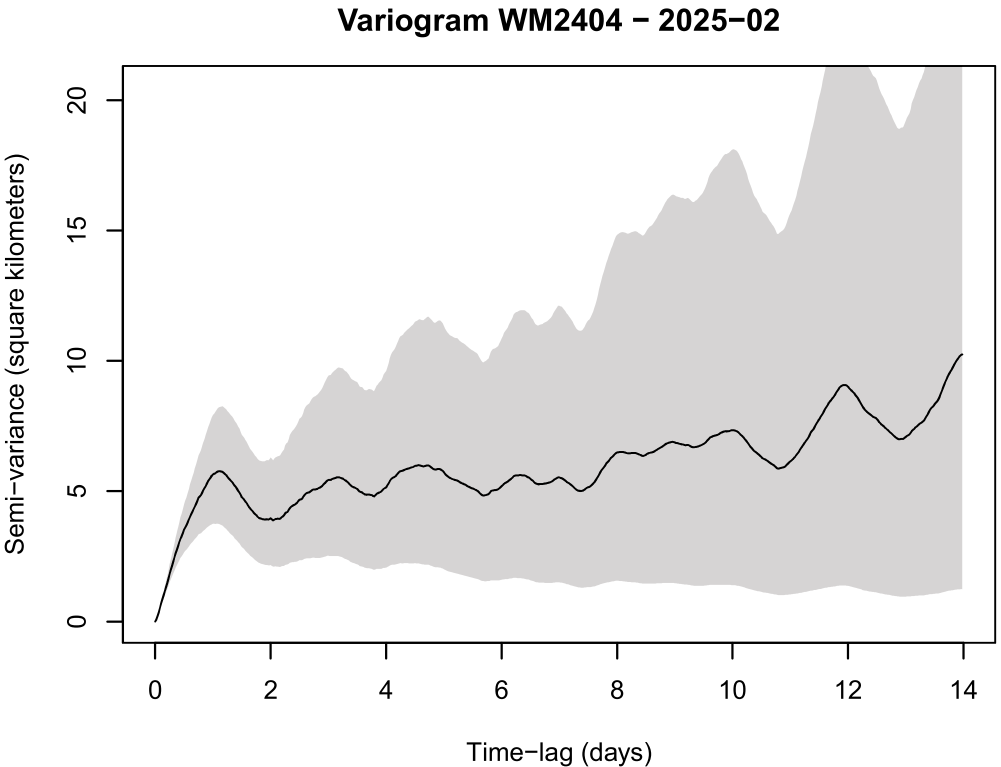
Variogram WM2404 − 2025−02
Semi−variance (square kilometers)
0
5
10
15
20
Time−lag (days)
0
2
4
6
8
10
12
14

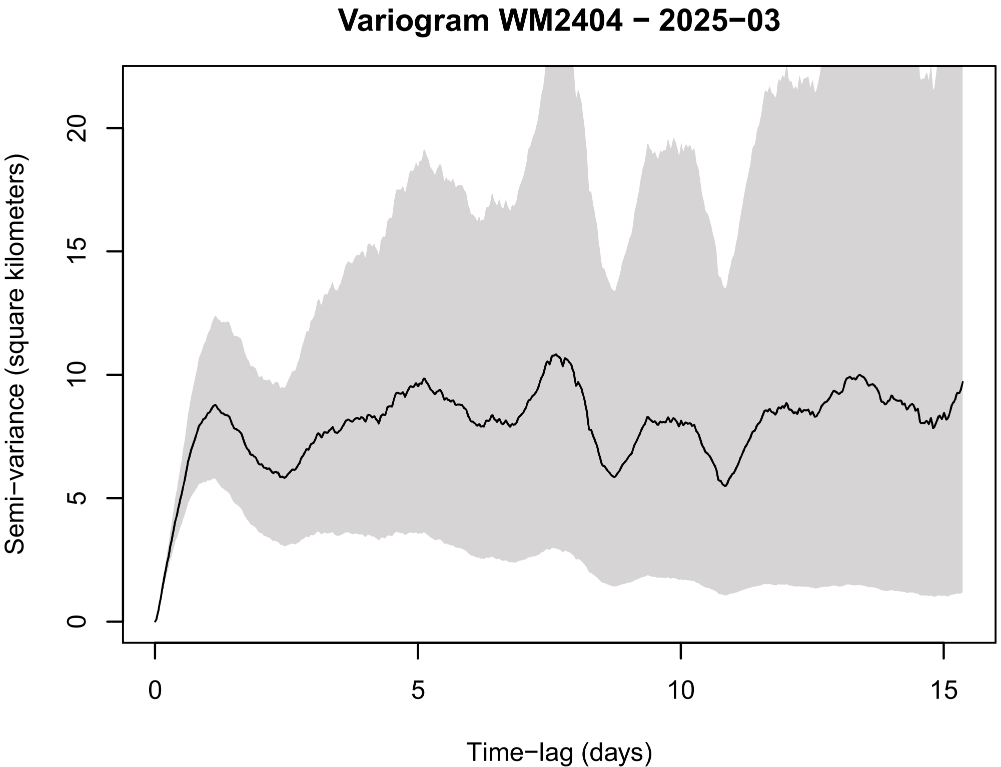
Variogram WM2404 − 2025−03
Semi−variance (square kilometers)
0
5
10
15
20
Time−lag (days)
0
5
10
15

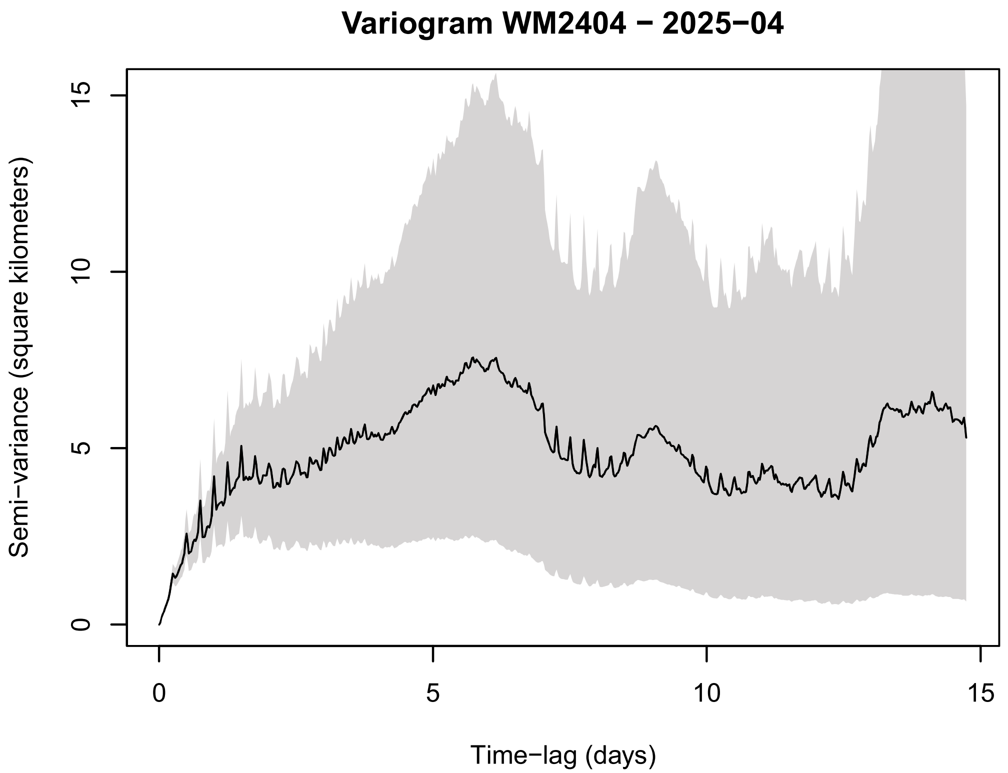
Variogram WM2404 − 2025−04
Semi−variance (square kilometers)
0
5
10
15
Time−lag (days)
0
5
10
15
